# LIFERING, an E3 ligase, ubiquitinates the acyl-CoA dehydrogenase-like protein IBR3, and coordinates cell proliferation and death upon DNA damage

**DOI:** 10.1101/2024.10.18.619078

**Authors:** Brigitta M. Kállai, Edit Németh, Aladár Pettkó-Szandtner, Zaki Ahmad, Fruzsina Nagy, László Bögre, Zoltán Magyar, Tamás Mészáros, Beatrix M. Horváth

## Abstract

Plants rapidly respond to environmental changes to ensure an optimal balance between growth and survival with intact genome. Here, we show that a DNA damage response gene, which we named *LIFERING*, is under the direct and independent regulation of ATM/ATR-SOG1 and RBR pathways. We demonstrate that LIFERING upon DNA damage plays an essential role in maintaining the balance between cell proliferation and cell death. Downstream of RBR, it is required to maintain proliferation, and in response to DNA damaging agents it initiates a cell death accompanied by rapid elongation and differentiation of the transit amplifying cells in the root meristem. LIFERING is a RING-between-RING E3 ligase; its ligase activity is dependent on the Cys392 residue located in the BRcat domain, responsible for the coordination of zinc ion binding. Using proximity labelling, we identified the acyl-CoA dehydrogenase-like protein IBR3, which is involved in the conversion of indole-3-butyric acid (IBA) to indole-3-acetic acid (IAA), as one of its targets. We provide evidence that IBR3 is ubiquitinated at several Lys residues in the presence of LIFERING. Our findings show that LIFERING rapidly responds to DNA damage and via ubiquitination of IBR3, is likely to be involved in the regulation of free-auxin level, indicating a link between DNA damage response and auxin regulation in the root.

## INTRODUCTION

During their entire lifecycle, plants are exposed both to exogenous and endogenous stresses and toxic compounds. Although sunlight is required for photosynthesis, the short-wave electromagnetic radiation causes cellular and DNA damage. Furthermore, other abiotic stresses, such as drought, salinity, and heavy metal contaminants also induce similar genotoxic effects (Yoshiyama *et al*., 2013a; Pedroza-Garcia *et al*., 2022). In addition, endogenous damage, spontaneously generated during DNA metabolism, replication and repair can also have an impact on the genome integrity. (Szurman-Zubrzycka *et al*., 2023).

Protection of genome integrity is especially crucial in the stem cell niches. The plant RETINOBLASTOMA-RELATED (RBR) plays a central role in this process by two distinct mechanisms: (i) promoting the exit from cell proliferation to differentiation (Harashima & Sugimoto, 2016) (Cruz-Ramirez *et al*., 2013), and (ii) directly regulating the expression of DNA damage response genes (Horvath *et al*., 2017). RBR performs these functions in complex with the three canonical E2Fs (E2FA, E2FB, and E2FC) and other core components of the DREAM complex (dimerization partner (DP), retinoblastoma (RB)-like, E2F and MuvB), which are essential to establish transient and stable cellular quiescence and DNA damage response (Lang *et al*., 2021; Gombos *et al*., 2023). In addition to the transcriptional control of genes involved in S-phase progression, replication and cell-cycle checkpoint, RBR, primarily in conjunction with E2FA, directly regulates the expression of numerous damage response and repair genes, including *AtBRCA1* and *AtRAD51* (Biedermann *et al*., 2017; Horvath *et al*., 2017). This allows RBR to act rapidly to endogenous replication-induced DNA damage and initiates an ATM/ATR-SOG1 independent response pathway.

To mitigate the consequences of DNA damage, plants similarly to other organisms have evolved mechanisms to sense DNA lesions and initiate the DNA damage response (DDR) pathways (LaRocque & McVey, 2023). The DDR signalling cascade is conserved in animals and plants (Clay & Fox, 2021; Herbst *et al*., 2024); the double-strand breaks (DSB) activate the ATAXIA-TELANGIECTASIA MUTATED (ATM) kinase by the MRN complex (MRE11-RAD50-NBS1), while the ATAXIA-TELANGIECTASIA-AND-RAD3-RELATED (ATR) is mainly recruited to single stranded breaks (SSB) by the complex of Replication Protein (RPA) and ATR Interacting Protein (ATRIP). Though both ATM and ATR have distinct functions during signalling in response to genotoxic agents such as γ-irradiation, they show additive and complementary roles during DNA replication, cell cycle progression and repair (Nisa *et al*., 2019; Herbst *et al*., 2024).

In plants, the activity of the ATM/ATR-driven signalling pathway merges with the phosphorylation of the plant-specific transcription factor, SUPPRESSOR OF GAMMA RESPONSE 1 (SOG1), which is considered to be the functional analogue of the mammalian p53 (Yoshiyama *et al*., 2013a; Yoshiyama *et al*., 2013b). Upon genotoxic stress, in parallel to RBR and E2Fs, the activated SOG1 regulates the expression of numerous DNA repair and cell cycle genes to coordinate repair with cell cycle progression (Bourbousse *et al*., 2018; Ogita *et al*., 2018).

Halted cell cycle resumes when the lesions are repaired via various pathways depending on the nature of the lesion (Raina *et al*., 2021; Szurman-Zubrzycka *et al*., 2023). However, in the case of severe damage, proliferating cells escape from further division and initiate processes of (i) cell death, especially in the stem and daughter cells (Furukawa *et al*., 2010; Horvath *et al*., 2017), (ii) endoreduplication (Adachi *et al*., 2011; Lang & Schnittger, 2020), and/or (iii) differentiation. RBR plays a key role to establish the threshold at which cell death is initiated upon genotoxic stress (Horvath *et al*., 2017).

Besides transcriptional regulation, posttranslational modifications, such as phosphorylation, ubiquitination and SUMOylation, also play an essential role in the regulation of the DNA damage response (Su *et al*., 2020). Among the characterised plant-specific E3 ubiquitin ligases involved in DDR, the DNA damage response mutant 1 (DDRM1) and 2 (DDRM2) play a role in the early steps of damage recognition and recruitment of repair proteins to the damaged site. DDRM1 monoubiquitinates the phosphorylated SOG1, thereby promoting its stability (Wang *et al*., 2022). Subsequently, SOG1 induces the expression of DDRM2, binding of which to AtRAD51 is required for its recruitment to the damaged DNA site labelled with γ-H2AX (Yu *et al*., 2023). Upon DDR, the RBR-induced *KNOTEN1* (KNO1) also partially co-localises with γ-H2AX and is stabilised by the de-ubiquitinases, UBP12 and UBP13 (Chen *et al*., 2023). After stabilisation, KNO1 mediates the K63-linked ubiquitination and autophagic degradation of RMI1 protein, a structural component of the Bloom syndrome complex, thereby increasing homologous recombination frequency (Chen *et al*., 2023).

Studying the role of RBR in the protection of genome integrity, we found that the *At5g60250* transcript exhibited significantly higher expression levels than any other transcripts upon silencing of *RBR* (Horvath *et al*., 2017). With genome-wide transcriptional analyses, *At5g60250* was also detected to be upregulated in response to a plethora of genotoxic conditions, including γ-radiation and oxidative stress (Culligan *et al*., 2006; Yi *et al*., 2014), radiomimetic drugs like hydroxyurea and bleomycin (Cools *et al*., 2010; Yi *et al*., 2014), metal toxicity (Sjogren *et al*., 2015; Chen *et al*., 2019), as well as due to mutations in genes, such as *fas-1* and *fas-2* (Schonrock *et al*., 2006), and upon overexpression of AAR7 (Lee *et al*., 2007). Despite extensive knowledge on the expression of *At5g60250* in various conditions, the role in DDR and its catalytic activity remains unknown.

Here we provide evidence that *At5g60250* hereafter named as LIFERING, is directly regulated by RBR in an independent pathway to ATM/ATR-SOG1. Our genetic analysis shows that upon genotoxic stress LIFERING is required to maintain meristematic activity by preventing the exit to differentiation as well as to regulate stem cell divisions downstream of RBR. We demonstrate that LIFERING is a functional ubiquitin E3 ligase; it is capable of autoubiquitination, and its activity depends on the Cys392 residue involved in the coordination of zinc ion binding. Using proximity-based labelling and protein interaction studies, we show that the most prominent interacting protein of LIFERING, the acyl-CoA dehydrogenase-like protein, IBR3 is involved in the conversion of indole-3-butyric acid (IBA) to indole-3-acetic acid (IAA). Both by *in vitro* and *in vivo* assays, we provide evidence that LIFERING ubiquitinates IBR3.

Based on these data, we suggest that the ubiquitination of IBR3 via LIFERING may connect the DDR and auxin responsive pathways upon DNA damage, to protect cells via auxin-driven cell death response.

## MATERIALS AND METHODS

### Plant material and growth conditions

Seeds were sterilised and grown as described earlier (Blilou *et al*., 2005) except that seedlings used for qRT-PCR and microscopical analysis were germinated on 1.2 % plant agar. *Arabidopsis thaliana* ecotype Columbia 0 (Col-0) was used as wild type; the T-DNA insertion line, *lifering-1* (SM_3_34488) was obtained from the Nottingham *Arabidopsis* Stock Centre. The transgenic lines, *e2fa-1* (MPIZ_244, (Berckmans *et al*., 2011) *sog1-1*, (Yoshiyama *et al*., 2009) *Atbrca1-1* (Reidt et al., 2006)*, rRBr* (Blilou *et al*., 2005), *amiRBR* (Cruz-Ramirez *et al*., 2012) and *amiRBR*,*sog1*-*1* (Horvath *et al*., 2017) were described earlier. SM_3_34488, is an JIC SM line, ordered via NASC. The T-DNA insertions, mutations were confirmed by PCR based genotyping or sequencing and gene silencing was demonstrated via gene expressional studies and phenotyping. To study the *lifering-1* (pgLIFERING-GFP) phenotype more than ten independent transformant were generated. The T-DNA insertion mutations were confirmed by PCR based genotyping or sequencing while gene silencing was demonstrated via gene expressional studies and phenotyping. *GFP* expression was followed by microscopical analysis. The list of primer pairs used in these experiments are listed in Supporting Information Table S1.

### Chemical treatments and induction studies

To induce DNA damage response, 5-6 days-old seedlings (das) were transferred to tissue culture plates, containing fresh MS liquid medium without or with 10 μg ml^-1^ mitomycin C (MMC), and treated for a short period of 1-4 h or 16 h. Kinase inhibitory assays were carried out as described earlier (Horvath *et al*., 2017). In each treatment, the relevant controls and mutants were studied simultaneously, the experiments were repeated at least three times (n=biological repeat) with 15-20 (*N*=sample size) replicates. Although the level of MMC induction varied between the different biological experiments, the ratio of the relevant induction between the treated controls and samples in interest were comparable. Counting the number of propidium-iodide (PI)-stained cells in the columella stem cells (CSC) and lateral root cap initials (LRC) and their daughter cells was used to quantify cell death in the distal stem cell niche. In addition, cell death was also determined of the root-tip measuring the contiguously PI-stained cell area in the proximal meristematic vasculature directly adjacent to the QC.

### Fluorescence microscopy and data processing

Leica SP2 or Olympus IX81-FV1000 inverted laser-scanning microscope equipped with different laser lines were used for phenotypic analysis of roots stained in 5 µg/ml propidium-iodide (PI), at 543nm for PI and at 488nm for GFP. For qualitative and quantitative comparison, images were taken in a single median section and recorded with identical microscope settings in all cases. Confocal images were converted to tiff files and processed with ImageJ software. Cell length was measured on the longitudinal axis in the middle of the cell, in the epidermis and/or cortex cell layers. The zone of the transit amplifying cells was defined by the boundary of cells with gradually increasing length. The distance from the QC to the first differentiating cell determined the start of differentiation.

### Cloning

The coding sequence of *LIFERING* (accession number AT5G60250) was amplified from the RIKEN RAFL22-04-P06 clone, then inserted into pEU3-NII-HxHLICNot and pRT-HA-TurboID-LIC using LIC (Bardoczy *et al*., 2008) and using *Nco*I and *Not*I restriction endonucleases into pRT-3xHA. R392 residue of the mutant LIFERING ORF (designated as C392R) was restored to C392 (designated as wild-type, WT) by *in vitro* mutagenesis (Laible & Boonrod, 2009). The coding sequence of IBR3 (accession number AT3G06810) was amplified from pACT2CS cDNA library (Nemeth *et al*., 1998) and inserted into pEU3-NII-GLICNot and pRT-9xMyc vectors as described above. Details on the vectors and primers used for cloning are provided in Supporting Information Table S2. Coding sequences of the constructs were verified by Sanger sequencing (Eurofins Genomics)

### *In vitro* protein expression and ubiquitination

The pEU3 vector constructs were used for *in vitro* transcription (Nagy *et al*., 2020) and 2.5 μl of crude mRNA-containing reaction mixture was added to 10 μl of W7240 wheat germ extract (CellFree Sciences). The bilayer *in vitro* translation was prepared according to the manufacturer’s recommendation, except that both layers were supplemented to contain 10 μM ZnCl_2_ and incubation was done for 20 hours at 20°C.

The *in vitro* ubiquitination reactions were performed as described previously (Takahashi *et al*., 2009) with minor modifications. The WT and C392R 12xHis-LIFERING-containing translation mixtures were diluted 1.25 times in the final volume of the ubiquitination reactions to contain 10 mM Tris-HCl pH 7.5, 5 mM MgCl_2_ and 3 mM ATP. FLAG-ubiquitin was produced in *E. coli* cells as described in the Supporting Information and was added at 4 μM final concentration where indicated. The mixtures were incubated for 3–4 hours at 30°C either alone to test for the ubiquitination activity of LIFERING or were added to bead-bound GST-IBR3 to ubiquitinate it.

12xHis-LIFERING was purified using Dynabeads™ His-Tag Isolation and Pulldown Beads (Thermo Scientific). 230 µl ubiquitination reaction mixture was added to 15 µl beads equilibrated and resuspended in 230 µl 2X B/W buffer (100 mM sodium phosphate pH 8, 600 mM NaCl, 0.02% Tween-20), and incubated on a rotator for 1 h at room temperature. The beads were washed three times with 300 µl 1X B/W buffer for 5 minutes each, twice with 1.5 ml 1X B/W buffer and six times with 1.5 ml PBS.

GST-IBR3 was purified using Pierce Glutathione Beads (Thermo Scientific) prior to the ubiquitination reactions. 200 µl GST-IBR3-containing translation mixture was added to 30 µl beads equilibrated and resuspended in 1 ml Glut B/W buffer (125 mM Tris-HCl pH 8, 150 mM NaCl, 0.5% Triton-X), and incubated on a rotator for 1 hour at room temperature. Beads were washed five times with 300 µl Glut B/W buffer and then resuspended in 100 µl ubiquitination reaction mixtures containing 12xHis-LIFERING proteins. After incubation, the unbound fraction was retained and the bead-bound GST-IBR3 proteins were washed three times with 300 µl Glut B/W buffer for 5 minutes each, twice with 1.5 ml Glut B/W buffer and six times with 1.5 ml PBS. The reactions were analysed by Western blot and the bead-bound proteins were subjected to mass spectrometry as described in the Supporting Information.

### Transient protoplast protein expression

C-terminally 3xHA-tagged wild-type and mutant C392R LIFERING and 9xMyc-tagged IBR3 were expressed driven by the *35S* promoter of pRT vectors through PEG-mediated transfection of 2×10^5^-5×10^5^ protoplasts (Yoo *et al*., 2007) isolated from Columbia (Col-0) *Arabidopsis thaliana* root cell suspension (Mathur & Koncz, 1998). Transfections and co-transfections were performed by using 5 μg of each vector. After 16 hours of incubation in the dark at room temperature, the cells were lysed by RIPA buffer and boiled in SDS-DTT sample buffer. The extracts were centrifuged at 13000 rpm for 10 minutes and the supernatants were analysed using Western blot. To study the stability of IBR3, 2 μg of each vector was used for the transfections, and after 16 hours incubation in the dark at room temperature, cycloheximide (CHX) and MG132 were added to a final concentration of 50 μM and further incubated in the dark for the indicated times, followed by lysis as previously described.

The details of the proximity labelling experiment can be found in the Supporting Information.

### SDS-PAGE and Western blot

For the SDS-PAGE analysis, the samples were mixed with SDS-DTT sample buffer and heated at 95°C for 5 minutes. Proteins were separated on SDS-PAGE or AnyKD (Bio-Rad) gels with Laemmli running buffer, then transferred to Immobilon®-FL low fluorescence PVDF membrane (Merck Millipore). The Dunn carbonate buffer system (Dunn, 1986) was used for protein transfer at 30 V for 16 hours at 4 °C. The Odyssey CLx Imager (LI-COR Biosciences) was used with automatic intensity settings for two-colours detection of FLAG-tagged and His-tagged proteins, Single detection of GST-, 3xHA-, and 9xMyc-tagged proteins and Revert 700 Total Protein Stain were carried out by following the manufacturer’s detection protocols. The list of antibodies used are provided in Supporting Information Table S3.

## RESULTS

### The DNA damage response (DDR) gene, *LIFERING,* is regulated both by the RETINOBLASTOMA-RELATED and the ATM/ATR-SOG1 pathways

The elevated expression of *LIFERING* upon RBR silencing was shown by micro-array analysis (Horvath *et al*., 2017). To corroborate these data, we followed its expression by qRT-PCR both in *rRBr* root-tips where *RBR* is silenced in the root meristem (Fig. 1a), and in *amiRBR* seedlings where the post-embryonic RBR levels are reduced constitutively using the *35S* promoter (Fig. 1b). The results obtained were in accordance with the microarray data, indicating that the transcriptional level of *LIFERING* was significantly higher in both *rRBr* and *amiRBR* lines. As *LIFERING* is co-expressed with *AtBRCA1* (Horvath *et al*., 2017), a gene known to be an essential component of DDR and homologous recombination, we compared the expressional changes of *LIFERING* and *AtBRCA1* and found a significantly higher level of induction of LIFERING than of *AtBRCA1* (Fig. 1a and b).

**Figure 1.**
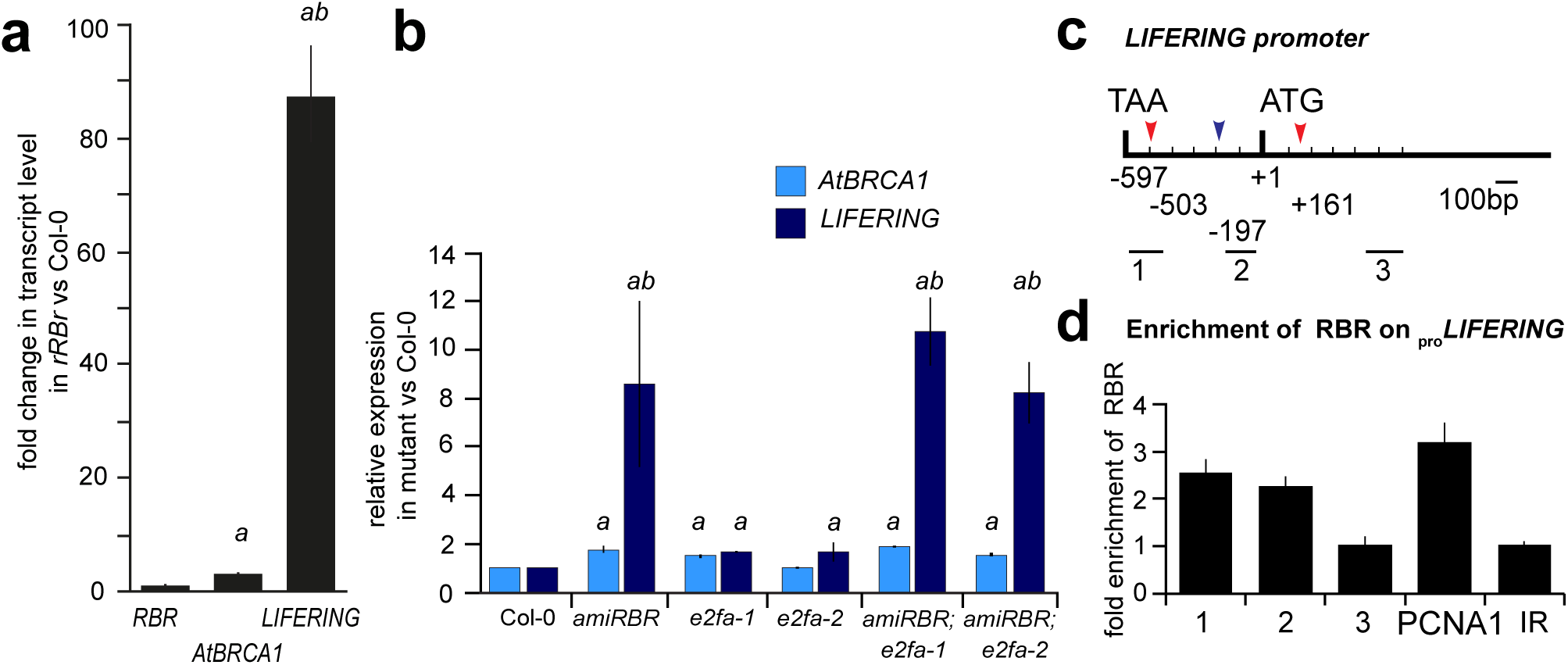
The expression of *LIFERING* is regulated by RBR through E2FA. **(a)** Bar chart shows the gene expression of *LIFERING* (At5g60250) compared to *RBR* and *AtBRCA1* expression determined by qRT-PCR. RNA was isolated from root-tips of 4-day old *rBRr* and Col-0 seedlings. Gene expression is given as the ratio between *rRBr* and Col-0. Fold change is given as *AtBRCA1* gene expression = 1. **(b)** Relative transcript level of *LIFERING* and *AtBRCA1* as a control, upon *RBR* silencing in *amiRBR* and in the single: *e2fa-1, e2fa-2* and double mutants compared to Col-0, where the level of expression was set to 1 for both genes. **(c)** Schematic representation of the *LIFERING* promoter. The positions of the following elements are indicated: start codon (ATG; +1), stop codon of the neighboring transcript (TAA; −597), putative E2F elements (red arrow-heads; −503:taggcccgcgaattt and +161:cggcgaaaa) and SOG1 binding site (black arrow; −197^-^-169) as suggested by Bourbousse *et al* (2018). Black horizontal lines (1, 2, 3) indicate the position and length of the amplified regions in the qPCR analysis, **(d)** Chromatin-Immuno Precipitation (ChIP) using RBR antibody; the chart shows fold enrichment calculated as a ratio of chromatin bound to the numbered sections of the promoter in (**c**) with or without antibody. The *PCNA1* promoter was used as a positive control, while IR (an intergenic region between At3g03360-70) as a negative control. Note that the enrichment of RBR on the *LIFERING* and *PCNA1* promoters are comparable. Data information: In (**a-c**): *a*: p<0.05 significance of the *AtBRCA1* expression in the mutant compared to Col-0 and *ab*: p<0.001 significance of the *LIFERING* expression in the mutant compared to Col-0 and to *AtBRCA1* calculated by the Student’s *t*-test. Error bar: +/− StDev, *n*=3, *N*>100.

To address whether RBR regulates *LIFERING* gene expression in conjunction with E2FA, we quantified its expression and found that it was elevated in both *e2fa-1* and *e2fa-2* mutants, but at a lower level than in *amiRBR* (Fig. 1b). The de-repression of *LIFERING* in the homozygous double mutant lines, *amiRBR;e2fa-1* and *amiRBR;e2fa-2* did not exceed the de-repression detected in *amiRBR* (Fig. 1b). These results suggest that RBR acts partly through E2FA in a co-repressor complex to regulate *LIFERING* expression.

The presence of the canonical E2F binding sites within a 1kb region from the translational start site (Fig. 1c, ATG) further suggested that RBR may regulate *LIFERING* expression through E2F proteins. To confirm whether RBR indeed directly regulates *LIFERING* expression, chromatin immunoprecipitation (ChIP)-PCR assays were carried out on root tissues using an RBR-antibody (Horvath *et al*., 2006). Significant enrichment of RBR was detected in distinct regions of the *LIFERING* promoter. The enrichment on fragment 1 (Fig. 1c) was comparable to that detected on the *PCNA1* promoter, (Fig. 1d). A slightly lower level of enrichment was also observed in the neighbouring region (fragment 2), likely to be due to the heterogeneous size of sonicated fragments in the range of 300 to 500 bp.

To investigate the regulation of *LIFERING* expression upon DNA damage, *Arabidopsis thaliana* Col-0 seedlings were treated with the DNA damage agent, mitomycin C (MMC) and the induction of *LIFERING* transcription was analysed using qRT-PCR. Upon MMC treatment, *LIFERING* expression was significantly higher than that of *AtBRCA1* (16h, 10μg ml^-1^, Fig. 2a), consistent with the difference observed in mRNA levels of the same genes upon RBR silencing. To assess whether *LIFERING* induction depends on the activity of the ATM/ATR kinases, we inhibited the ATM/ATR driven pathways with specific chemicals developed to block the activity of these kinases (IATM, KU55933 and IATR, VE-821) and previously shown to be effective in plants (Horvath *et al*., 2017). While *AtBRCA1* expression was equally reduced by both the ATM and ATR inhibitors, *LIFERING* expression was more sensitive to the inhibition of ATR. The simultaneous inhibition of the ATM and ATR kinases almost completely abolished the expression of both *LIFERING* and *AtBRCA1* (Fig. 2b).

**Figure 2.**
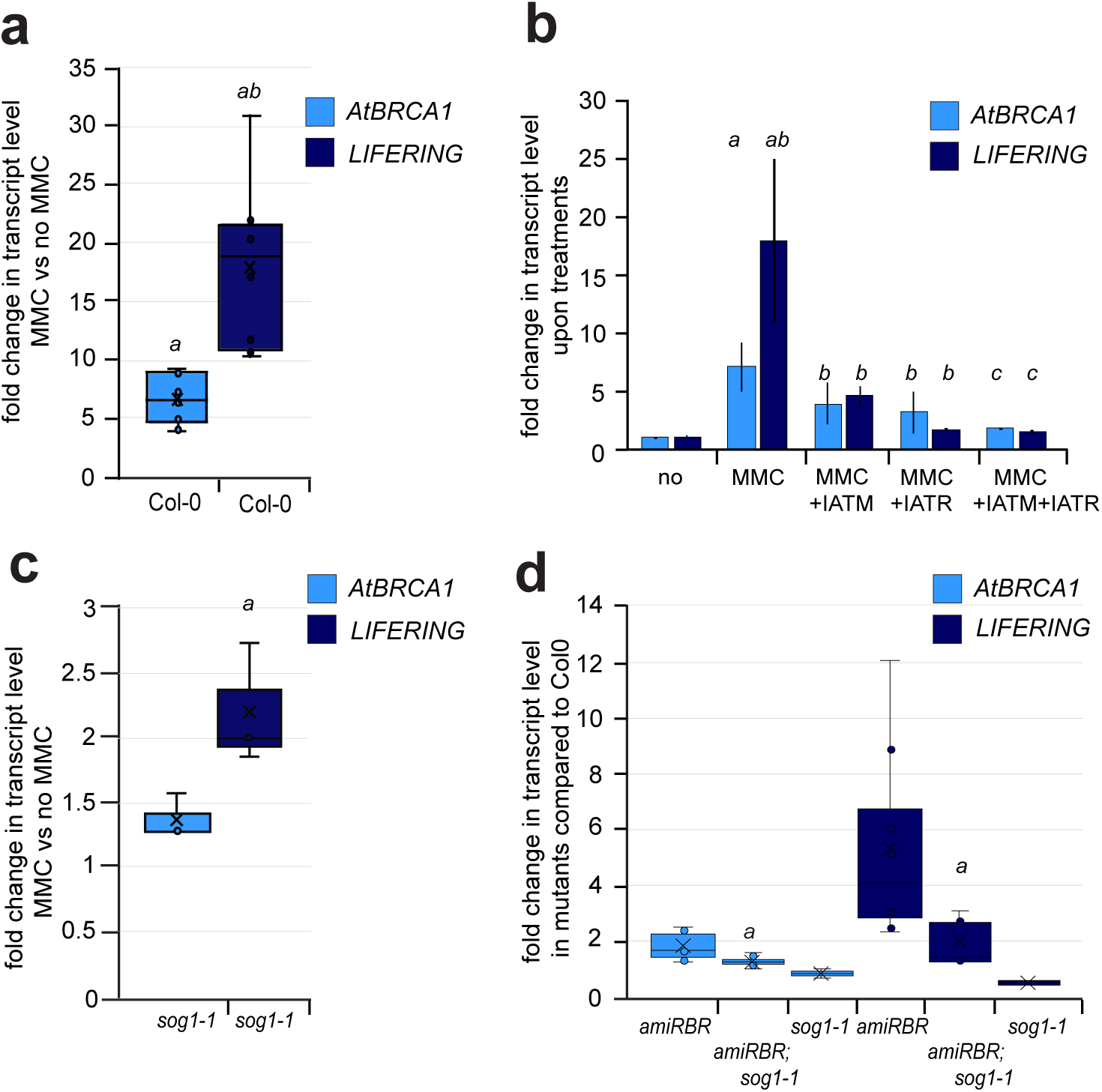
*LIFERING* expression is independently regulated by ATM/ATR-SOG1 and RBR pathways. (**a**) Chart of relative gene expression of *LIFERING* upon MMC (10mg ml^-1^) treatment for 16h.The qRT-PCR was carried out on Col-0 seedlings. Relative expression of *AtBRCA1* is shown as a comparison. The values represent fold changes normalised to the values of the relevant genes in non-treated samples. *a*: p<0.01 significance compared to the non-treated control, *ab:*p<0.01 significance *LIFERING* to the non-treated control and of *AtBRCA1* transcript levels.. (b) Effect of the ATM and ATR inhibitors (IATM and IATR, respectively) on the induction of *LIFERING* expression upon DNA damage (16h, 10mg ml^-1^ MMC). Relative transcript levels of *LIFERING* and *AtBRCA1* as a control, were calculated comparing the expression levels to the non-treated sample which was set to 1. *a*: p<0.05 significance between MMC and no-MMC treated samples, *b*: p<0.05 significant difference upon applying inhibitors individually and *c*: p<0.05 upon applying both inhibitors. (c) Chart of the relative expression of *LIFERING* in the *sog1-1* mutant upon MMC and non-MMC treatment (16h, 10mg ml^-1^) in comparison to *AtBRCA1* expression. The values represent fold changes normalised to the values of the relevant gene in non-treated samples where the expression was set to 1. *a*: p<0.05 significant difference compared to the non-treated control. (d) Effect of *sog1-1* mutation in the *amiRBR* background (*amiRBR;sog1-1*) both on *LIFERING* and *AtBRCA1* expression. Relative transcript levels were calculated comparing the expression levels to the relevant transcripts in Col-0 seedlings, in which the values were set to 1. *a*: p<0.05 significant difference in the double mutant compared to both single mutants. Data information: In (**a**-**d**), experiments were carried out on 5-6 das seedlings, *n*>3, *N*>100. Significance was calculated by the Student’s *t*-test, error bars: +/− StDev.

As SOG1 regulates DDR gene expression upon DNA damage, we tested the expression of *LIFERING* in the *sog1-1* mutant after MMC treatment. Comparing the relative expression of *LIFERING* with and without genotoxic agent in the *sog1-1* mutant, we detected 1.5-2 times higher relative gene expression of *LIFERING* than *AtBRCA1* (16h, 10μg ml^-1^, Fig. 2c). This indicates that the SOG1-independent induction is stronger in the case of *LIFERING* than *AtBRCA1*.

Lastly, we tested whether the regulation of *LIFERING* expression upon *RBR* silencing is independent of SOG1 function. To this end, we followed its transcript level in the double *amiRBR,sog1-1* and the single mutants as controls. In the *sog1-1* mutant, the level of *LIFERING* expression was only partially reduced in the *amiRBR* background, suggesting that the SOG1 and RBR control *LFR* expression independently (Fig. 2d).

### Role of LIFERING in cell death vs cell differentiation upon DNA damage

To investigate the role of LIFERING in DDR, first, we studied the phenotypic changes of the *lifering-1* mutant (SM_3_34488) upon genotoxic stress. For this, we studied whether the mutation in the 2^nd^ exon leads to full knock-out of the relevant transcript by RT-PCR. Indeed, no induction of the *LIFERING* transcript was detected in the mutant either in the presence or absence of MMC in the mutant compared to the control (MMC, 16h, 10μg ml^-1^, Fig. S1a and b).

Under normal growth conditions, root development and growth of the *lifering-1* mutant was comparable to the wild-type (Fig. S1c and d). No difference was detected either in the structure of the root or in the organisation of the meristem (Fig. 3a and Fig. S1e). To be able to establish its role in the cellular processes upon DDR, first we followed the cell death response in the mutants, *lifering-1, Atbrca1-3* and Col-0 roots, in 5-6 days-old seedlings (das) upon MMC treatment (14-16h, 10μg ml^-1^). As shown in Fig. 3, the cell death response was significantly weaker in the *lifering-1* mutant, compared to Col-0 and *Atbrca1-3*, both in the proximal and distal root meristems (Fig. 3a-c). It is important to note that high proportion of *lifering-1* seedlings did not display any cell death response. To verify this response, we counted the number of the roots with zero, one or more than one dead cell in the stem cell niche. Only 15% of the studied seedlings developed cell death in more than one cell in the *lifering-1* SCN compared to the 61% of Col-0 (Table S4). To provide evidence that this phenotype is exclusively linked to the mutation in *LIFERING*, we complemented the mutant with the LIFERING-GFP construct, in which the genomic region of *LIFERING* was fused to GFP and expressed under its own promoter (pgLIFERING-GFP). In the three independent transformants, both the transcriptional and cell death responses were comparable to those of Col-0 upon MMC treatment (Fig. S2a-c). The fluorescence signal was detected only after inducing DNA damage, predominantly in the cytoplasm and was located along the entire meristem (Fig. S2d).

**Figure 3.**
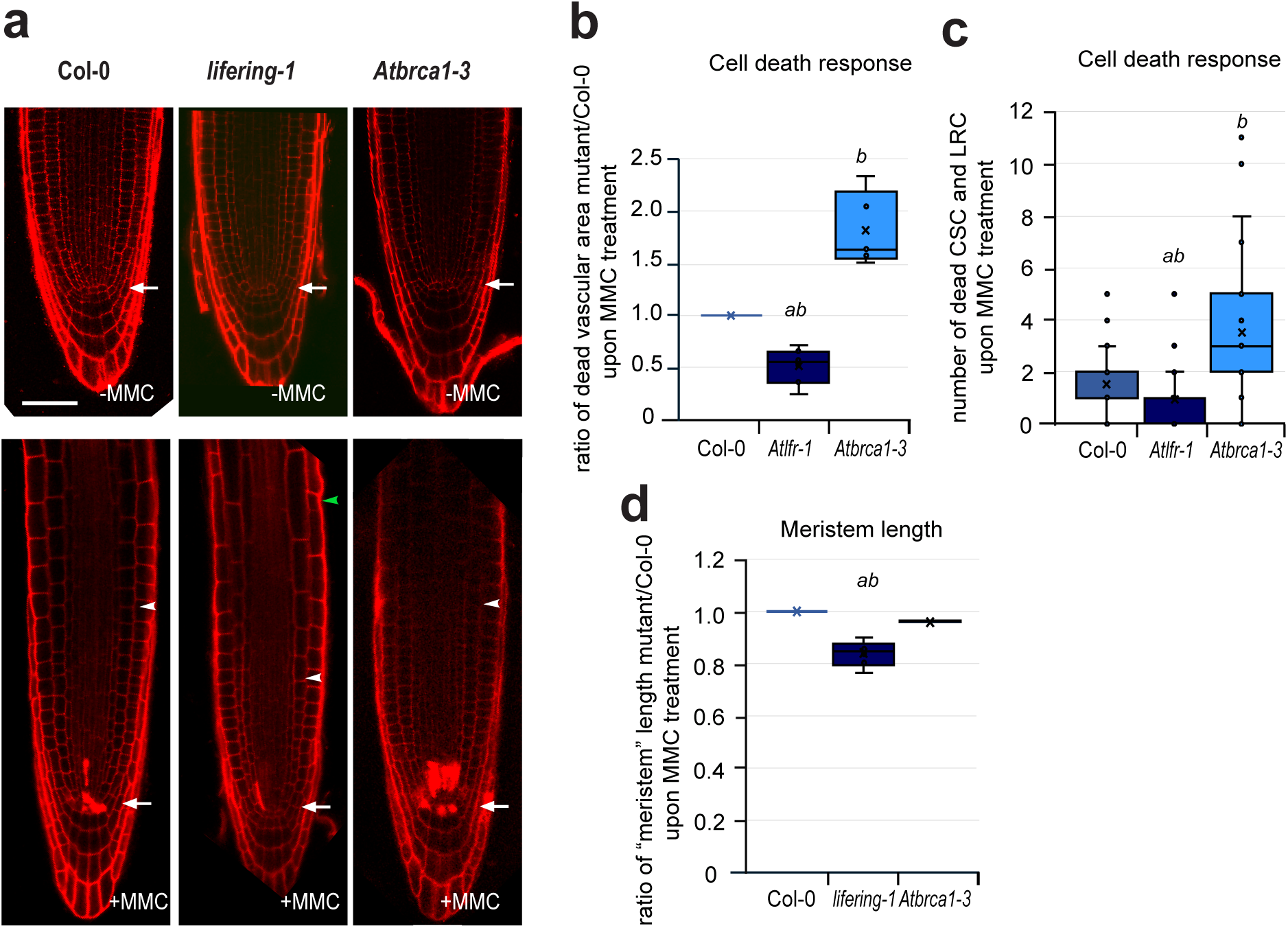
LIFERING is involved in DNA damage response upon genotoxic stress. (**a**) Representative confocal microscopy (CM) images of propidium-iodide (PI)-stained root-tips of Col-0, *lifering-1* and *Atbrca1-3* mutants; seedlings were grown without (-MMC, top line) and at 5-6 days, treated with MMC (+MMC, bottom line) for 16 h. Arrows indicate the position of the QC, white arrowheads in the cortical cell layer point towards the transition zone, while the green arrowhead shows the position of the first roothair forming epidermal cell in the MMC-treated *lifering-1* roots. Scale bar: 50μm. See Figure S1 for further comparison of Col0 and *lifering-1* root phenotype with and without MMC treatment. (**b**) Cell death response was quantified in the proximal meristem by measuring the area of dead cells (PI-stained) in Col-0 and in the mutants, *lifering-1*, *Atbrca1-3*. The plot shows the ratio of PI-stained area in the relevant mutant and Col-0. The PI-stained area was measured in each experiment (*n*>4) from *N*>15 mutants and Col-0. Although, the mean of cell death area varied between the different biological repeats, the ratio between the mutant and Col-0 (setting the area of the wild-type to 1), allowing comparisons between independent experiments. (**c**) Quantification of death response in the distal stem cell region by counting the dead columella stem and daughter cells (CSC) in addition to the lateral root initials and their descendants (LRC) in the median section as shown in (**a**). Data of *n*>5, *N*±90 seedlings. Further information about the distribution of the stem cell niche with 0, 1 and n>1 collapsed cells is given in Table S4. (**d**) The ratio of the length of the meristematic region between the mutants and Col-0 calculated from *n*>4, *N*>10 in each experiments. Length of meristem is defined here as distance between the QC and the cortical cell boundary at the transition. Data Information: in (**b** and **c**), the box plot analysis shows both the median and mean. Significance was calculated using Student’s *t*-test, *ab*: p<0.00001 indicates significant difference comparing *lifering-1* to Col-0 and *Atbrca1-3*, while *b*: p<0.00001 between *Atbrca1-3* and Col-0. In (**d**) *ab*:p<0.00001 shows significance of *lifering-1* both to Col-0 and *Atbrca1-3*.

Enhanced endoreplication followed by terminal differentiation is a further response to genotoxic stress protects genome integrity. To determine whether LIFERING also involved in these processes, we measured the distance between the QC and the boundary of the transition between the zones of the transit amplifying cells and elongation in the *lifering-1* mutant and compared to Col-0. As Fig. 3a illustrates, upon MMC treatment in *lifering-1*, the root meristem size is reduced compared to Col-0 and the *Atbrca1-3* mutant (Fig. 3a), suggesting that LIFERING has a role in maintaining cell proliferation and preventing early exit to cell elongation upon genotoxic stress. Treatment with genotoxic agent also led to root hair differentiation closer to the root-tip in the *lifering-1* than in Col-0, further supporting this notion.

In conclusion, due to the lack of LIFERING function, the balance between multiple DNA damage responses is shifted. Reduced cell death response is compensated for, by faster exit from cell cycle and premature cell differentiation.

### LIFERING acts downstream of RBR to maintain cell proliferation

Since LIFERING is directly repressed by RBR, we hypothesised that LIFERING is required for cell death induction and over-proliferation when *RBR* expression is silenced. To investigate this, we focused on a well-characterised phenotype of cell death and over-proliferation in the distal stem and daughter cells of the root tip (Fig. 4).

**Figure 4.**
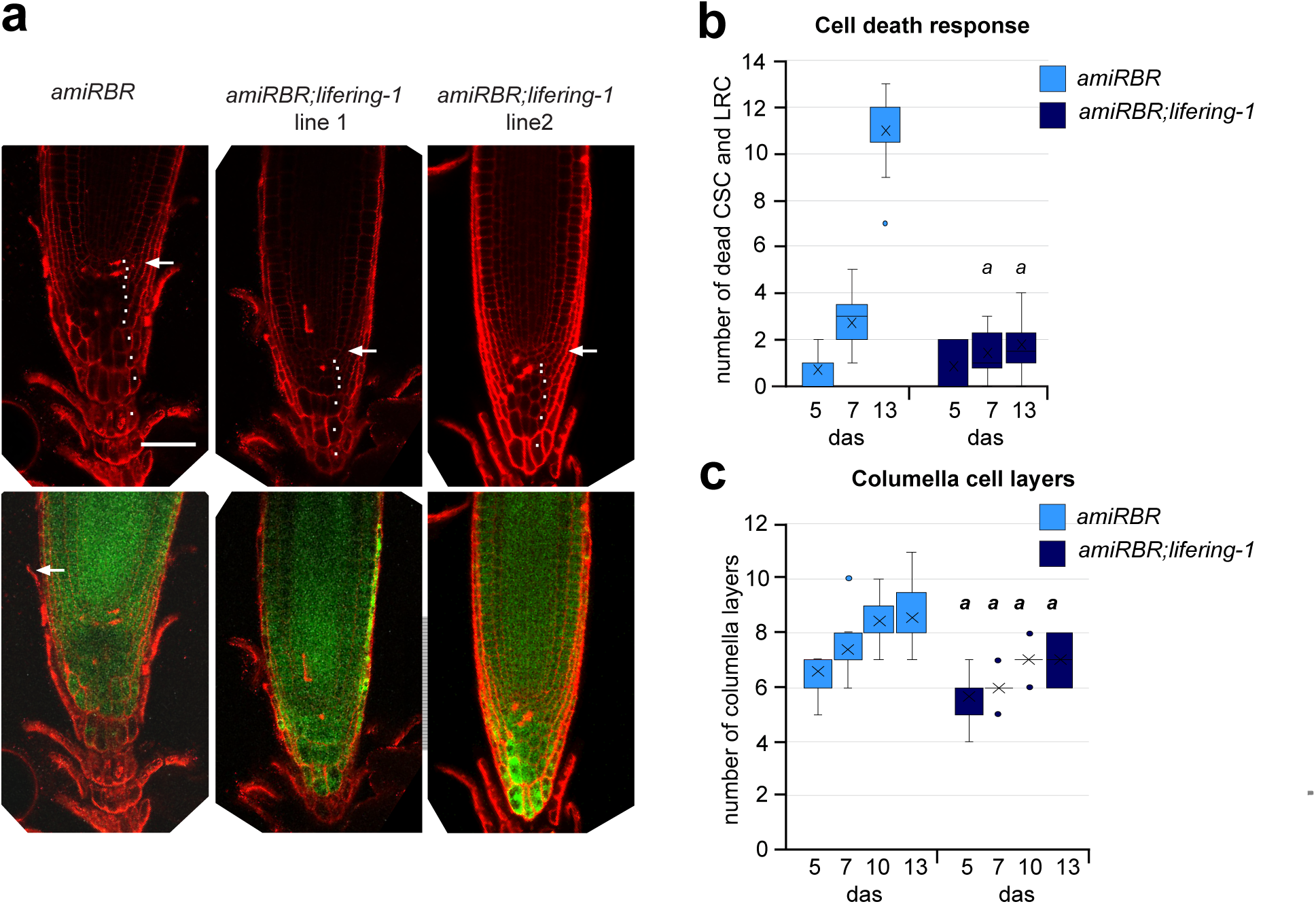
LIFERING affects cell proliferation downstream of RBR’s function. (a) The representative CM images illustrate the cell death response (PI stained cells) and the number of columella and stem cell layers (white dots). PI-stained root-tips seedlings (12-13 das) from *amiRBR* and from the independent introgressed *amiRBR;lifering-1* lines 1 and 2 are shown. White arrows point to QC position in each image. The images with GFP signals show the presence and level of silencing in the mutant lines. Scale bar: 50μm, (b) Cell death response of *amiRBR* and *amiRBR;lifering-1* seedlings at 5, 7 and 13 das. Total number of dead columella stem and daughter cells (CSC), lateral root cap initials and their descendants (LRC) were counted in the median sections as shown in (**a**). The boxplot shows the combined data of the introgressed lines, *n*>3 biological repeats, *N*>15, in each experiment. *a*: p<0.05 significant difference between the introgessed lines and *amiRBR* at a given timepoint. (c) Quantification of the number of columella stem and differentiated cell layers of 5, 7, 10 and 13 days-old roots from *amiRBR*, *amiRBR;lifering-1* lines 1 and 2. The boxplot shows the data of *n*>3, *N*>15 for each mutant in each experiment. *a*: p<0.01 significant difference between the introgessed lines and *amiRBR* at a given timepoint.

To this end, we introgressed the *lifering-1* mutant into *amiRBR* and studied the double homozygous descendants of two independent crosses verified by genotyping, the presence of GFP fluorescence, a readout for *RBR* silencing (Fig 4a) and by the lack of *LIFERING* expression upon MMC treatment (Fig. S3).

To quantify cell death response, we counted the number of dead cells in the distal stem cell niche and daughter cells (CSC and LRC) at various stages of root growth. Both cell death and over-proliferation of stem cells progressively accumulated over time. Correspondingly, at 5 das, the number of dead cells was comparable between the *amiRBR* and *amiRBR;lifering-1* lines, however, at later time points, the number of cells with cell death response started to stagnate and became significantly fewer in the *amiRBR;lifering-1* lines compared to *amiRBR* (Fig. 4a and b). The observed suppression became even more apparent at 13 das, when the cell death was limited to only few cells in the distal stem cell niche (Fig. 4b).

Similar to the suppression of cell death, the excess number of columella cell layers and disorganization of the stem cell area that progressively evolved during the period of 5 to 13 das in the *amiRBR* were also repressed in the *amiRBR;lifering-1* lines (Fig. 4c).

Based on these data, we concluded that LIFERING functions downstream of RBR both for the initiation of cell death and for maintaining cell proliferation.

### LIFERING is a plant-specific E3-ubiquitin ligase

To understand how LIFERING regulates cell proliferation and cell death, we investigated its biochemical activity. LIFERING is annotated as a putative E3 ubiquitin ligase. Our systematic comparisons to known E3 ligases revealed that LIFERING shows protein sequence homology and structural similarity to the cluster of RING-between-RING (RBR) family of E3 ubiquitin ligases. It contains a conserved three-domain module comprising RING1, BRcat (benign-catalytic) and Rcat (required-for-catalysis), along with an N-terminal reverse transcriptase-like domain (Fig. 5a) (Marin, 2010; Spratt *et al*., 2014). Accordingly, it has been suggested that LIFERING (At5g60250) belongs to the Plant II family of RING-between-RING class of E3 ubiquitin ligases (Marin, 2010).

**Figure 5.**
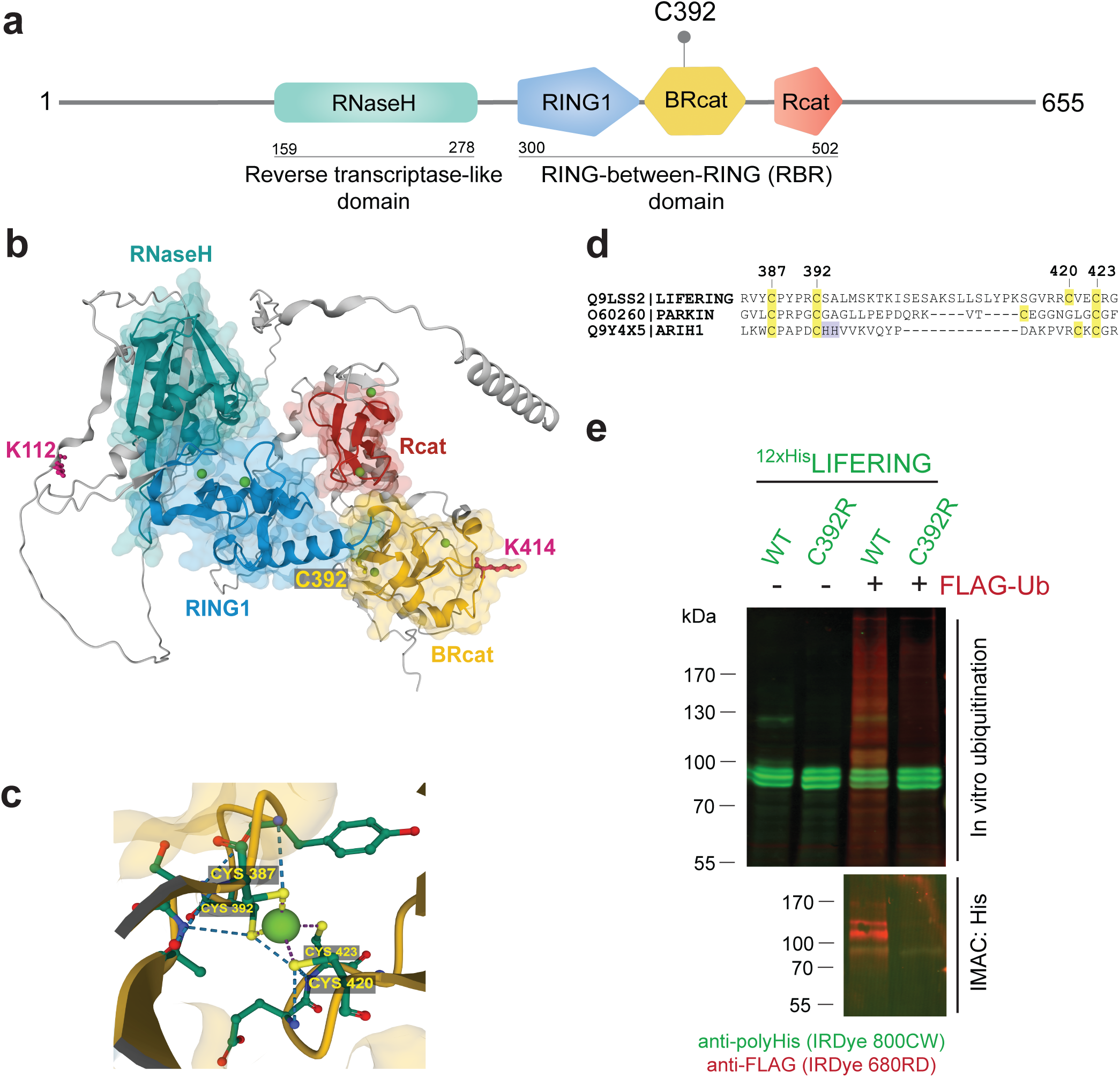
LIFERING is an E3 ubiquitin ligase. **(a)** Conserved domains of the LIFERING protein (UniProt: Q9LSS2) based on the InterPro database annotations. The Cys392 residue of the Q9LSS2 protein variant, which is Arg392 in the Q681M6 variant, is indicated. **(b)** Prediction of the three-dimensional protein structure and ligand-binding of LIFERING using AlphaFold and AlphaFill, respectively. The domains identified in (**a**) are shown with the corresponding colours (green: RNaseH domain, blue: RING1, yellow: BRcat, red: Rcat), and the six zinc ions (green beads) predicted to be bound. The ubiquitinated lysine residues identified by mass spectrometry are shown in magenta, and the Cys392 residue in the BRcat domain is illustrated in yellow. **(c)** Detailed representation of the first zinc-binding motif within the BRcat domain of LIFERING. **(d)** Sequence alignment of the first zinc-binding motif in the BRcat domain of LIFERING, Parkin and HHARI (ARIH1). Zinc-binding cysteine residues are highlighted yellow. The numbers above indicate the position of the residues in LIFERING coding sequence. **(e)** Western blot analysis of LIFERING *in vitro* ubiquitination reactions. The reactions were conducted using a combination of FLAG-tagged ubiquitin and total translation mixtures of the *in vitro* translated wild-type (WT) and C392R variants of 12xHis-LIFERING, as indicated. The E1 and E2 enzymes were sourced from endogenous wheat germ extract proteins used for *in vitro* translation. The ubiquitinated 12xHis-LIFERING proteins were isolated by pull-down (PD) using Dynabeads™ His-Tag Isolation and Pulldown beads and subsequently analysed by mass spectrometry. The proteins were separated by SDS-PAGE under reducing conditions. Multiplex detection was carried out using mouse anti-polyHis with IRDye 800CW goat anti-mouse conjugate (green) and rabbit anti-FLAG with IRDye 680RD goat anti-rabbit conjugate (red).

Associated with the At5g60250 genomic region, two different protein variants, Q9LSS2 and Q681M6 has been deposited in the UniProt database, having the genomic sequence-based and the transcript-derived amino acid sequences, respectively. The two protein variants show six amino acids differences at the N-terminal region, and deviate also at position 392 where the transcript-derived amino acid sequence codes arginine instead of the default cysteine (Fig. S4). Using AlphaFold computational protein structure prediction combined with the AlphaFill algorithm (Fig. 5b), we assumed that the cysteine at position 392, along with cysteines at positions 387, 420, and 423, are part of one of the two zinc-binding sites of the BRcat domain and are liable to engage in zinc-binding (Fig. 5c) (Jumper *et al*., 2021; Varadi *et al*., 2022; Hekkelman *et al*., 2023). To investigate the importance of this cysteine, we performed a multiple sequence alignment of this zinc-binding region in the BRcat domain of LIFERING with the two most widely studied human RING-between-RING (RBR) E3 ligases, Parkin and HHARI. We identified that Cys392 in LIFERING corresponds to the Cys281 of HHARI and Cys337 of Parkin (Fig. 5d) both of which are involved in coordination of zinc binding.

To study whether the LIFERING is a functional ubiquitin ligase, we cloned the *LIFERING* cDNA using the RIKEN clone (RAFL22-04-P06) into an *in vitro* translation vector fused with a 12xHis-tagged LIFERING. Subsequent sequencing of the coding sequence showed that the original cDNA clone and the newly constructed *in vitro* translation vector coding the 12xHis-LIFERING contained arginine at position 392 of the protein instead of cysteine. To convert arginine to cysteine, we performed an *in vitro* mutagenesis (*C*GC to *T*GC) and termed this variant the wild-type LIFERING.

To investigate the predicted ubiquitin ligase activity of LIFERING, we produced both the wild-type and mutant C392R LIFERING variants using a wheat germ-based *in vitro* translation system supplemented with zinc. We assessed their ubiquitin ligase activity through *in vitro* protein ubiquitination assays. To this end, we supplemented the crude translation mixtures with FLAG-tagged ubiquitin (FLAG-Ub), while the control reaction was performed without ubiquitin. To detect the expression of the two LIFERING variants and the ubiquitination status, a polyhistidine-, and a FLAG-specific antibodies, respectively, were used in Western blot analysis (Fig. 5e). Although the wild-type and the C392R LIFERING proteins were detected in equal amounts in both the FLAG-Ub supplemented and control reactions, we observed a significantly higher level of ubiquitination in the protein extracts containing the wild-type protein compared to the mutant variant (Fig. 5e, lanes 3 and 4).

From these results we conclude that LIFERING exhibits ubiquitin ligase activity, which is critically dependent on the presence of Cys at position 392.

To investigate whether LIFERING undergoes autoubiquitination, we isolated both the wild-type and the mutant 12xHis-labelled LIFERING proteins from the ubiquitination reaction mixture (Fig. 5e, lanes 3 and 4) using immobilized metal affinity chromatography (IMAC) and performed an additional Western blot analysis using anti-FLAG antibodies. As Fig. 5e shows, we detected a ladder-like pattern of ubiquitination exclusively in the presence of the wild-type LIFERING. The proteins bound to the IMAC beads were further analysed using mass spectrometry to reveal the position of the ubiquitinated amino acids. As a result of this analysis, isopeptide-linked ubiquitins were detected at Lys112 and Lys414 exclusively in the samples containing the wild-type LIFERING variant (Fig. S5), indicating that LIFERING can undergo autoubiquitination in the wheat germ-based *in vitro* translation system.

### LIFERING ubiquitinates IBR3, the enzyme involved in the indole-3-acetic acid synthesis

To understand LIFERING’s function, we set out to identify its target substrates for ubiquitination. We leveraged the TurboID-based biotin proximity labelling technique (Branon *et al*., 2018). The wild-type and mutant LIFERING proteins were cloned in CaMV 35S promoter-driven vectors coding for TurboID at the N-terminal of the fusion proteins and *Arabidopsis* root cell suspension-derived protoplasts were transfected with these constructs. Using streptavidin-coated magnetic beads, the biotinylated proteins were isolated and subsequently identified by protein mass spectrometry. The spectral count data from numerous independently performed experiments with wild-type and mutant samples were analysed using SAINTexpress software (Teo *et al*., 2014). Among the numerous potential targets, the acyl-CoA dehydrogenase IBR3 (At3g06810) was repeatedly identified as an interactor with both wild-type and mutant LIFERING proteins (FoldChange >100, Table S5) relative to the control runs with TurboID alone.

To validate the proximity labelling results, first we generated the GST-IBR3 fusion protein through *in vitro* translation, then we investigated in an *in vitro* ubiquitination assay the functional interaction between LIFERING and IBR3. The ubiquitination reactions were carried out with the crude translation mixtures containing wild-type or mutant 12xHis-LIFERING in the presence of GST-tagged IBR3 bound to the glutathione beads. The results of the pull-down assay were analysed by Western blotting using different antibodies. Through application of polyHis-specific antibody, we found that both the wild-type and mutant LIFERING could interact with IBR3, and using FLAG- and GST-specific antibodies, we observed ubiquitination of the glutathione bound IBR3 exclusively by the wild-type LIFERING (Fig. 6a). The position of ubiquitination of IBR3 by LIFERING was also determined through mass spectrometry. Analysis of the bead-bound IBR3 treated either by the wild-type or mutant LIFERING corroborated the results obtained by Western blot. Four lysine residues of IBR3 (Lys170, 213, 311, and 536) were identified as ubiquitinated amino acids in the wild-type LIFERING containing sample. In contrast, no ubiquitination was detected in the reaction mixture containing the mutant C392R LIFERING (Fig. S6).

**Figure 6.**
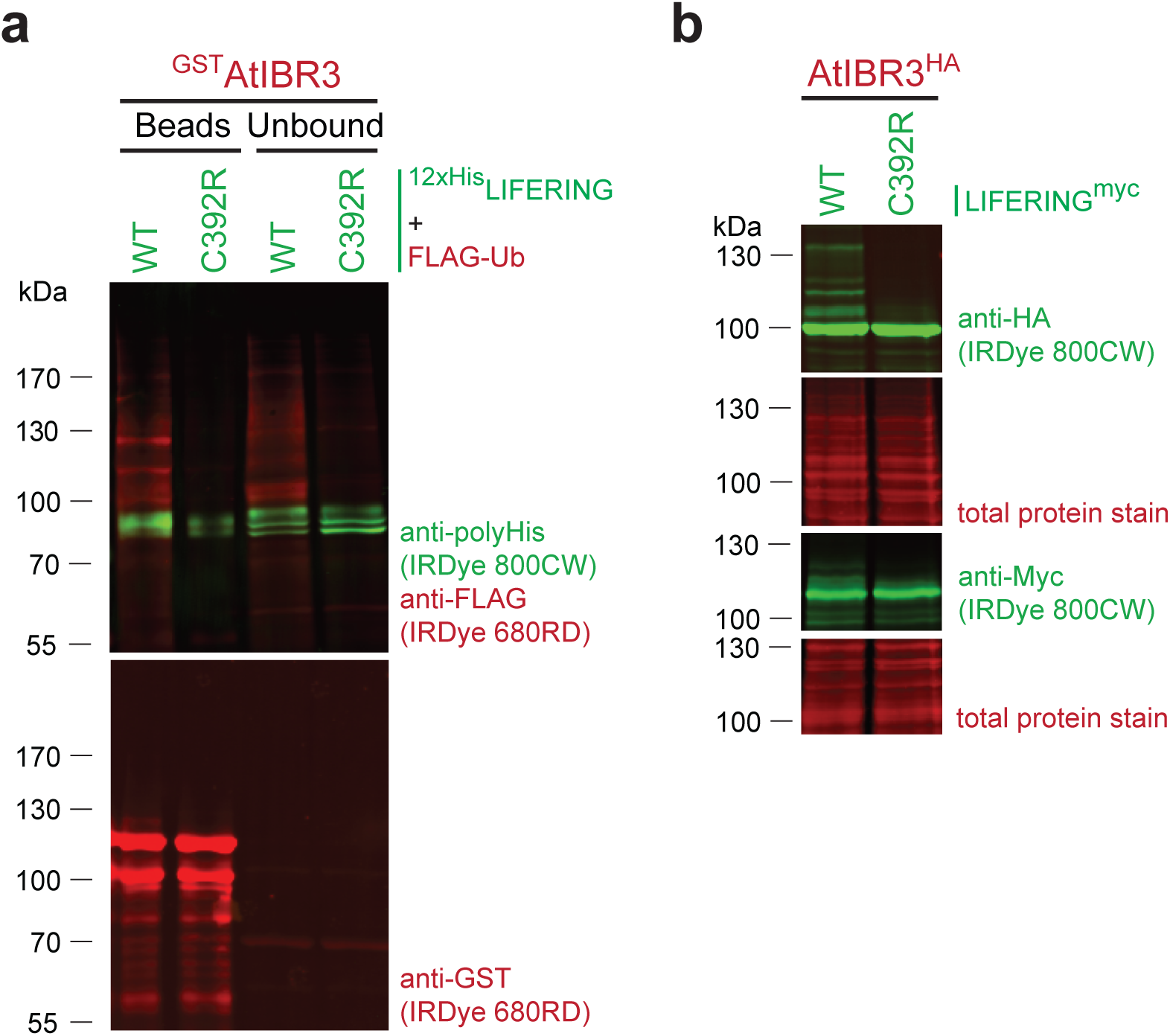
LIFERING ubiquitinates AtIBR3 *in vitro* and *in vivo*. **(a)** Western blot analysis of glutathione beads-bound AtIBR3 *in vitro* ubiquitination reactions in the presence of wild-type (WT) and C392R variants of LIFERING. FLAG-tagged ubiquitin and total translation mixtures of the *in vitro* translated wild-rype (WT) and C392R variants of 12xHis-LIFERING were added to the bead-bound AtIBR3, as indicated. Subsequently, the unbound fraction and washed beads were analysed following the *in vitro* ubiquitination reaction. The proteins were separated by SDS-PAGE under reducing conditions. Multiplex detection was carried out using mouse anti-polyHis antibody and rabbit anti-FLAG antibody. In the bottom part of (**a**), the singleplex detection of the bead-bound GST-IBR3 using rabbit anti-GST antibody is shown. **(b)** Western blot analysis of *Arabidopsis* root cell suspension protoplasts co-transfected with wild-type or mutant Myc-tagged LIFERING and HA-tagged AtIBR3-encoding vectors. Singleplex detection was carried out using mouse anti-HA and mouse anti-Myc, and the membranes were stained with Revert 700 Total Protein Stain to check for loading consistency across the wells. The secondary antibodies used in (**a** and **b**) were IRDye 680RD goat anti-rabbit conjugate (red) and IRDye 800CW goat anti-mouse conjugate (green).

To determine whether LIFERING can also catalyse the ubiquitination of IBR3 *in vivo*, we co-transfected protoplasts derived from *Arabidopsis thaliana* root cell suspensions with different combinations of C-terminal Myc-tagged wild-type and mutant LIFERING variants and HA-tagged IBR3-encoding vectors. Immunoblot analysis of the protein extracts revealed that both the Myc- and HA-specific antibodies decorated the overexpressed proteins, confirming the production of the proteins of interest. A ladder-like pattern of IBR3 was also clearly visible when co-transfected with the wild-type LIFERING and using anti-HA antibody. Furthermore, the wild-type LIFERING also exhibited a ladder-like pattern not detected with the mutant variant by using the anti-Myc antibody (Fig. 6b). These *in vivo* observations were in line with the *in vitro* results, implying that only the wild-type Cys392-containing LIFERING is capable of autoubiquitination and the ubiquitination of IBR3.

Ubiquitination of proteins can lead to different outcomes, i.e. mono- and multi/mono-ubiquitination can often regulate protein activity and localisation, whereas polyubiquitination is conventionally regarded as a signal for proteasomal degradation. To study whether ubiquitination of IBR3 by LIFERING results in its hydrolysis in the proteasome, we transfected the protoplasts with IBR3-encoding vector alone and with wild-type LIFERING-encoding vector together. After overnight incubation in the dark, the cells were treated with cycloheximide (CHX) and MG132 to block *de novo* protein synthesis and proteasomal activity, respectively. According to the Western blot analysis, IBR3 is a highly stable protein, with only a negligible difference in its protein levels detected even after 24 hours incubation in the presence of CHX. The combined addition of CHX and MG132 had no discernible effect on the protein amounts detected either (Fig. S6). The same phenomenon was observed with the co-transfected protoplast-derived samples. While the sign of ubiquitination, the ladder-like pattern of IBR3 was present, there was only an insignificant decrease in its level (Fig. S6). Our findings suggest that the ubiquitination of IBR3 by LIFERING may serve a distinct, yet to be identified function rather than proteasomal degradation.

## DISCUSSION

The importance of RING domain proteins in plants is highlighted by their genomic over-representation compared to other eukaryotes (Kosarev *et al*., 2002). Here, we show that LIFERING, a member of this protein family known to be rapidly upregulated upon many stress signals and is a catalytically active E3 ubiquitin ligase that ubiquitinates IBR3, the acyl-CoA dehydrogenase-like protein.

Structurally, LIFERING has a RING1-BRcat-Rcat signature and has been assigned to the Plant II subfamily of RING-between-RING-type E3 ubiquitin ligases (Marin, 2010). In addition to *in silico* analysis, we also performed biochemical experiments and demonstrated that LIFERING is an active ubiquitin ligase; its activity depends on the presence of zinc ions and Cys392 which corresponds to one of the zinc coordinating cysteines of Parkin, the human RBR type E3 ligase, one of the closest mammalian homologues of LIFERING. Previous research demonstrated that the biological activity of Parkin is altered by mutation of zinc-binding cysteines (Beasley *et al*., 2007), which is in alignment with our observation.

LIFERING has an essential function to maintain genome stability *in planta*. It is a universal DNA damage and stress response gene; highly induced independently of whether the damage is caused by exogenous genotoxic agents or endogenous stresses due to mutations in genes involved in DNA replication and repair and their regulation, irrespective of the nature of the damage in the DNA structure. A genome-wide study to identify direct targets of SOG1 in response to γ-radiation also uncovered LIFERING as a target (Bourbousse *et al*., 2018). Here we show that upstream to SOG1, LIFERING expression is regulated by ATM/ATR signalling. Independently and parallel to this, it is directly repressed by RBR, partly through E2FA. As a response to DNA damage, besides induction of gene expression, cells mitigate the effects of dividing with damaged DNA by inhibition of cell cycle and induction of cell death. LIFERING has a role in triggering both processes. It is essential to induce cell death response both upon exo- and endogenous DNA damage.

The so-far identified E3 ligases involved in DDR either play a role in the recognition of the damage site and/or its repair or in the regulation of cell cycle progression (Herbst *et al*., 2024), and structurally do not belong to the RING1-BRcat-Rcat class of E3 ubiquitin ligases. The limited data available on the members (22) of the largest Plant II subfamily of *Arabidopsis thaliana* RING-between-RING-type ligases suggest that they are mainly involved in phytohormone mediated regulation of stress responses (Wang *et al*., 2020). RING FINGER OF SEED LONGEVITY1 (RSL1) (Bueso *et al*., 2014a; Bueso *et al*., 2014b), RING Finger ABA-Related1 (RFA1), and RFA4 (Fernandez *et al*., 2020) target ABA receptors for degradation via proteasomal or vacuolar pathways. Similar to the members of PlantII subfamily, the ARIADNE12 (AtARI12) belongs to the Ariadne subfamily of plant RING-between-RING ubiquitin ligases (Mladek *et al*., 2003) and is also stress responsive. Its expression is strongly up-regulated by UV-B irradiation and depends on the activity of the E3 ligase *CONSTITUTIVELY PHOTOMORPHOGENIC1 (COP1*), in both white and UV-B light (Xie *et al*., 2015). As the increase of *AtARI12* mRNA levels did not correlate with the abundance of AtARI12 protein levels, moreover, using proteasome inhibitor MG132 had no effect on its stability, the authors proposed that AtARI12 protein levels are unlikely to be regulated by proteasome-mediated degradation (Xie *et al*., 2015).

In searching for putative substrates of LIFERING, we found that it readily ubiquitinates the acyl-CoA dehydrogenase-like protein, IBR3, an enzyme involved in the conversion of IBA to indole-3-acetic acid IAA. IBR3 was ubiquitinated both *in vitro* and in the protoplast transient overexpression system, visualised by the characteristic ladder-like pattern of the protein with different molecular weight. Furthermore, the ubiquitinated lysines of IBR3 protein were identified through mass spectrometry. The ubiquitin code can modify protein function in different ways. While K48-linked polyubiquitin chains mainly target proteins for proteasomal degradation, proteasome-independent processes are usually mediated by non-K48-linked ubiquitin chains affecting their stability, activity, interactions with other proteins and their subcellular localisation. In plants, the K63-linked chains are identified as the second most abundant modification and proposed to play proteasome-independent roles in hormonal responses and development, nutritional responses, biotic interactions and DNA repair and autophagy targeting (Romero-Barrios & Vert, 2018; Orosa-Puente & Spoel, 2022). Despite the application of proteosome and protein synthesis inhibitors (MG132 and CHX), no significant change in the IBR3 protein level could be detected, suggesting that ubiquitination of IBR3 by LIFERING likely has a different biological function than proteasomal degradation. The conversion of IBA to IAA is a multistep enzymatic reaction during which the auxin precursor IBA undergoes peroxisomal oxidation to release free IAA (Damodaran & Strader, 2019). The conversion plays an essential role in the spatial determination of the auxin level, and thus controls a distinct set of plant developmental and growth effects (Strader *et al*., 2011). It also provides a local auxin source in the root cap (Xuan *et al*., 2015), while along the root axis, the IBA-derived auxin is responsible, on the one hand, for setting the local auxin level to position lateral root formation (Xuan *et al*., 2015) and, on the other hand, in co-operation with cytokinin, for the formation of adventitious root (Damodaran & Strader, 2024).

In the root tip, IBR3 is localized to the lateral root cap; mostly to the lateral - and columella root cap cells, but also in the QC, dividing and young meristem cells, albeit at a lower level. In the differentiation zone, its expression was identified in the stele cells including the pericycle (Xuan et al., 2015; Ryu, KH et al., 2019). Importantly, the expression of IBR3 overlaps with that of LIFERING in the lateral - and columella root cap cells and the dividing meristematic cells, albeit on a significantly lower level in non-genotoxic conditions (Ryu, KH et al., 2019). In the triple IBA response mutants, *ibr1,ibr3,ibr10,* the overall IAA level is unchanged (Strader *et al*., 2010), while in the quadruple mutant, *ech2*,*ibr1,ibr3 ibr10*, the free auxin level is reduced with 20-25% in the root tip. As expected from the local auxin levels, the quadruple mutant develops a significantly shorter meristem (Strader *et al*., 2011), and an overall reduced root system with fewer number of lateral roots (Xuan *et al*., 2015). However, despite the morphological changes, no spontaneously developed cell death response was reported related to any of these mutants in normal growth conditions (Strader *et al*., 2011).

### How can we integrate our data in the existing knowledge on the connection between DNA damage response and hormonal regulation?

Root growth is dependent on the balance of the hormones, cytokinin and auxin (Salvi *et al*., 2020). Upon DNA damage, however, the balance between cytokinin and auxin levels shifts. Through SOG1, DNA damage induces cytokinin synthesis and activates its signalling around the transition zone, thereby inhibiting downward auxin flow. With the reduction in auxin level, the position of auxin minima changes towards the stem cell niche, resulting in the G2 arrest of the dividing cells. Independent of G2 arrest, reduced auxin signalling also leads to stem cell death. In parallel, increased cytokinin signalling triggers early onset of endoreduplication. The combined effects of G2 arrest, early transition from division to endoreduplication and stem cell death, reduce meristematic activity and thus inhibit root growth to safeguard genome integrity (Davis *et al*., 2016; Takahashi *et al*., 2021). The fact that, in the absence of functional LIFERING, there was no or significantly reduced cell death in the root meristem, either upon treatment with genotoxic agent or in silencing of RBR, especially in the stem cell niche, we suggest that LIFERING plays a role in setting the auxin level in the stem cell niche. Furthermore, closer endoreduplication in the meristem also indicates a shift in hormone balance.

Both the suppression of auxin signalling and auxin levels are prerequisites to the prompt local development of cell death response in the root meristem to protect genome integrity (Takahashi *et al*., 2021). Upon DNA damage, the otherwise lowly expressed *IAA5* and *IAA29* genes, coding for the transcriptional repressors of auxin signalling, are directly induced by SOG1 in the vascular stem cells and daughters. This way, they can locally reduce the auxin response and ensure cell death upon damage in the stem cell niche and surrounding vascular cell layer. In contrast, in the double mutant *iaa5-1, iaa29-1*, due to lack of suppression of the local auxin level signalling, spatial reduction of the cell death response was detected in the apical meristem when challenged with DNA damage agent (Takahashi *et al*., 2022). Similar to the *iaa5-1,iaa29-1*, the mutant line, *lifering-1* develops no or substantially weakened cell death response upon DNA damage and thus has significantly reduced number of dead cells in both the apical and distal meristems with no or only one dead cell either in the distal and/or in the vascular stem cell niche compared to the Col-0.

In line with the overlapping localisation of LIFERING and IBR3 and considering that LIFERING targets IBR3 for ubiquitination and consequently potentially inactivates it by an-as-yet-unknown mechanism, we propose that under normal growth conditions, in the absence of LIFERING in the root, the IBA to IAA conversion supplies auxin to support growth along the root meristem. In contrast, under stress, when *LIFERING* is rapidly induced, it attenuates the auxin supply, thereby leading to growth arrest and early differentiation. By setting the balance between cell proliferation and differentiation, LIFERING through auxin signalling also regulates cell death response upon DNA damage.

In conclusion, we propose a working model according to which we suggest that by adjusting the level of free auxin available in the root meristem, LIFERING initiates a cell death response to prevent further division with damaged DNA and to fine-tune the auxin level in the QC and the surrounding area. Further experimentation should provide evidence to prove this model.

## ACKNOWLEDGEMENT

We thank Hana Kourová and Pavla Binarová (Czech Academy of Sciences, Czech Republic) for their valuable advice during the experimentation. We thank Christian W.B. Bachem (Solynta, The Netherlands) for critically reading the manuscript.

## AUTHORS CONTRIBUTIONS

B.M.H., L.B., B.M.K. and T.M. conceived and designed the original research plan. B.M.H., E.N., F.N., and Z.A. carried out the expressional, phenotypic and microscopical studies on the mutant and the complementation lines. B.M.K. and T.M. designed and carried out the *in vitro* translation, protoplast transient expression, pull-down and immuno-blotting assays, B.M.K. performed the ubiquitination assays, the Turbo-ID proximity labelling and generated the vector constructs essential for the above experiments. B.M.K. provided material for and A.P-Sz. carried out mass spectrometry analysis of proximity labelling derived samples. A.P-Sz. And B.M.K. evaluated the MS data. Z.M. and L.B. shared material and discussed the experiments and data. The manuscript was written by B.M.H., B.M.K, and T.M., it was read and commented by all the authors.

## SUPPORTING INFORMATION

### Supporting Information on Materials and Methods

#### Cloning

The N-terminal 3xHA-TurboID-tagging pRT-3xHA-Turbo-LIC vector was created by replacing AtMPK9 in the pRT-HA-AtMPK9 vector (Nagy *et al*., 2015) with the 3xHA-TurboID coding sequence. Because the LIC cassette must contain unique *Ssp*I restriction site, the one found in the ampicillin resistance gene promoter of the original vector was removed through in vitro mutagenesis (Liable & Boonrod, 2009). Subsequently, AtMPK9 insert was removed by digestion with *Nco*I and *Bam*HI. A template containing the coding sequence for 3xHA-TurboID was provided in a pMA-RQ vector. 3xHA-TurboID was amplified by PCR with primers coding for the *Ssp*I-harbouring ligation-independent cloning (LIC) cassette (Bardóczy *et al*, 2008) and inserted into the pRT-HA vector between the *Nco*I and *Bam*HI restriction sites.

To produce FLAG-tagged ubiquitin, the coding sequence of the ubiquitin monomer was optimized for wheat germ codons and checked for *E. coli* incompatible codons. The obtained sequence was chemically synthesized with LIC-compatible overhangs by Integrated DNA Technologies. Ubiquitin was amplified by PCR and inserted into the wheat-germ in vitro translation compatible pEU3-NII-FLAGLICNot vector (Nagy *et al*, 2020) using LIC. To produce larger amount of ubiquitin using the bacterial protein expression system, the FLAG-ubiquitin coding DNA was amplified by PCR from the pEU3-NII-FLAGLICNot_Ubiquitin vector described above and subcloned into the pET28a vector using *Nco*I and *Nde*I restriction endonucleases.

#### Bacterial FLAG-ubiquitin expression

FLAG-ubiquitin (FLAG-Ub) was expressed in BL21(DE3) competent *E. coli* cells (New England BioLabs) via autoinduction as previously described (Studier, 2005). A positive colony was inoculated into MDG non-inducing medium with 50 μg/ml kanamycin. After 5 hours of growth at 37°C, the starter culture was used to inoculate ZYM-5052 autoinducing medium with 50 μg/ml kanamycin at a 1:400 dilution and grown overnight at 37°C. The bacterial culture was centrifuged at 4000 rpm, 4°C for 20 minutes, and the resulting pellet was resuspended in 1/5– 1/6 volume of extraction buffer (300 mM NaCl, 10 mM imidazole, 0.1% Triton-X) relative to the original culture volume. The resuspended cells were sonicated for 5 cycles of 30 seconds each (30s on, 30 s off. The cell lysate was then centrifuged at 13000 rpm and 4°C for 20 minutes.

The resulting supernatant was heated in 500 μl aliquots at 85°C for 5 minutes and chilled on ice. Following centrifugation at 13000 rpm for 10 minutes at 4°C, the soluble fraction of FLAG-Ub was purified using the flow-through from a 30K MWCO Amicon centrifugal filter (EMD Millipore) and concentrated with a 3K MWCO Amicon centrifugal filter (EMD Millipore). The concentration and purity of FLAG-Ub were determined by Tricine-SDS-PAGE (Haider *et al*, 2012) and densitometry analysis (data not shown). The concentration of the FLAG-Ub stock solution was adjusted to 200 μM, and aliquots were stored at −20°C.

#### Transient protein expression and proximity labelling

The Columbia (Col-0) *Arabidopsis thaliana* root-cell suspension culture was maintained according to Mathur and Koncz (Mathur & Koncz, 1998). Protoplast isolation and PEG-mediated protoplast transfection were conducted as previously described with slight modifications (Yoo *et al*., 2007). To transfect 2×10^5^–5×10^5^ protoplasts, 1 μg of HA-TurboID and HA-TurboID-LIFERING (wild-type and C392R) coding pRT plasmid DNAs were used. The transfected cells were cultured in 24-well plates for 16 hours in the dark at room temperature. Through empirical testing we identified the optimal biotin concentration and incubation time of biotin labelling as 50 μM and 3–4 hours, respectively (data not shown). Proximity labelling was then carried out by adding d-biotin at a final concentration of 50 μM, zeocin at a final concentration of 0.05 μg/μl, when indicated, and incubating for 4 hours in the dark at room temperature. For each sample, 2×10^6^ cells were harvested and lysed using a modified ice-cold RIPA buffer (50 mM Tris-HCl pH 7.8, 150 mM NaCl, 1% IGEPAL® CA-630, 0.5% sodium deoxycholate, 0.1% SDS, 1X protease inhibitor cocktail, 15 mM EGTA, 1 mM DTT, 1 mM NaF, 0.5 mM activated Na_3_VO_4_, 0.5 mM β-glycerophosphate, 1X PhosStop, 50 μM MG132) and incubation in an ultrasonic bath for 2 minutes. The extracts were centrifuged at 13000 rpm, 4°C for 15 minutes and the supernatants were desalted with 7K MWCO Zeba™ Spin Desalting Column to remove free biotin and analysed by mass spectrometry.

#### Mass spectrometry sample preparation and analysis

In case of TurboID experiments, the biotinylated proteins were purified from the cleared extracts by using μMACS^TM^ Streptavidin beads (Milteny Biotec). They were then digested in column with trypsin and analysed in a single run by mass spectrometry. The digestion mixtures were acidified and transferred to a single-use trapping mini-column (Evotip; 1/8 of the samples) and then analysed with a data-dependent LC-MS/MS method using an Evosep One (LC: 15 SPD; MS1: R:120,000) on-line coupled to a linear ion trap-Orbitrap (Orbitrap-Fusion Lumos, Thermo Fisher Scientific) mass spectrometer operating in positive ion mode. Data acquisition was carried out in data-dependent fashion, multiply charged ions were selected in cycle-time from each MS survey scan for ion-trap HCD fragmentation (MS spectra were acquired in the Orbitrap (R=60000) while MSMS in the ion-trap). The raw data was converted into peak lists using Proteome Discoverer (v1.4) and searched against the Uniprot Arabidopsis thaliana database (downloaded on 12 June 2019, containing 89,461 proteins) using our in-cloud Protein Prospector search engine (v5.15.1). Spectral counting was employed to estimate the relative abundance of individual proteins and were analysed by SAINTexpress (version 3.6; https://saint-apms.sourceforge.net/Main.html) using default parameters.

For the mass spectrometry analysis of in vitro ubiquitinated LIFERING and IBR3, 1/20 of the bead-bound protein samples were used for Gel-aided sample preparation (GASP) (Fischer & Kessler, 2015). The remaining LIFERING samples were concentrated using a 10kDa Amicon Ultra-0.5 Centrifugal Filter Unit and then mixed with 0.5 ml of Lysis Buffer (Miltenyi Biotec). They were then immunopurified using anti-FLAG antibody-coupled magnetic beads (MACS® Technology, Miltenyi Biotec, d=50 nm), digested in a column with trypsin, and analysed in a single run by mass spectrometry (Kobayashi *et al*, 2015, Hubner *et al*, 2010, Jankovics *et al*, 2018). The digestion mixtures were acidified and transferred to a single-use trapping mini-column (Evotip; 1/8 of the samples). Subsequently, they were analysed using a data-dependent LC-MS/MS method. The LC-MS/MS method was performed using an Evosep One (LC: 15 SPD; MS1: R :120,000) on-line coupled to a linear ion trap-Orbitrap (Orbitrap-Fusion Lumos, Thermo Fisher Scientific) mass spectrometer operating in positive ion mode. Data acquisition was performed in a data-dependent manner. Multiply charged ions were selected in cycle-time from each MS survey scan for ion-trap HCD fragmentation. MS spectra were acquired in the Orbitrap (R=60000), while MSMS was acquired in the ion-trap. The raw data was converted into peak lists using the in-house Proteome Discoverer (v 1.4). The peak lists were then searched against the Uniprot WHEAT and Arabidopsis thaliana database, as well as 6 user-defined proteins (downloaded on 20th July 2022, containing 280426 proteins), using our in-cloud Protein Prospector search engine (v5.15.1). with the following parameters: enzyme: trypsin with maximum 2 missed cleavage; mass accuracies: 5 ppm for precursor ions and 0.6 Da for fragment ions (both monoisotopic); fixed modification: carbamidomethylation of Cys residues; variable modifications: acetylation of protein N-termini; Met oxidation; cyclization of N-terminal Gln residues, GlyGly (Uncleaved K), LeuArgGlyGly (Uncleaved K), Phospho (STY) allowing maximum 2 variable modifications per peptide. Acceptance criteria: minimum scores: 22 and 15; maximum E values: 0.01 and 0.05 for protein and peptide identifications, respectively.

## Supporting Tables

**Table S1:**
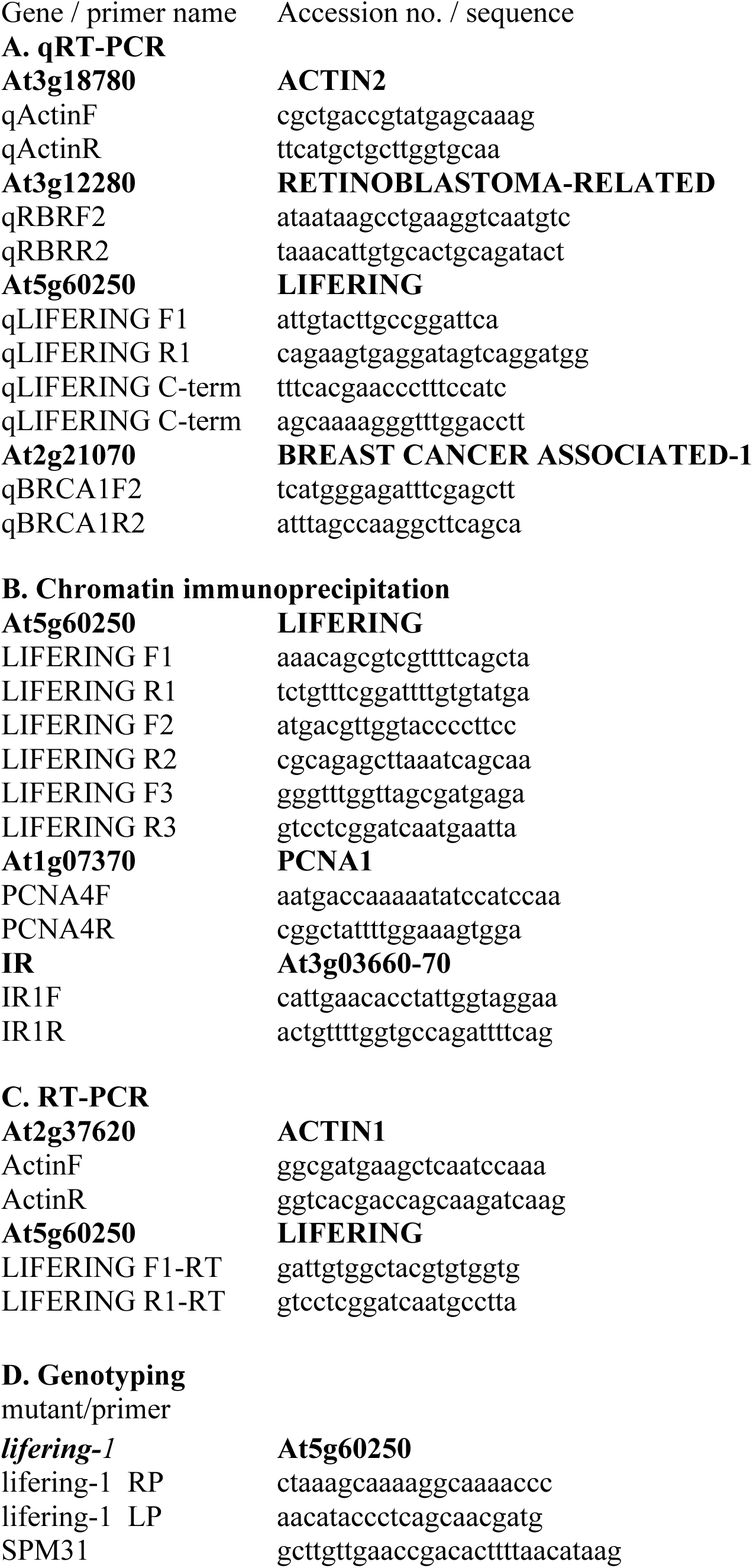
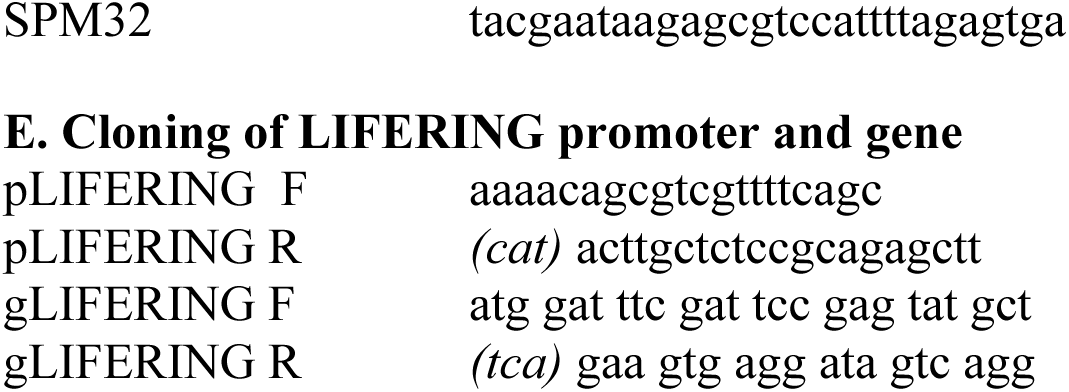
List of primers used for *in planta* expressional studies.

**Table S2:**
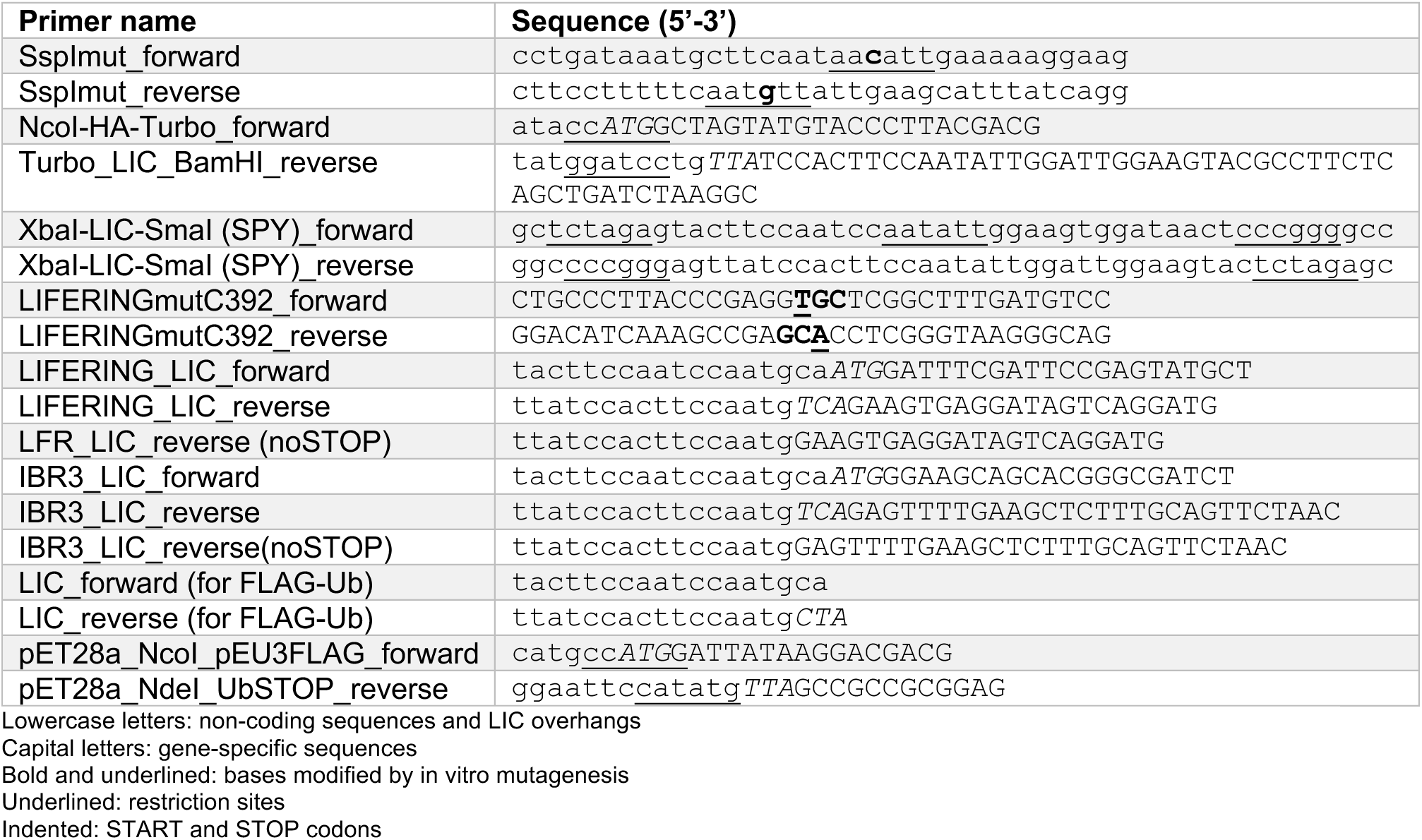
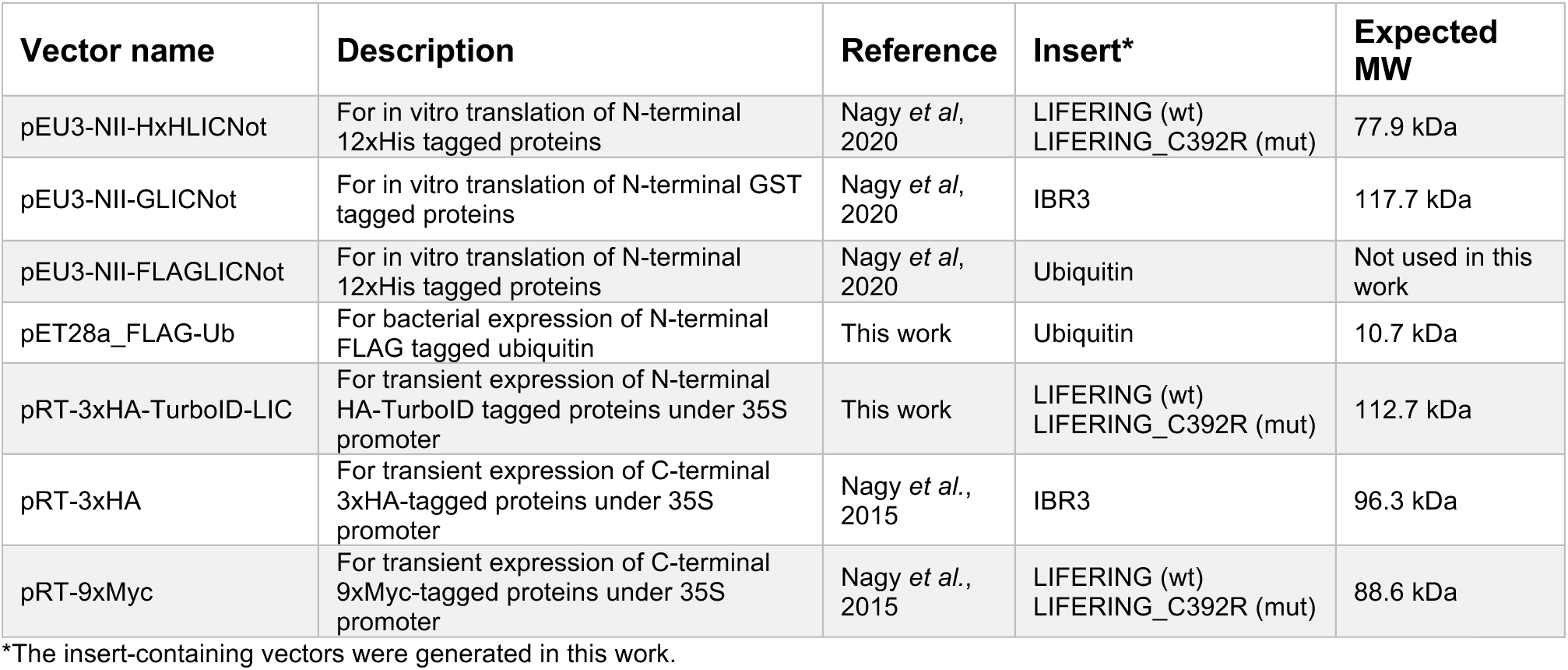
Primers and vectors used for *in vitro* translation and transient protein expression experiments.

**Table S3:** Antibodies used in this study.

| Antibody | Clone | Dilution | Source |
| --- | --- | --- | --- |
| Mouse anti-polyHistidine | HIS-1 | 1:2000 | Sigma |
| Rabbit anti-GST | 06-332 | 1:2000 | Upstate, Merck KGaA |
| Rabbit anti-FLAG | F7425 | 1:1000 | Sigma |
| Mouse anti-HA | HA-7 | 1:1000 | Sigma |
| Mouse anti-c-Myc | 9E10 | 1:1000 | Roche |
| IRDye® 800CW goat anti-mouse | N/A | 1:20000 | LI-COR Biosciences |
| IRDye® 680RD goat anti-rabbit | N/A | 1:20000 | LI-COR Biosciences |

**Table S4:**
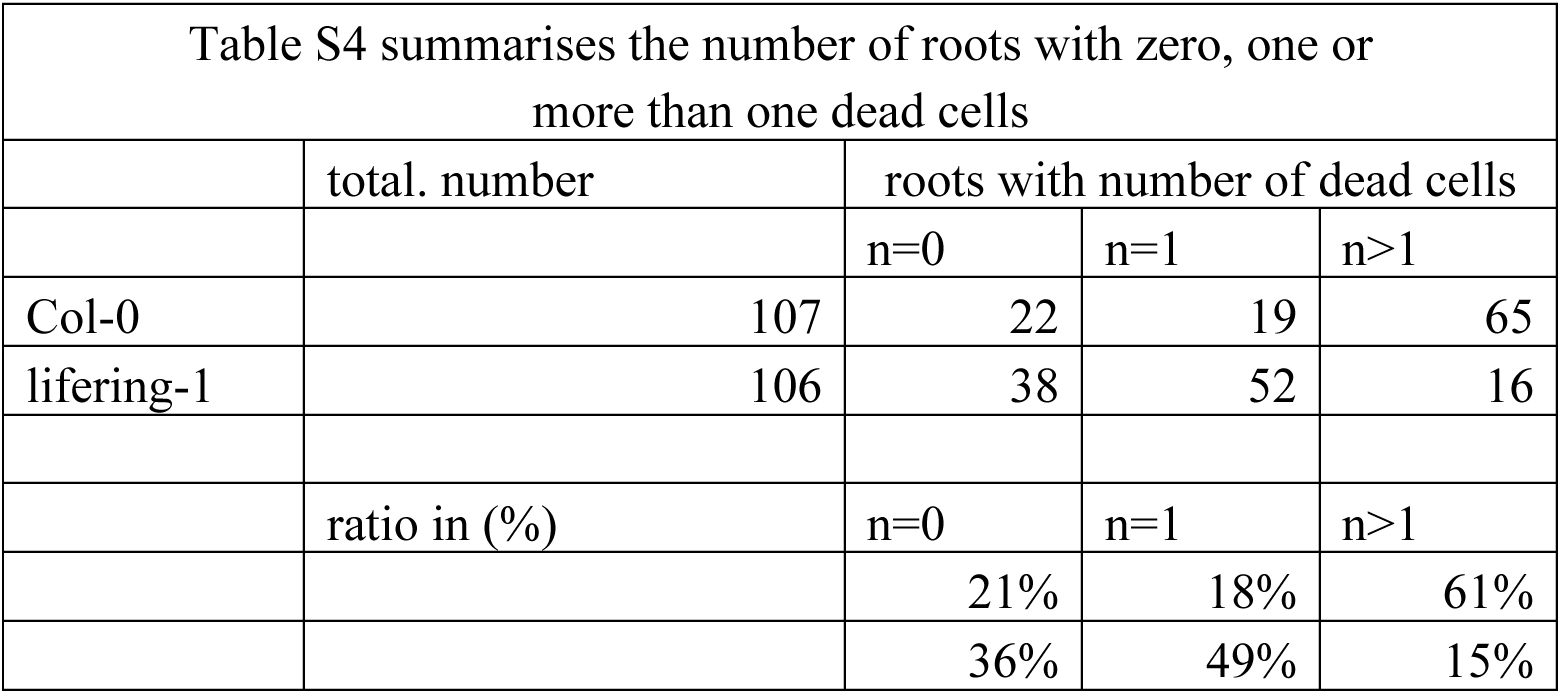
Summary of the roots with different number of dead cells.

| Table S4 summarises the number of roots with zero, one or more than one dead cells |  |  |  |  |
| --- | --- | --- | --- | --- |
|  | total. number | roots with number of dead cells |  |  |
|  |  | n=0 | n=1 | n>1 |
| Col-0 | 107 | 22 | 19 | 65 |
| lifering-1 | 106 | 38 | 52 | 16 |
|  | ratio in (%) | n=0 | n=1 | n>1 |
|  |  | 21% | 18% | 61% |
|  |  | 36% | 49% | 15% |

**Supplementary Table S5.** Analysis of mass spectrometry spectral count data from TurboID proximity labelling experiments using SAINTexpress (v3.6). RINGwt is the wild-type LIFERING protein with Cys392, RINGmut is the mutant LIFERING with an arginine at position 392 of the protein coding sequence.

| Identifier | Bait name | Zeocin treatment | Replicate |
| --- | --- | --- | --- |
| RINGmut | TurboID-RINGmut | no | 1 |
| RINGmut_z | TurboID-RINGmut | yes | 1 |
| RINGwt | TurboID-RINGwt | no | 1 |
| RINGwt_z | TurboID-RINGwt | yes | 1 |
| TbID_1 | TurboID | no | 1 |
| TbID_2 | TurboID | yes | 2 |
| TbID_3 | TurboID | no | 3 |

**Figure S1.**
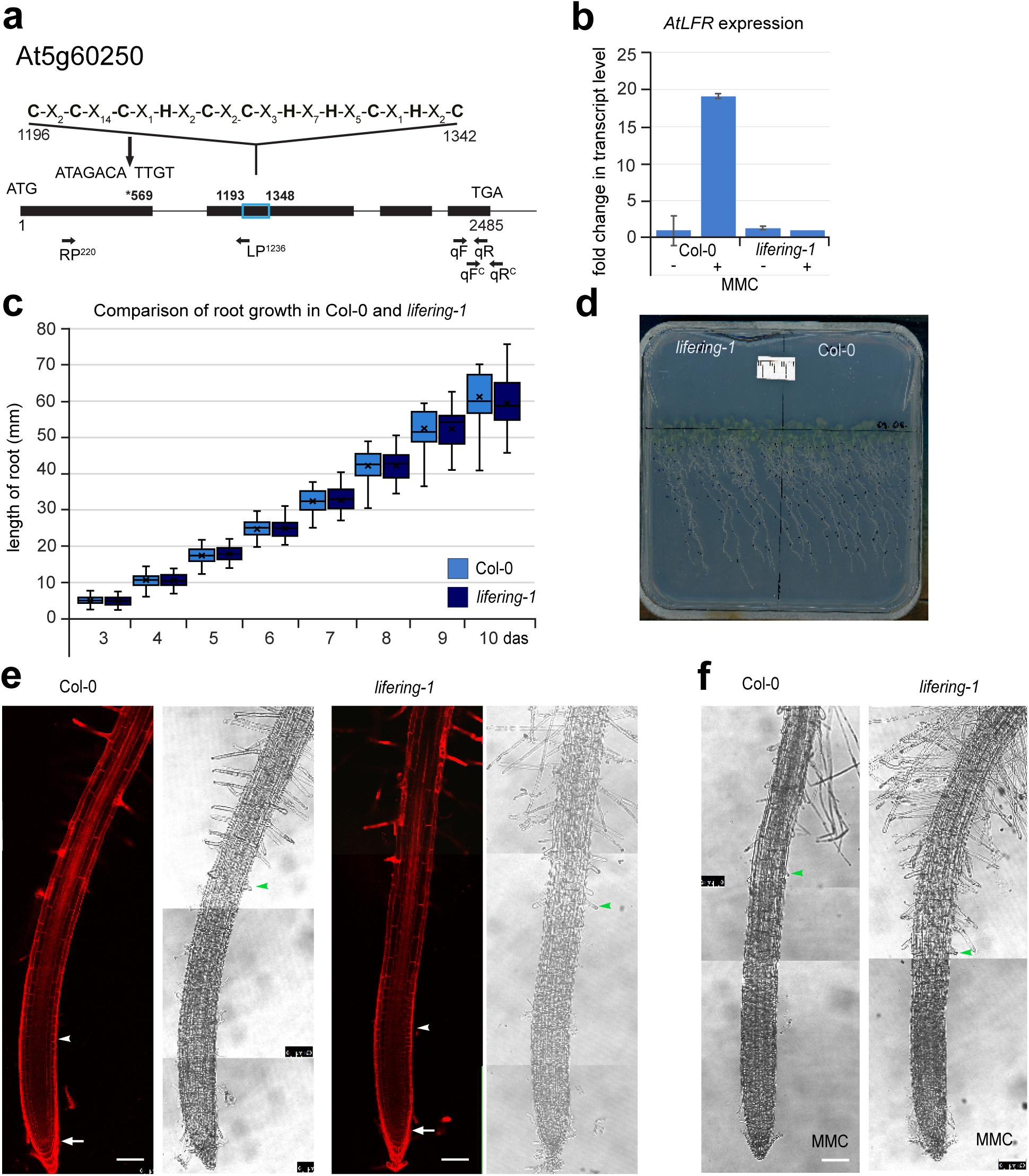
Expressional and phenotypic analysis of the *lifering-1* mutant. (**a**) The position of the insertion (*569) in the first exon of the coding region of the At5g60250 gene (*LIFERING*) in the insertion line, SM_3_34488 is shown. The RING-Ubox is labelled by the blue rectangle at position 1193-1348, the corresponding amino-acid sequence represents the C3HC4 RING-Between-RING finger domain at positions 1196-1342 of the genomic sequence. The primer pairs used for genotyping and expression analysis are located between positions: RP-LP: 220-1236 and qF-qR: 2304-2485, while for controlling the presence of the innate transcript in the lines for complementation, *lifering-1*(pgLIFERING): qF^C^-qR^C^ 2446-2548. See Table S1 for primer sequences and Figure S2 for the complementation of the mutant. (**b**) The level of *LIFERING* does not change upon induction with MMC treatment (- vs + MMC) in the mutant *lifering-1*, indicating that it is a full knock-out mutant. In contrast, in Col-0, upon MMC treatment, around 20 fold induction was detected. MMC treatment +/−16 h, 10μg ml^-1^, *n*=2. (**c**) Quantification of the root growth of Col-0 and *lifering-1* between 3 and 10 das. On a given day, no significant difference was detected between wild-type and mutant. (**d**) The image corresponds to the root growth depicted in (**c**). (**e**) The corresponding PI-stained confocal and differential interference contrast (DIC) microscopy images illustrate the root zonation of Col-0 and *lifering-1* mutant. The white arrows point to the position of QC, while the white arrowheads mark the start of the transition zone towards the elongation zone. (**f**) DIC images show the effect of MMC treatment (16-18h, 10μg ml^-1^), on differentiation in Col-0 and *lifering-1.* In both (**e** and **f**), green arrowheads label the first differentiated epidermis cell. Scale bar: 100μm.

**Figure S2.**
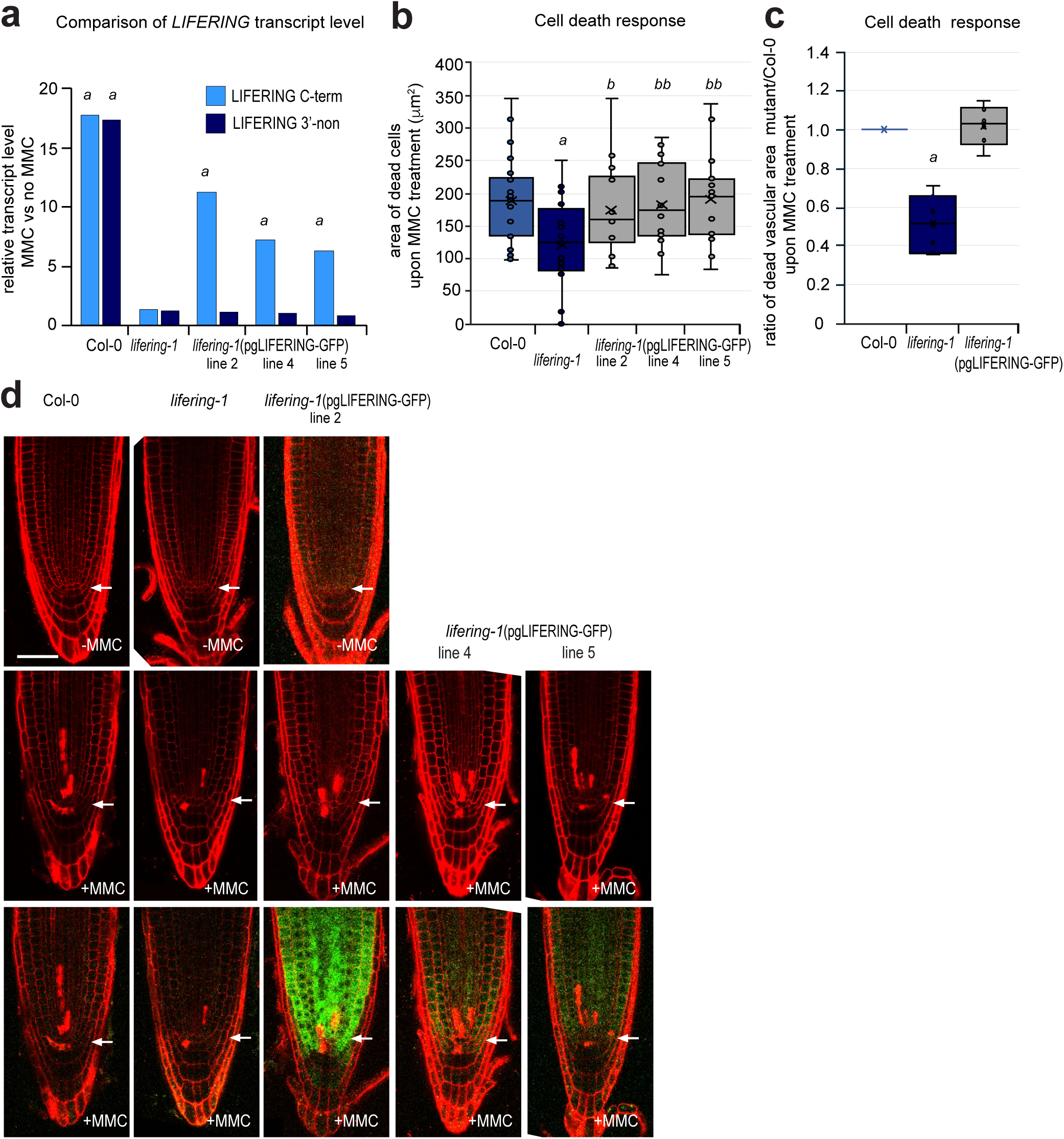
Introducing the *LIFERING* gene under its own promoter can complement the cell death response-phenotype of the *lifering-1* mutant. (**a**) The relative expression of *LIFERING* upon MMC treatment (10mg ml^-1^) after 16 h. qRT-PCR was carried out on Col-0, *lifering-1* and *lifering-1*(pgLIFERING:GFP) lines (line 2, 4 and 5) on 5-6 das seedlings. Primer combinations, were located to the C-terminal (C-term) and to the 3’ non-translated region (3’-non). Position of the primer pairs is indicated in Figure S1A. The values represent fold changes normalised to the values of the *LIFERING* gene in the non-treated samples where the expression was taken as 1. (**b**, **c**) Comparison of the cell death response upon MMC treatment (10mg ml^-1^, 16 h, 5-6das) in the *lifering-1* mutant and the complemented lines *lifering-1*(pgLIFERING:GFP).The cell death response was quantified by measuring the area of dead cells in the proximal meristem (mm^2^, PI-stained area). (**b**) The graph shows the combined measurement of *n*=3, *N*>10 in each experiment. In the box-plot analysis the median was also included. *a*: p<0.002 significance comparing the cell death response between *lifering-1* and Col-0, *b*: p<0.05, *bb*:p<0.001 between the introgressed lines and the mutant using Student’s *t*-test analysis. (**c**) illustrates ratio of cell death response of Col-0, *lifering-1* and *lifering-1*(pgLIFERING:GFP) where the measurements were combined for the three complemented lines. (**d**) Confocal microscopy (CM) images of PI-stained root-tips of Col-0, *lifering-1* and *lifering-1*(pgLIFERING:GFP) lines 2, 4 and 5. Seedlings were grown without (-MMC) and treated with MMC (+MMC) for 16 h. To compare the level of *LIFERING* gene induction, see the two sets of images below, red channel shows the cell death response, while red and green channels combined, the location of the signal. For the microscopic analysis same settings were used through out the entire experiment. Scale bar: 50μm, white arrow: QC position in each image.

**Figure S3.**
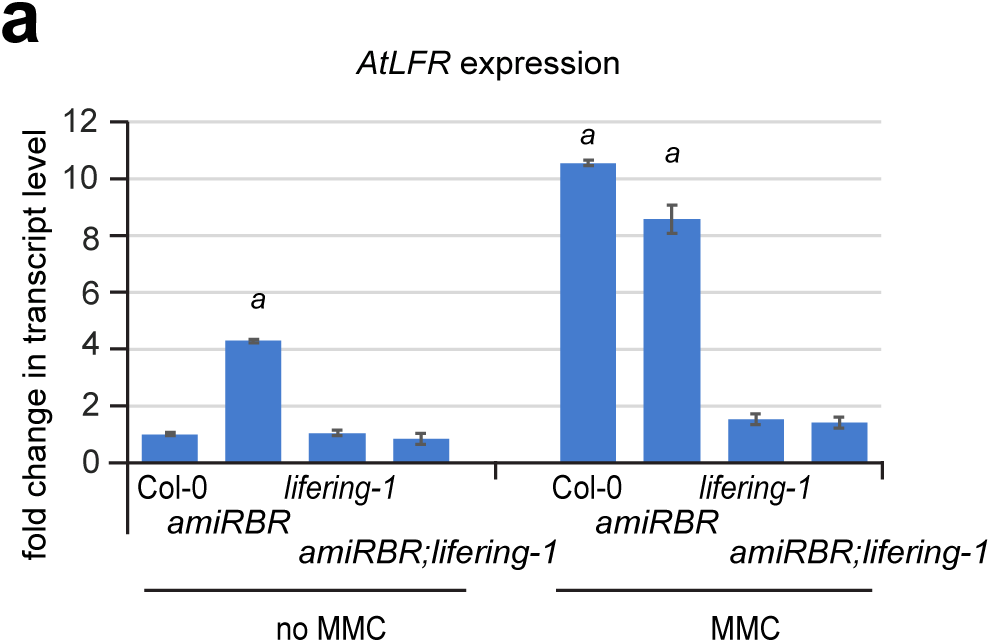
*LIFERING* expression in the double *amiRBR;lifering-1* compared to the single *amiRBR* and *lifering-1* mutants. (**a**) Comparison of the level of *LIFERING* expression in Col-0, in the single *amiRBR* and *lifering-1* mutants and in the double *amiRBR;lifering-1* mutant in the absence and presence of the genotoxic agent, MMC. No significant change was detected upon induction with MMC treatment in the *amiRBR;lifering-1*, indicating that the double mutant lacks the *LIFERING* transcript. MMC treatment +/−16 h, 10μg ml^-1^, *n*=3, *a*: p<0.01 significant difference between the mutant lines compared to non-treated Col-0 expression, where the level of *LIFERING* was taken as 1.

**Figure S4.**
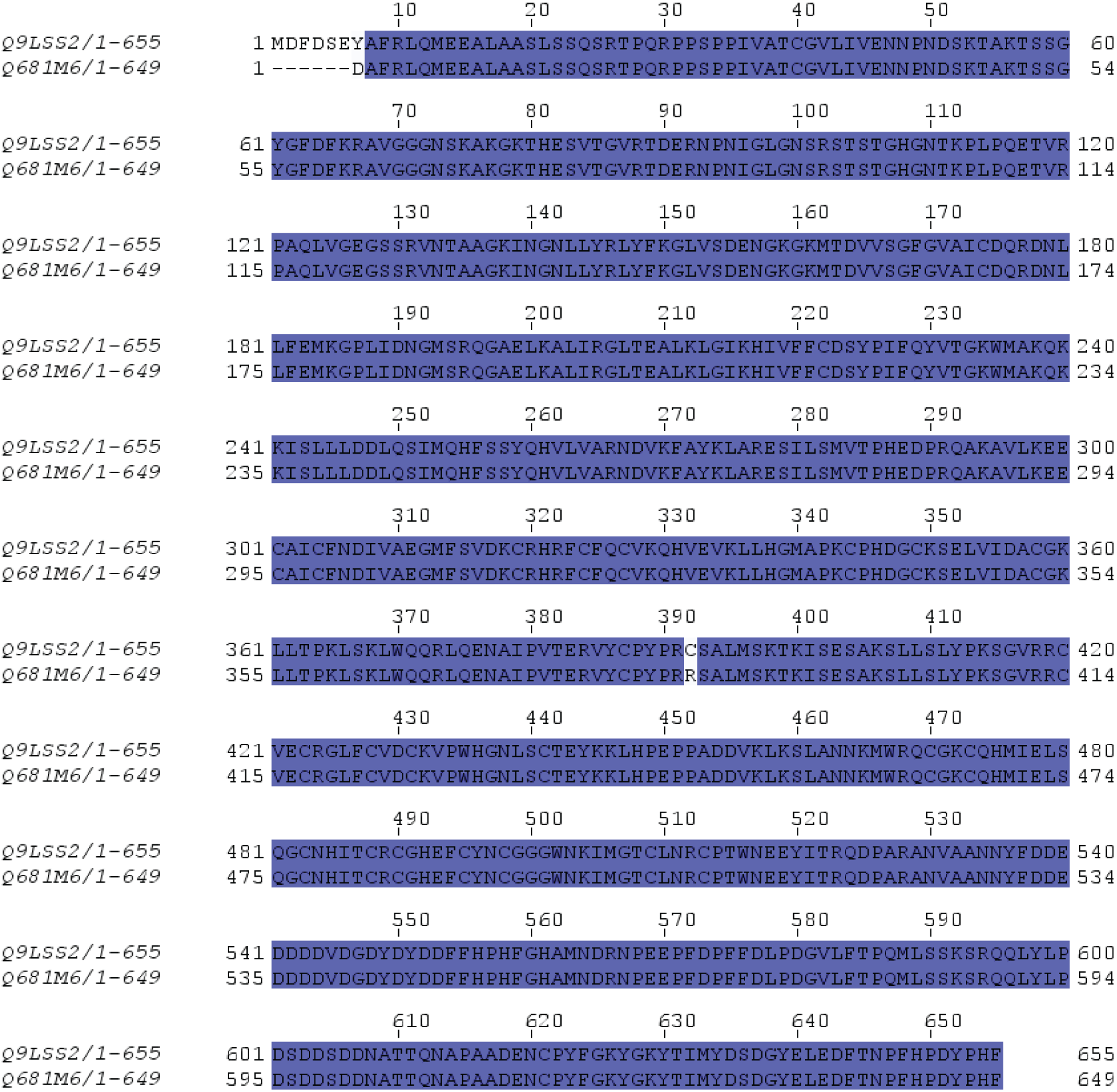
Aminoacid sequence alignment of UniProt Q9LSS2 and Q681M6. Amino acid sequence alignment of UniProt Q9LSS2 genomic sequence-based and Q681M6 transcript-derived sequences, the putative protein products of the *AT5G60250* gene.

**Figure S5.**
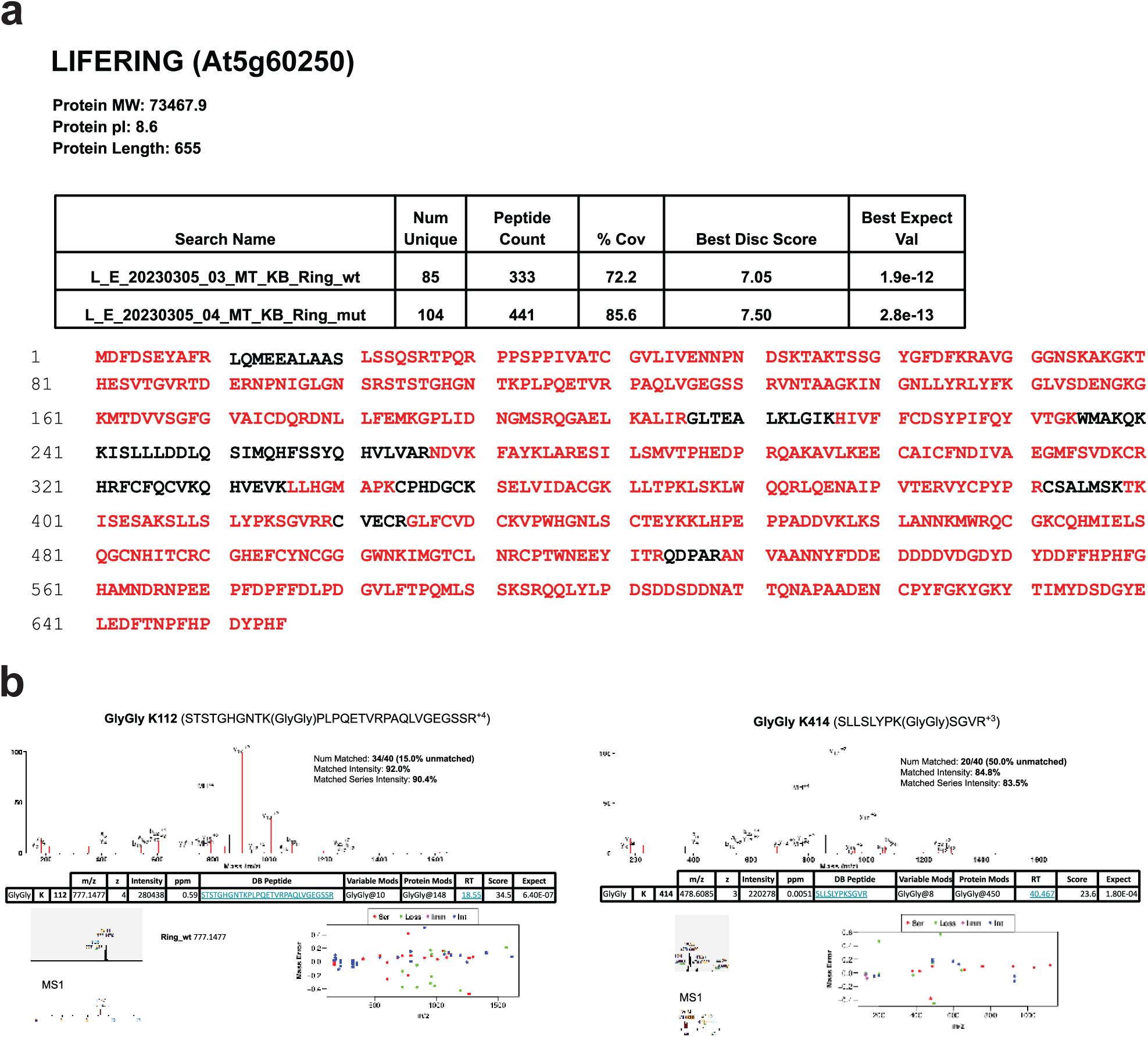
Mass spectrometry analysis of LIFERING (At5g60250) ubiqitination. The N-terminally 12xHis-tagged wild-type and mutant (C392R) LIFERING proteins were expressed using wheat germ extract-based *in vitro* translation. Following the *in vitro* ubiquitination reactions in the presence of FLAG-ubiquitin, the 12xHis-tagged proteins were isolated by immobilized metal-ion affinity chromatography (IMAC), and mass spectrometry was performed. Western blot analysis of the isolated proteins is shown in Fig. 5e. (**a**) the detected peptide fragments of LIFERING (Uniprot: Q9LSS2) labelled in red. (**b**) MS2 spectra of the annotated ubiquitinated peptides. LIFERING is intensively ubiquitinated on the K112 and K414 residues of the wild-type protein, but no MS1 signal, and therefore no ubiquitination, is detectable on the mutant protein.

**Figure S6.**
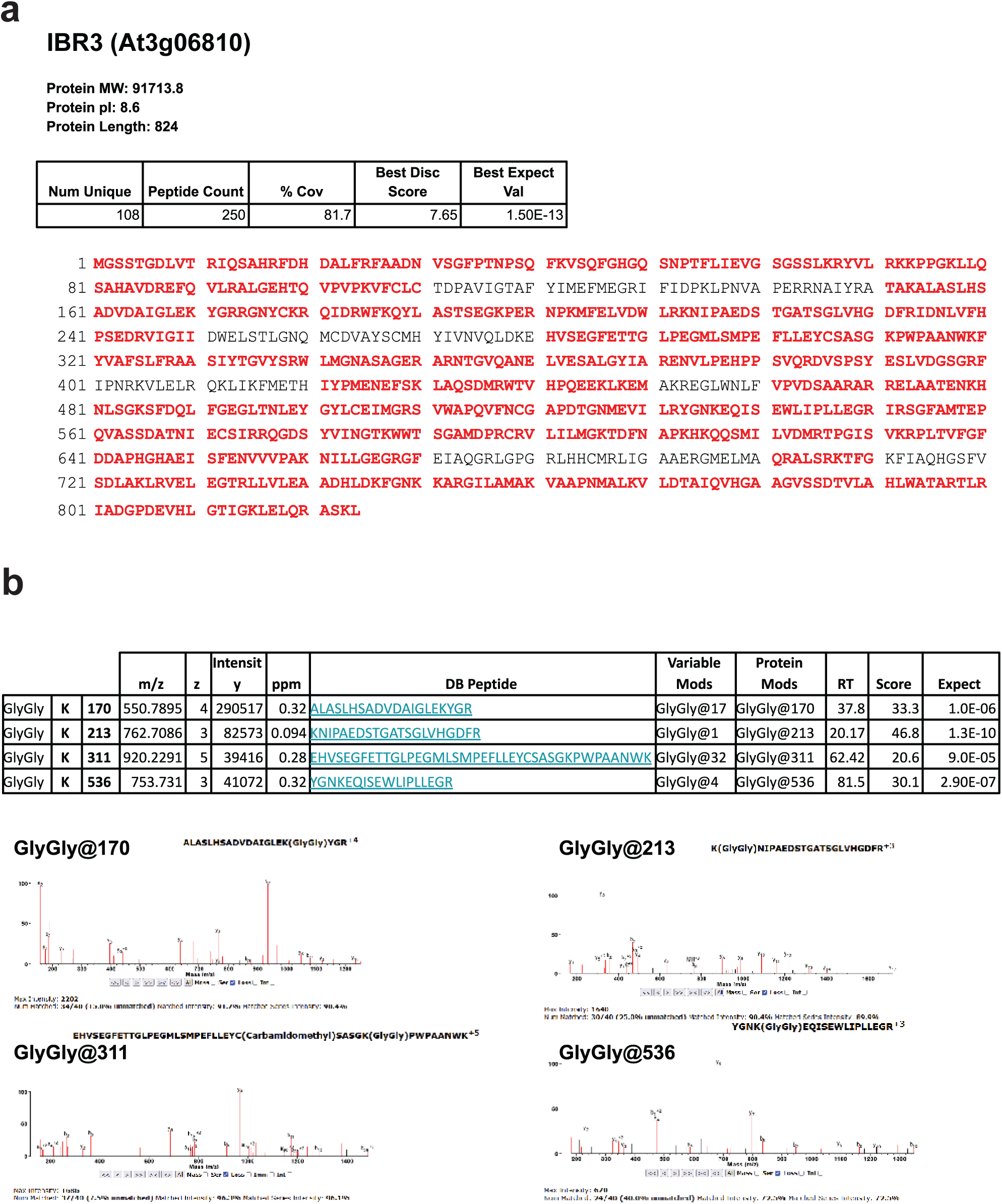
Mass spectrometry analysis of IBR3 (At3g06810) ubiquitination. The acyl-CoA dehydrogenase-like protein, IBR3, labelled with an N-terminal GST tag was expressed using wheat germ extract-based i*n vitro* translation and purified with glutathione beads. The bead-bound IBR3 proteins were subjected to *in vitro* ubiquitination reactions in the presence of total translation mixtures containing wild-type or C392R mutant LIFERING proteins and FLAG-ubiquitin. The bead-bound proteins were analysed by mass spectrometry. Western blot analysis of the isolated proteins is shown in Fig. 6a. **(a)** The detected peptide fragments of IBR3 (Uniprot: Q8RWZ3) labelled in red. **(b)** MS2 spectra of the annotated ubiquitinated peptides. IBR3 is ubiquitinated on the K170, K213, K311, and K536 residues exclusively by the wild-type LIFERING.

**Figure S7.**
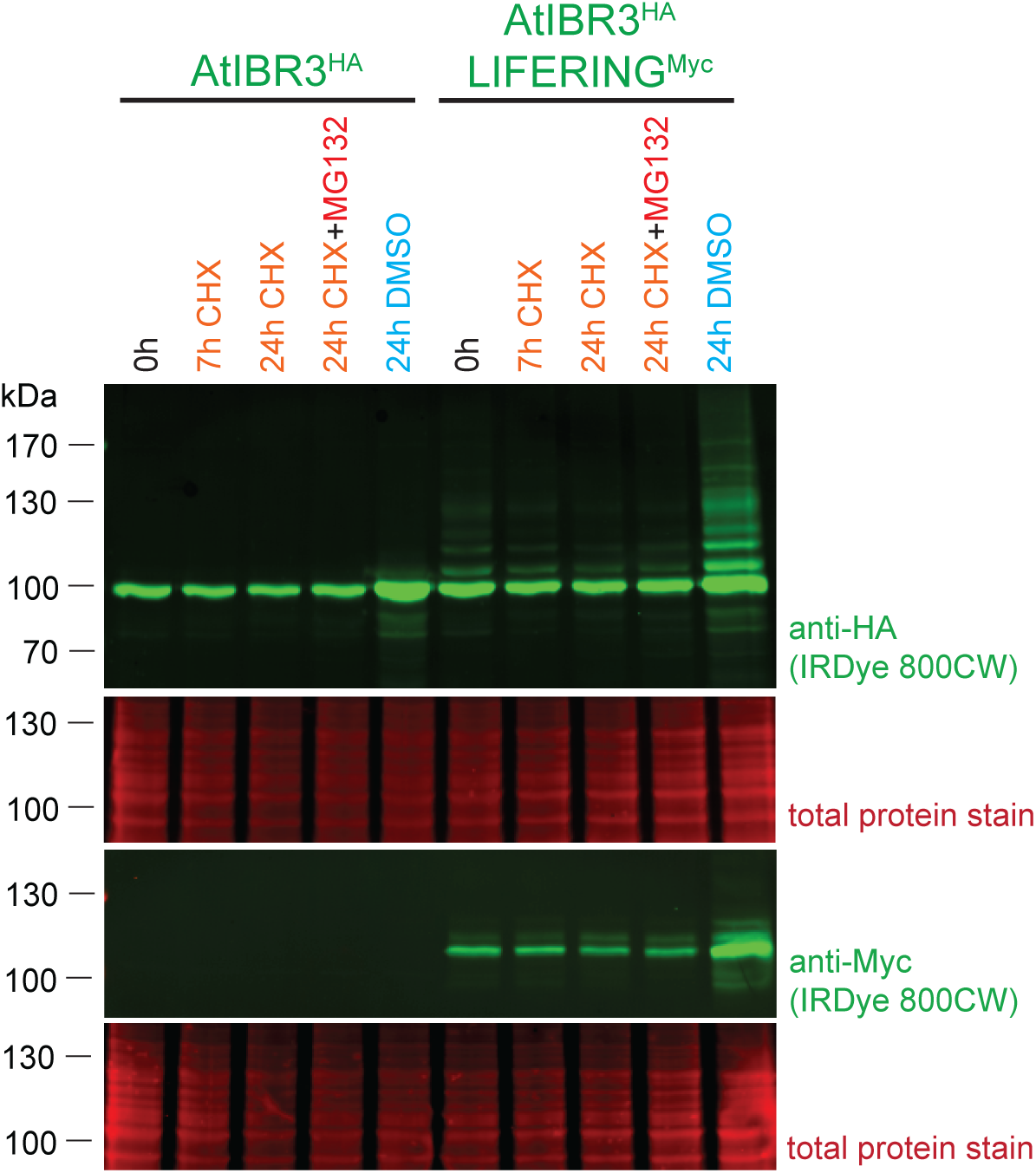
Effect of cycloheximide (CHX) and MG132 on the amount of AtIBR3 protein in *Arabidopsis* root cell suspension protoplasts. Western blot analysis of *Arabidopsis* root cell suspension protoplasts transfected with HA-tagged IBR3 or co-transfected with wild-type Myc-tagged AtLIFERING and HA-tagged AtIBR3-encoding vectors. Protoplasts were treated with 50 μM CHX and 50 μM MG132 for the indicated period is of time. Singleplex detection was performed using mouse anti-HA and mouse anti-Myc, and the membranes were stained with Revert 700 Total Protein Stain to check for loading consistency across the wells. The secondary antibody used was IRDye 800CW goat anti-mouse conjugate (green).

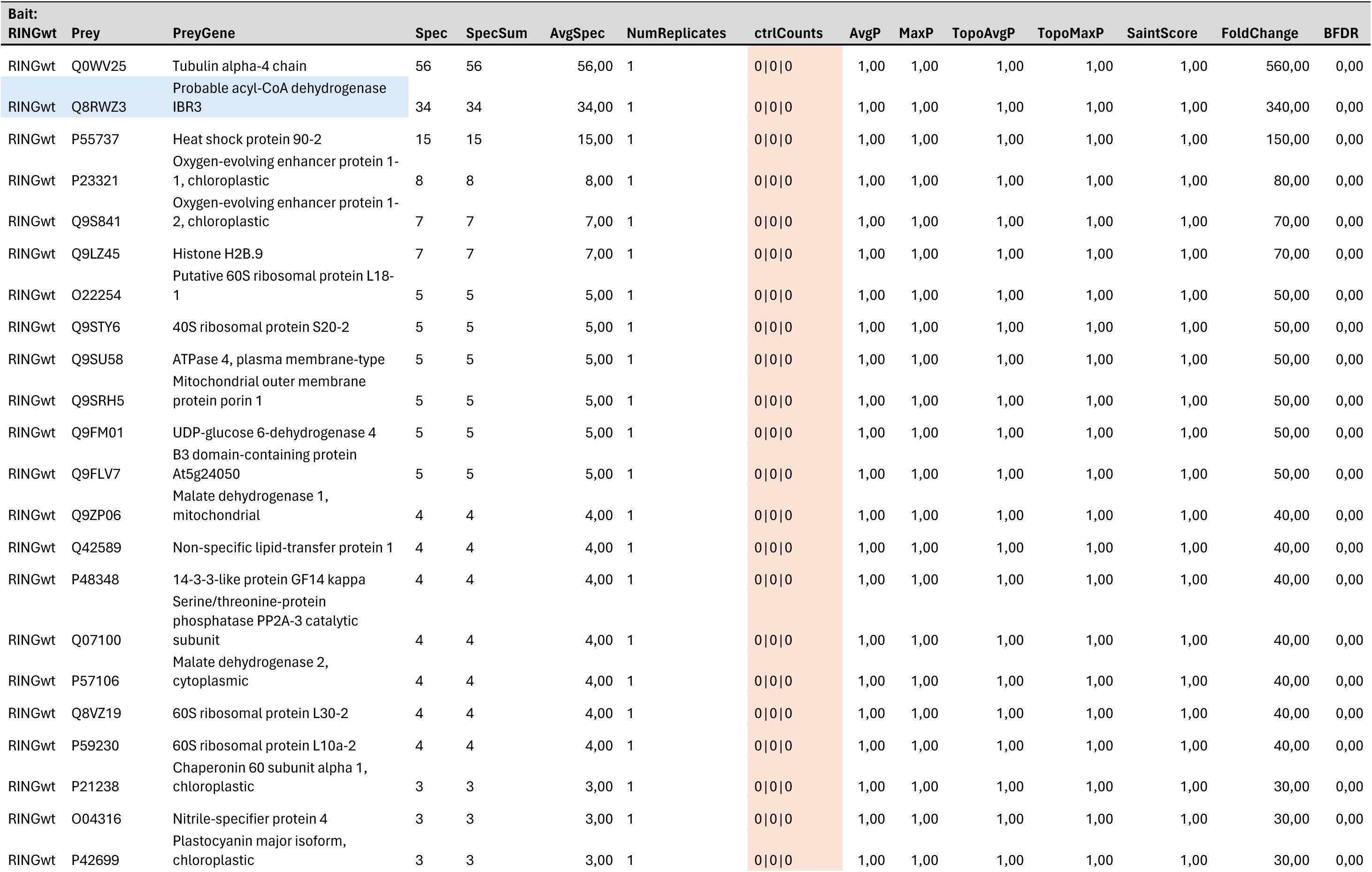
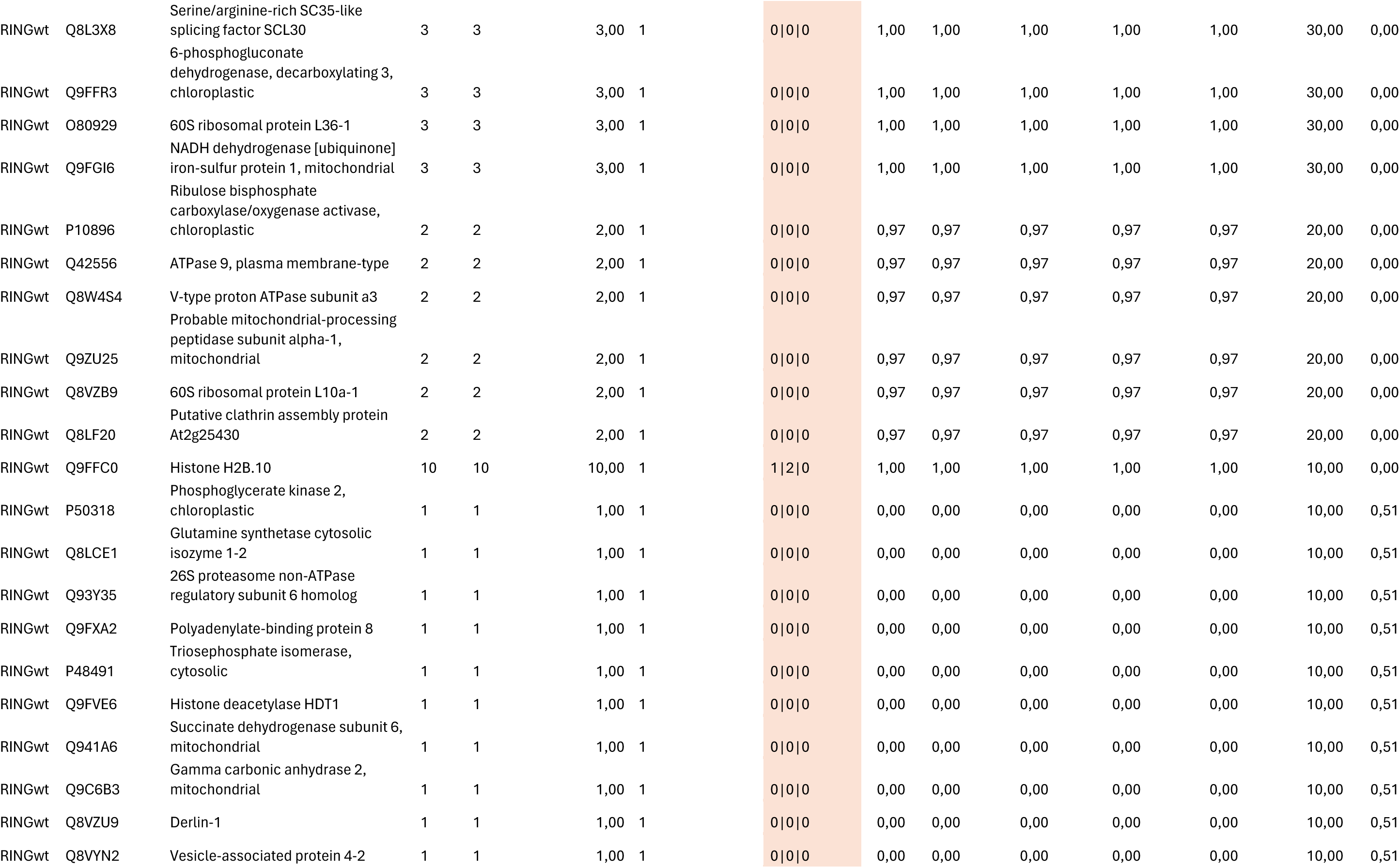
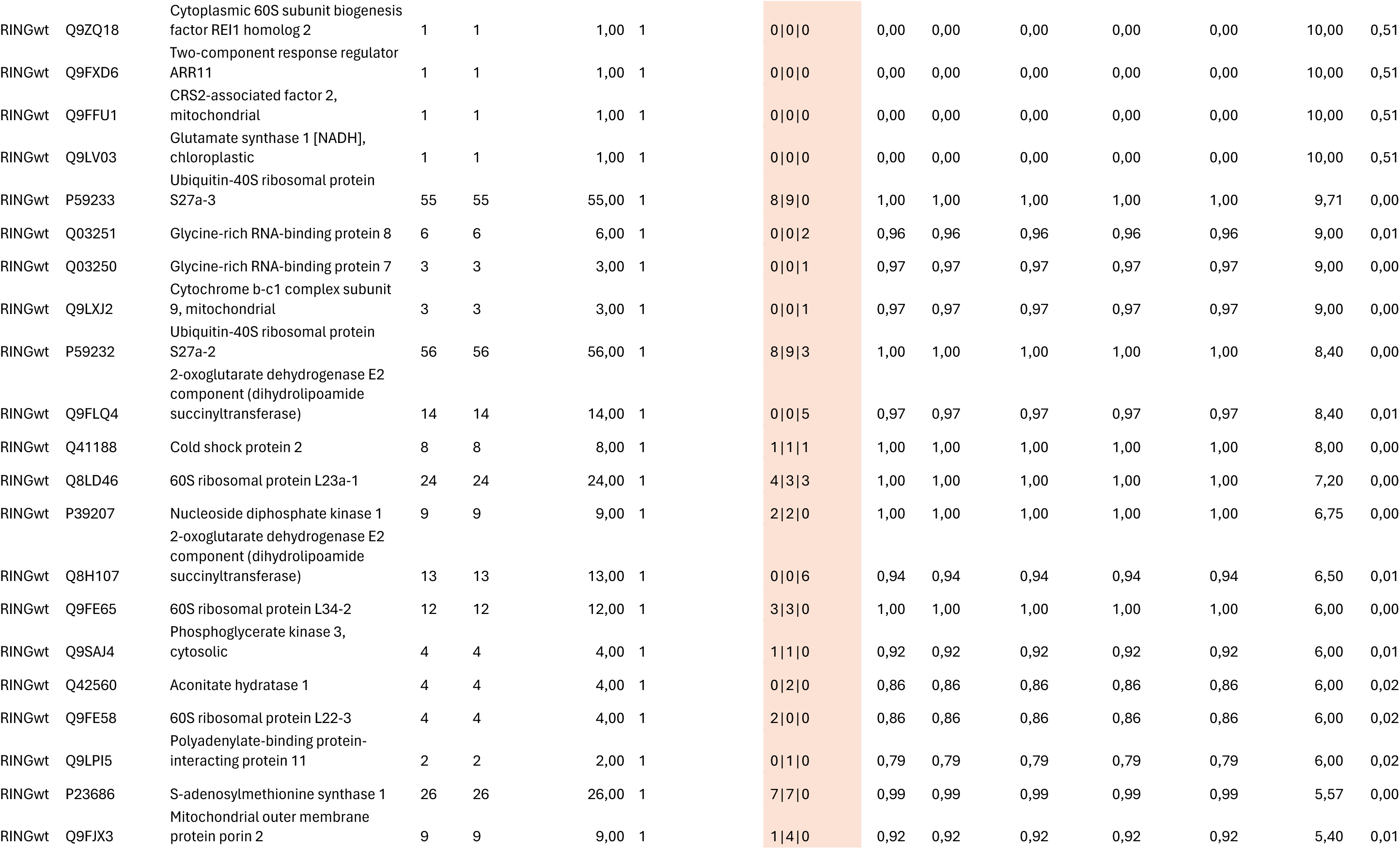
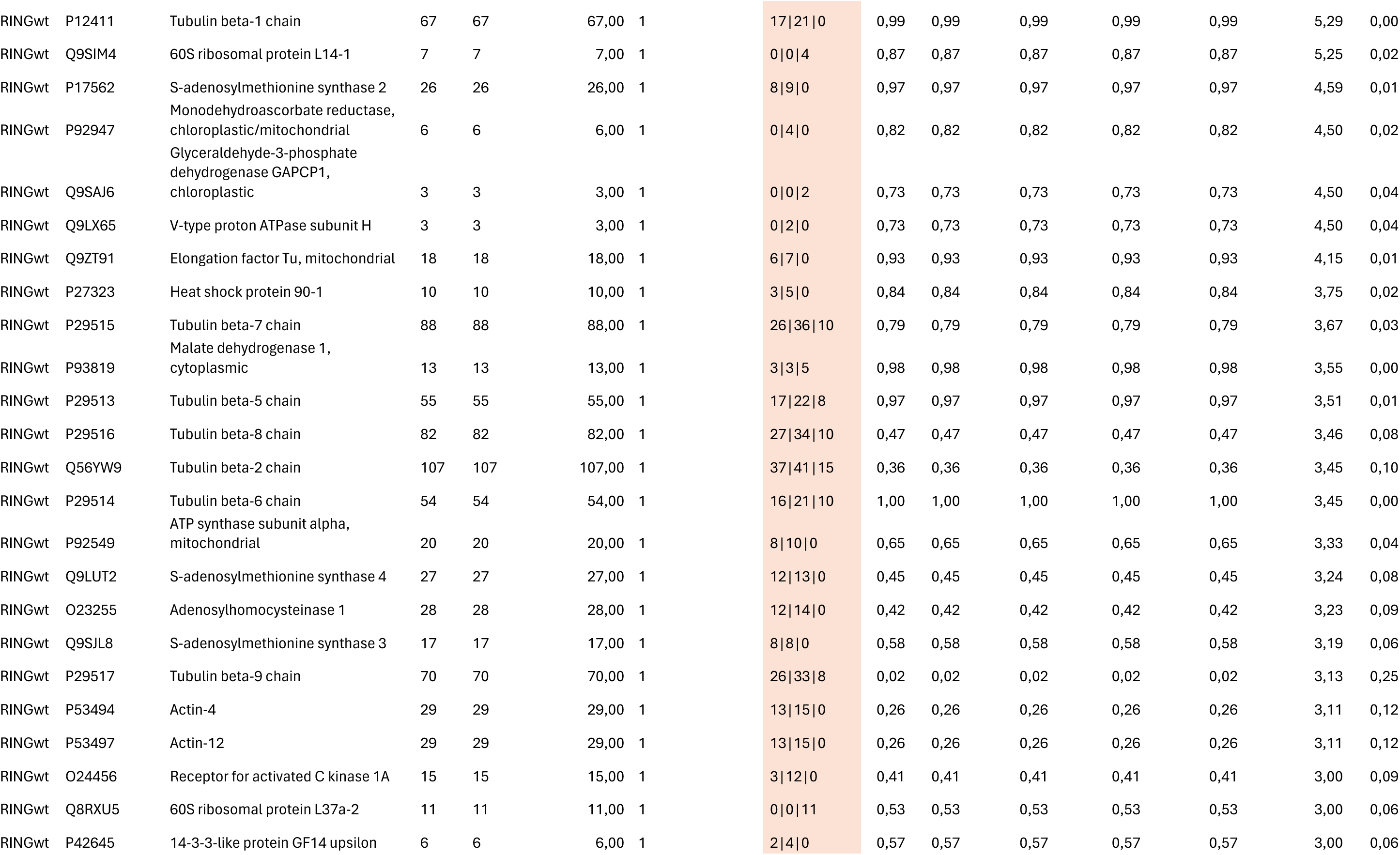
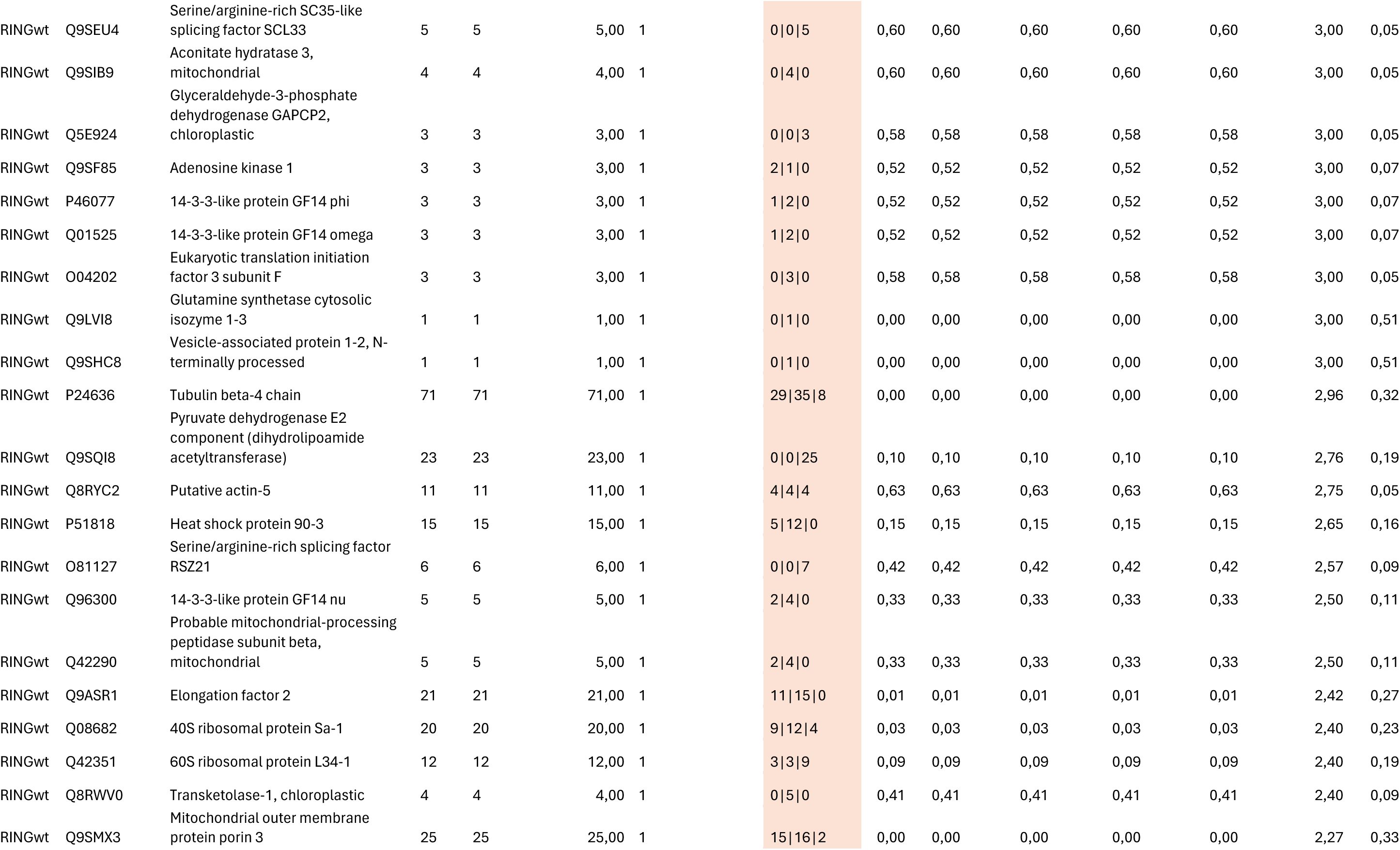
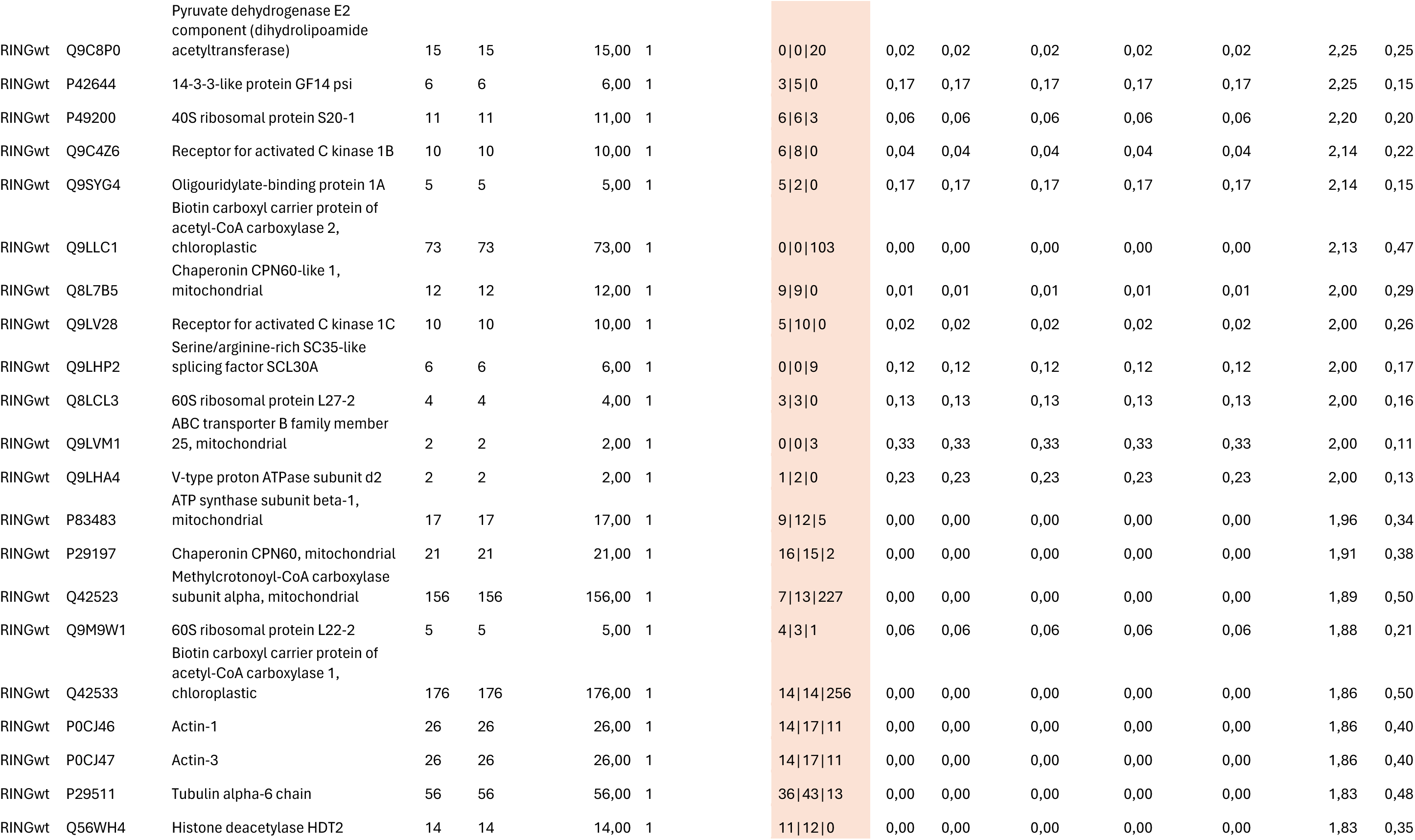
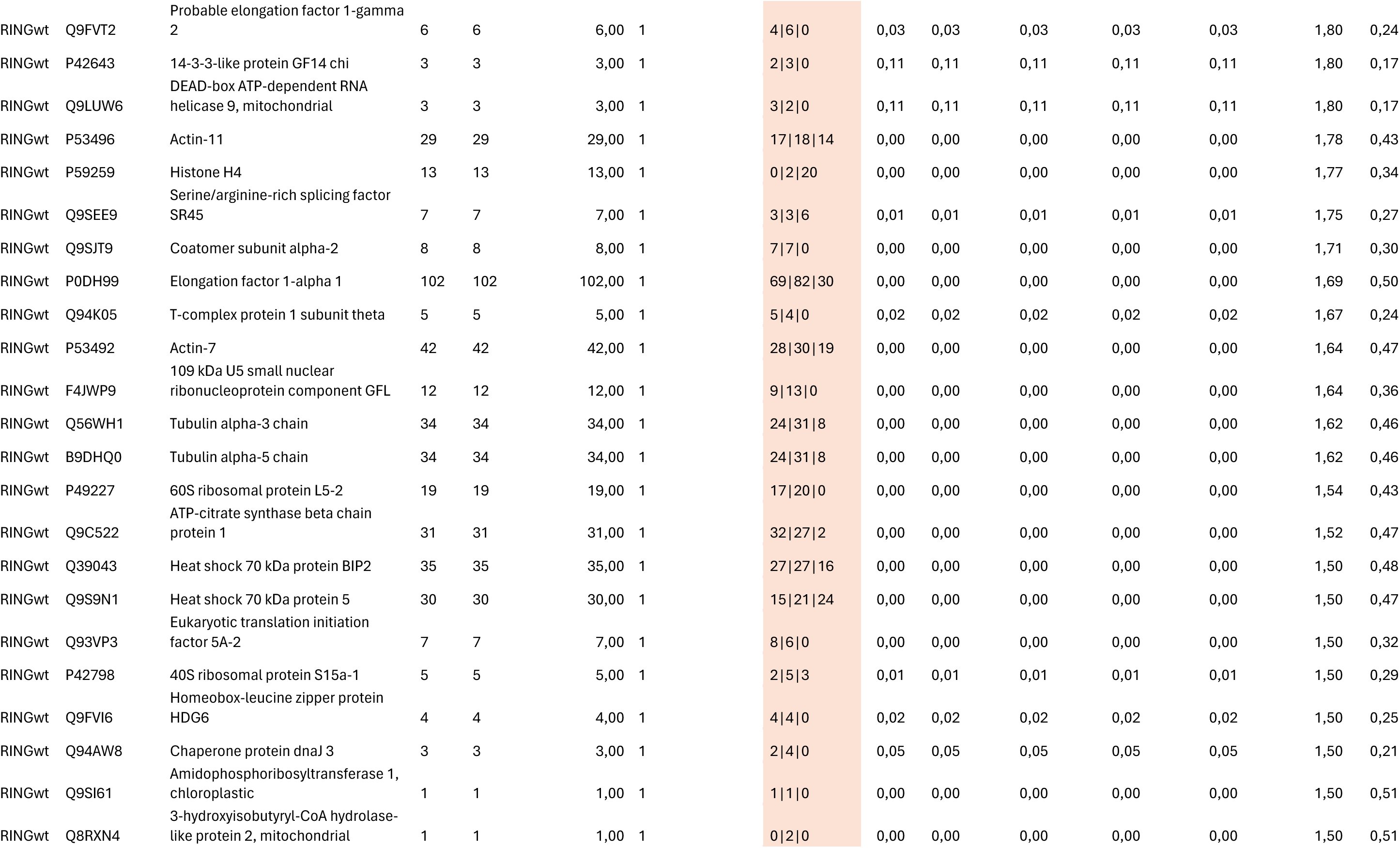
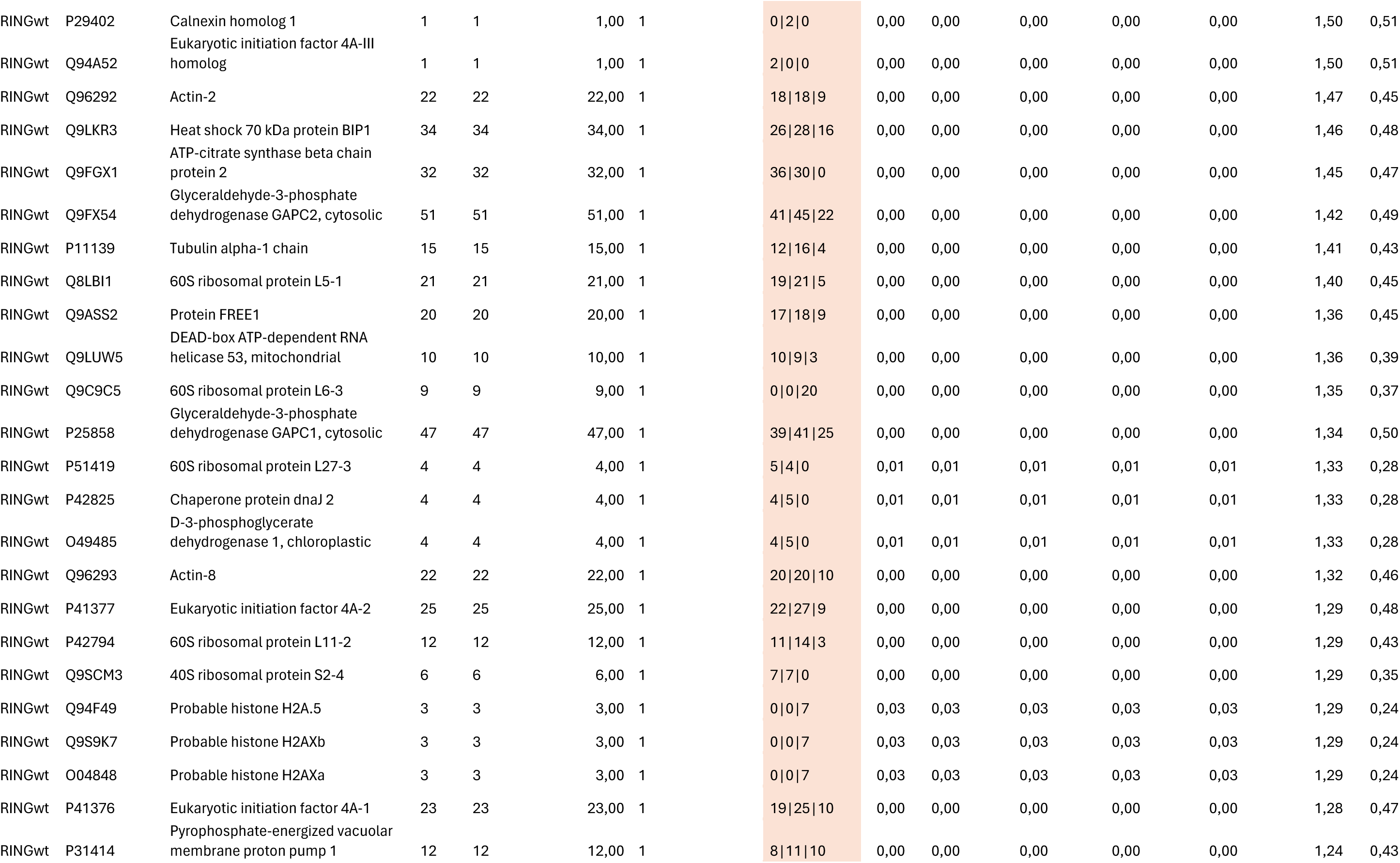
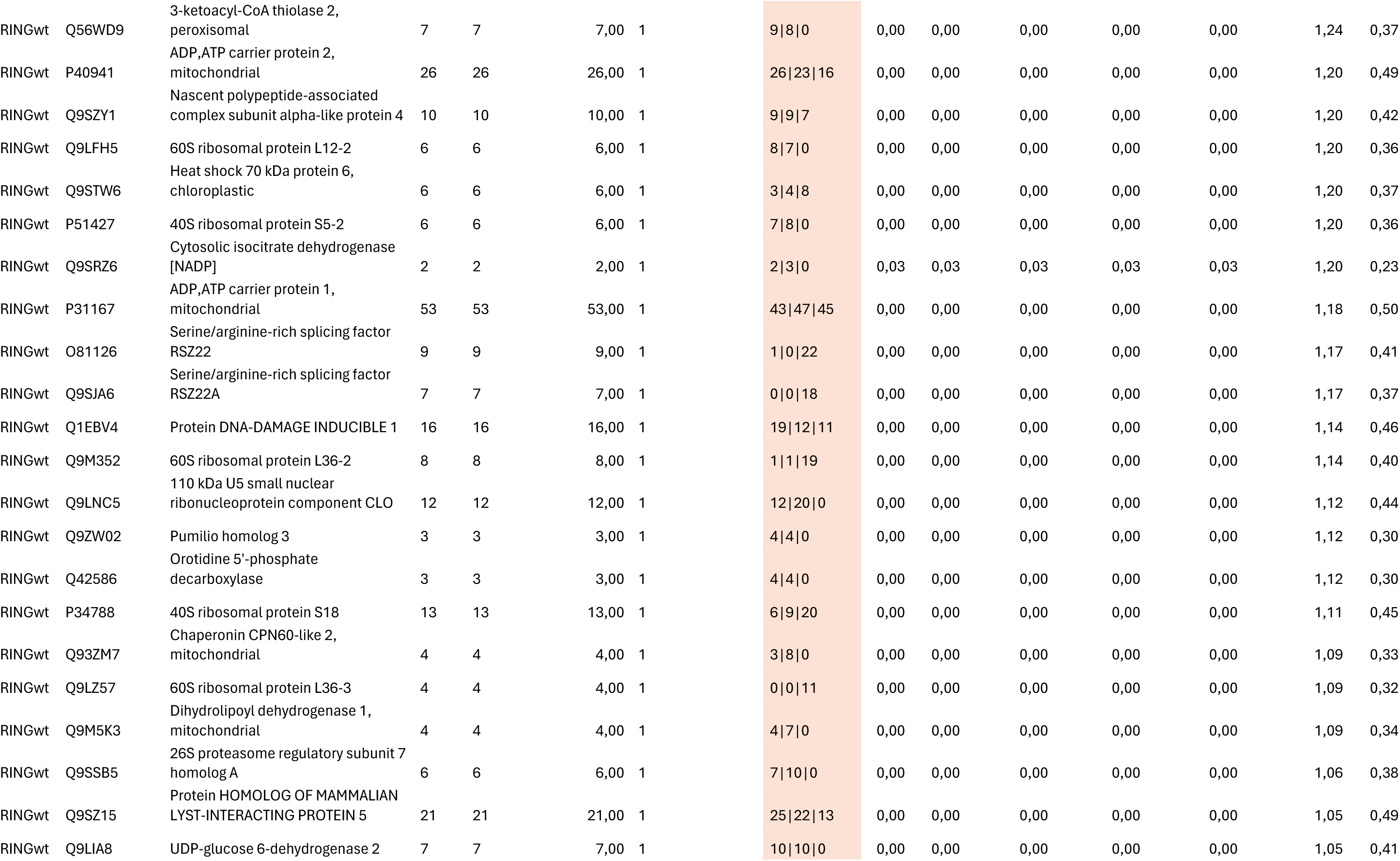
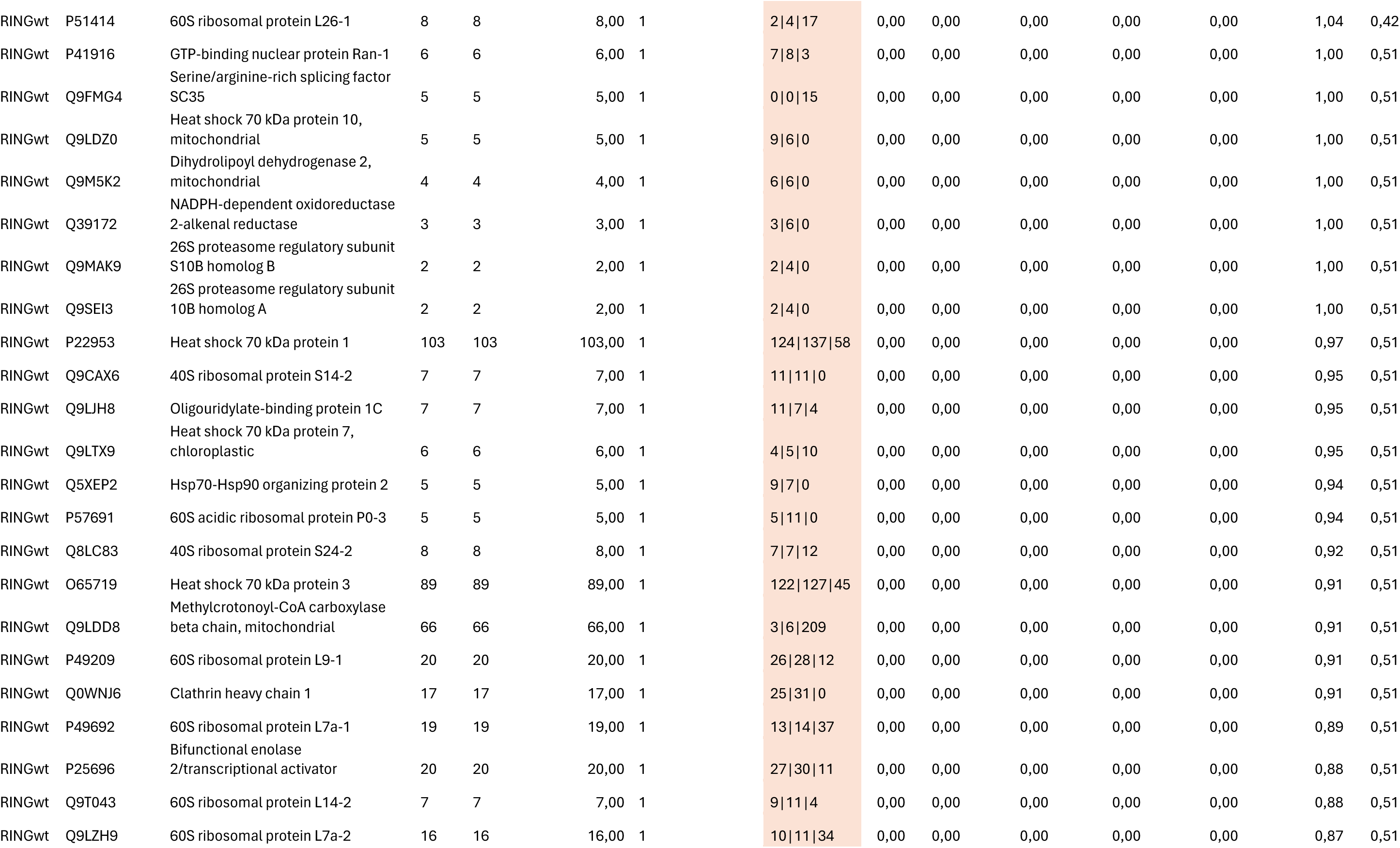
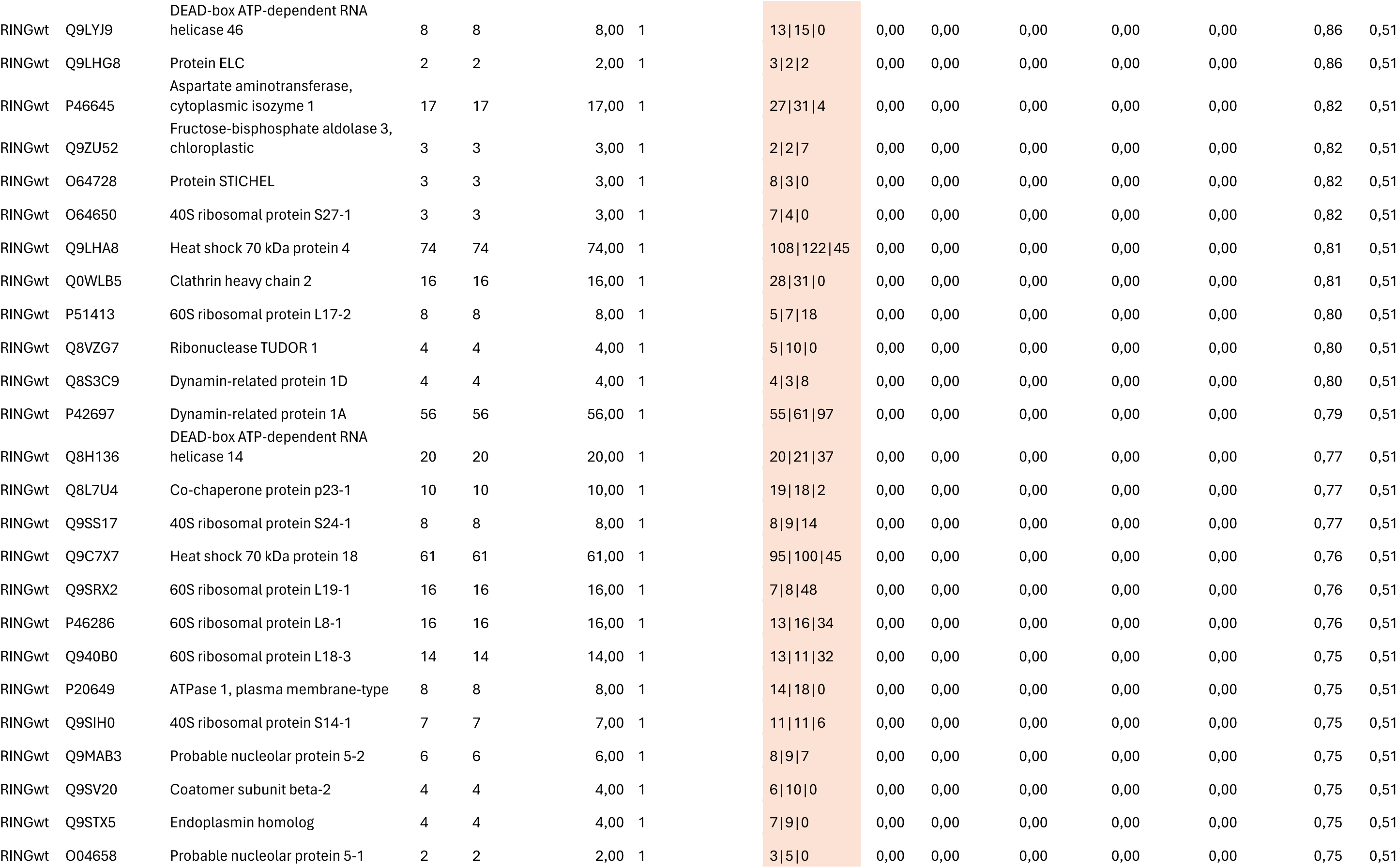
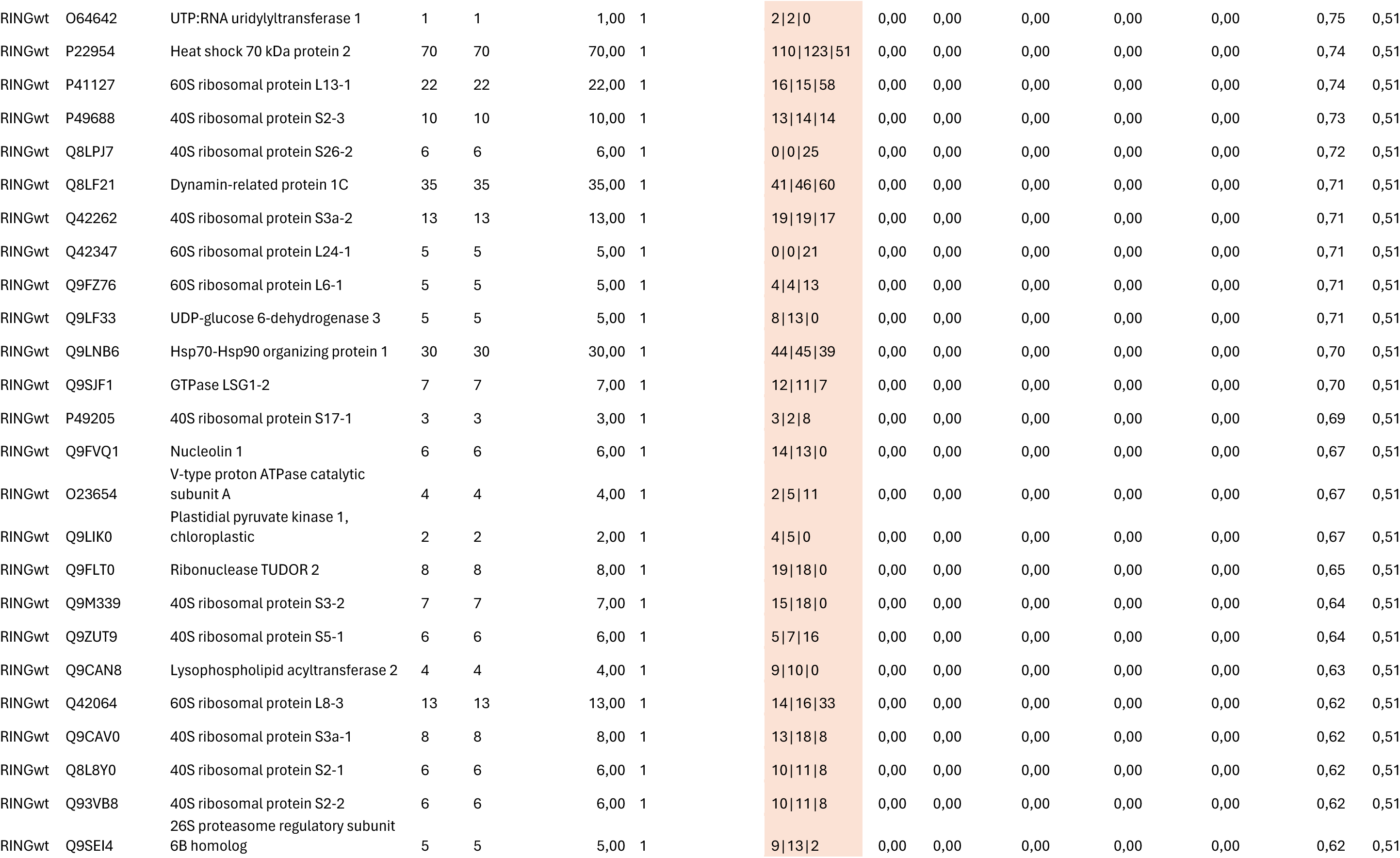
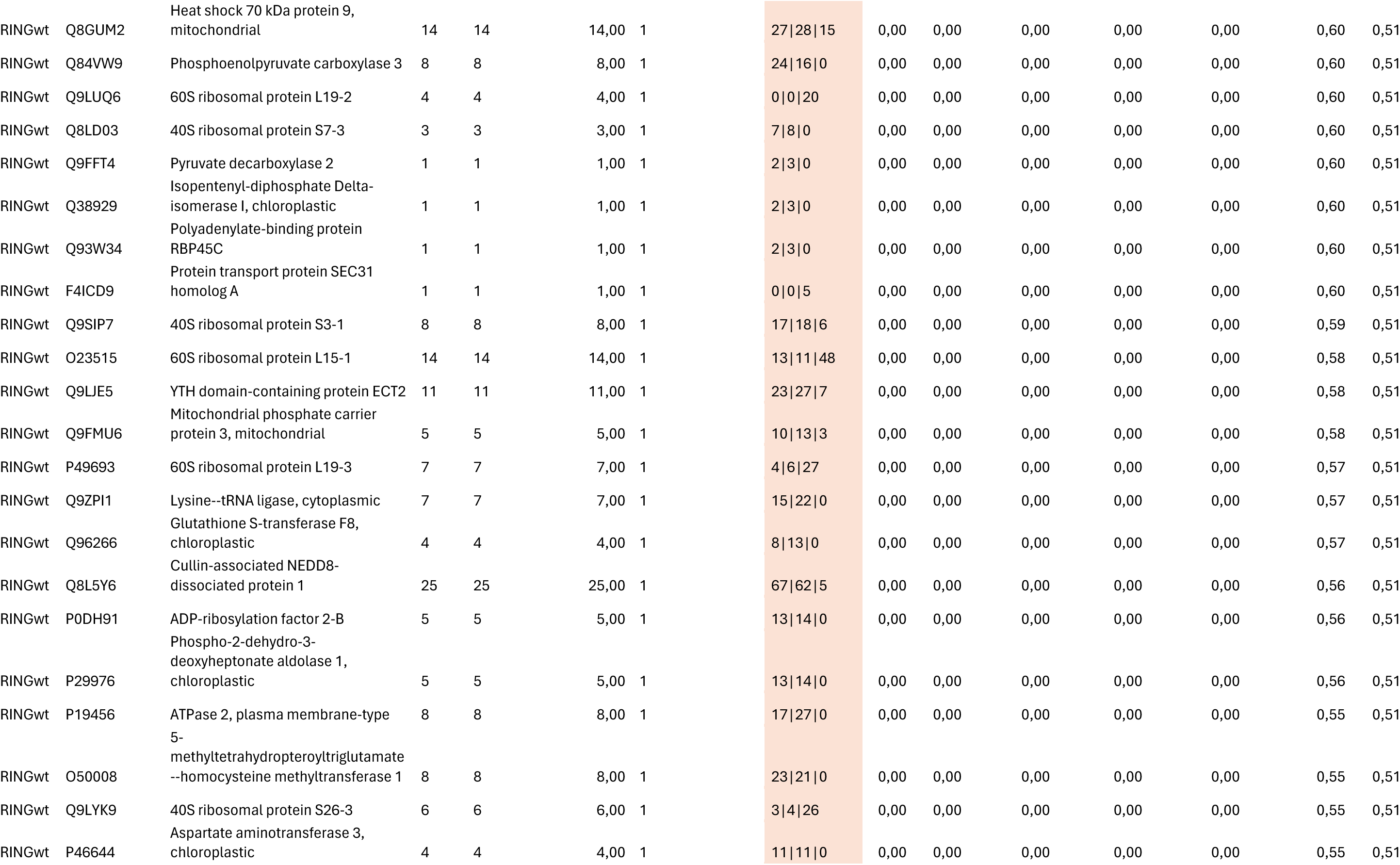
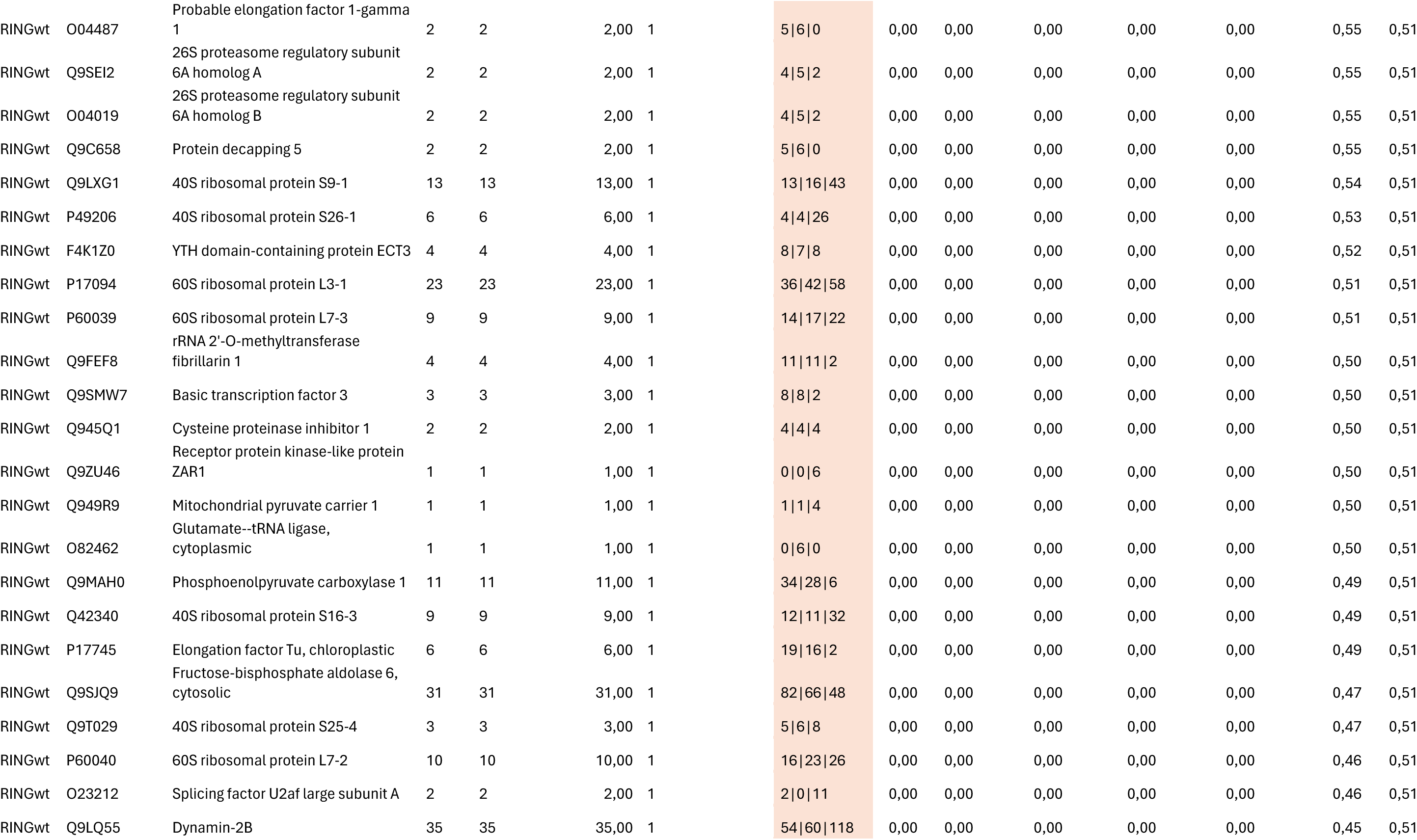
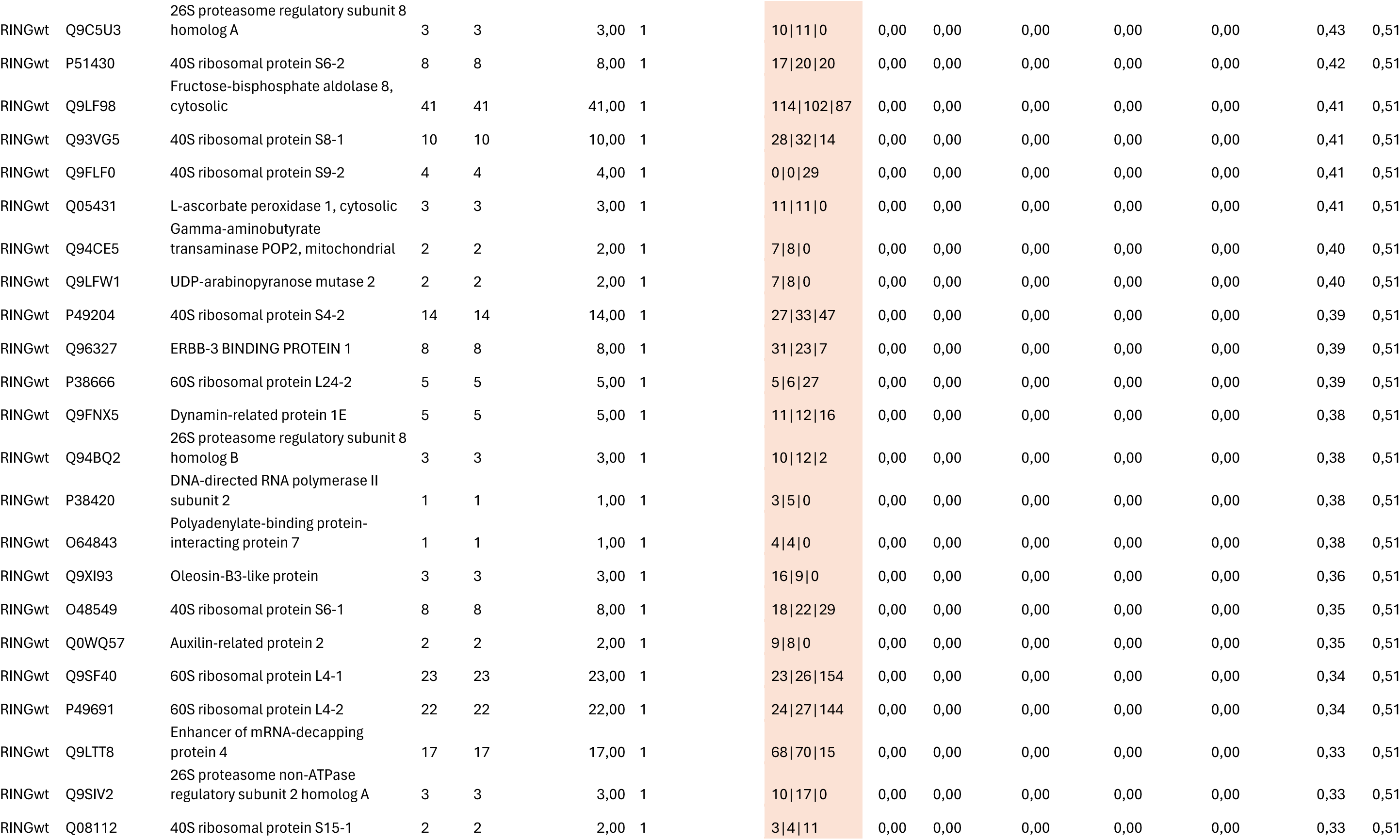
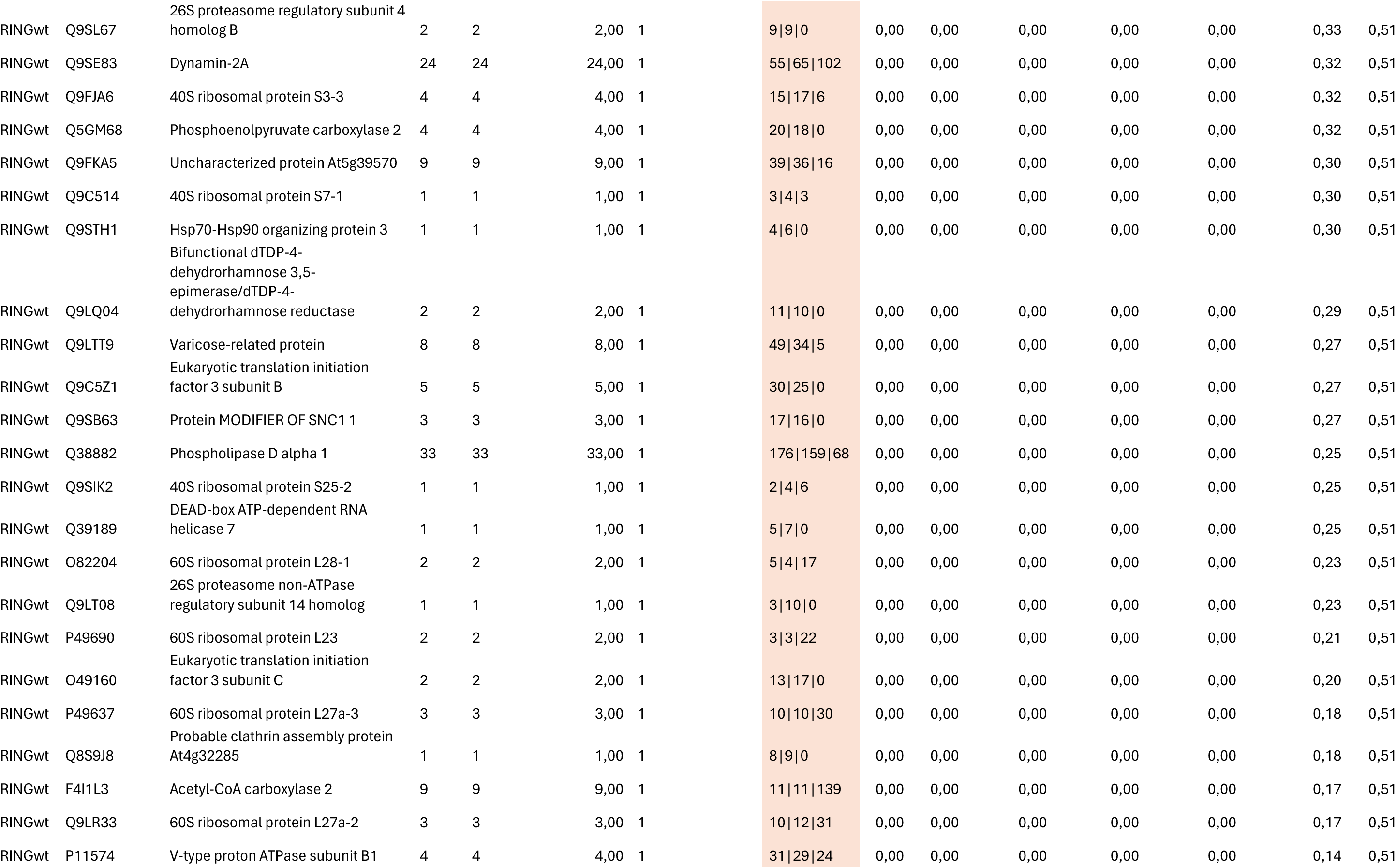
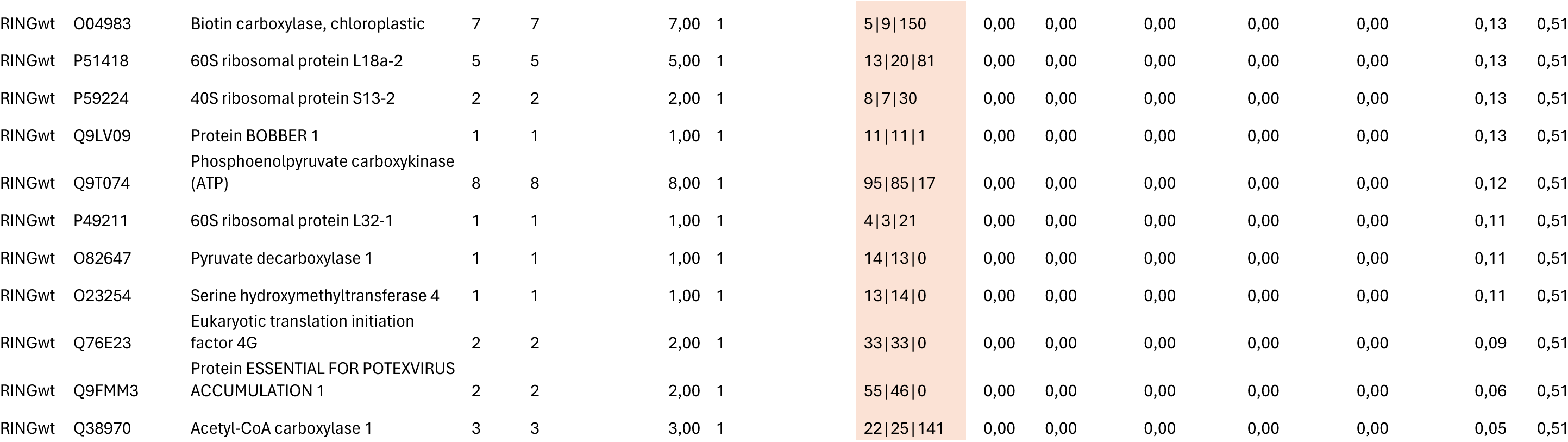
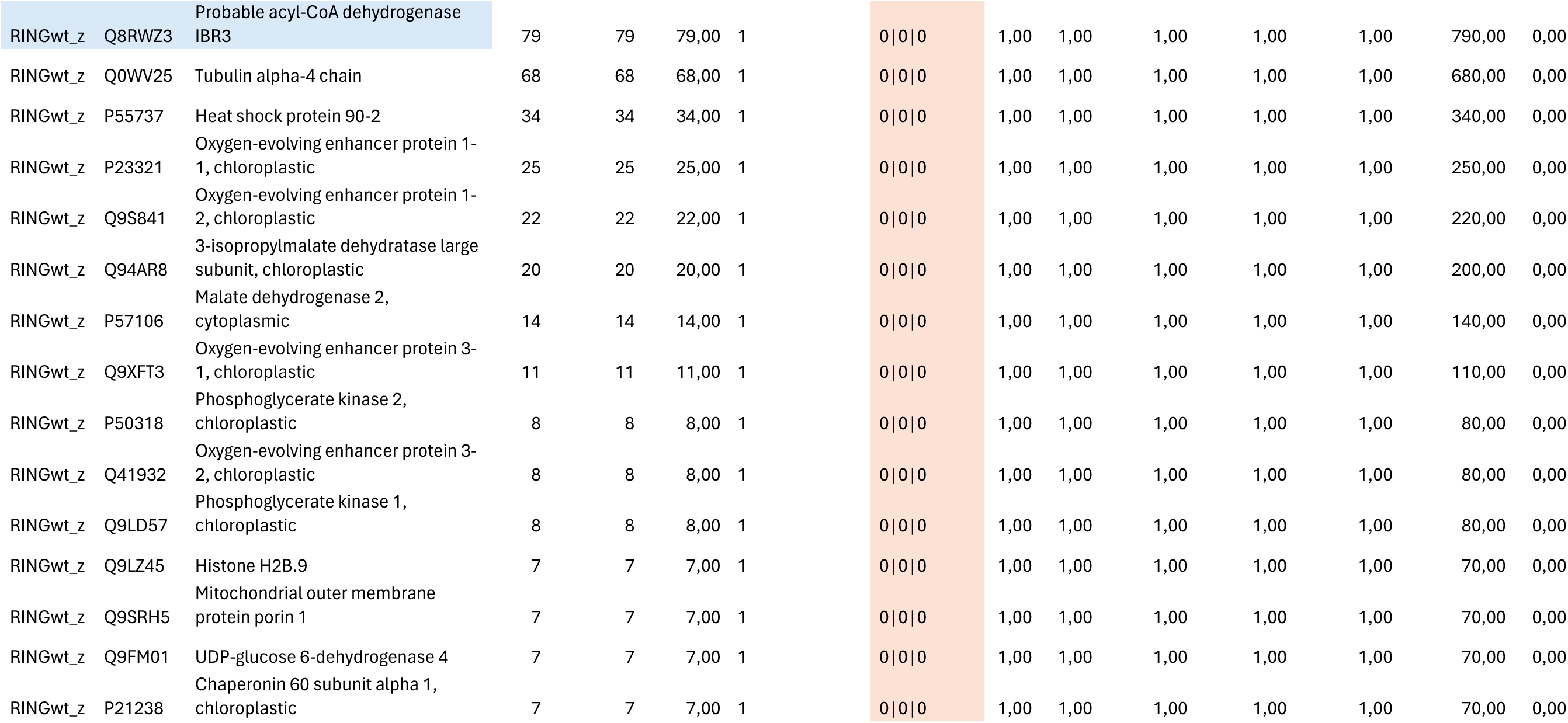
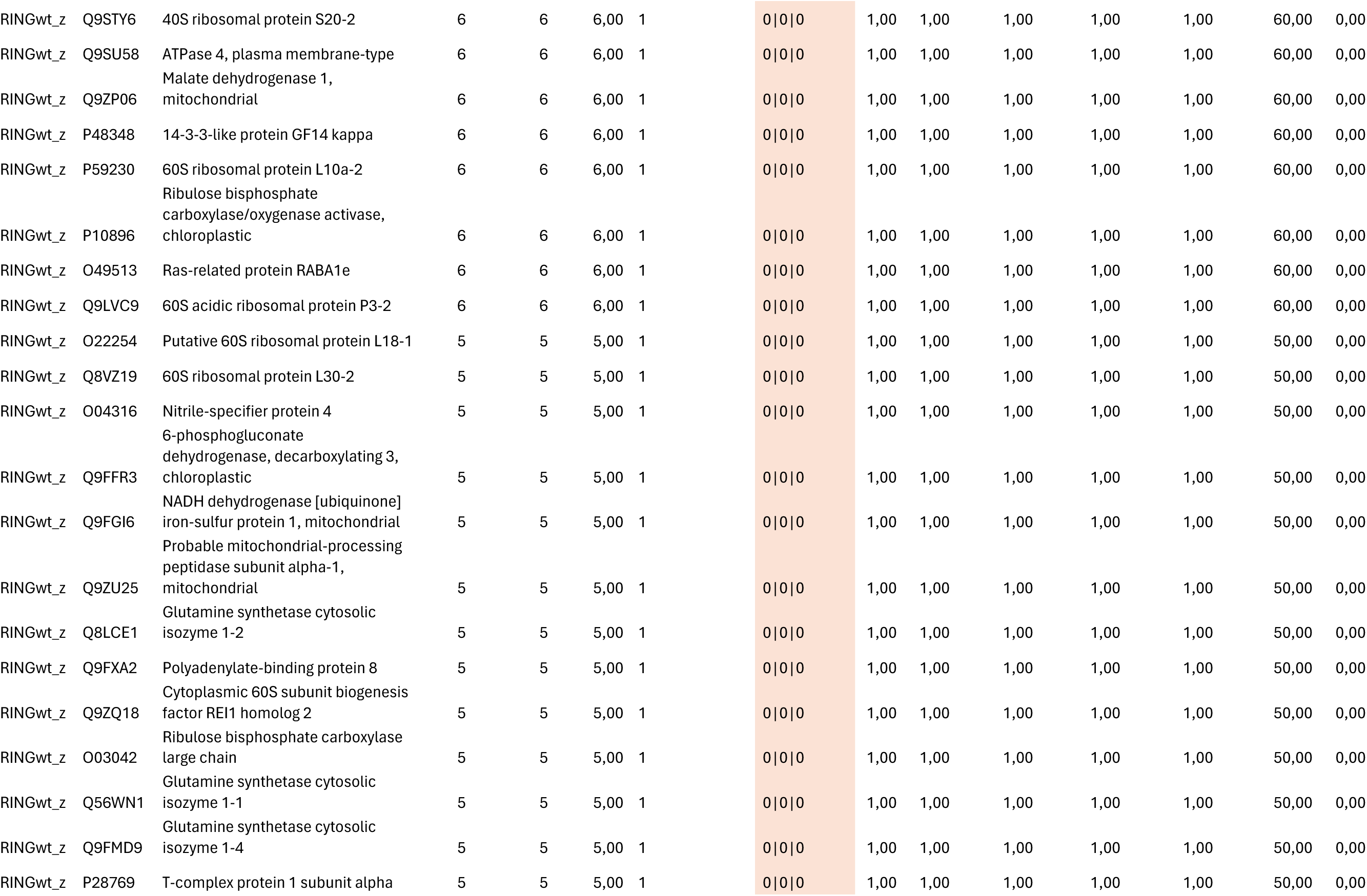
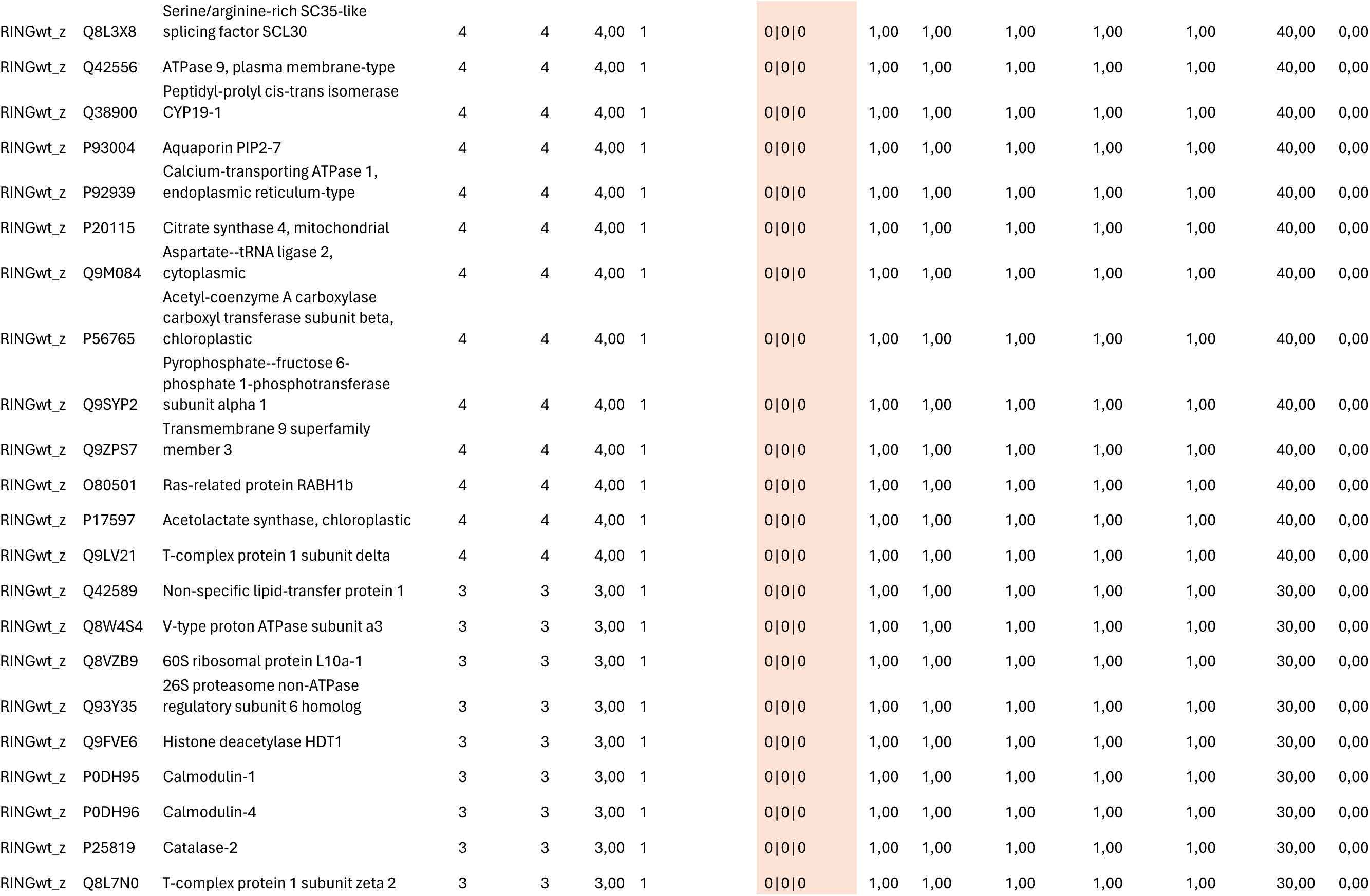
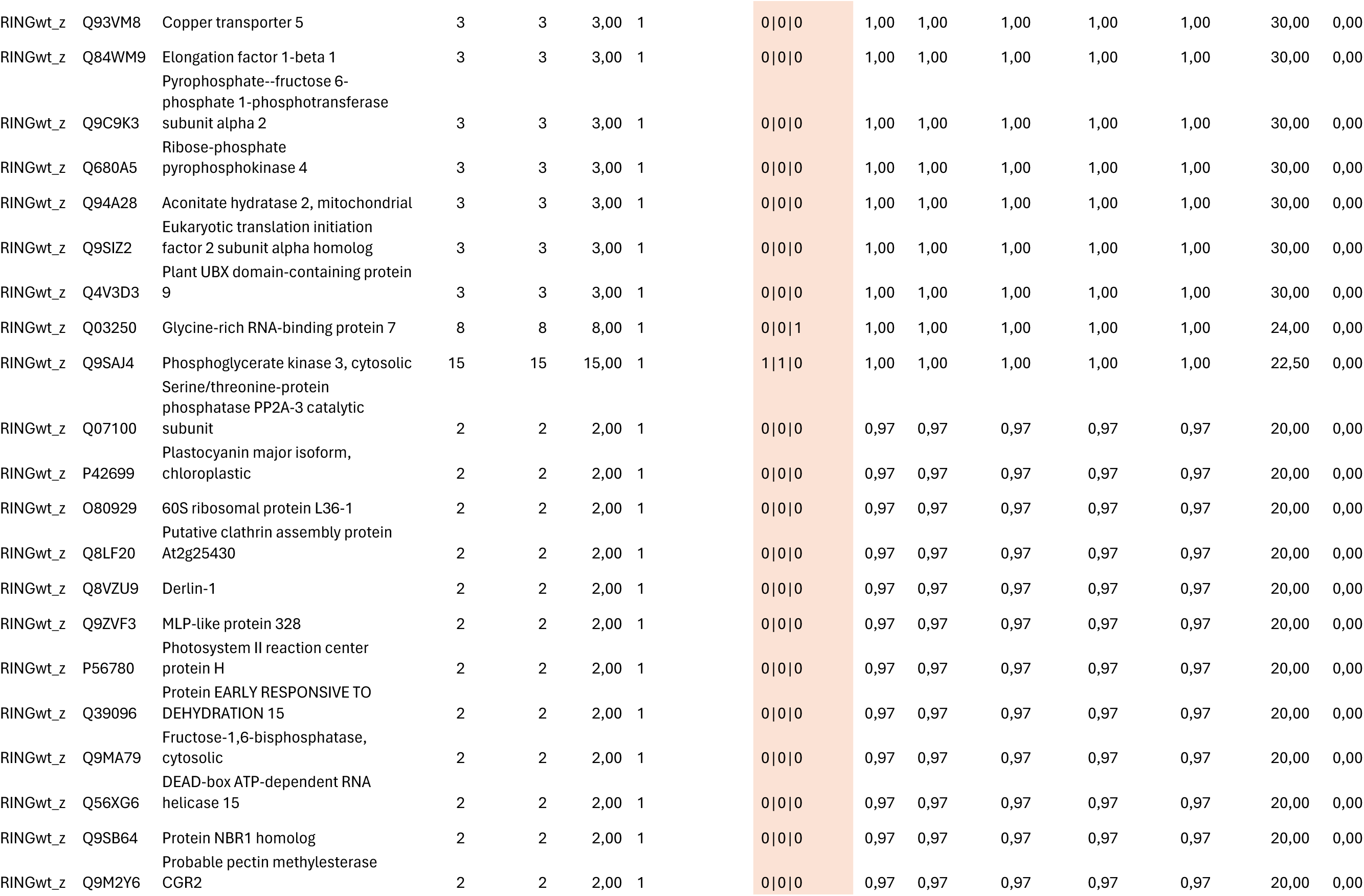
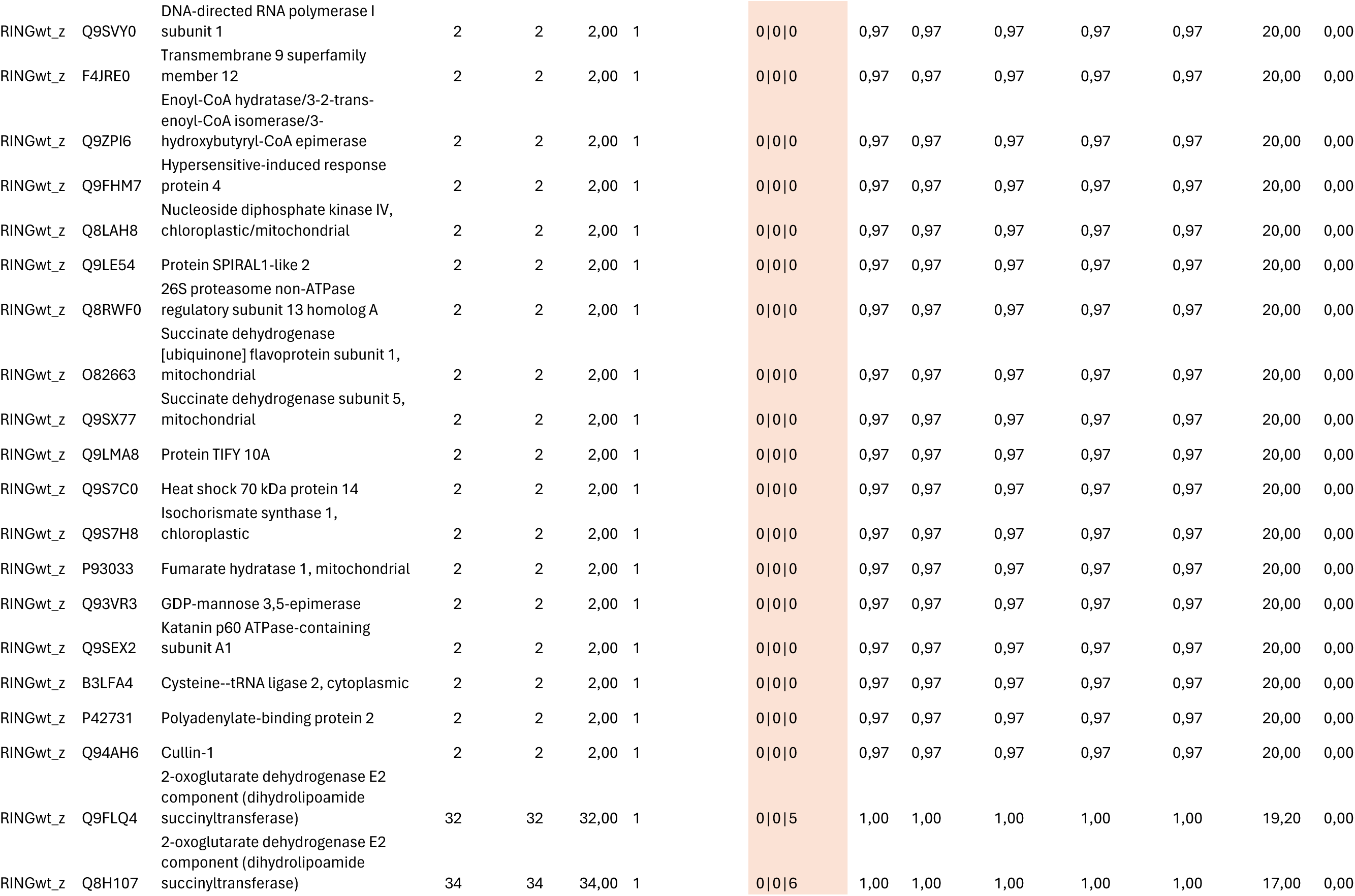
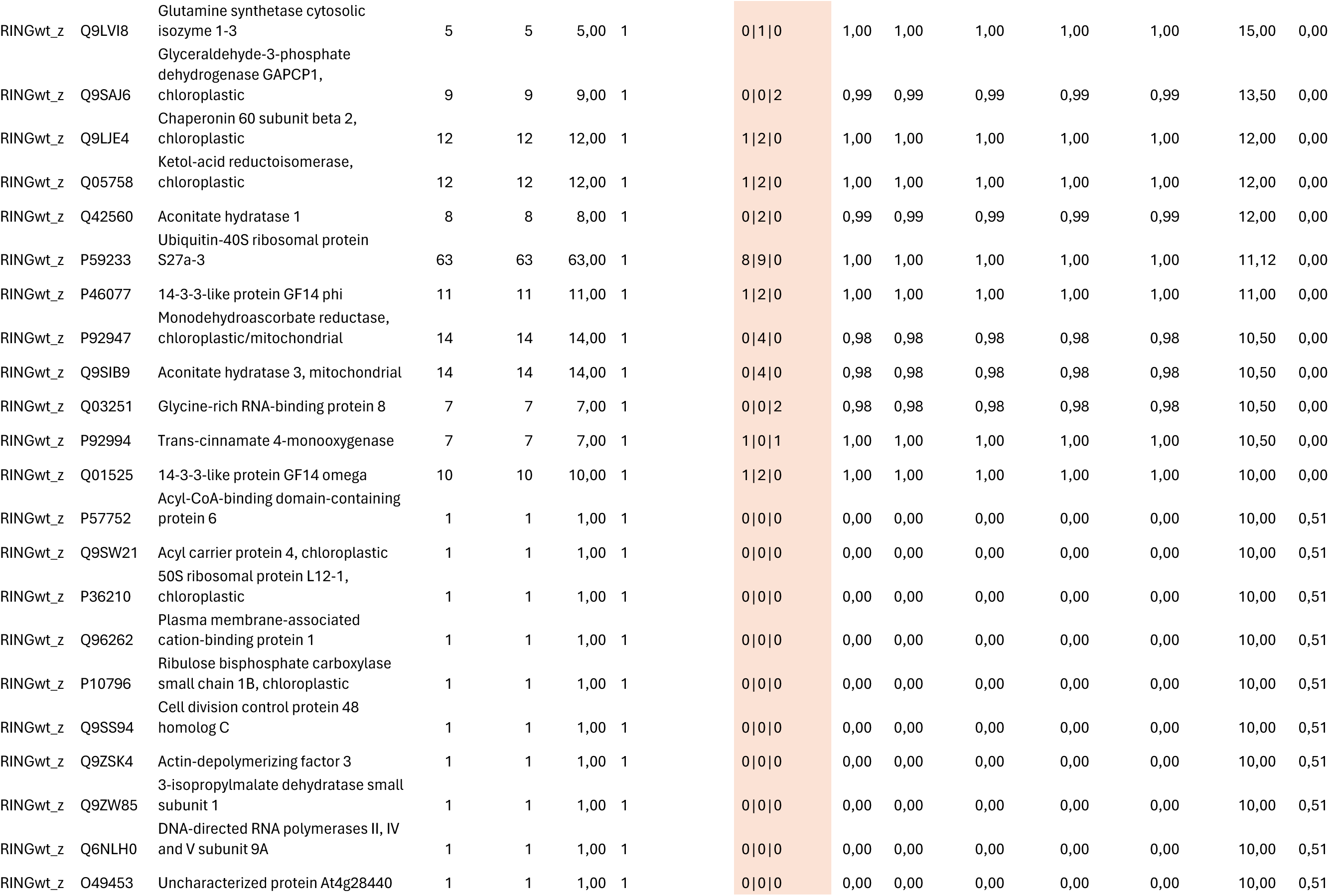
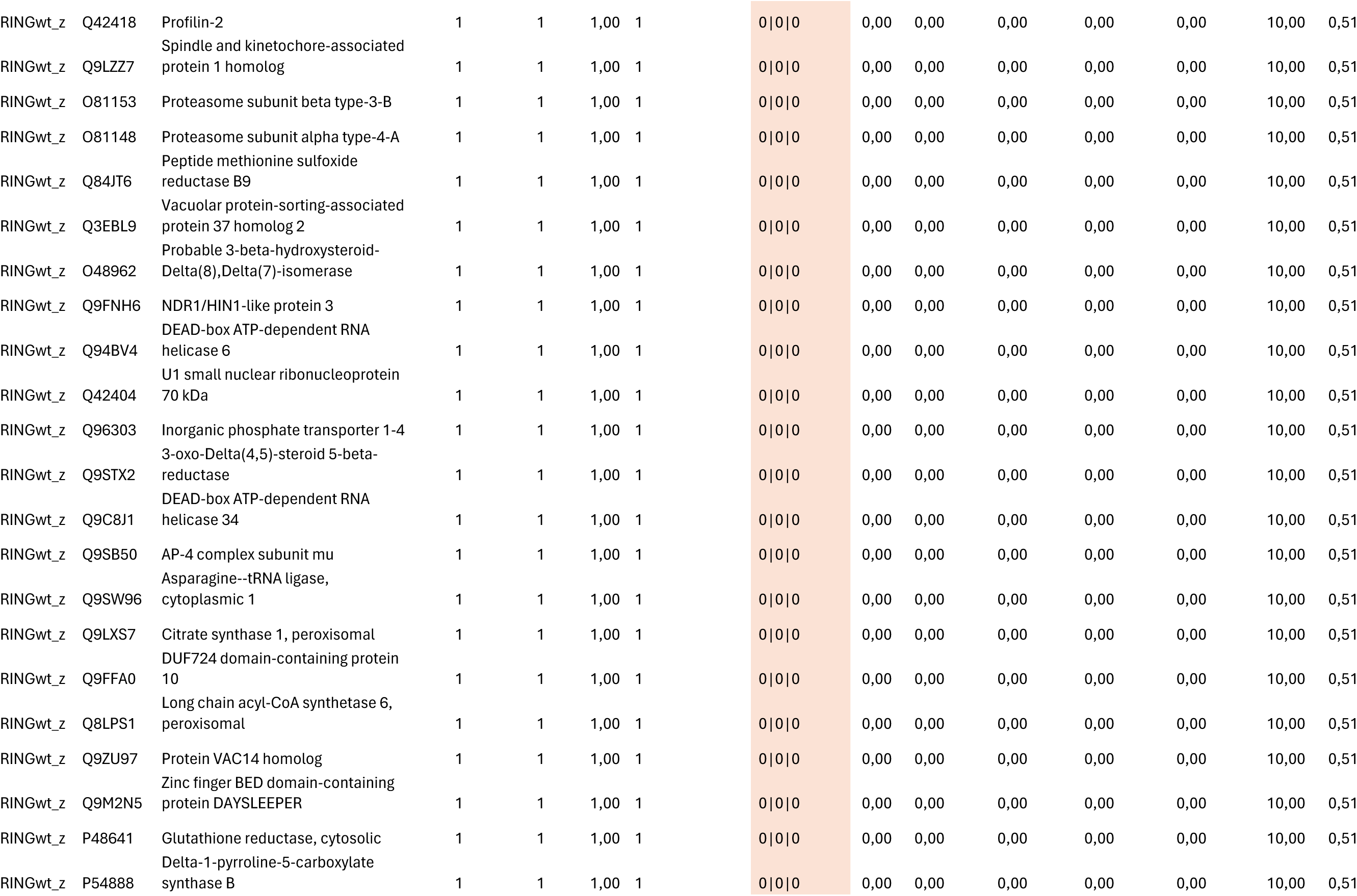
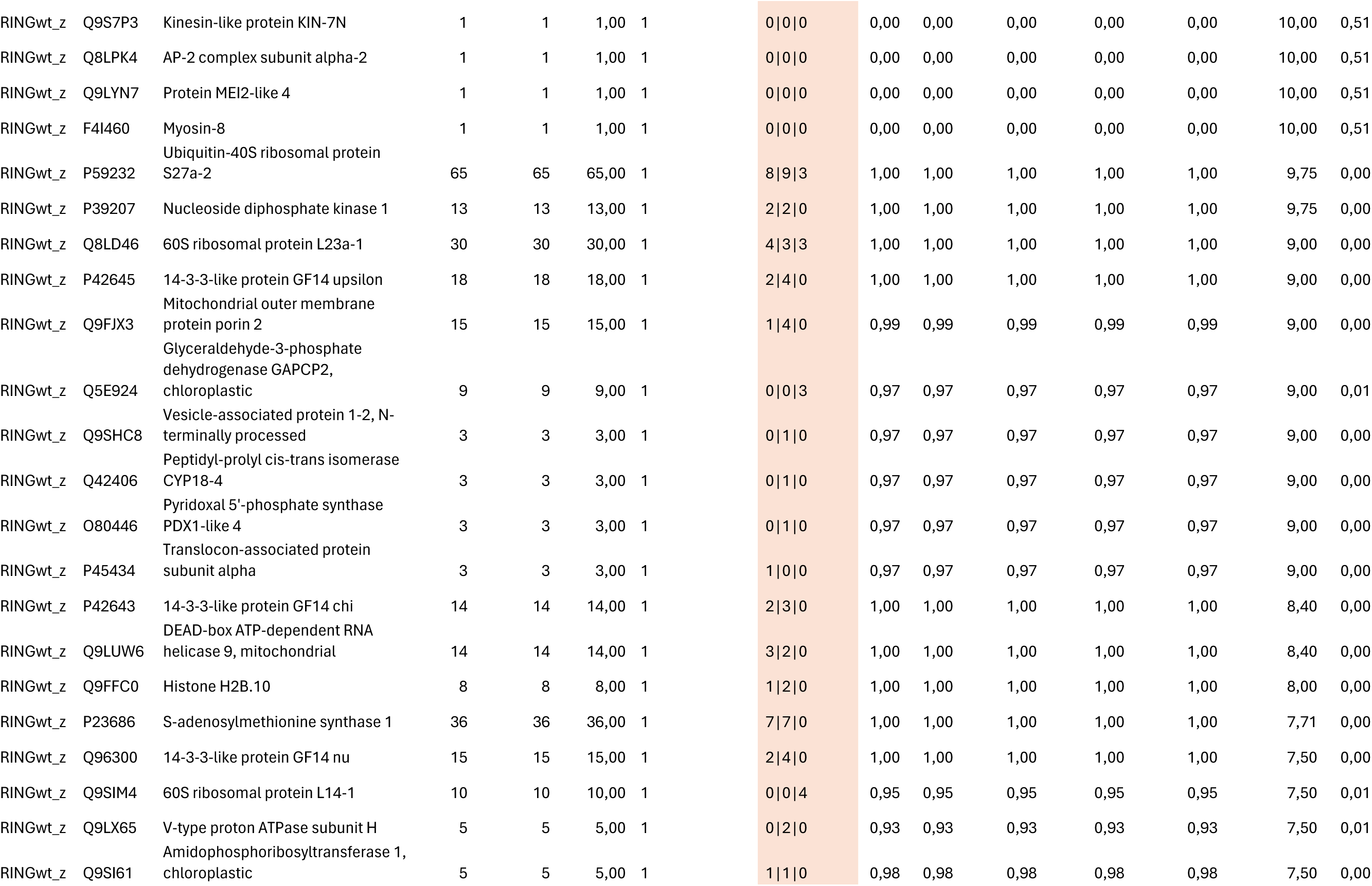
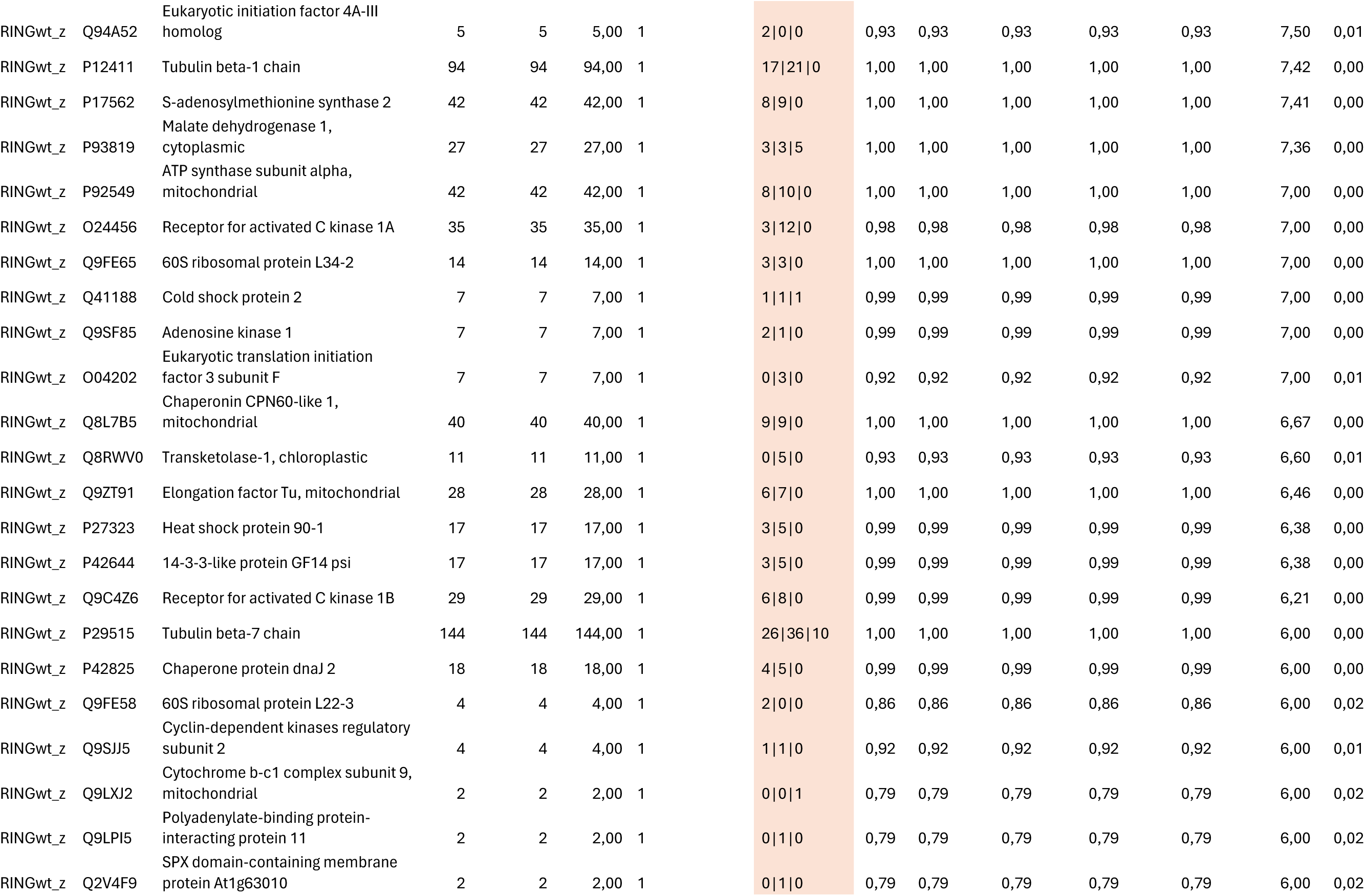
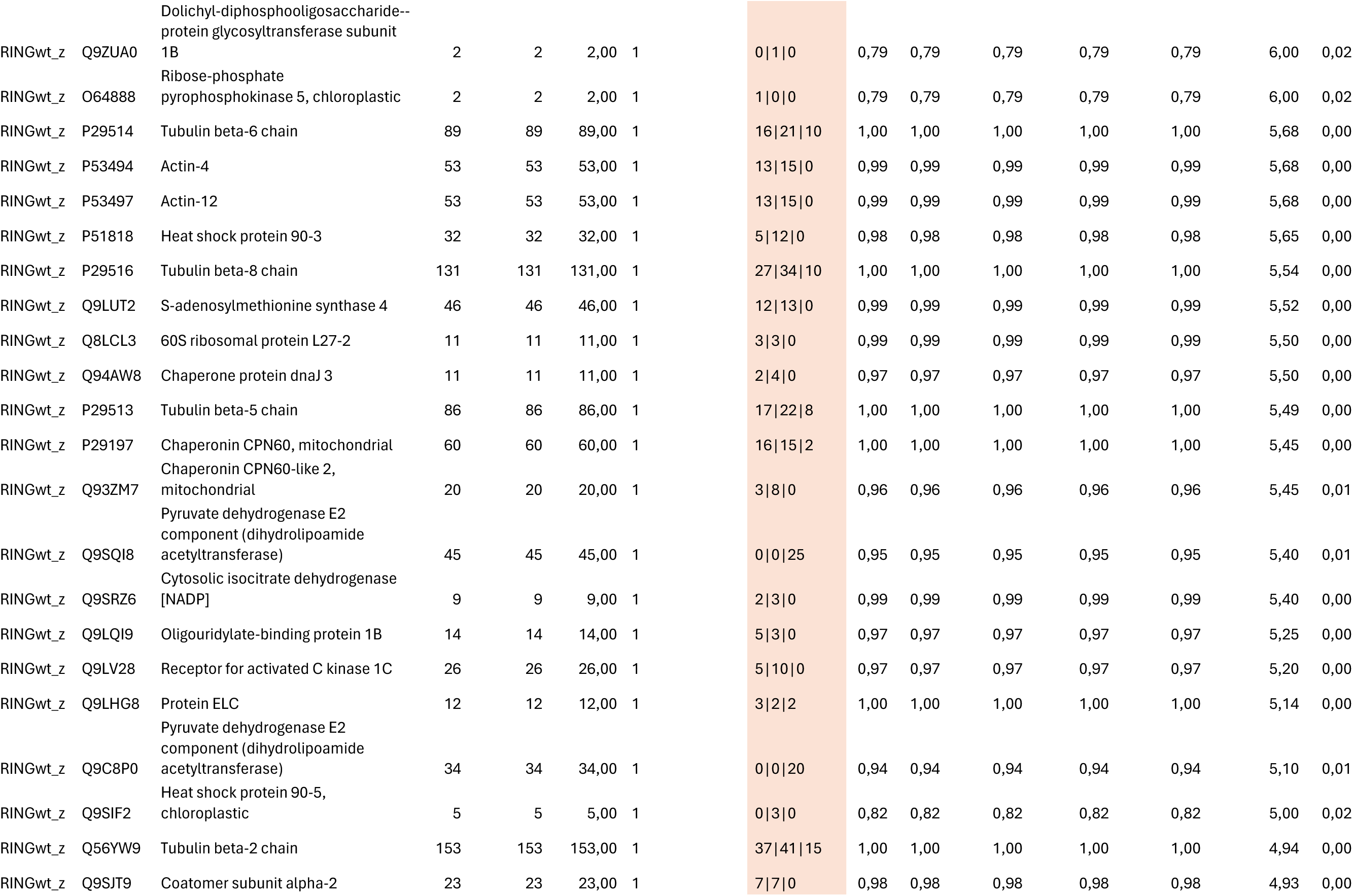
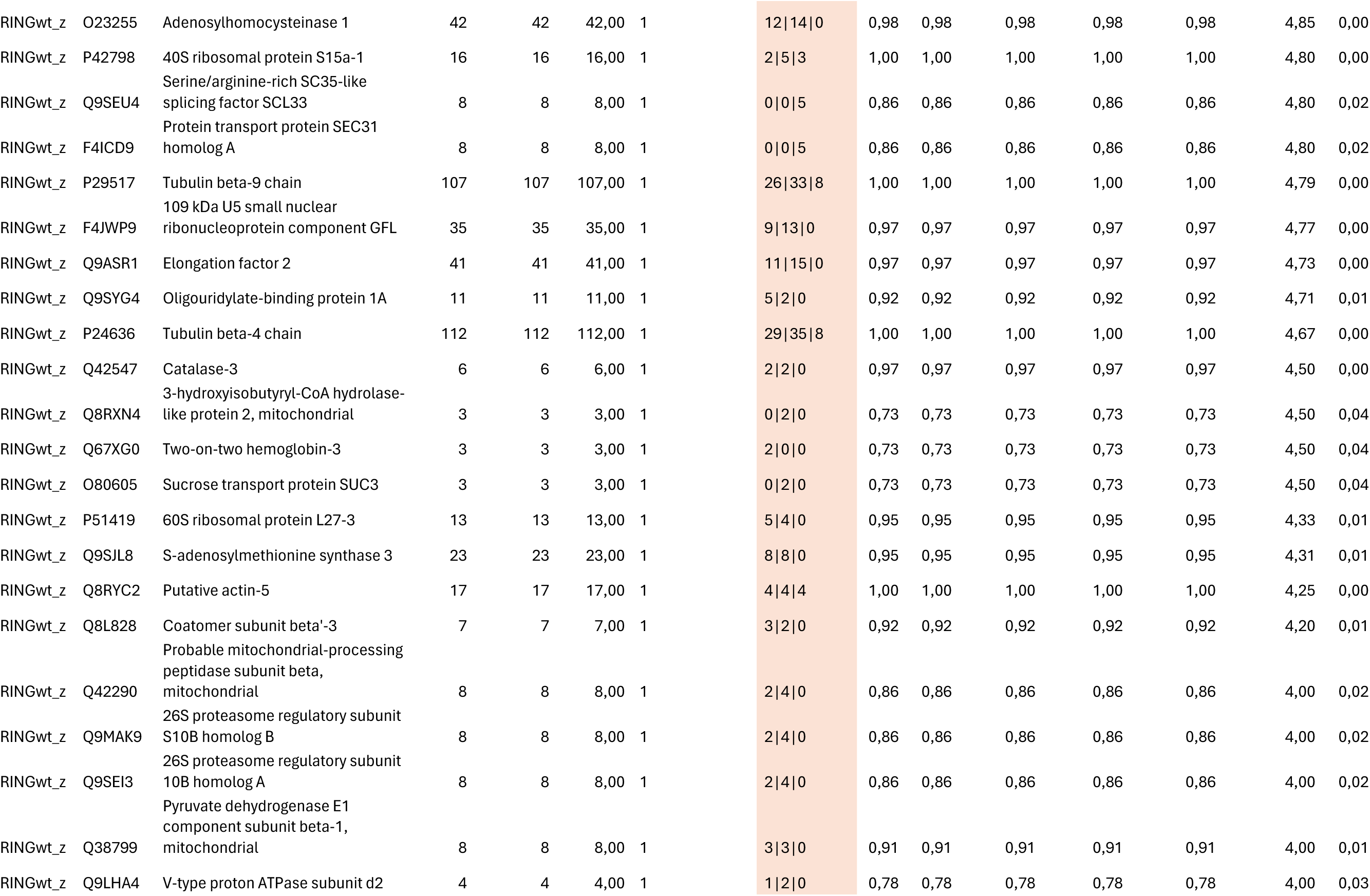
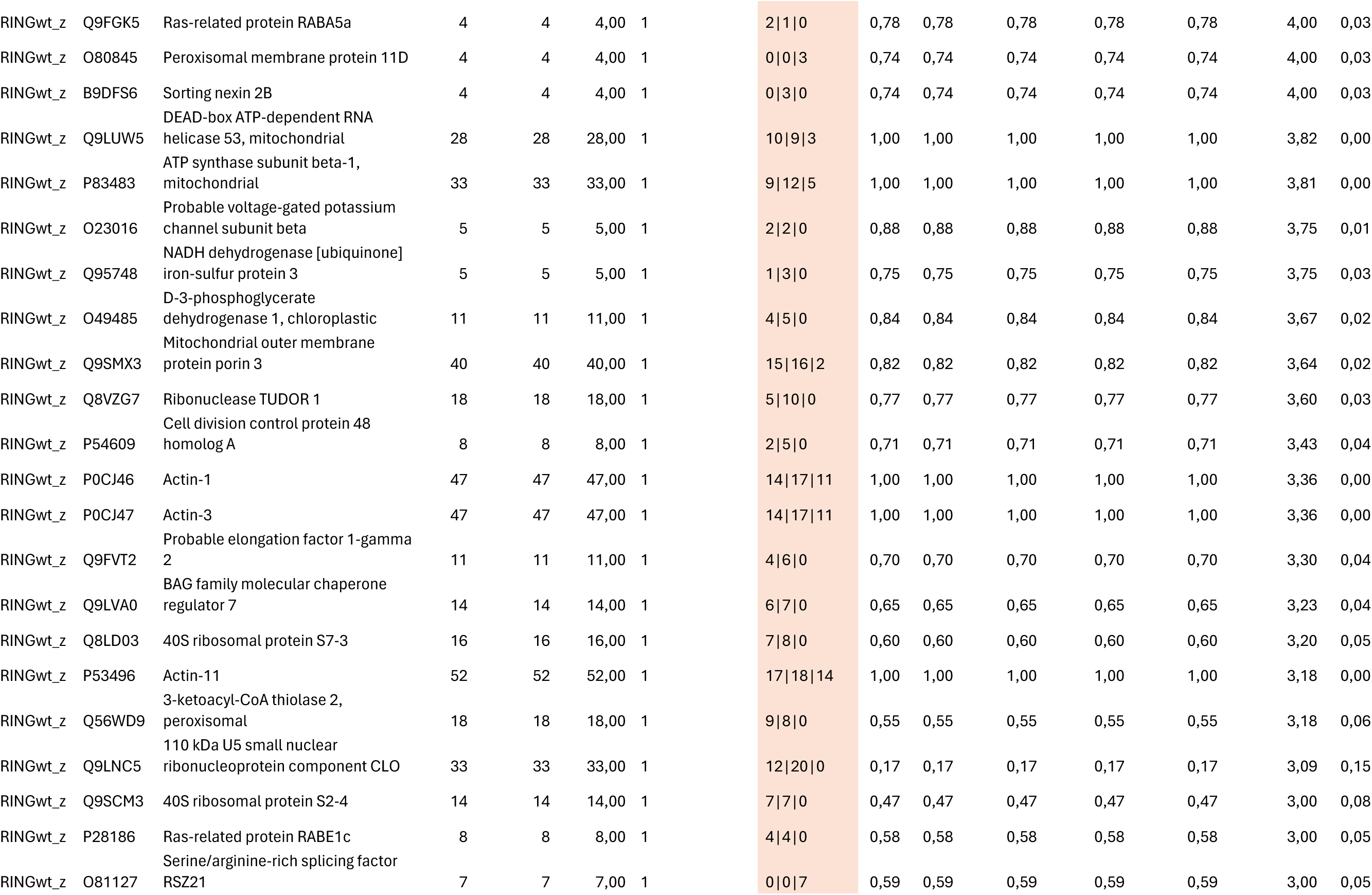
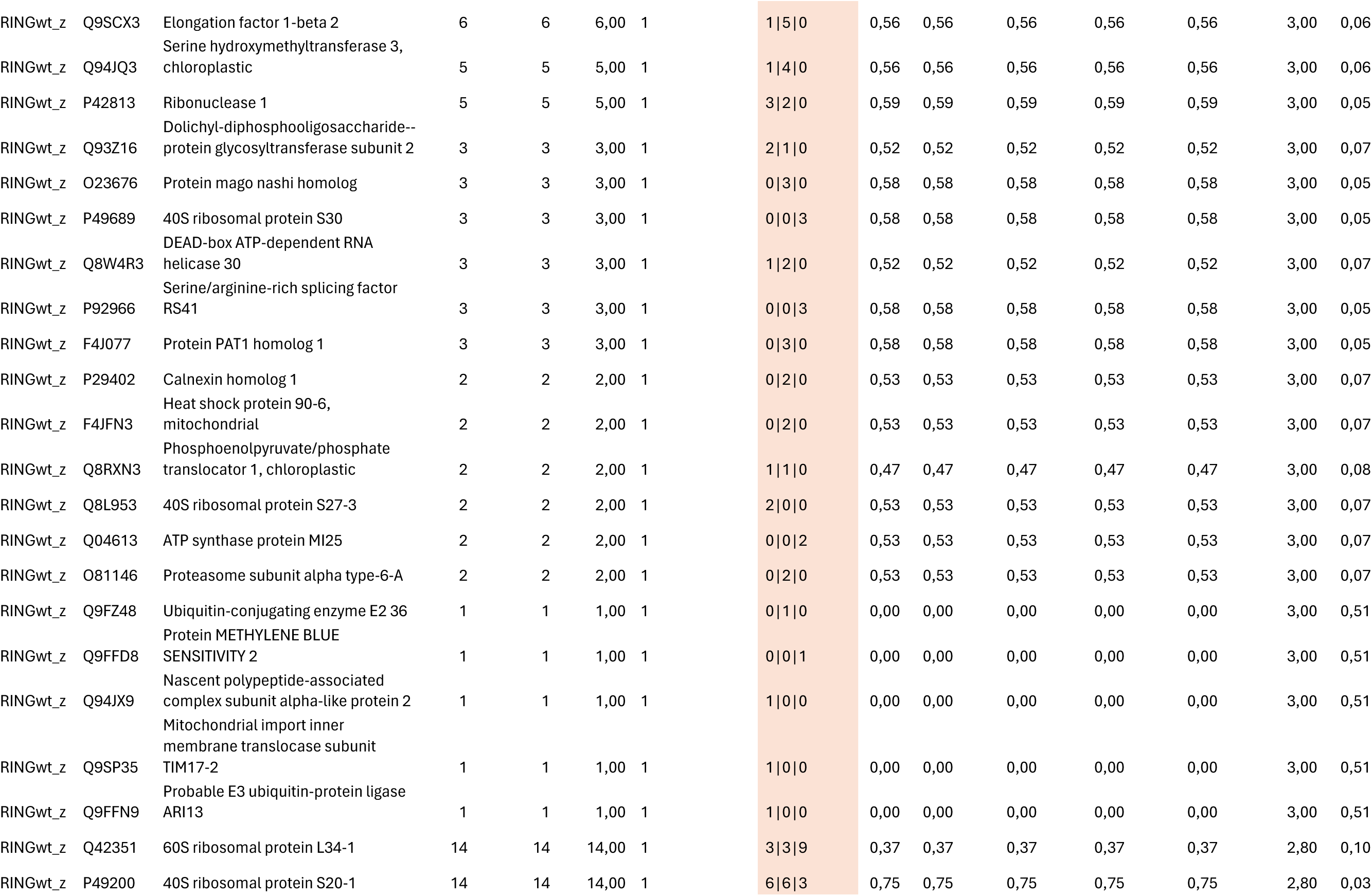
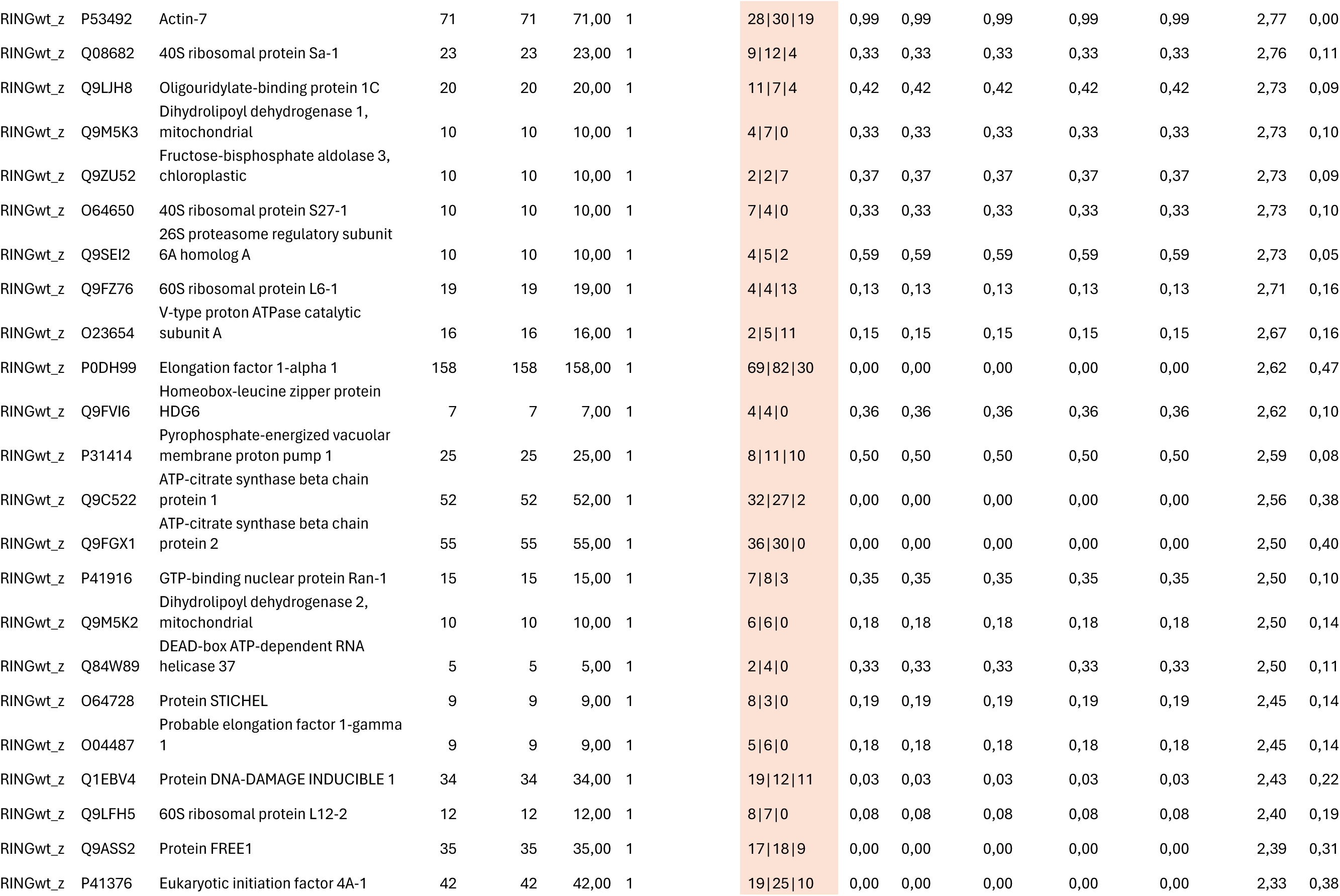
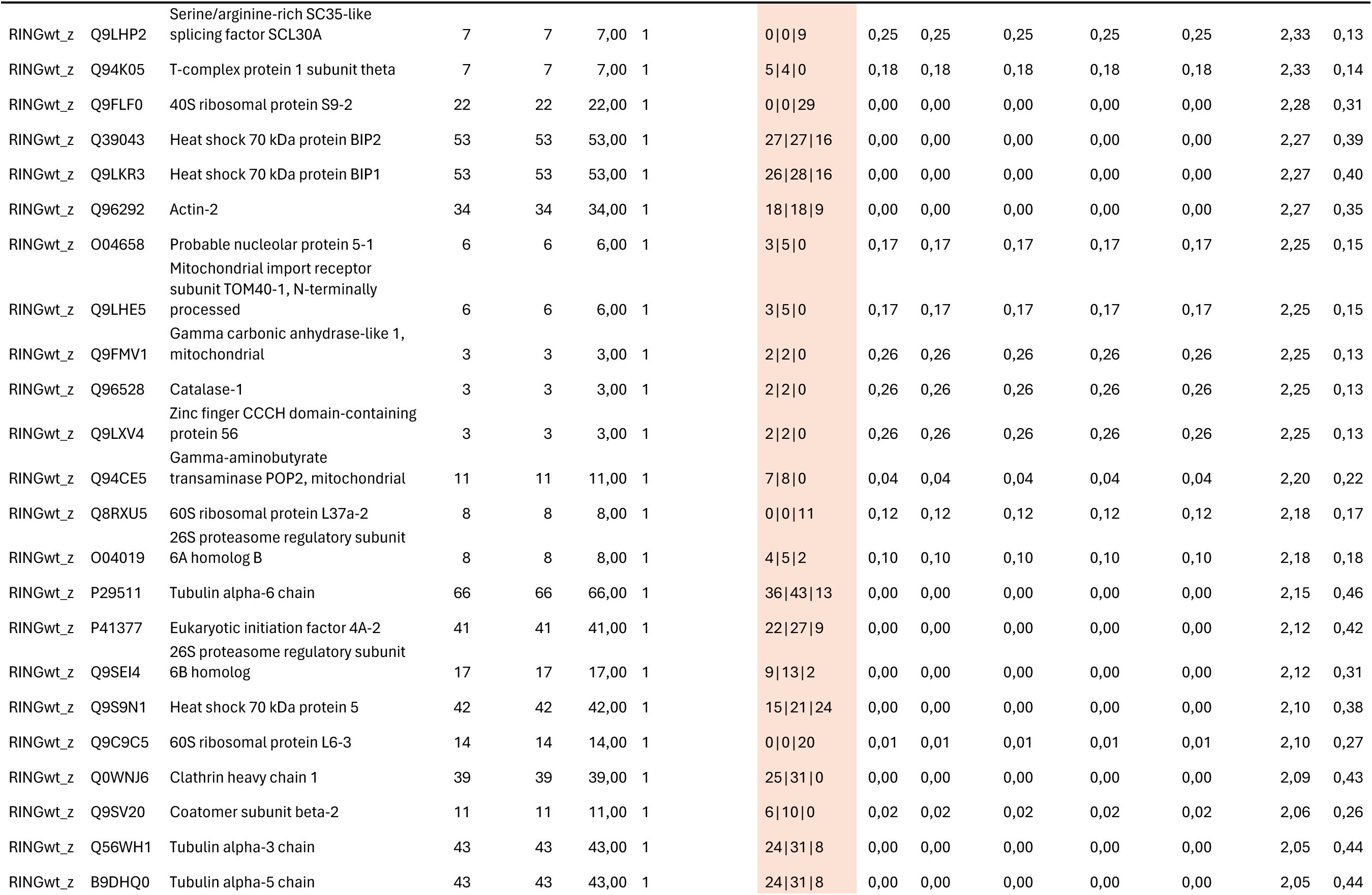
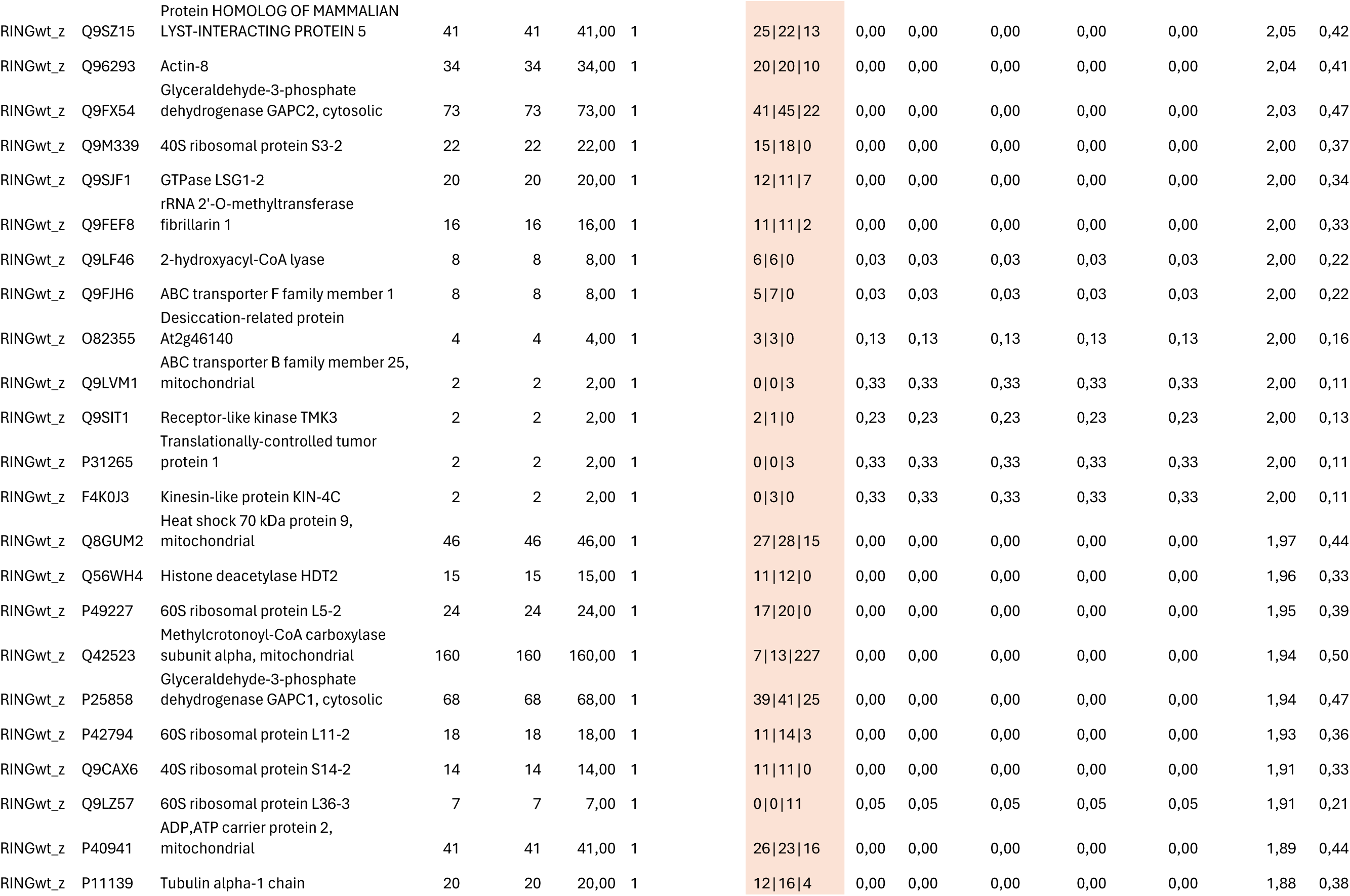
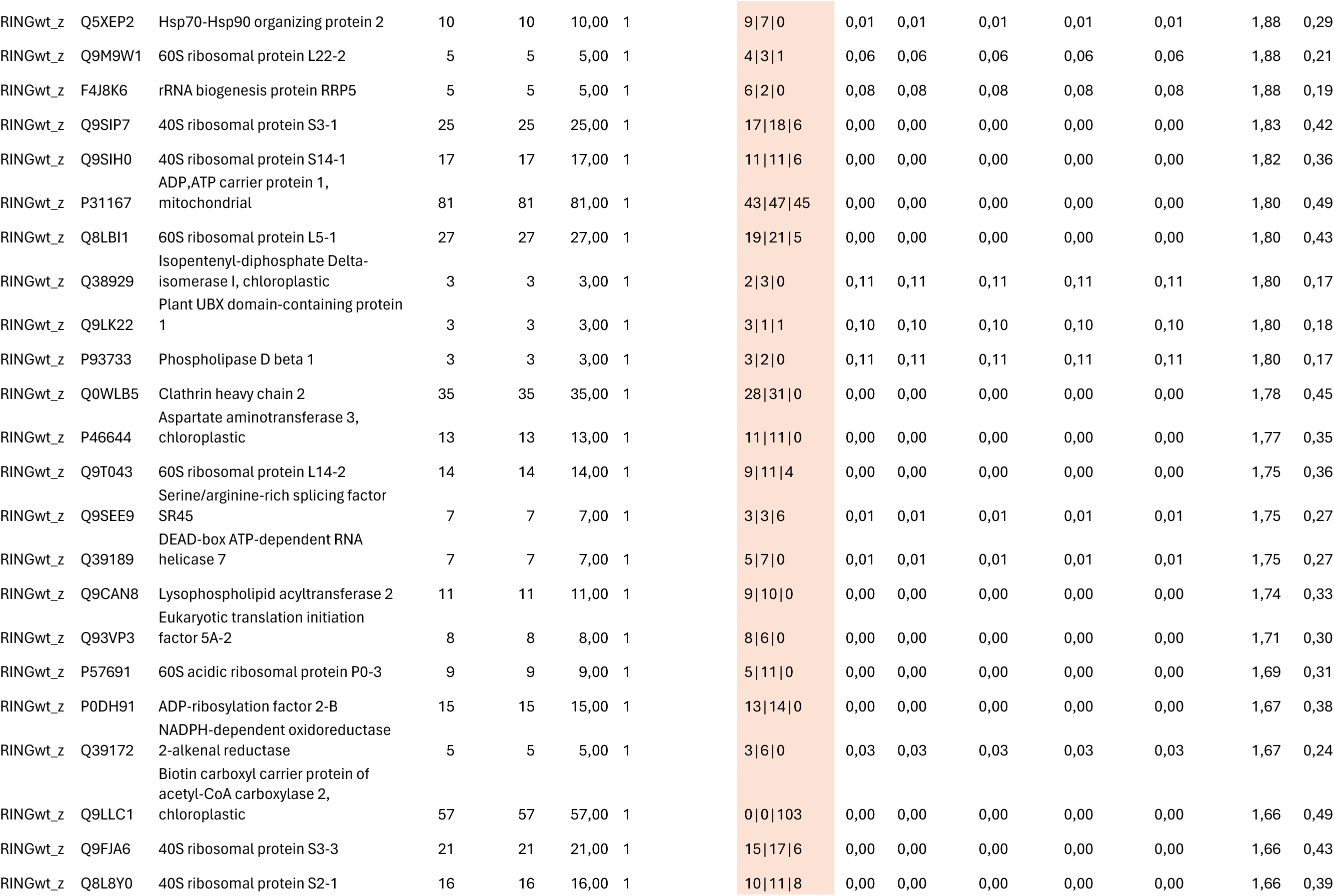
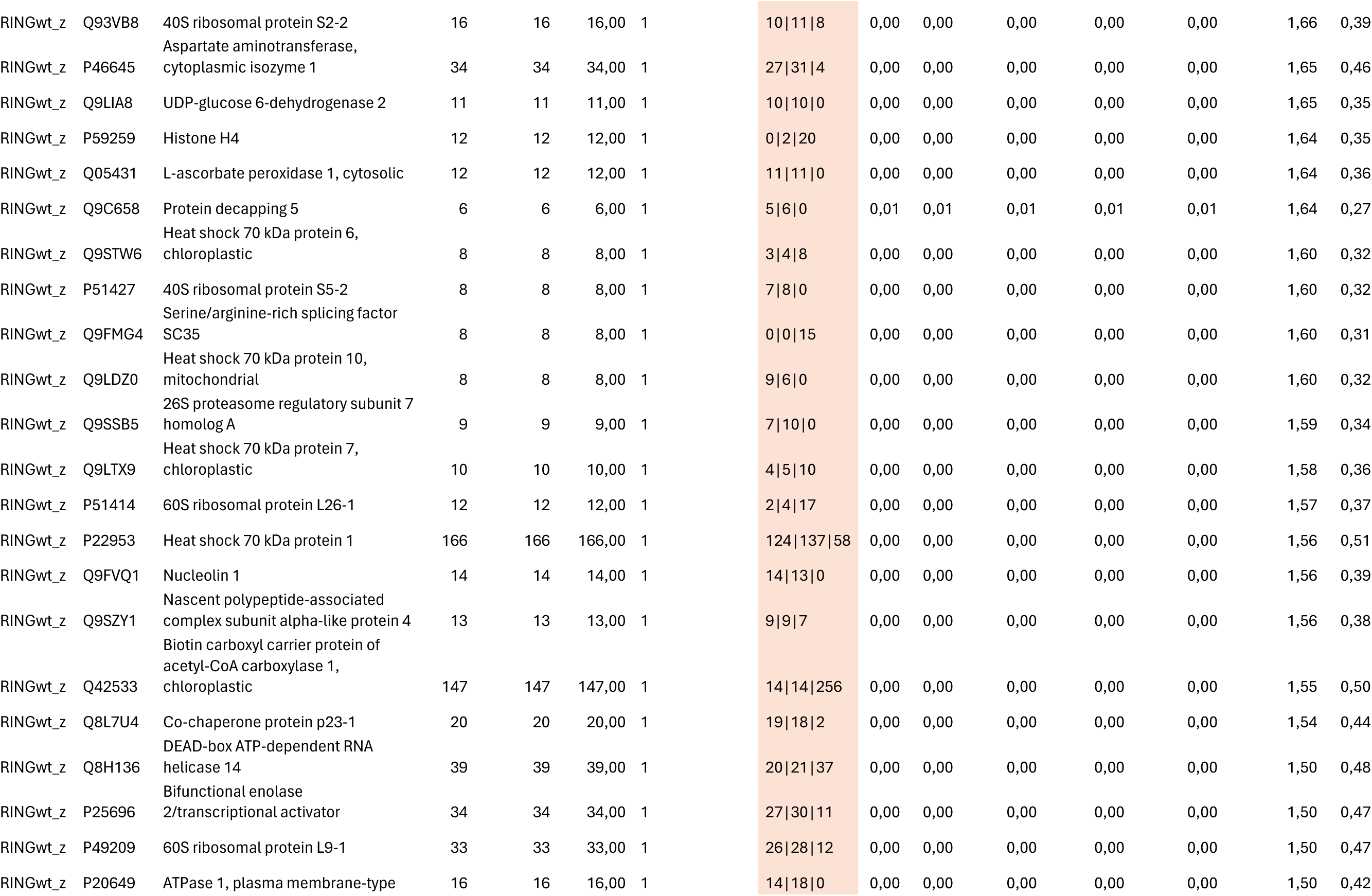
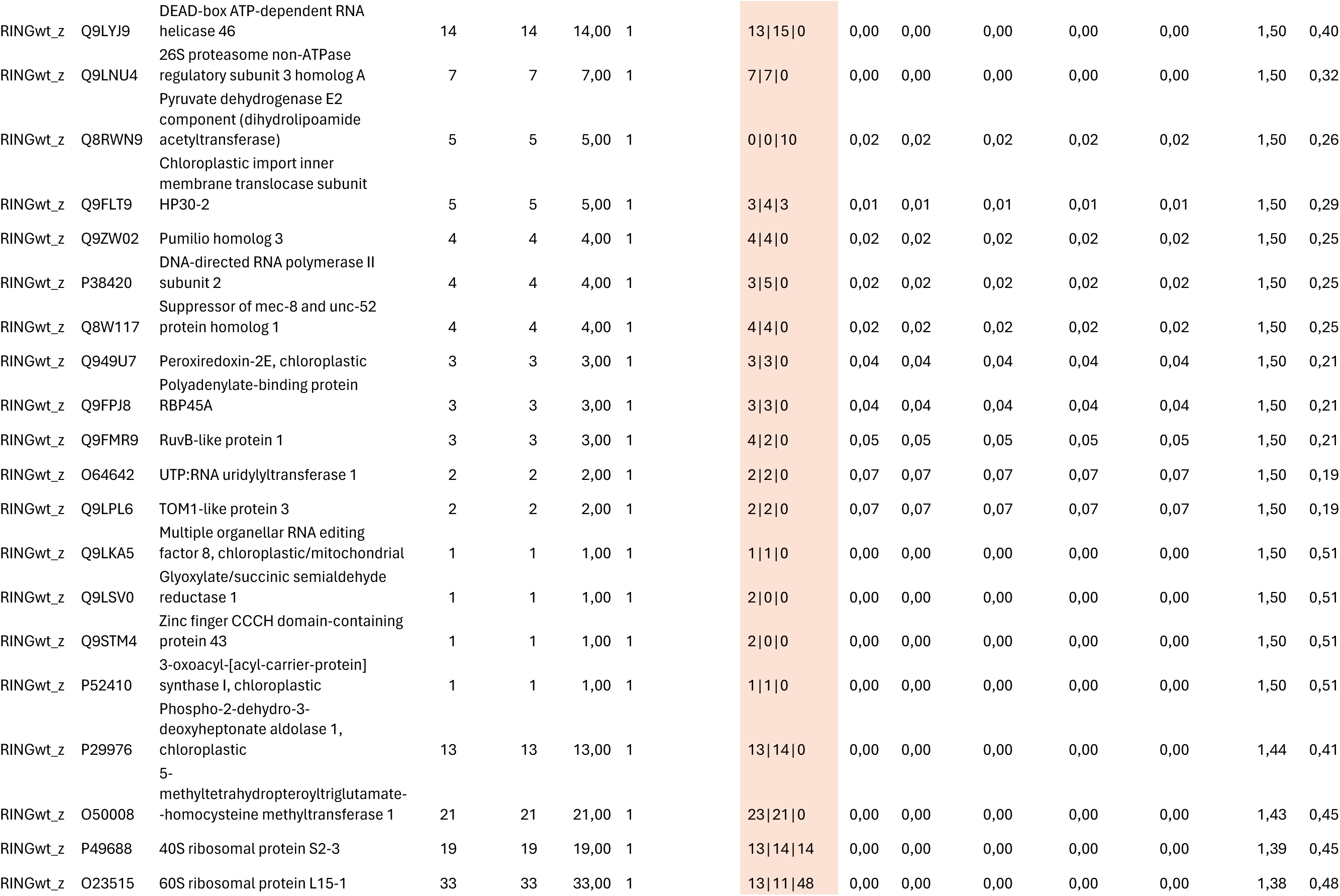
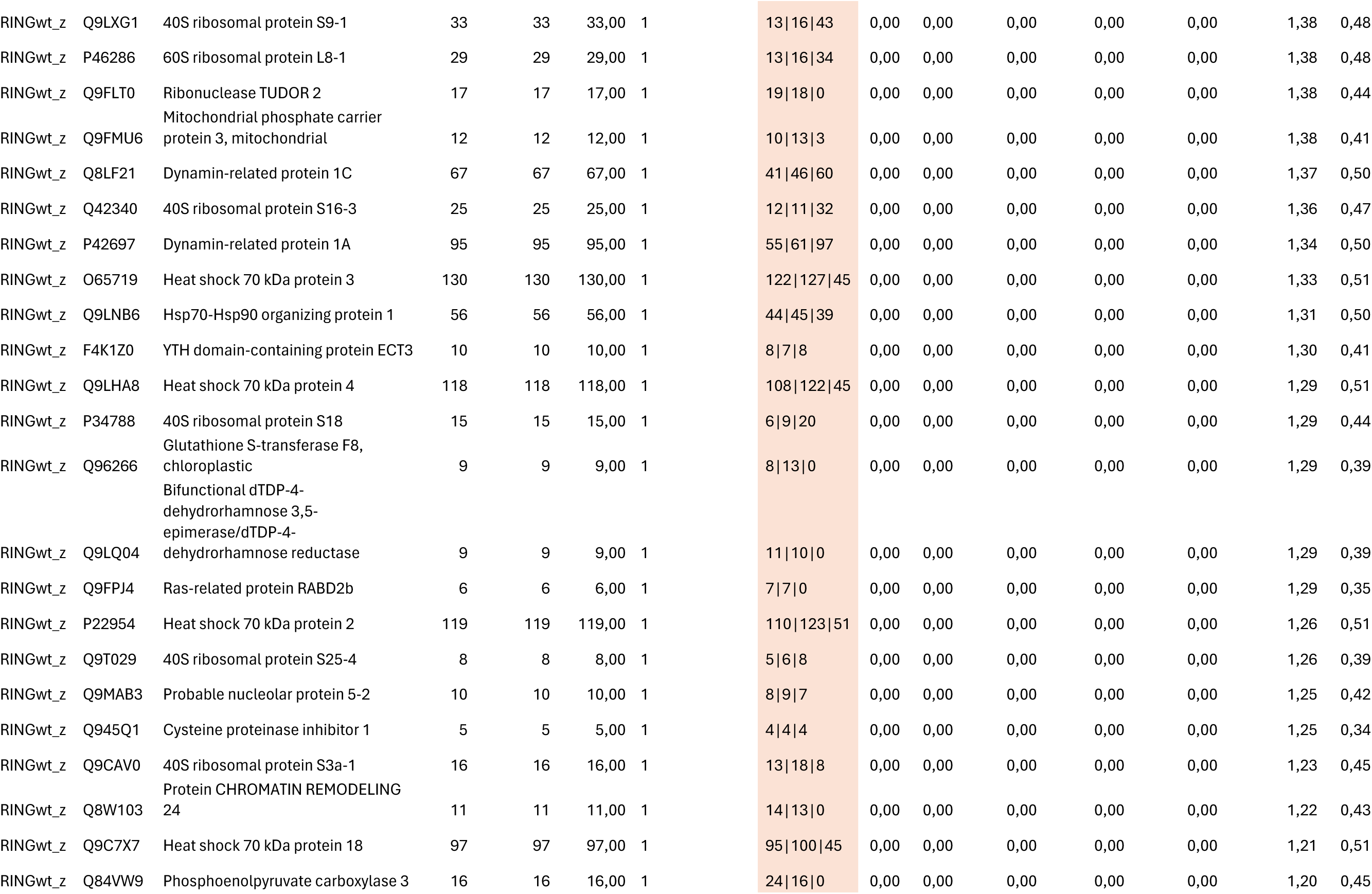
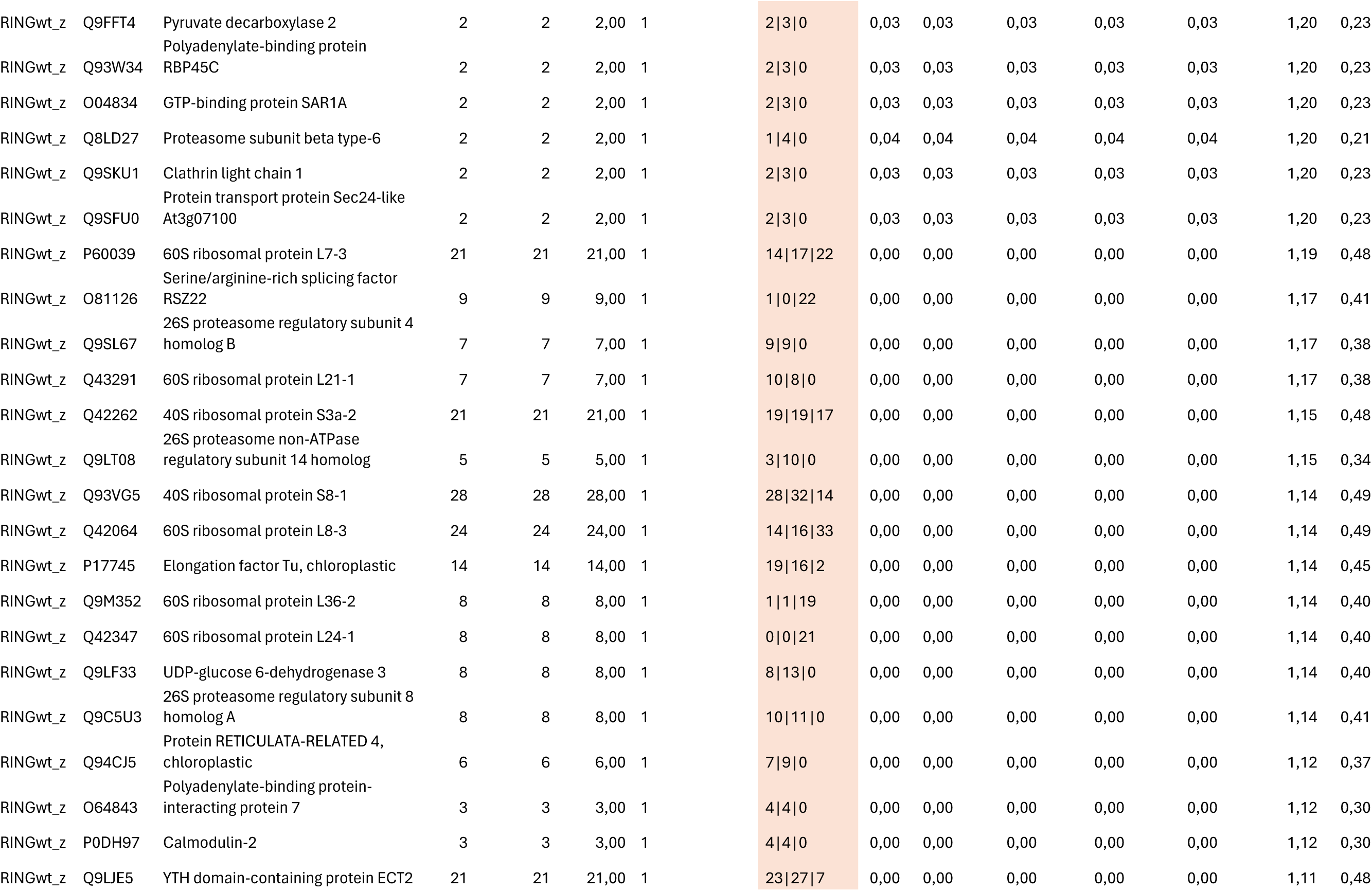
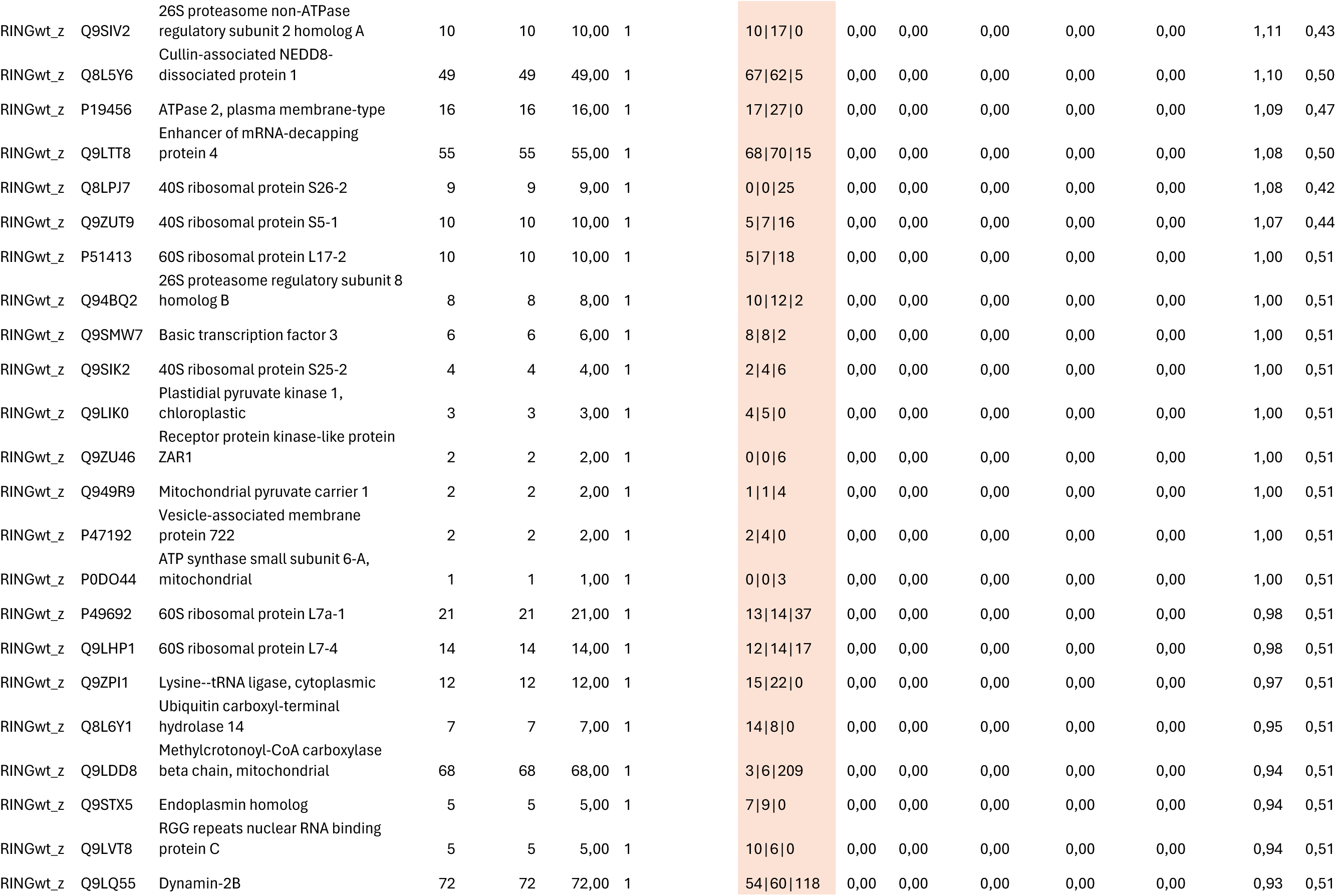
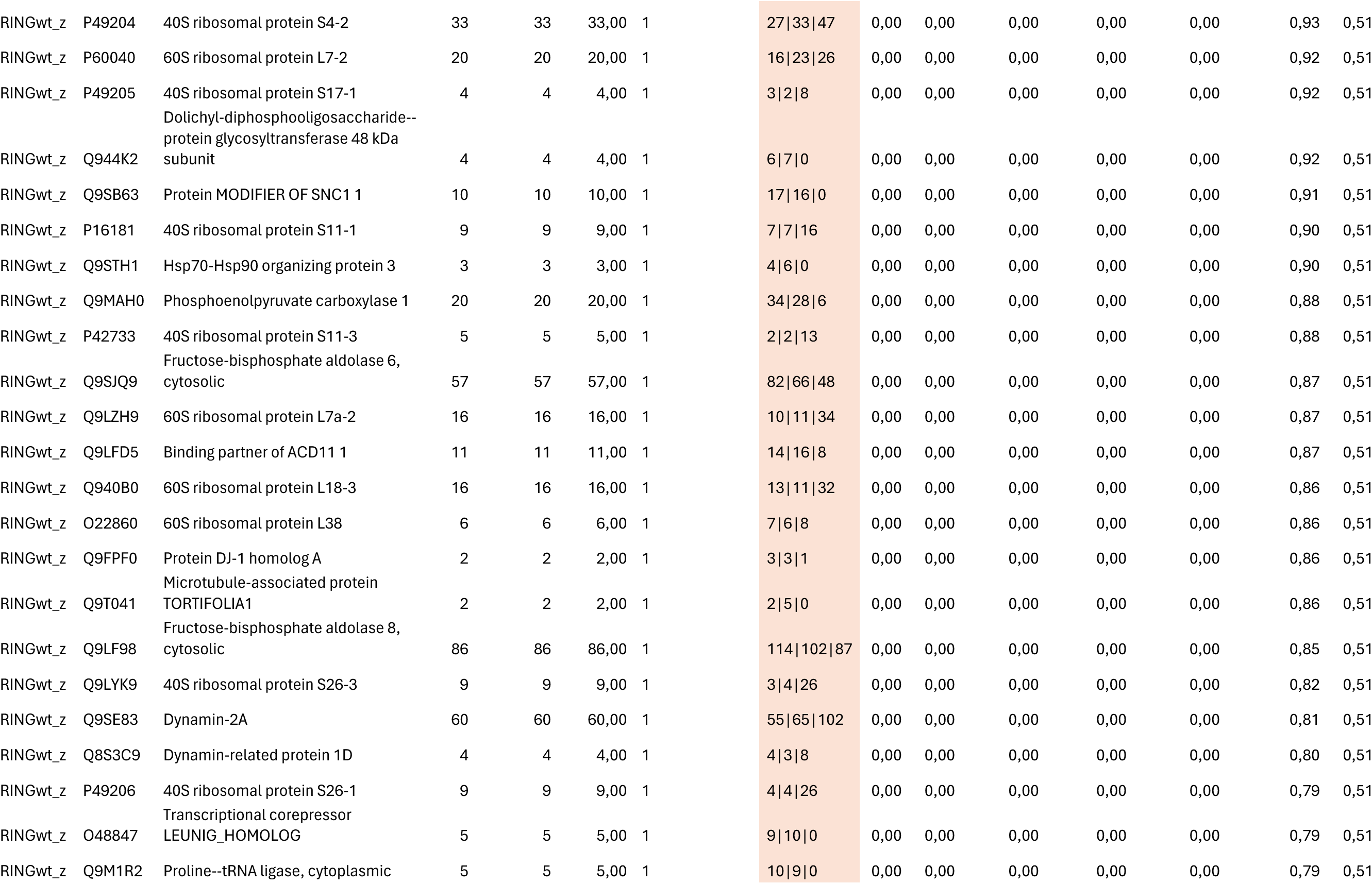
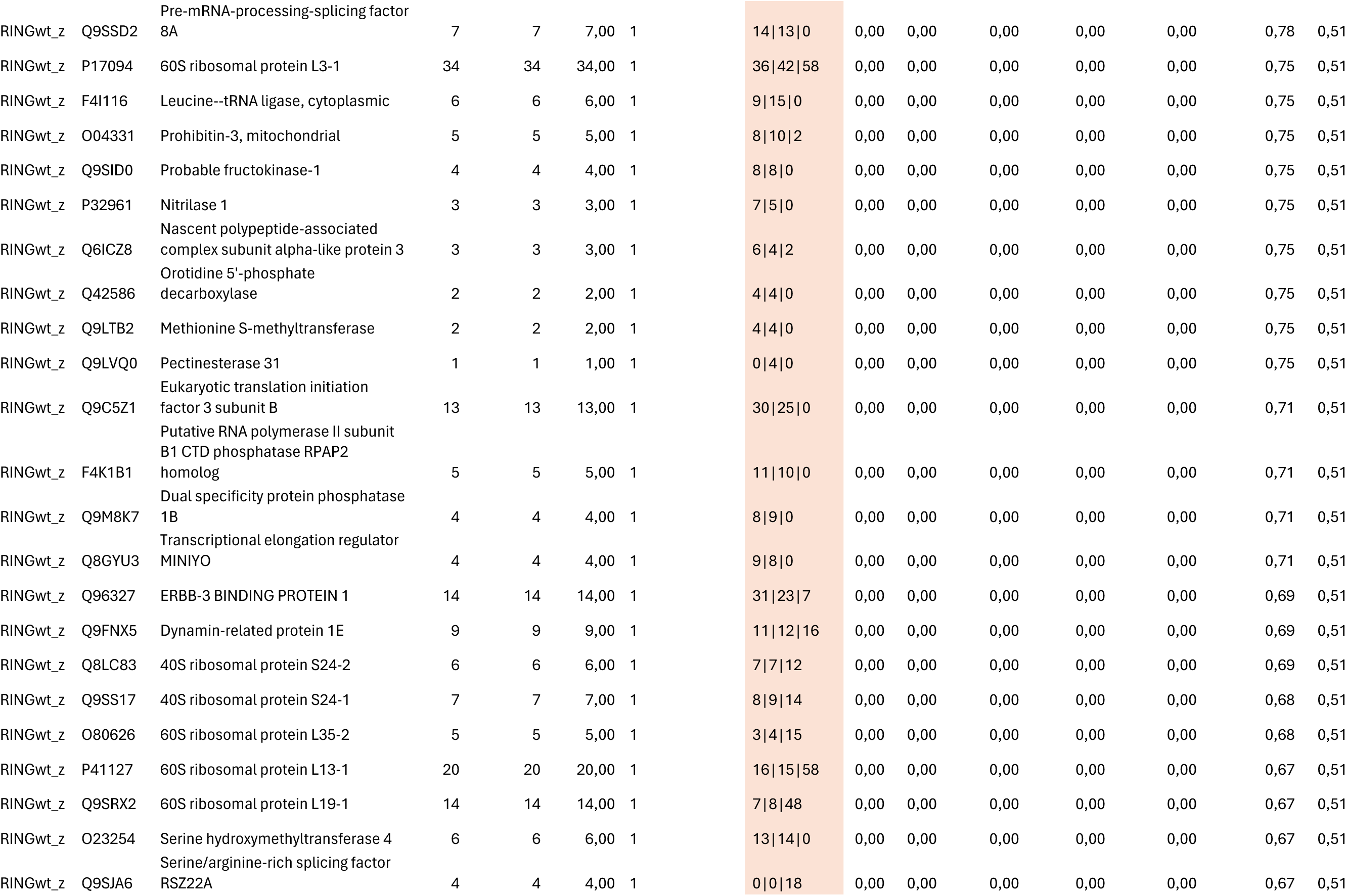
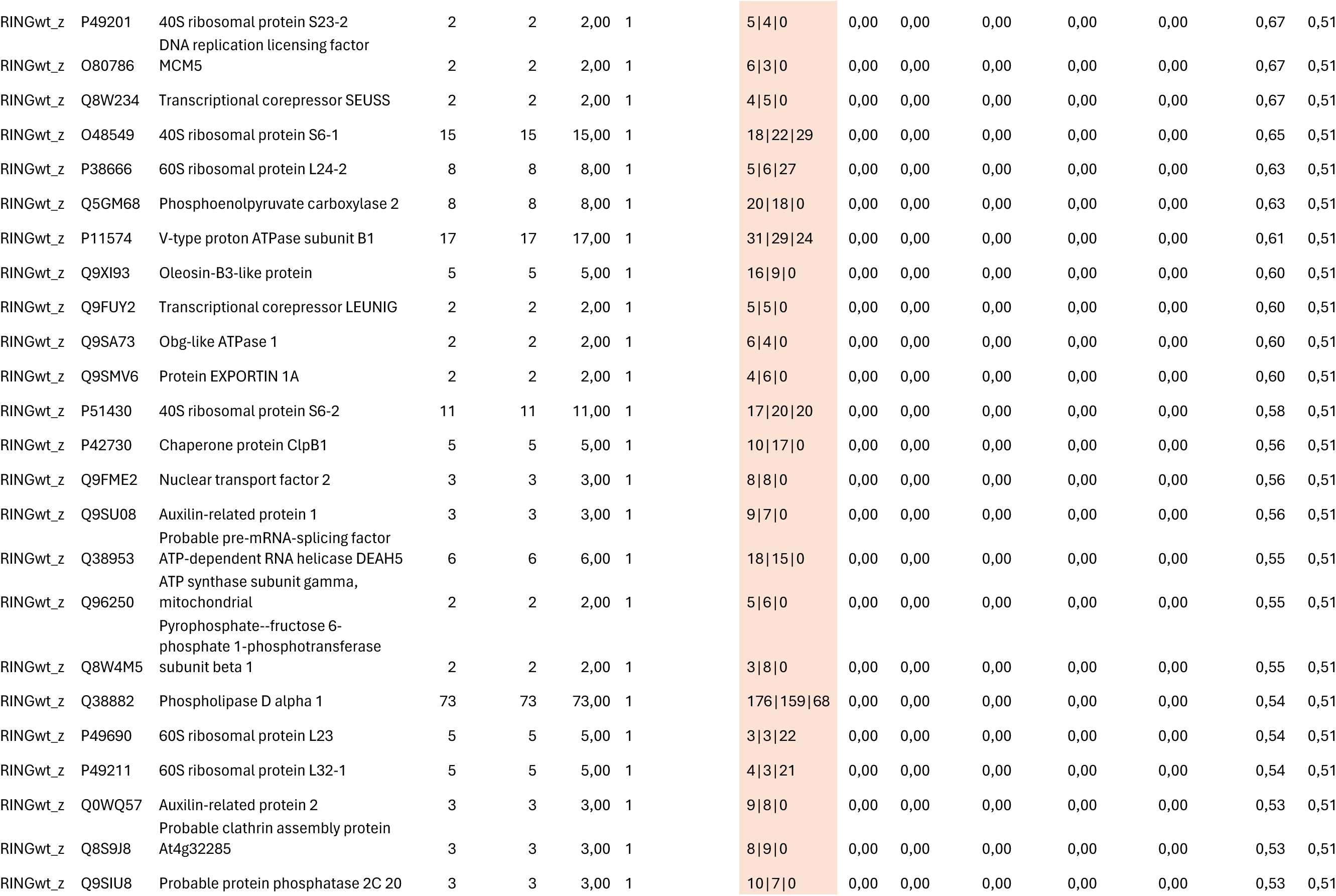
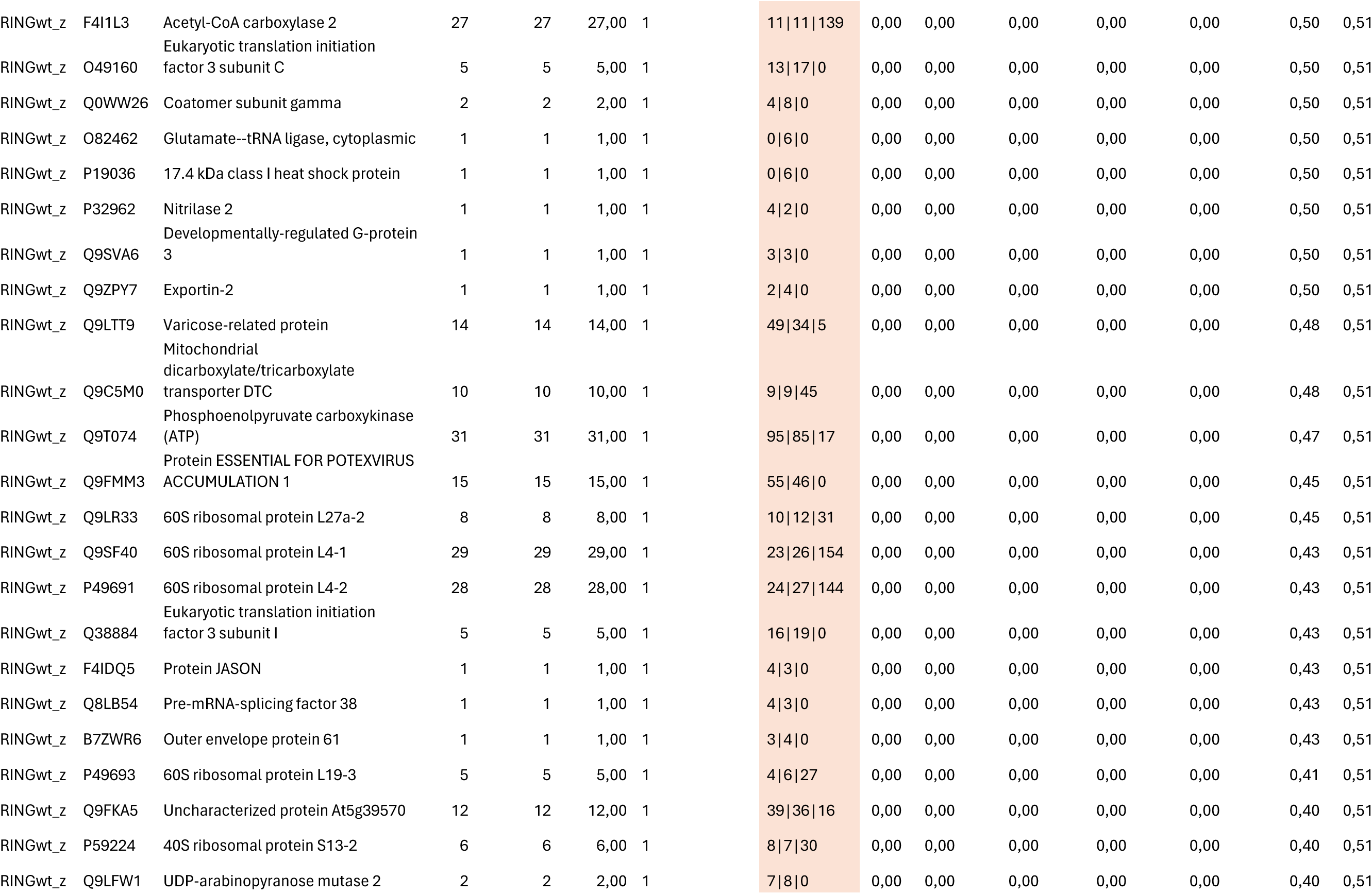
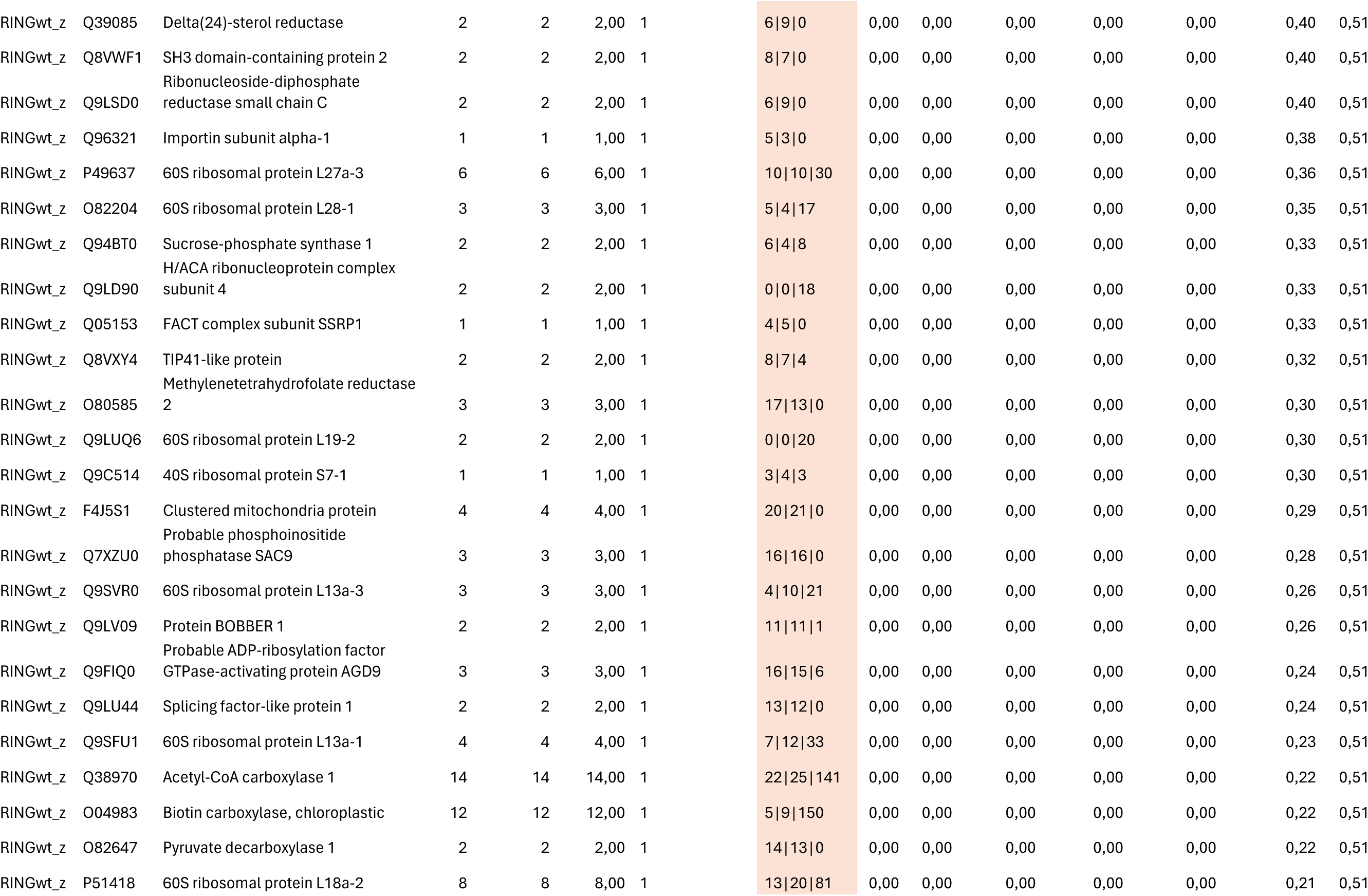
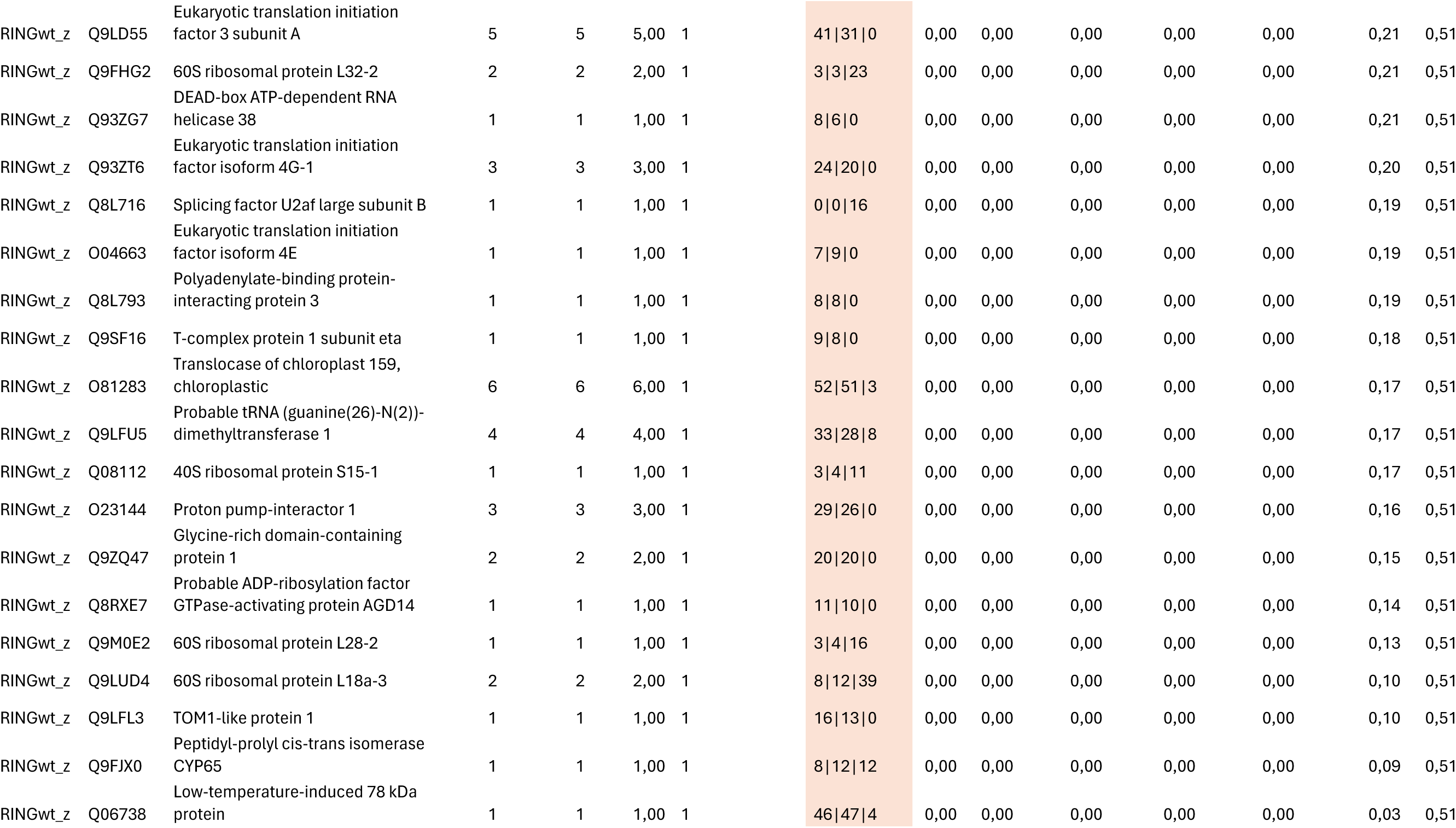
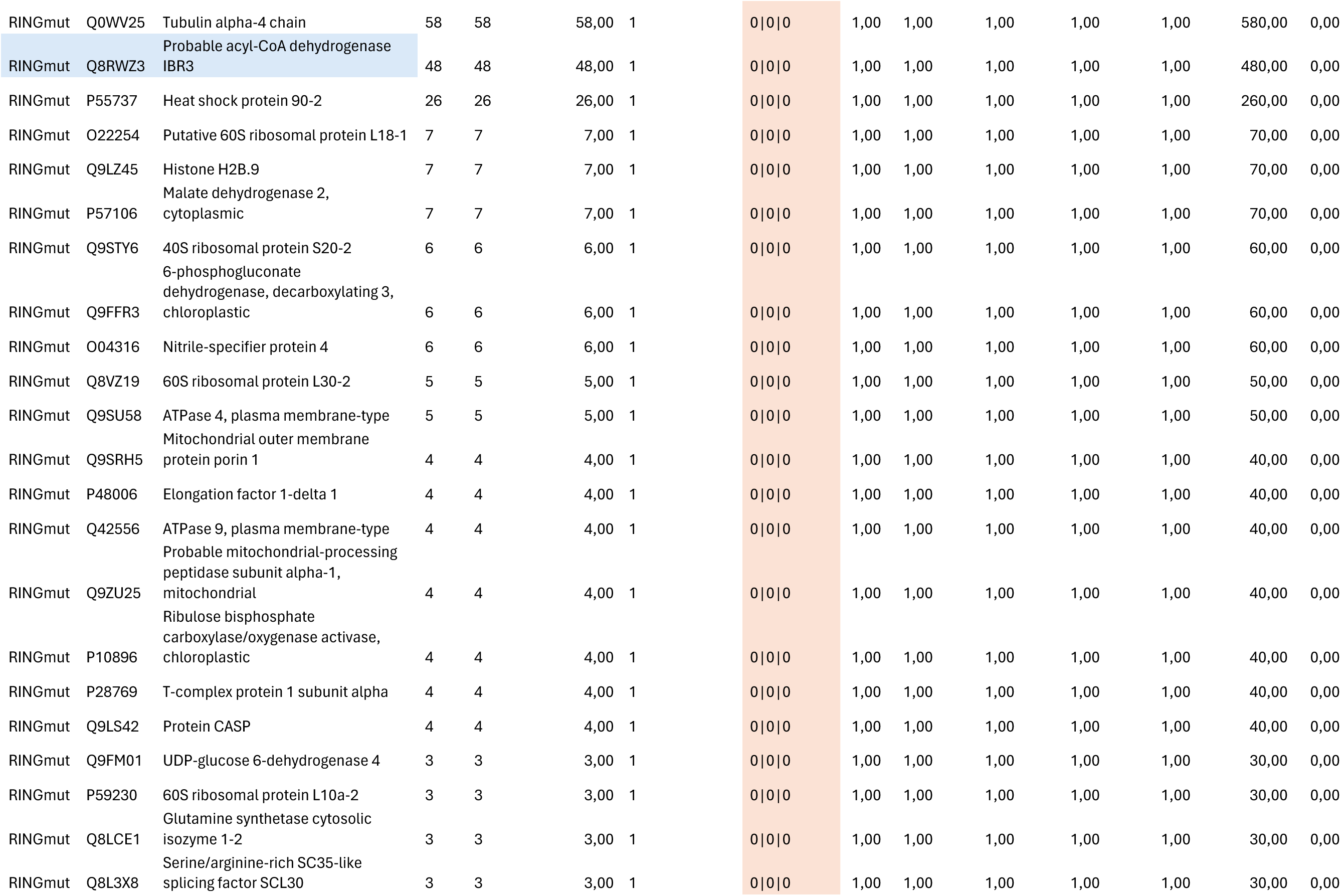
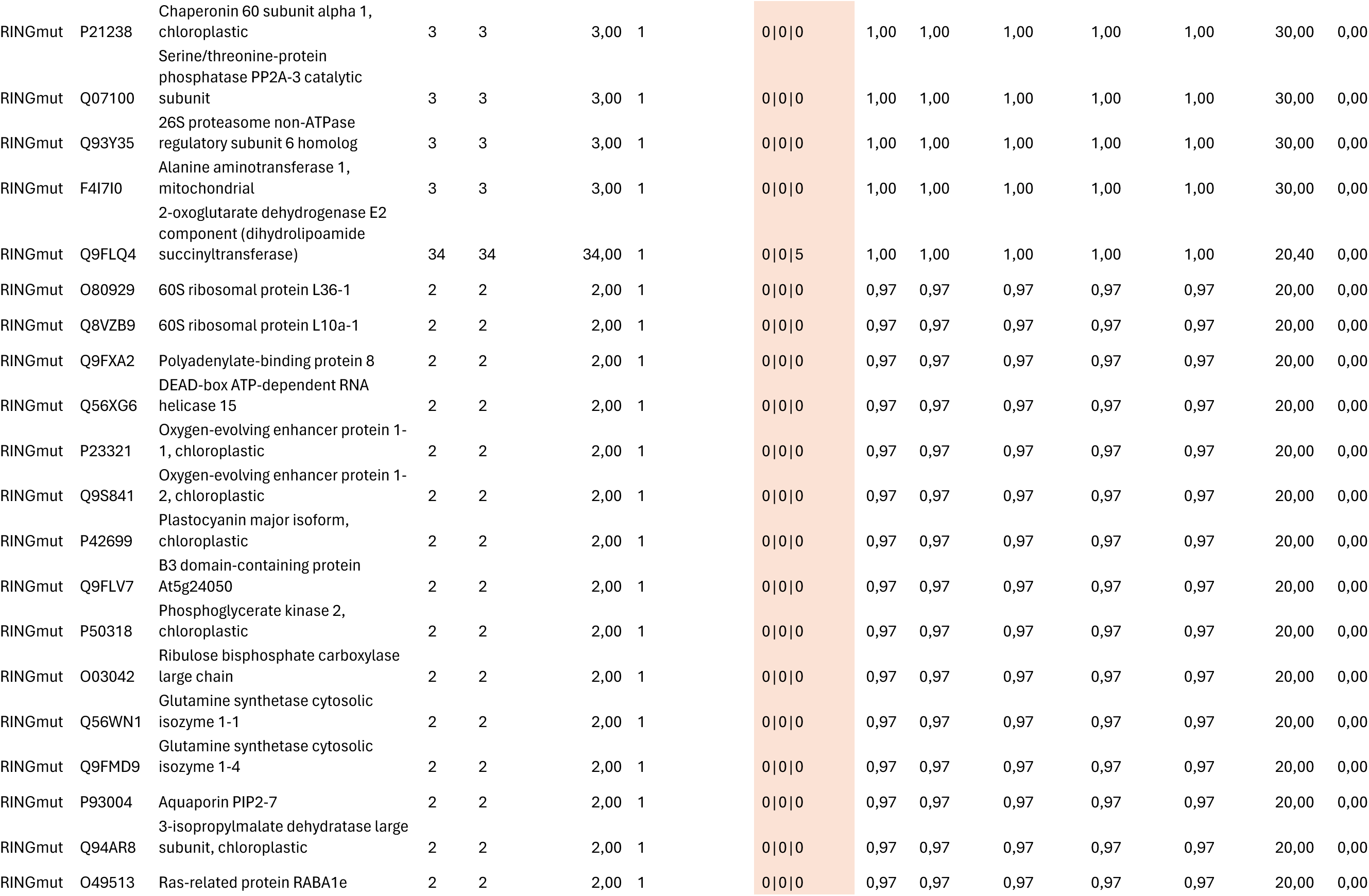
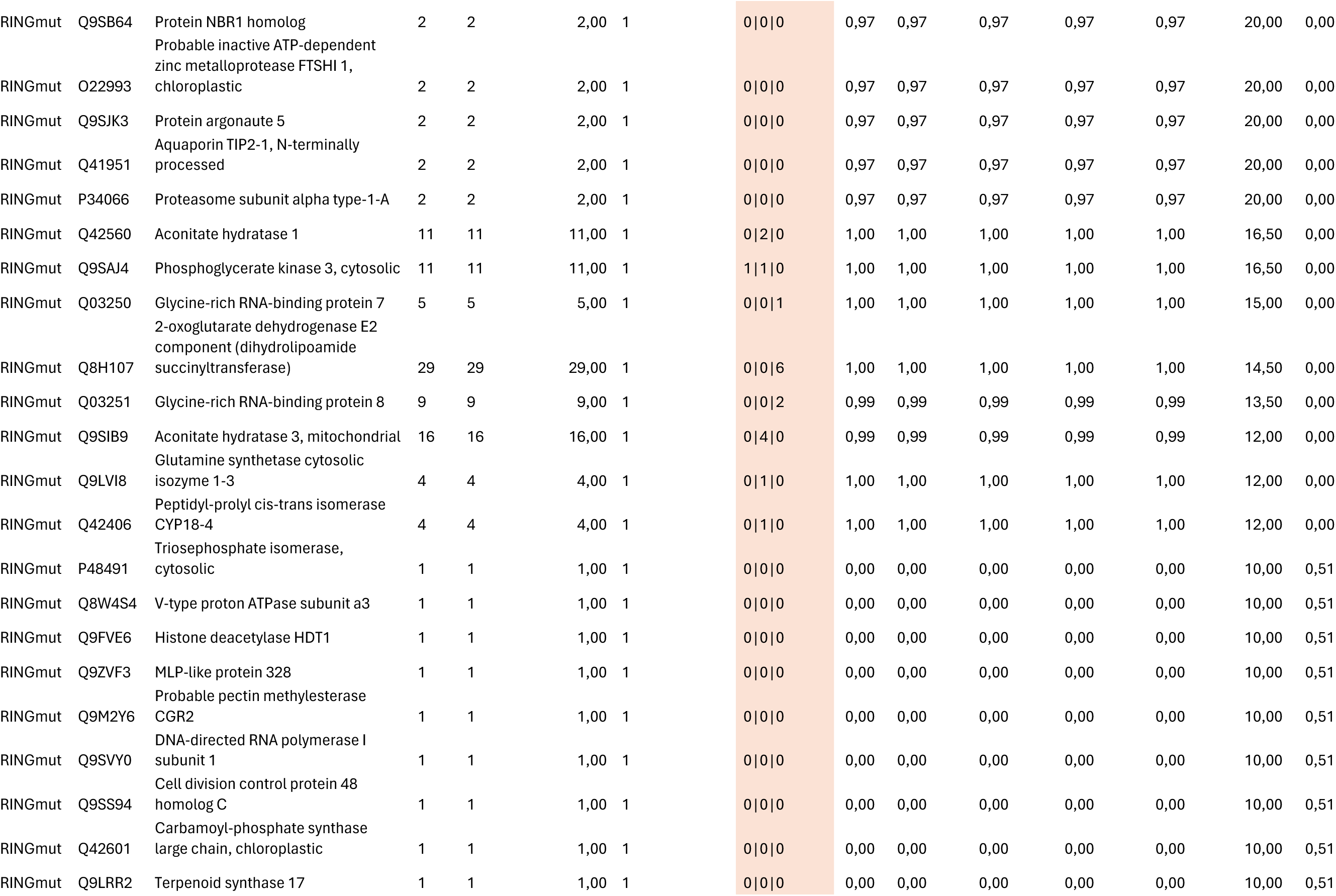
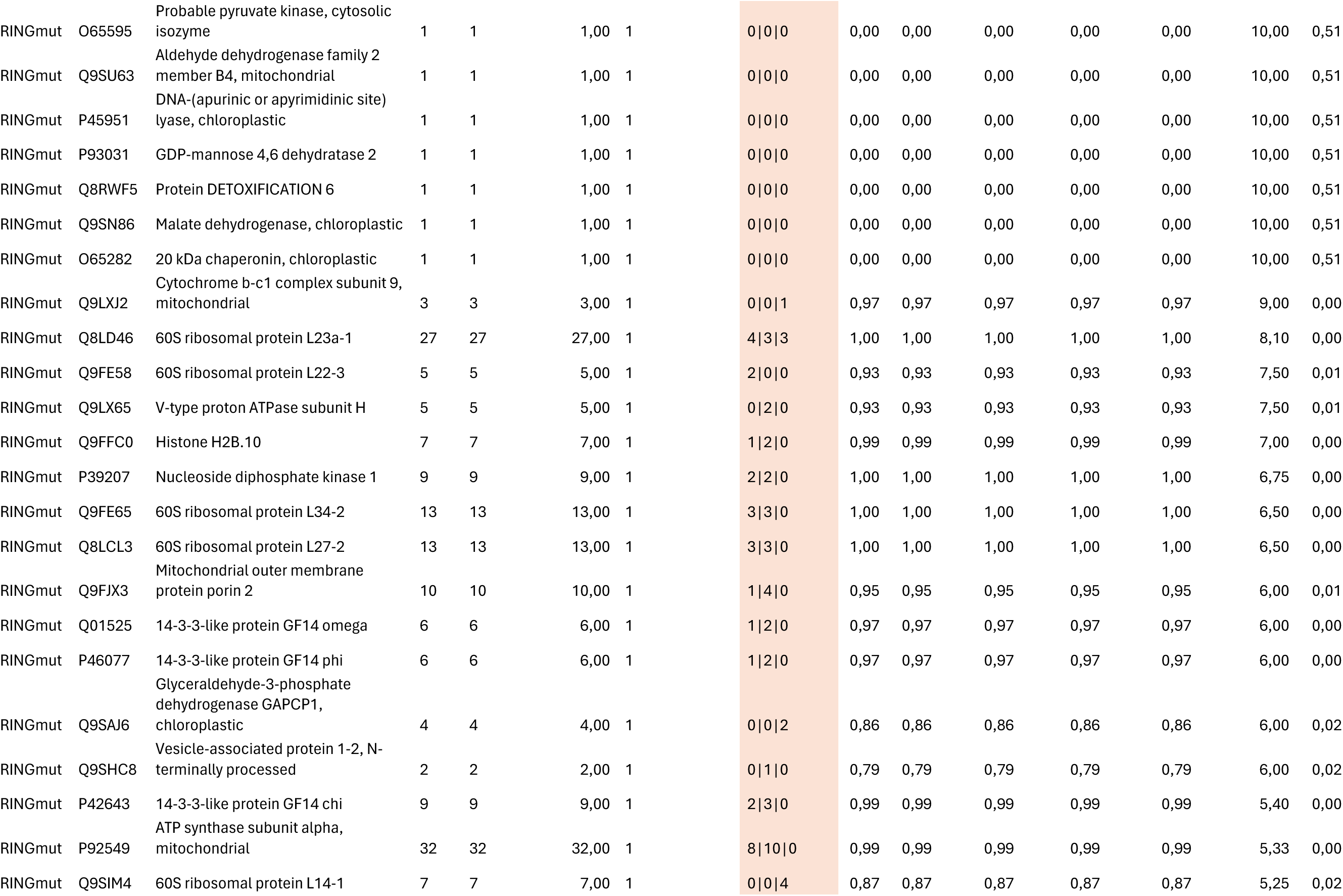
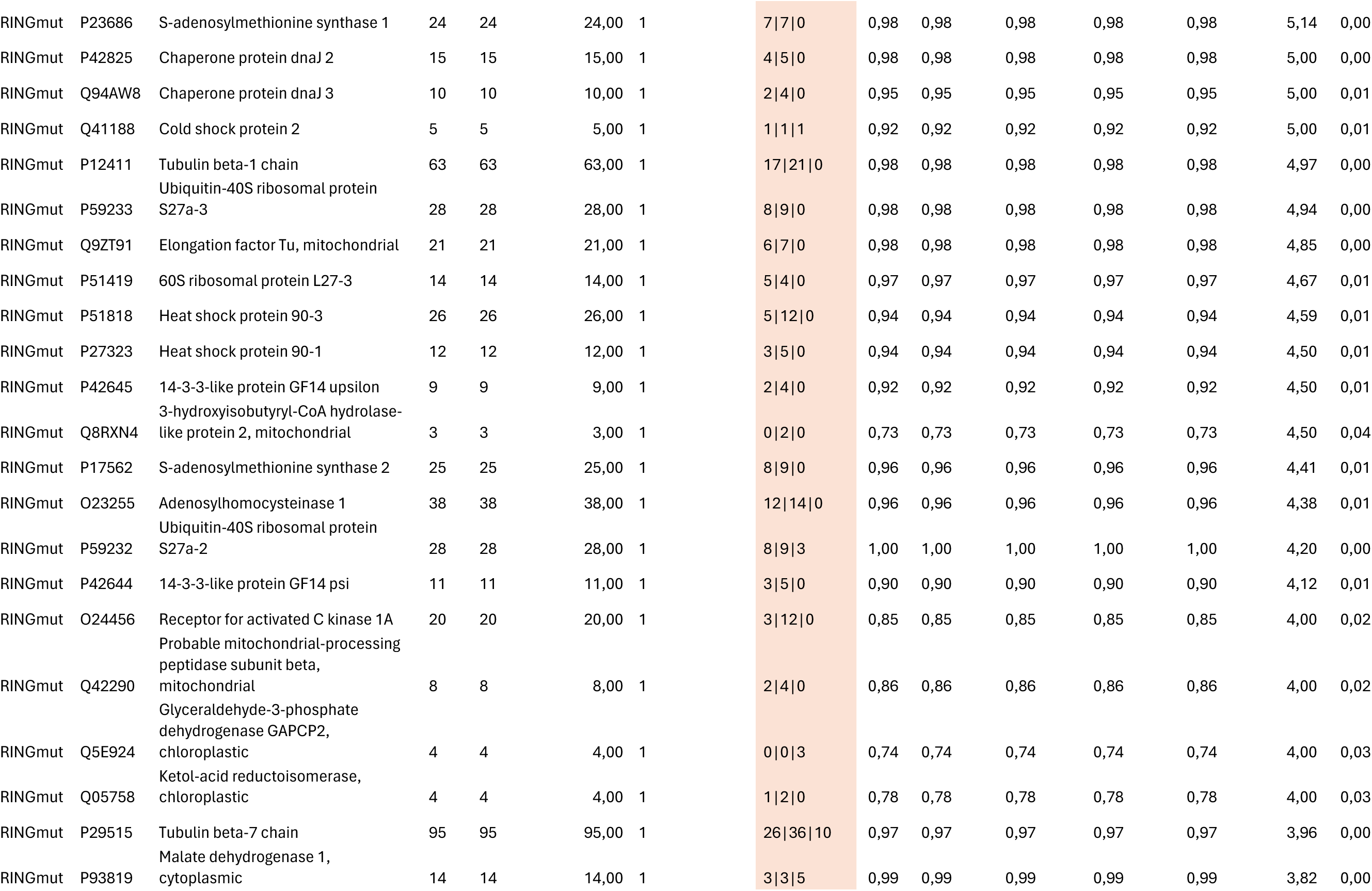
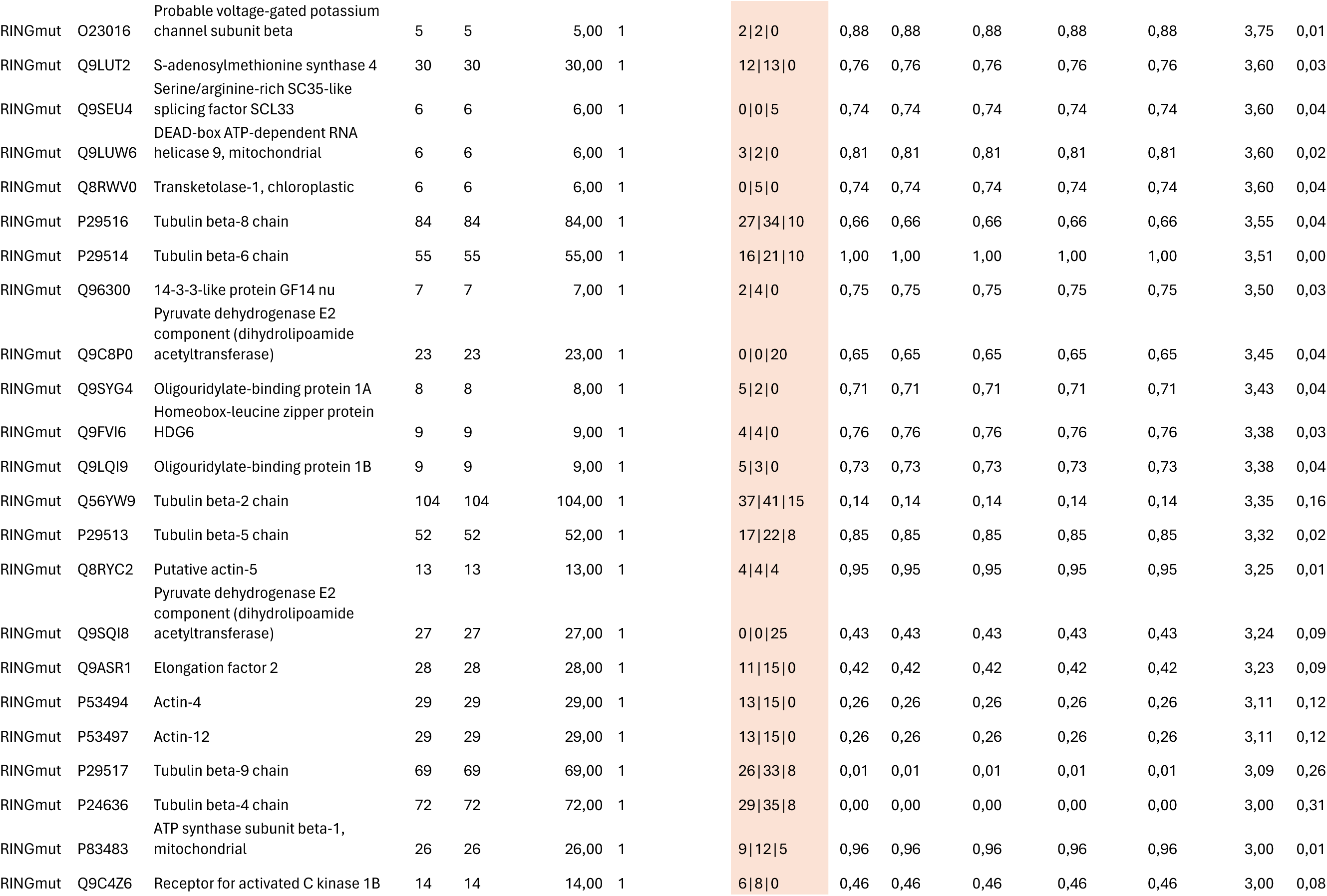
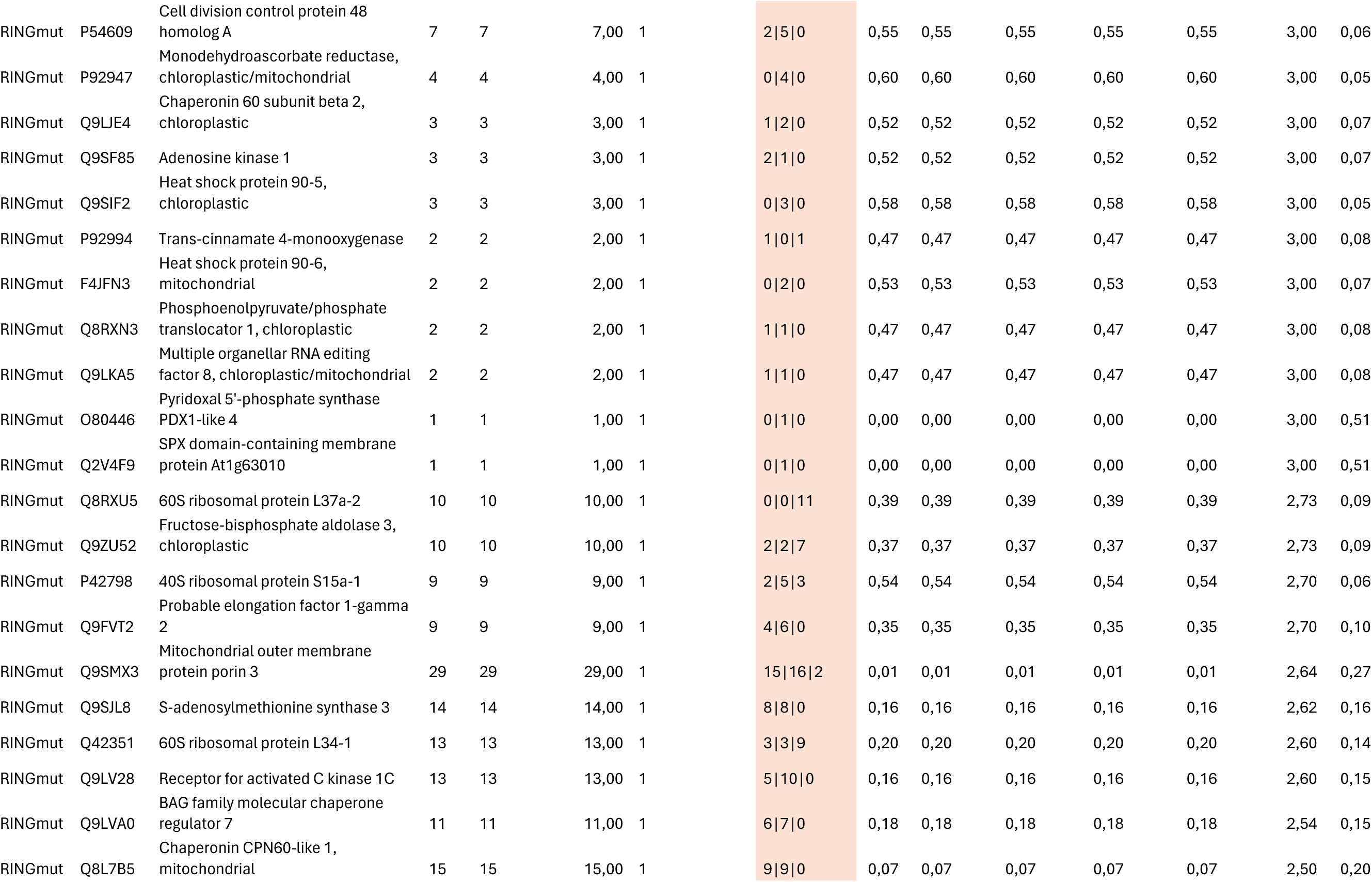
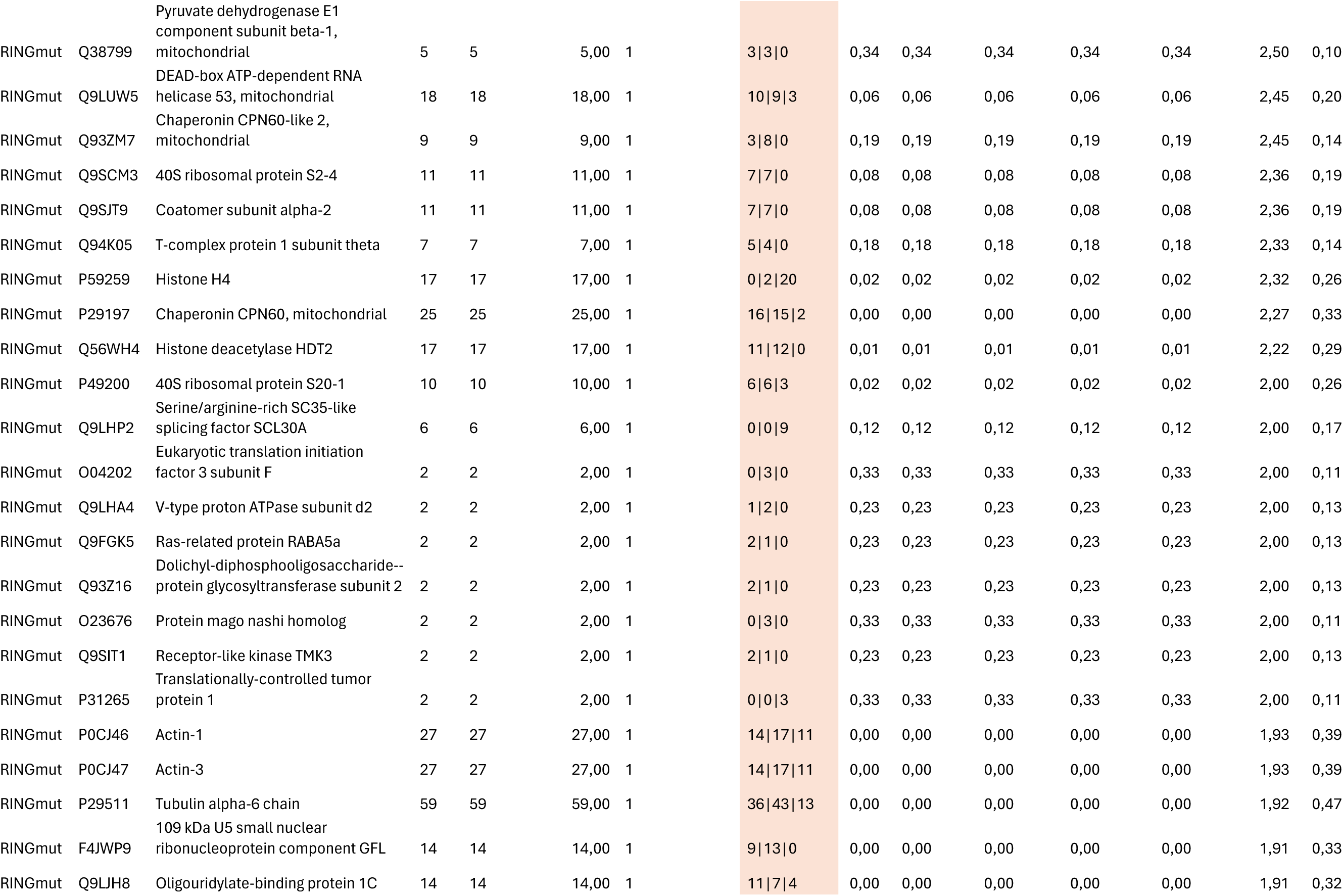
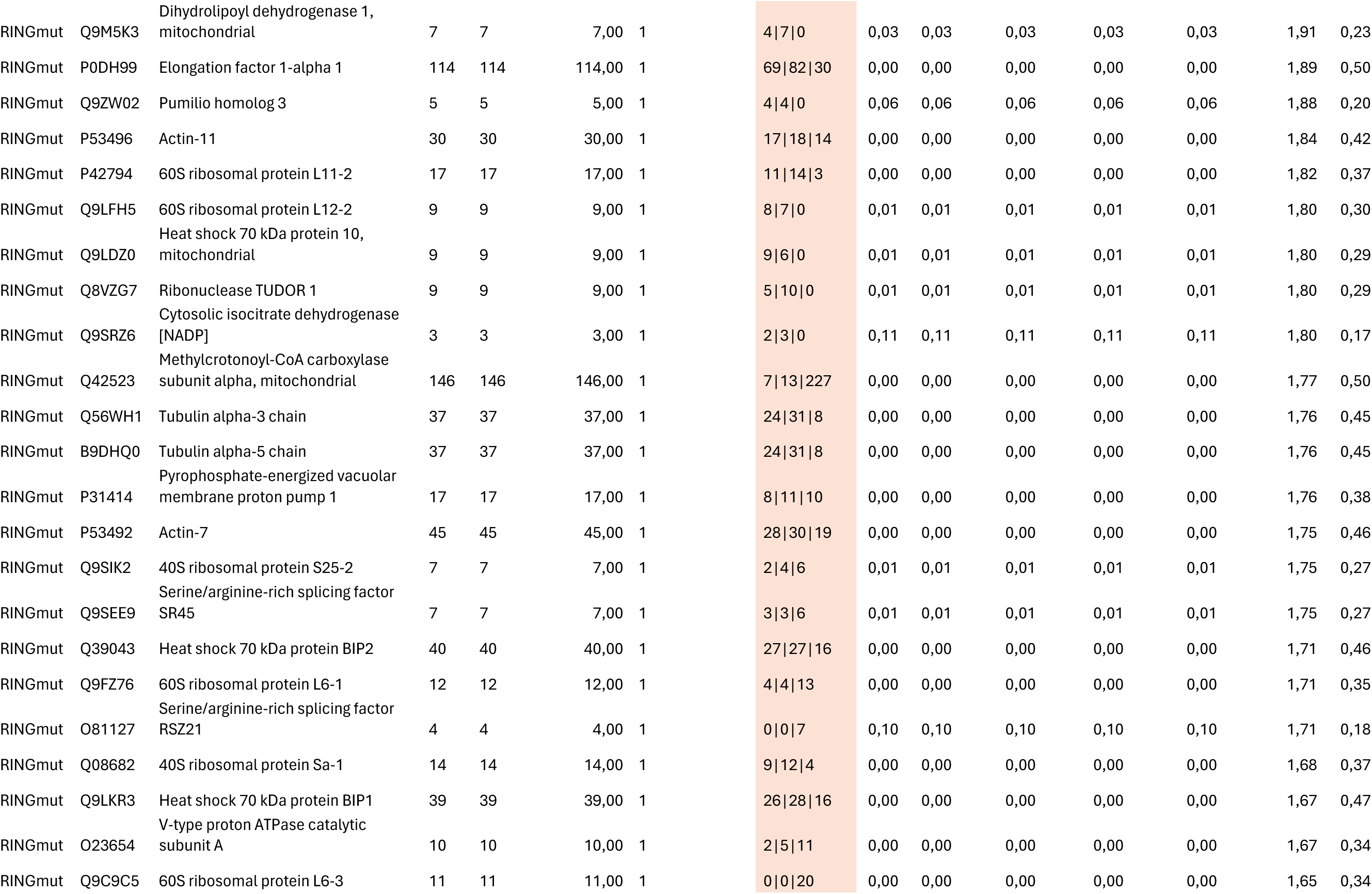
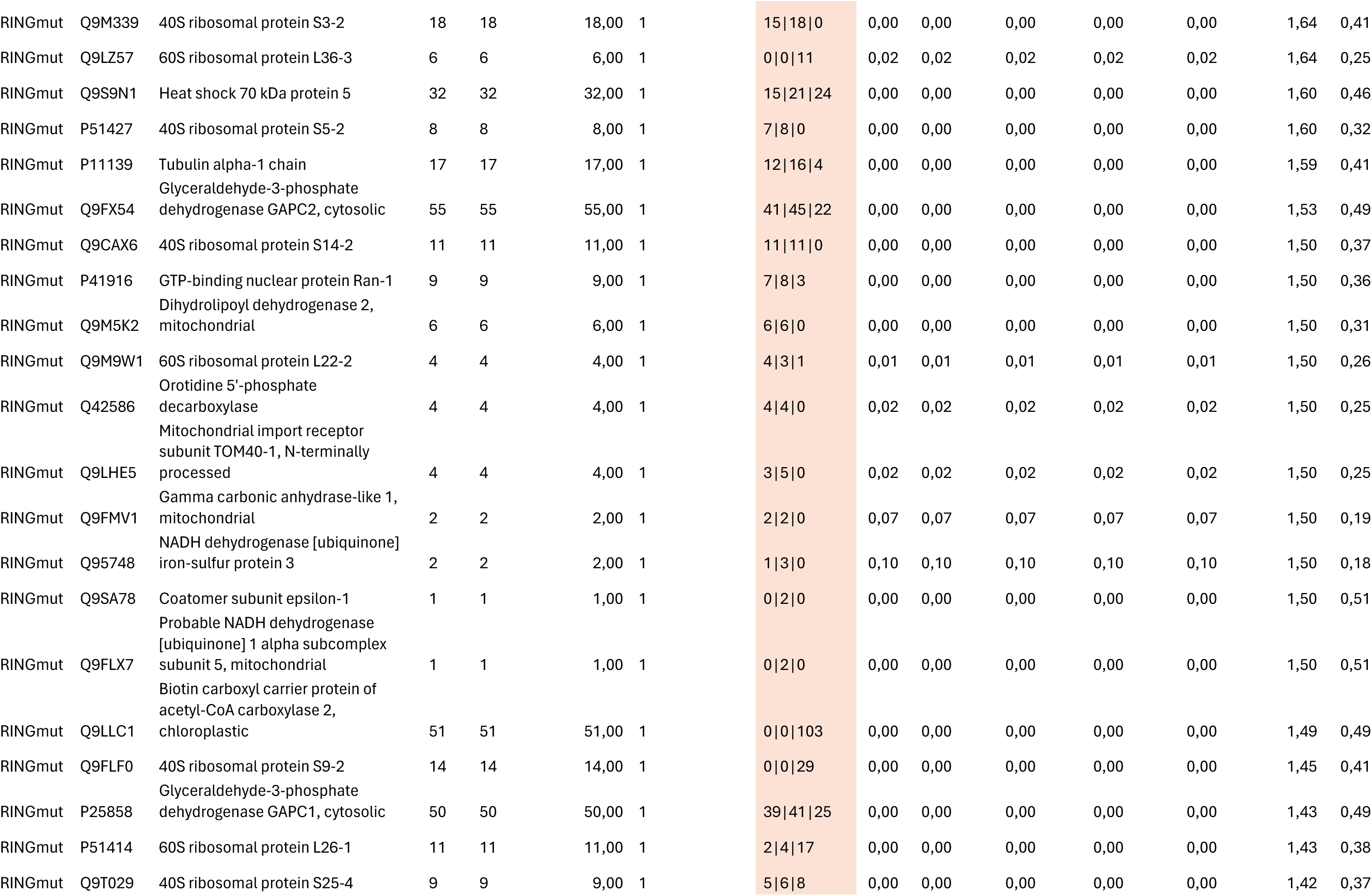
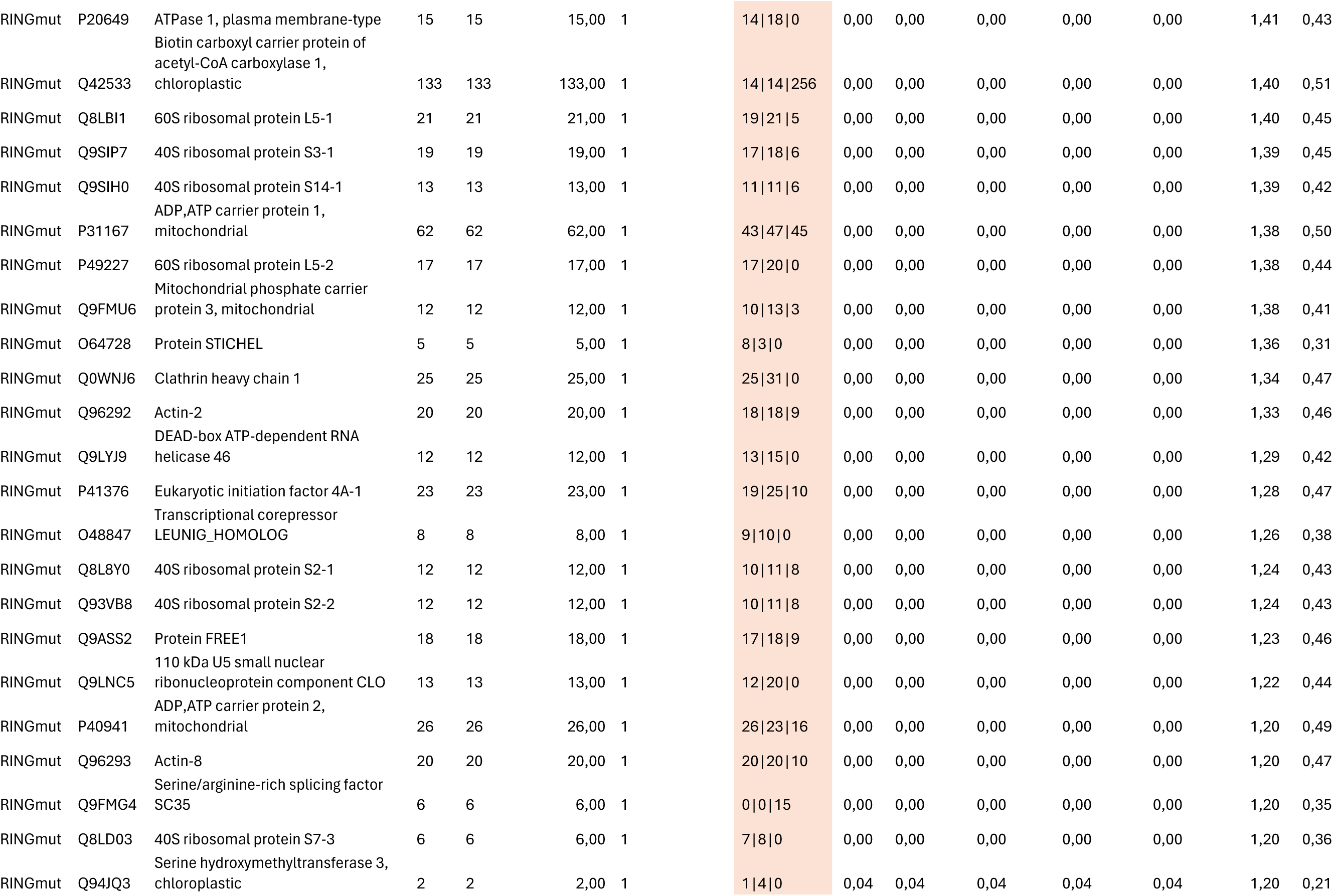
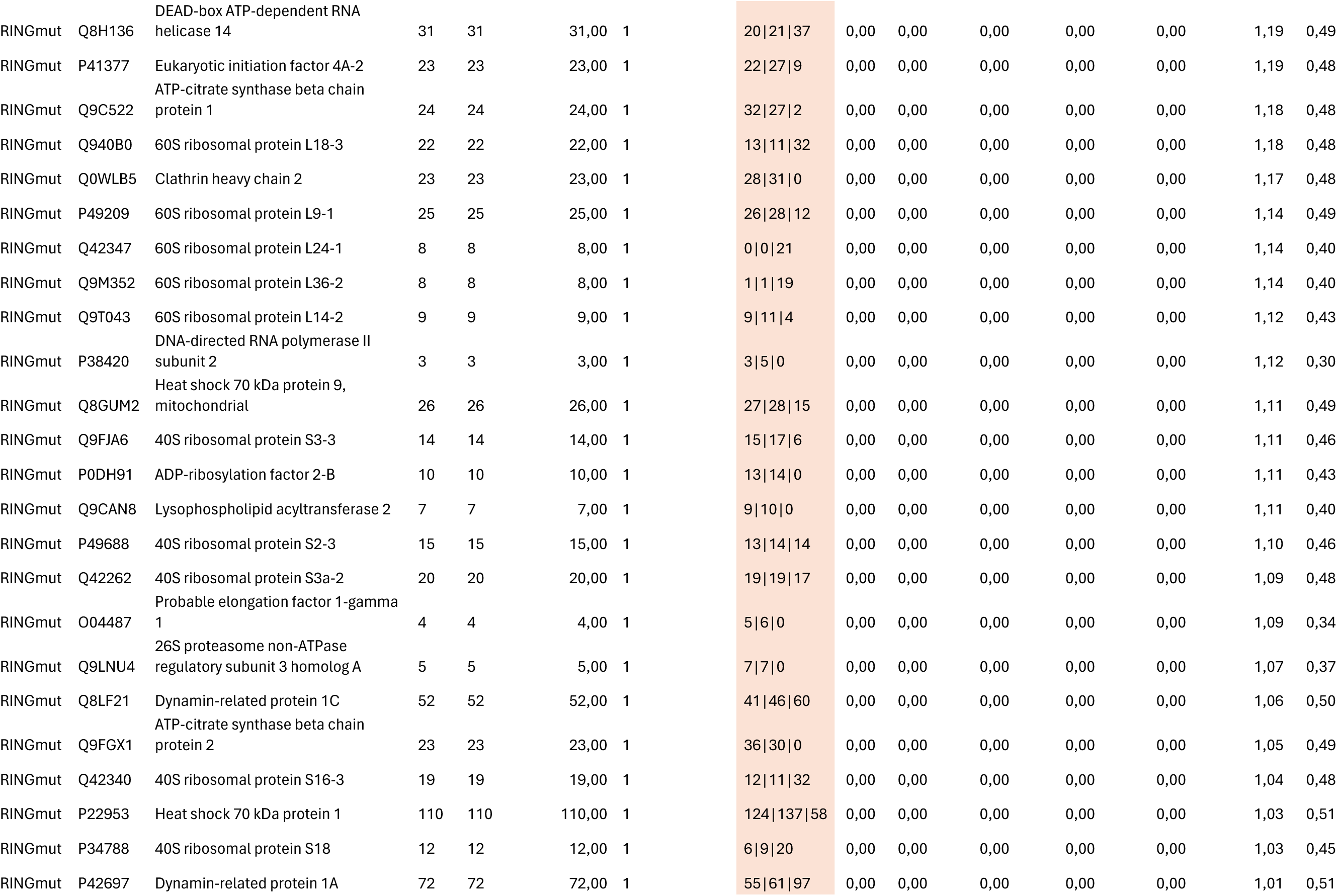
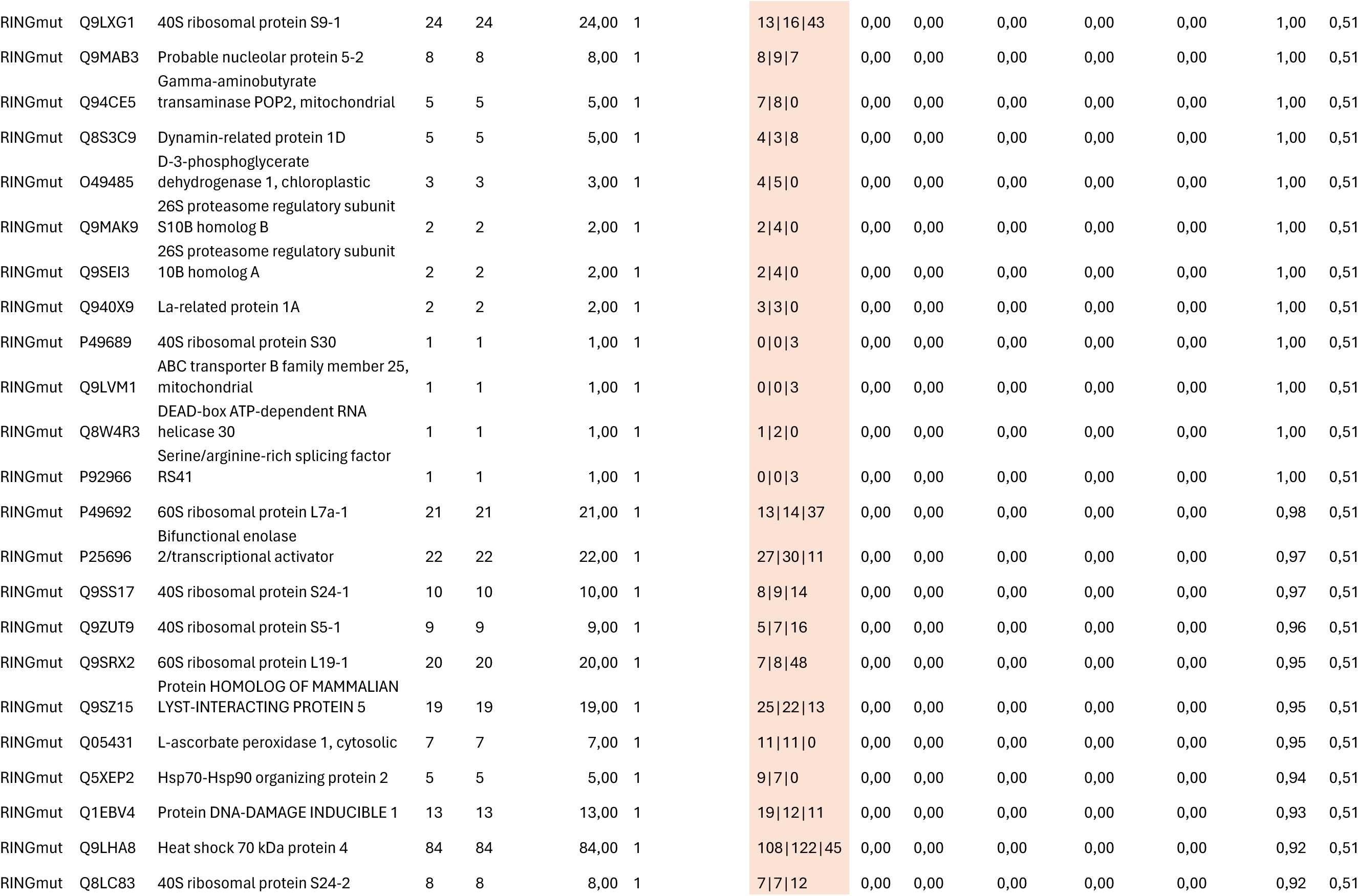
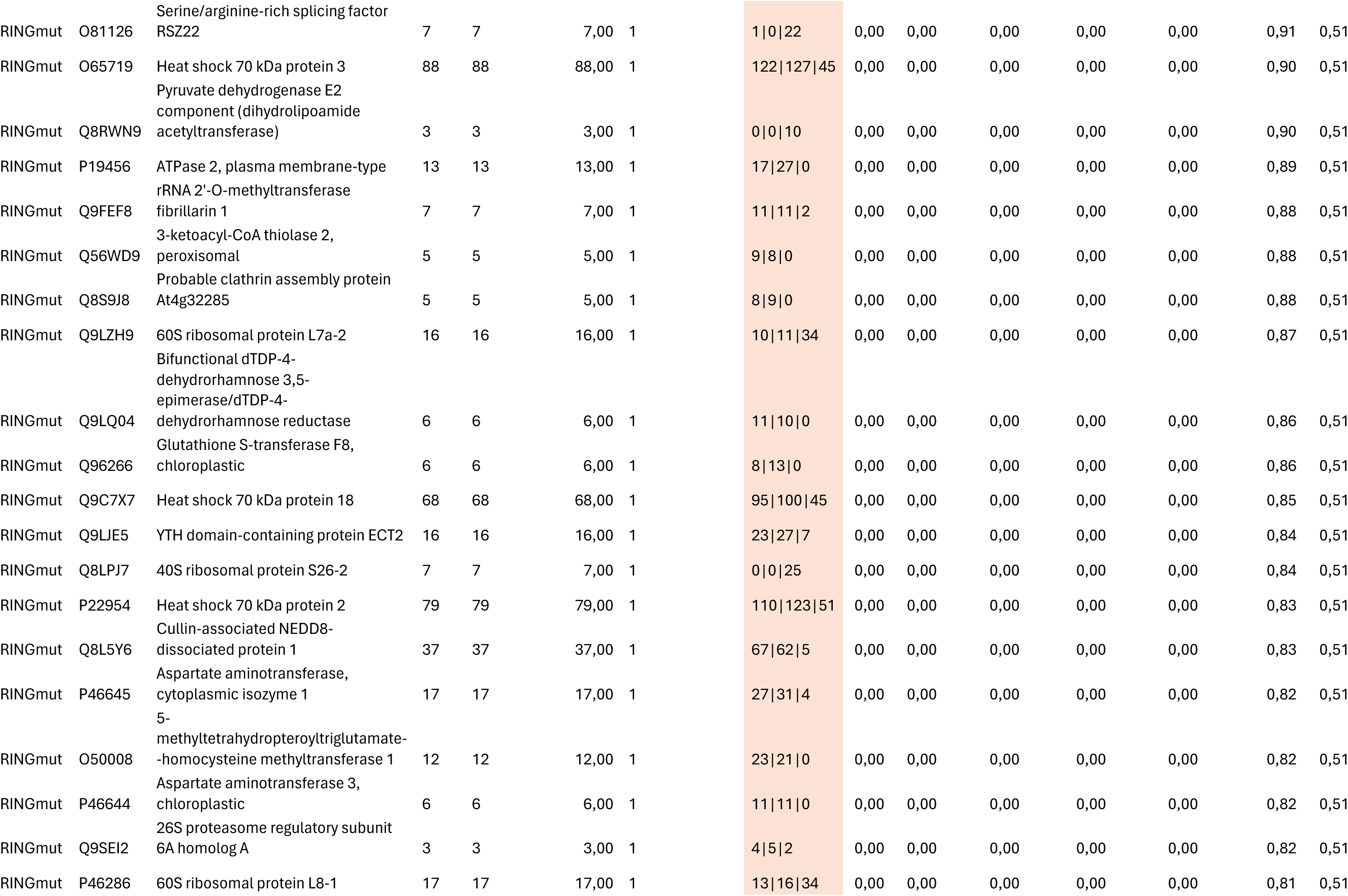
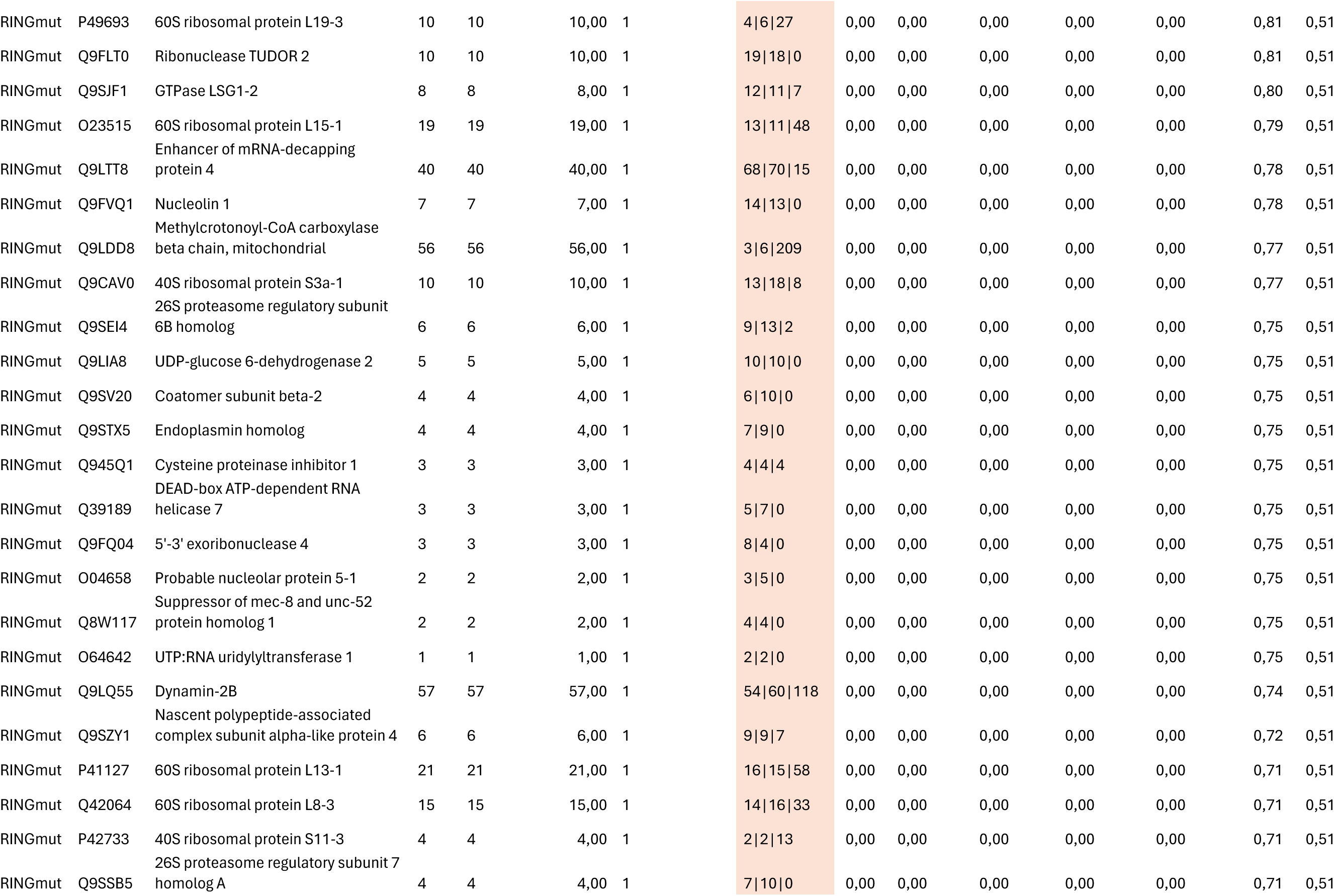
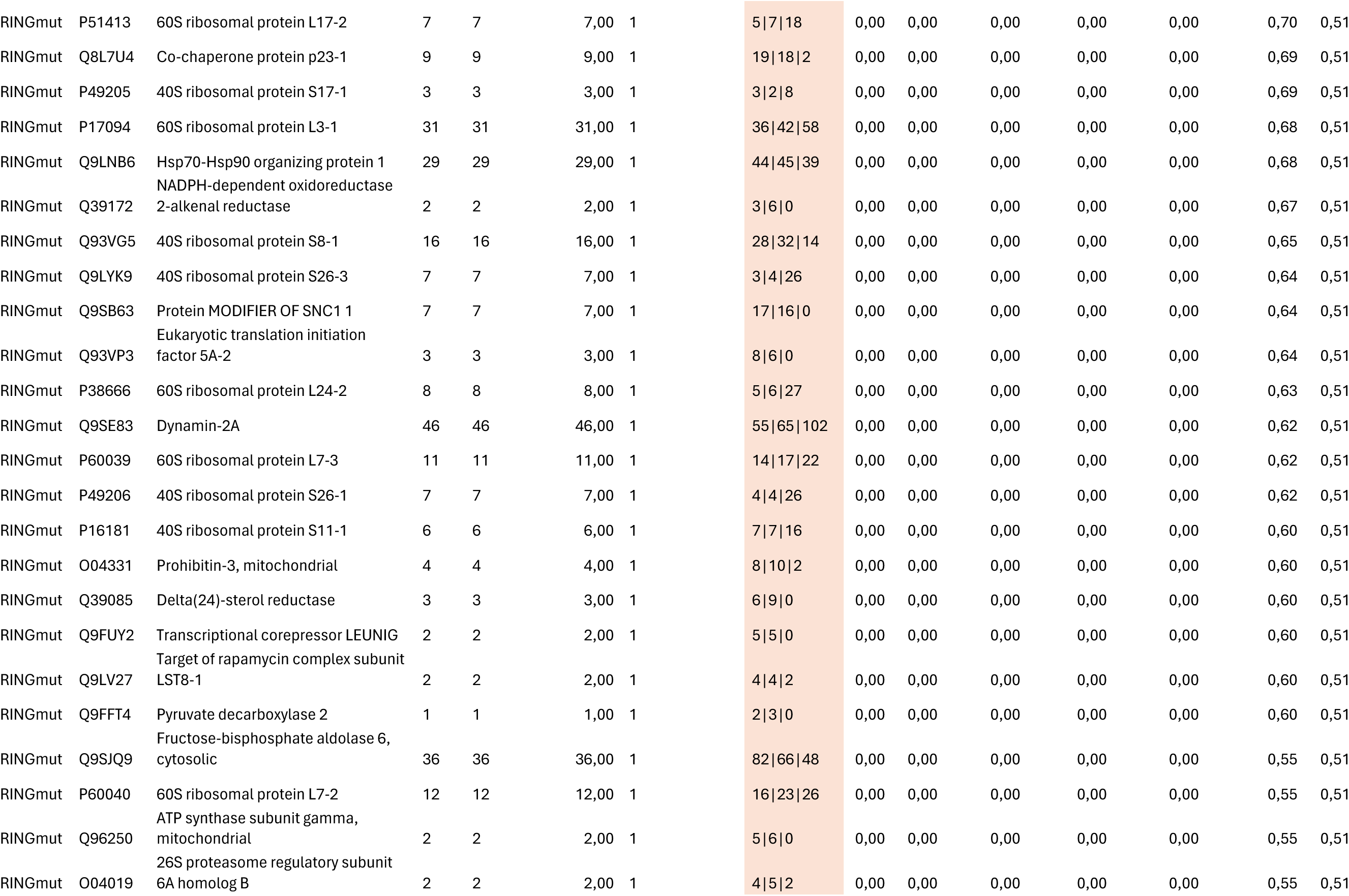
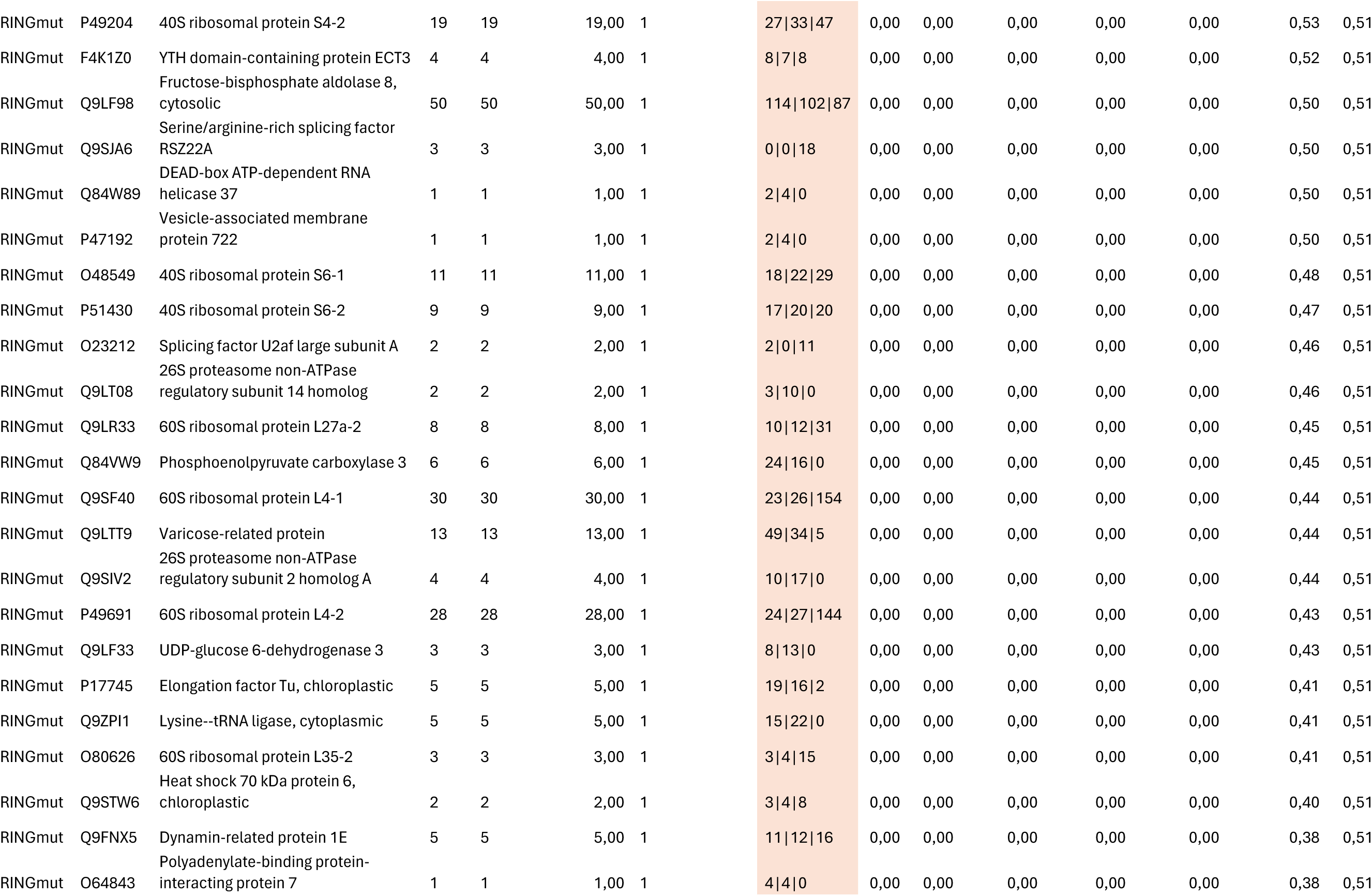
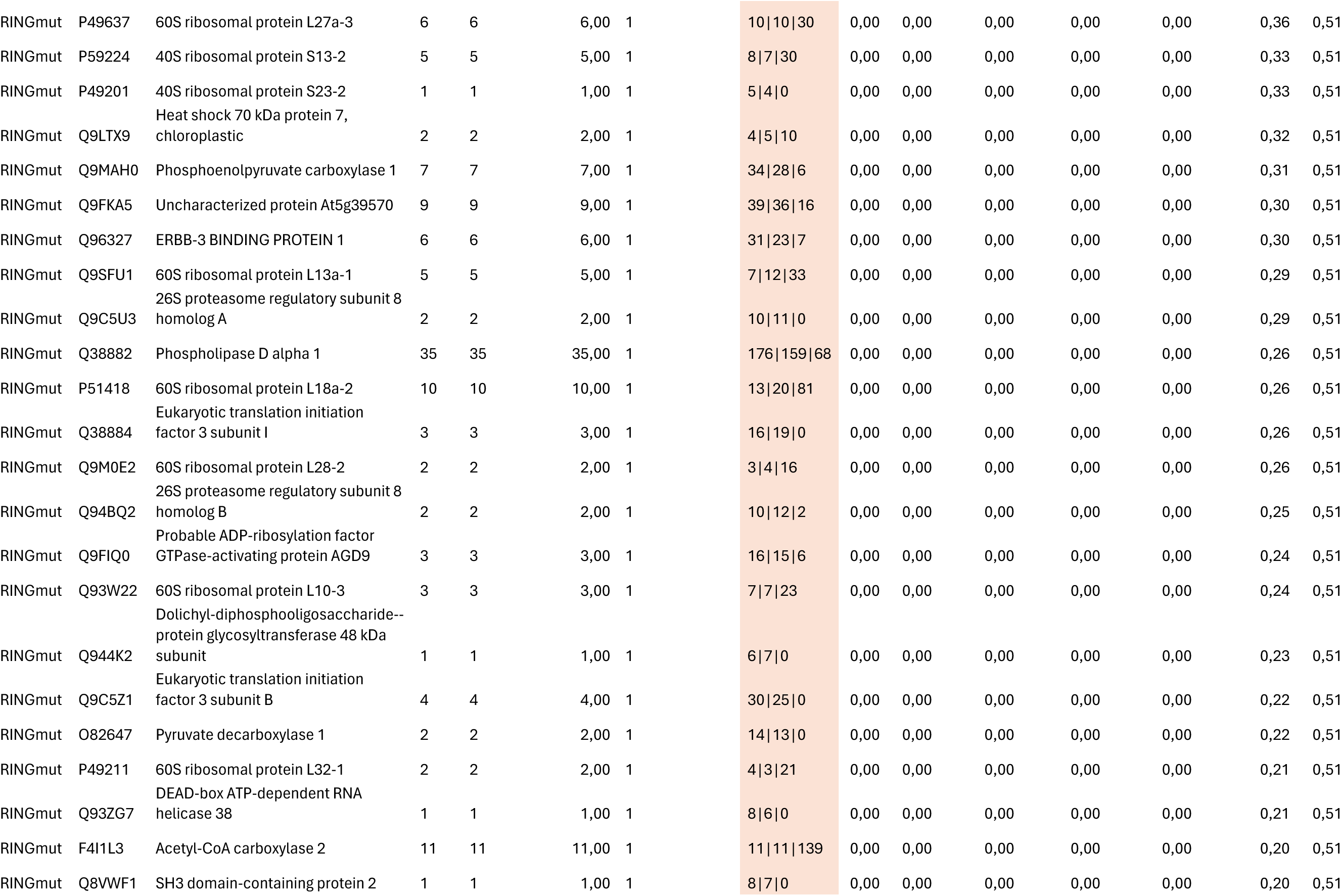
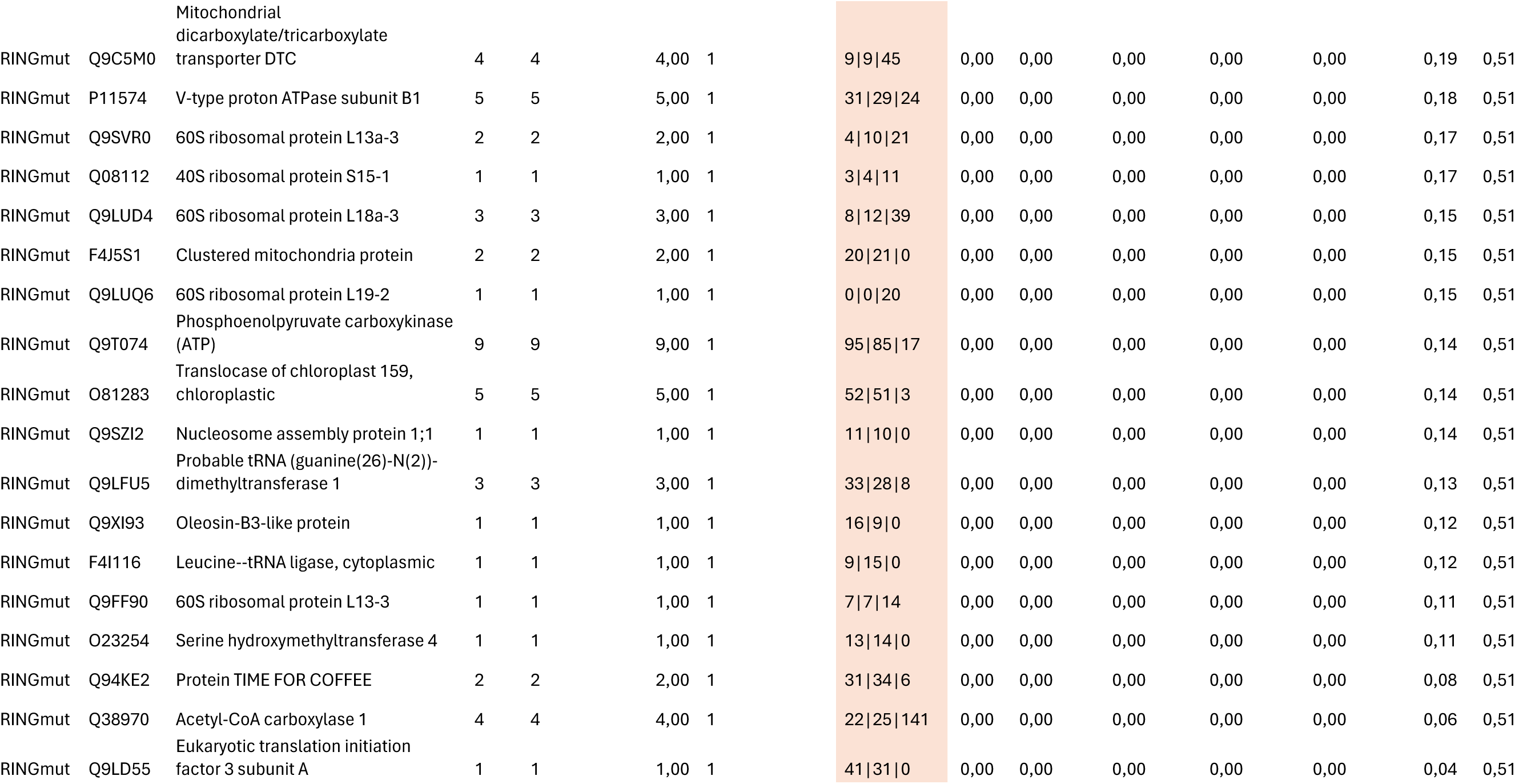
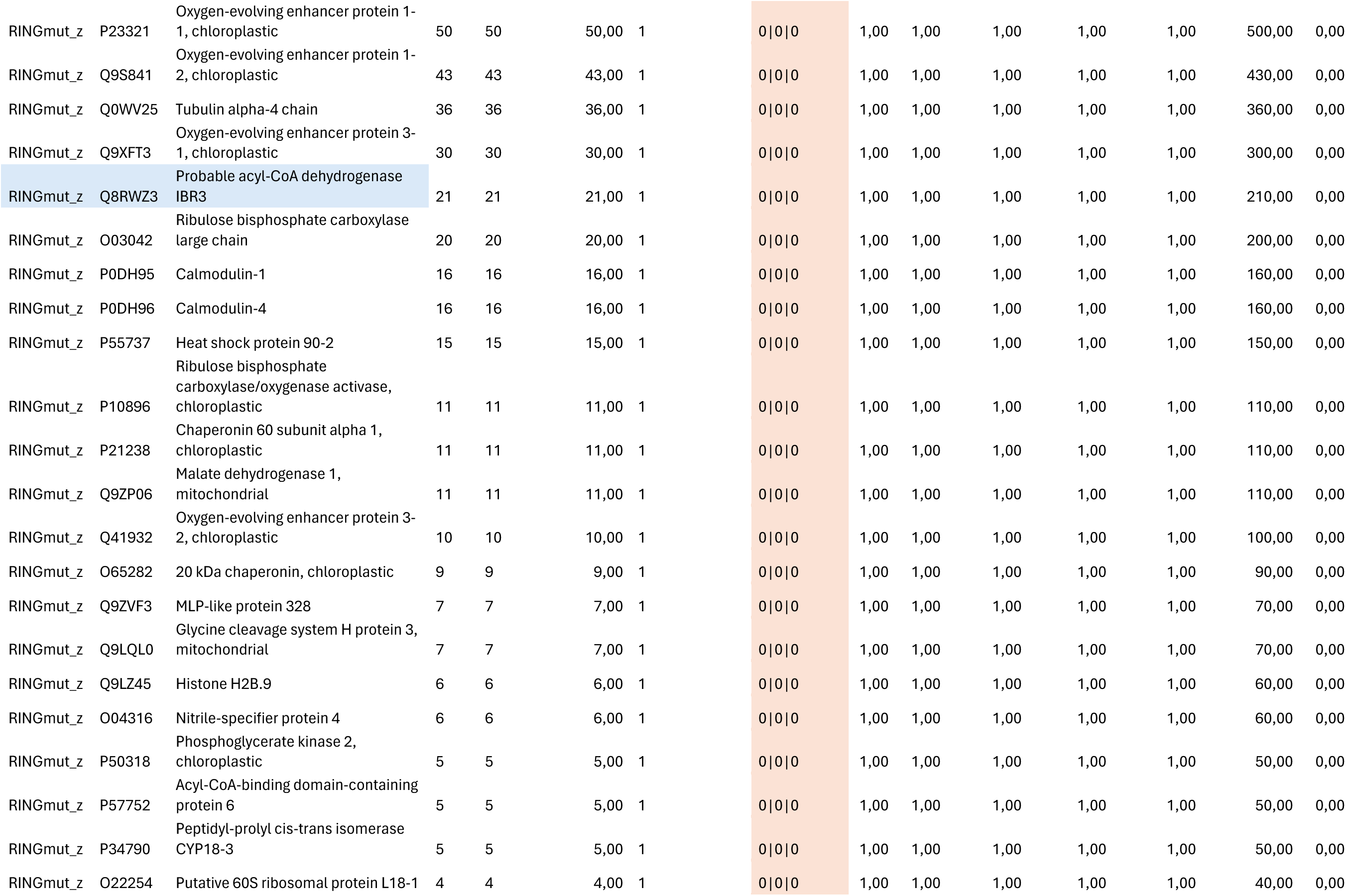
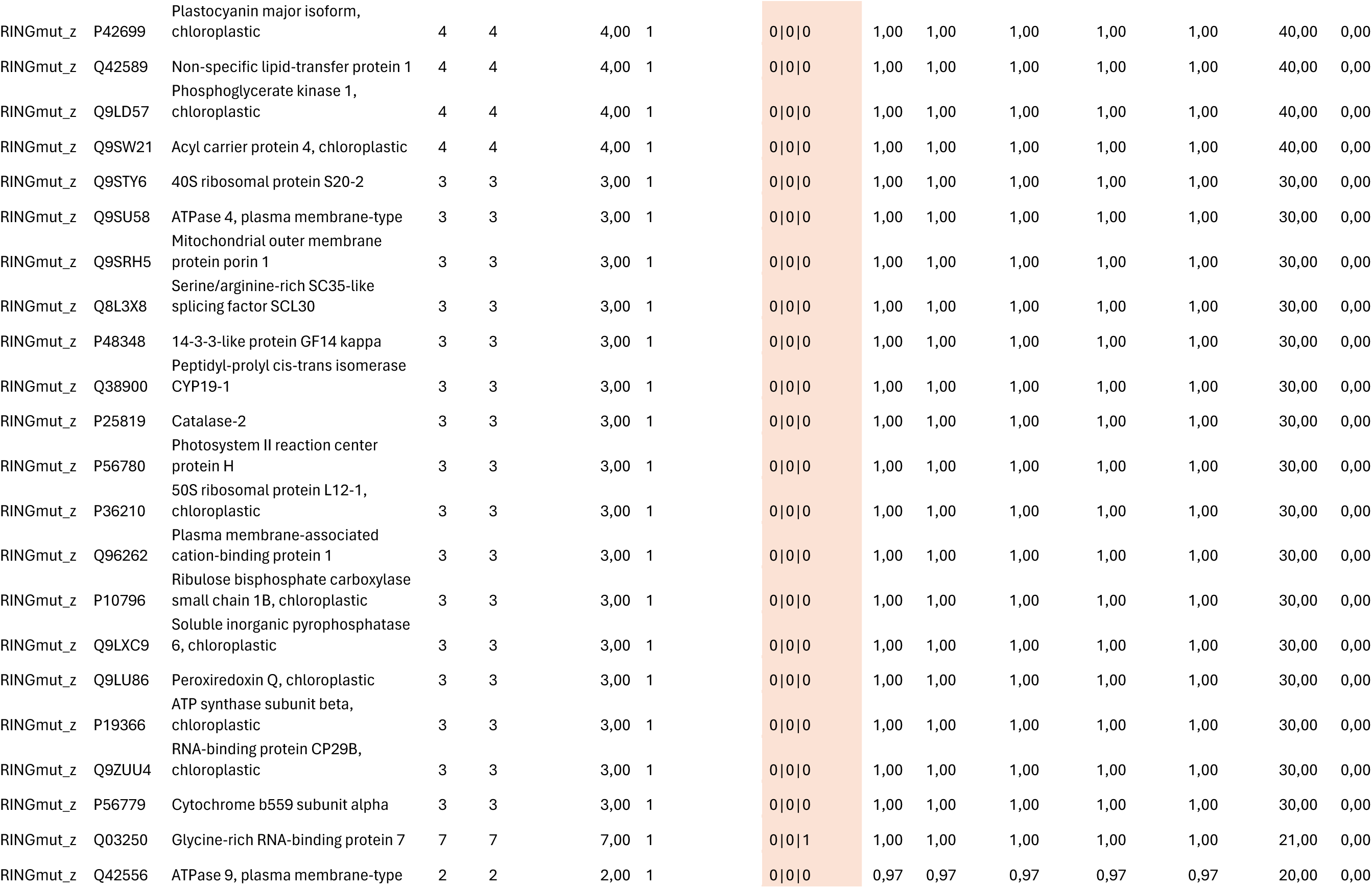
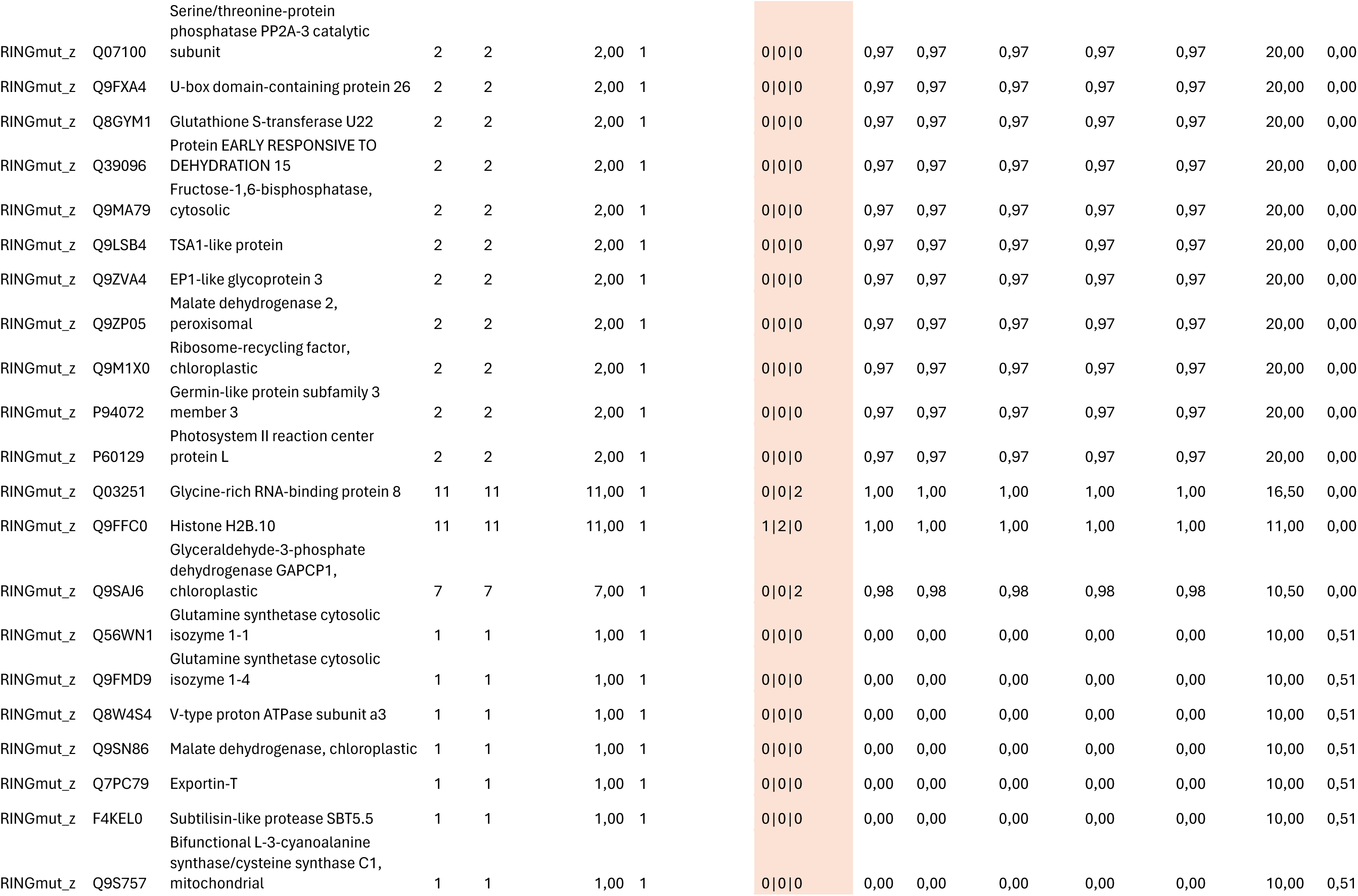
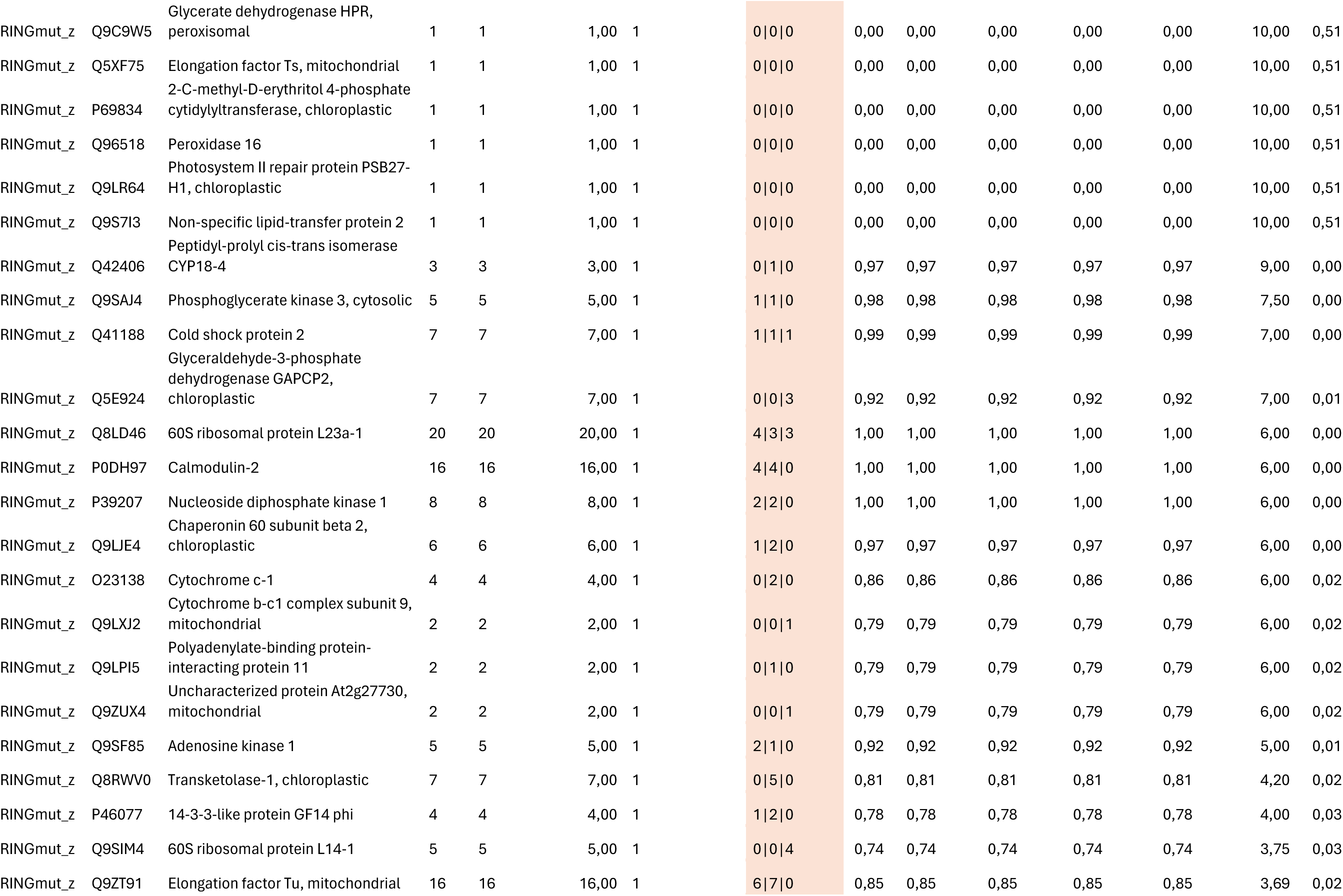
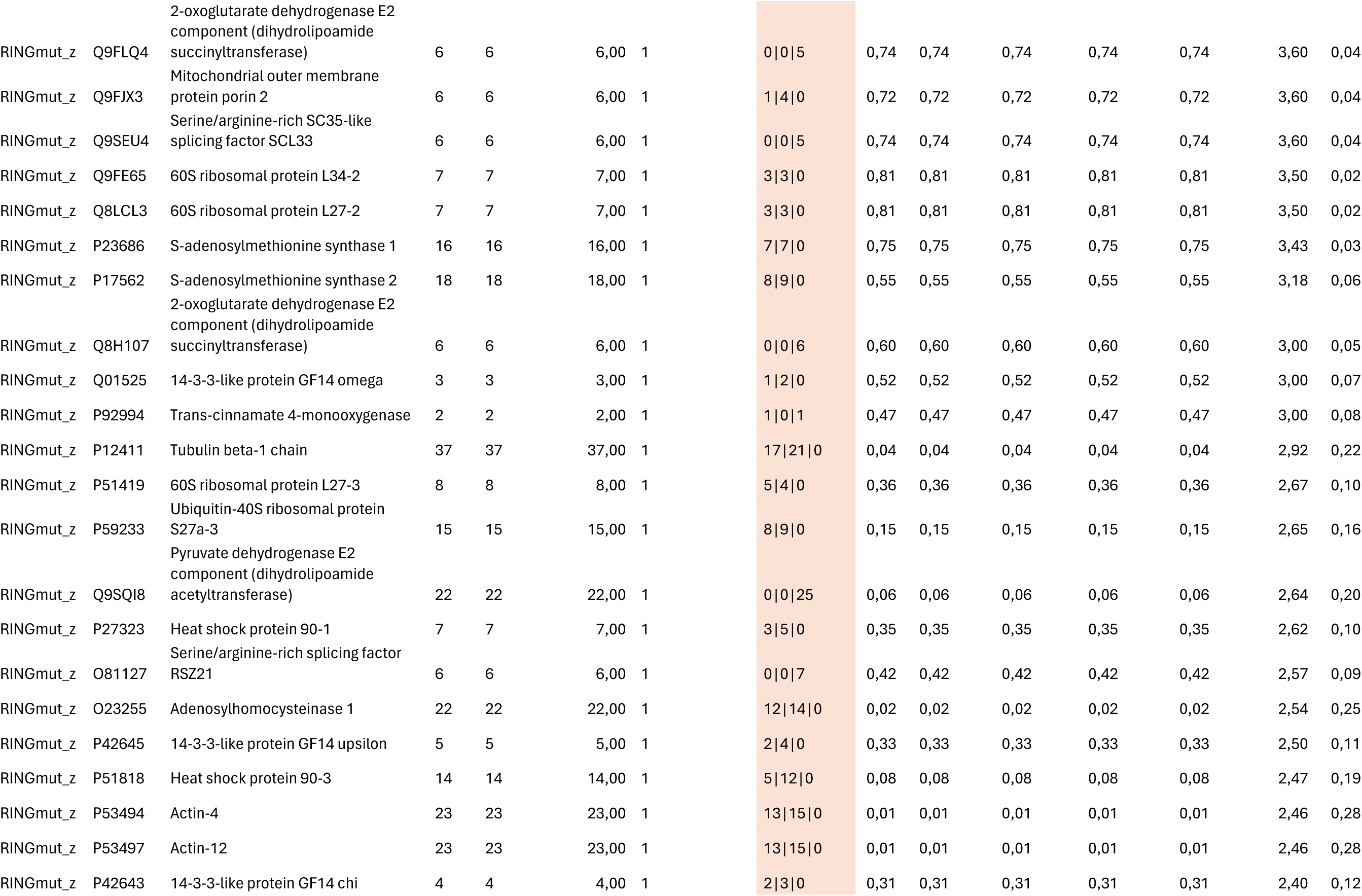
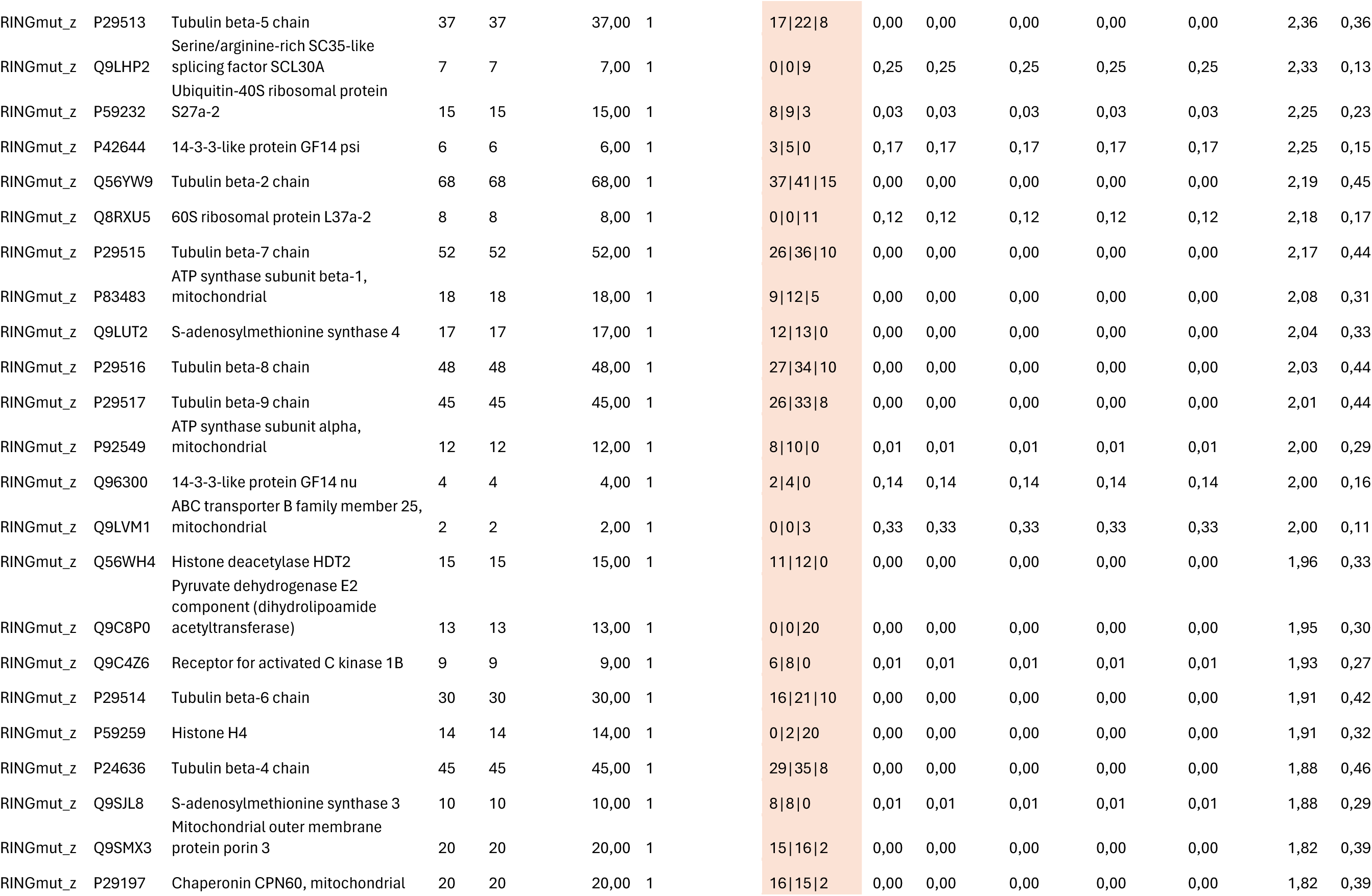
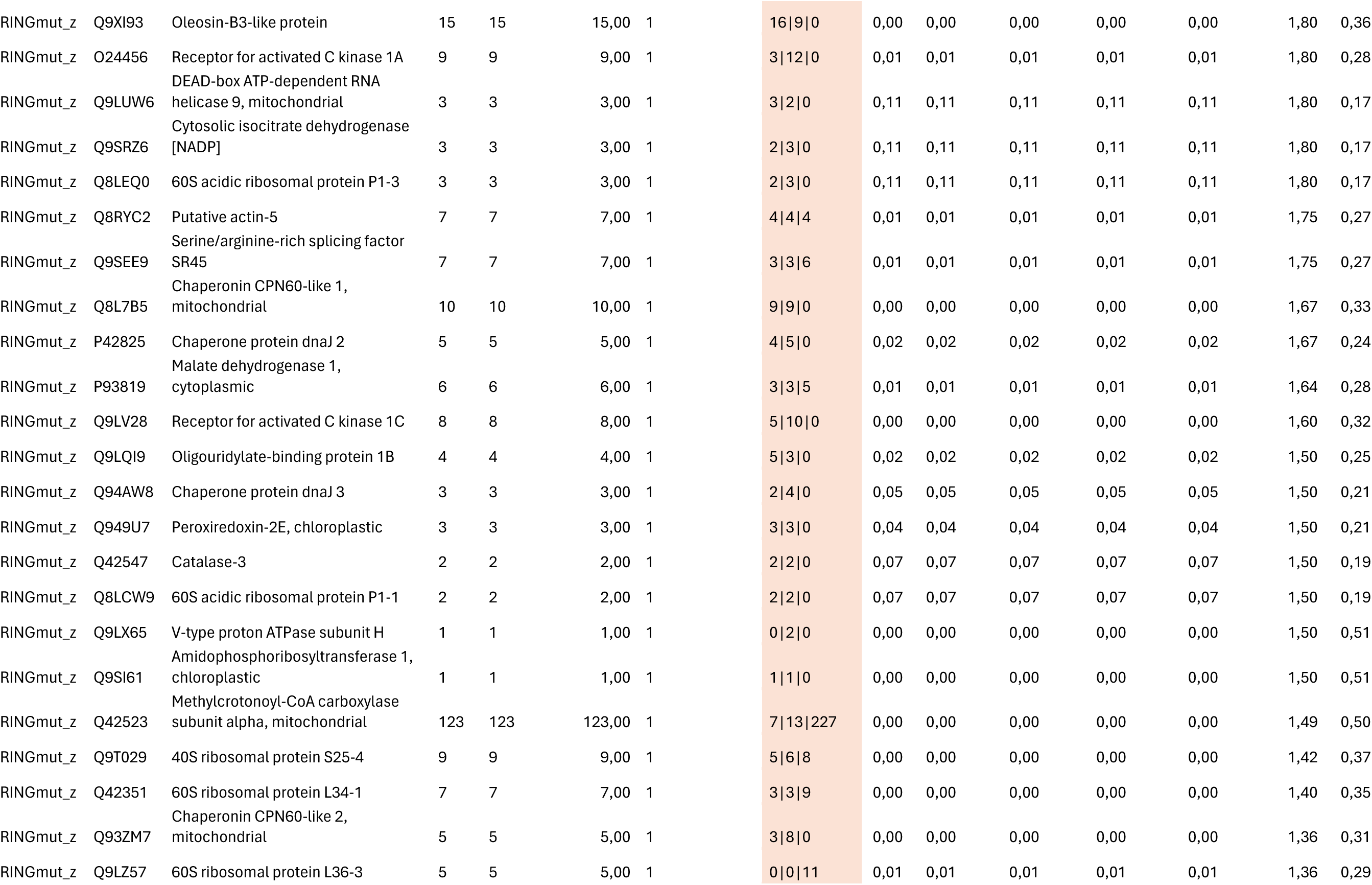
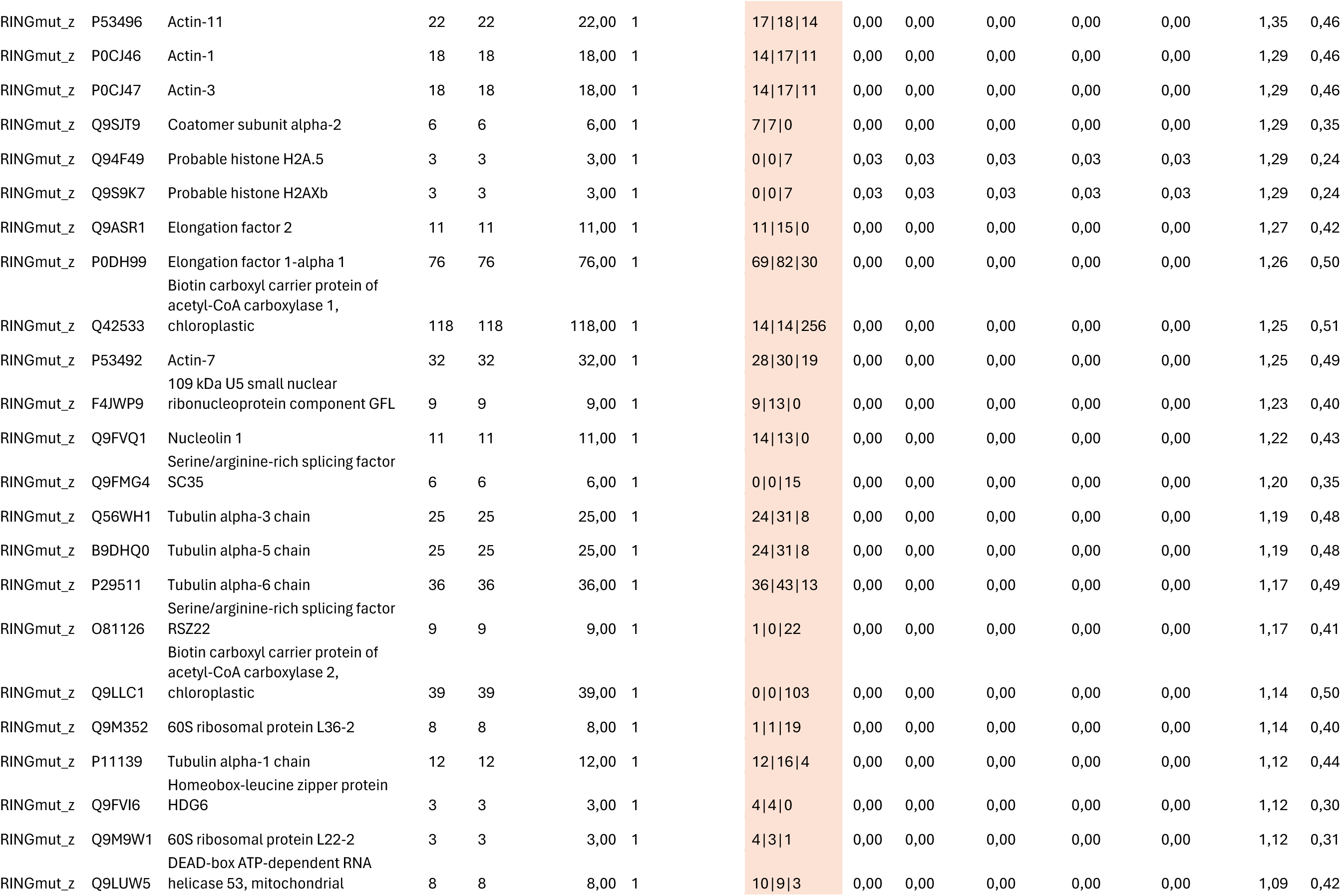
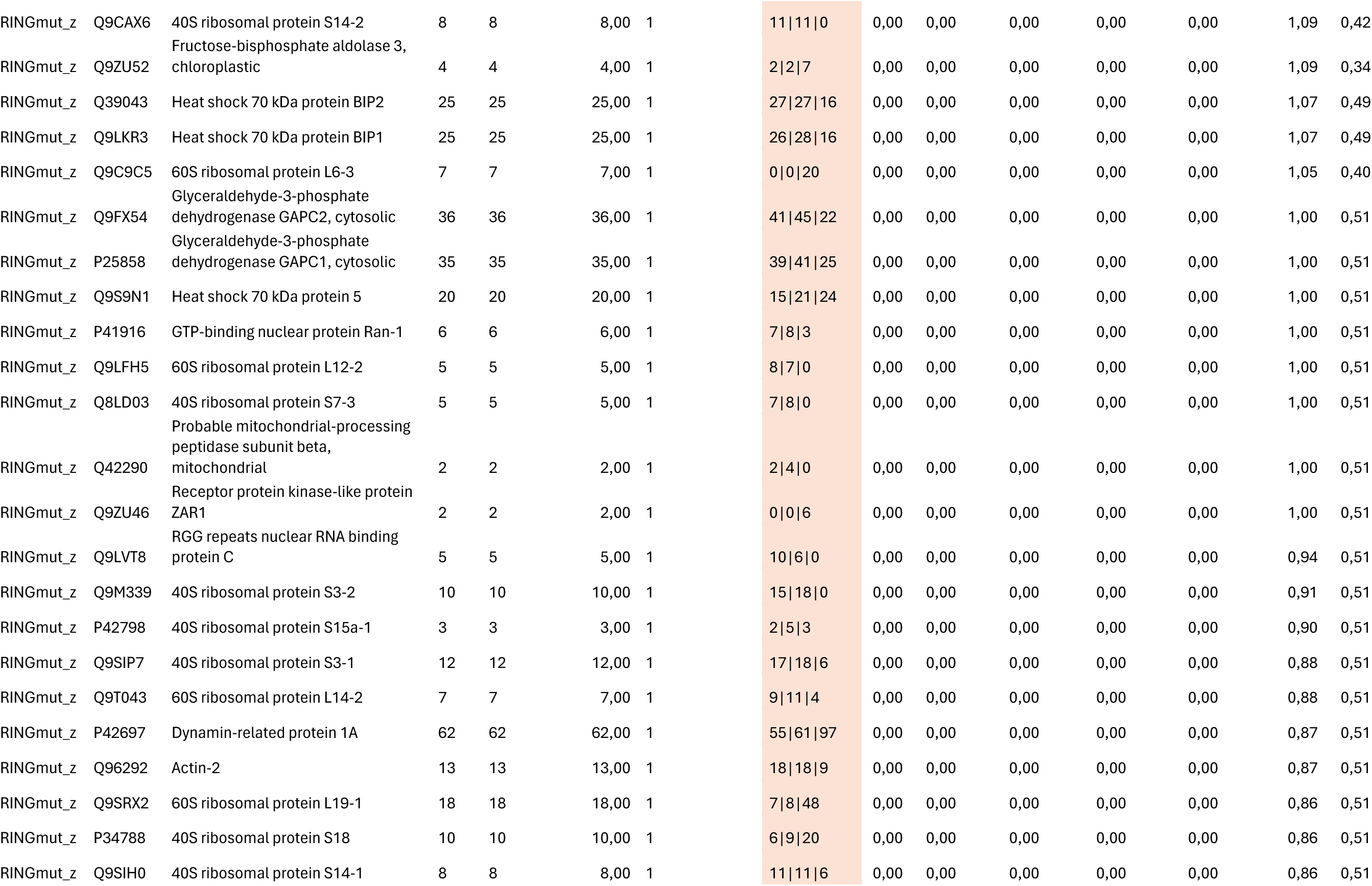
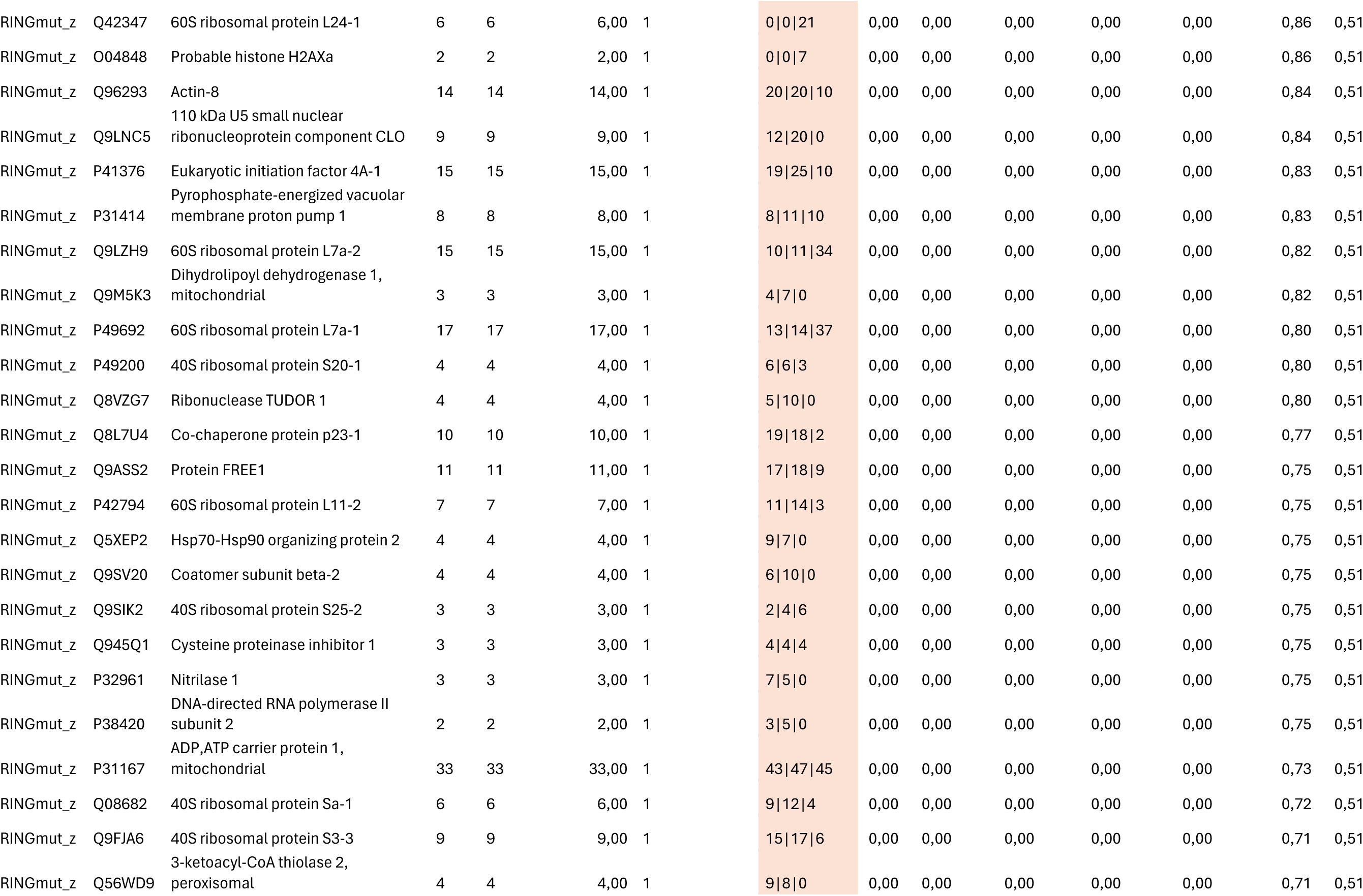
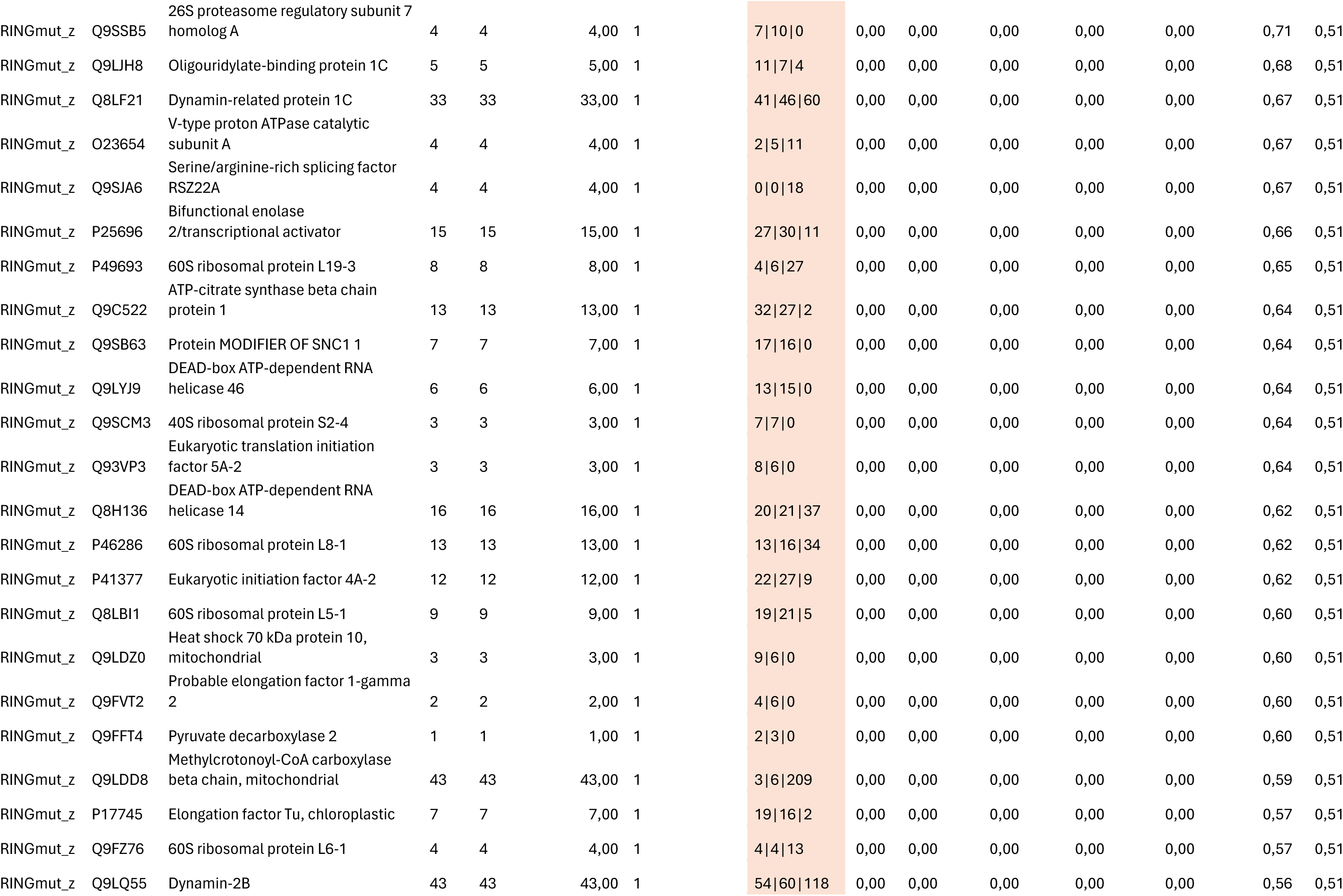
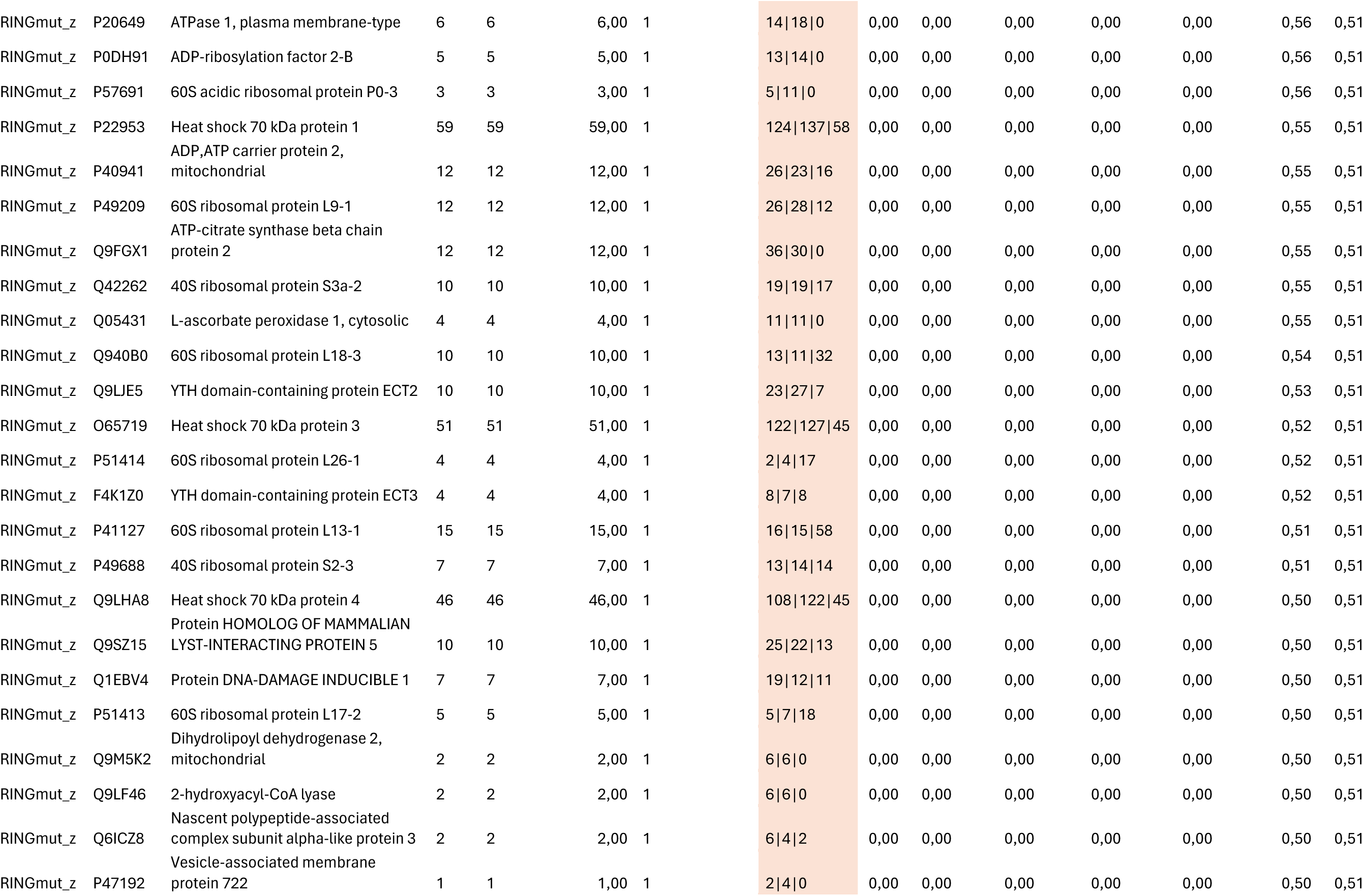
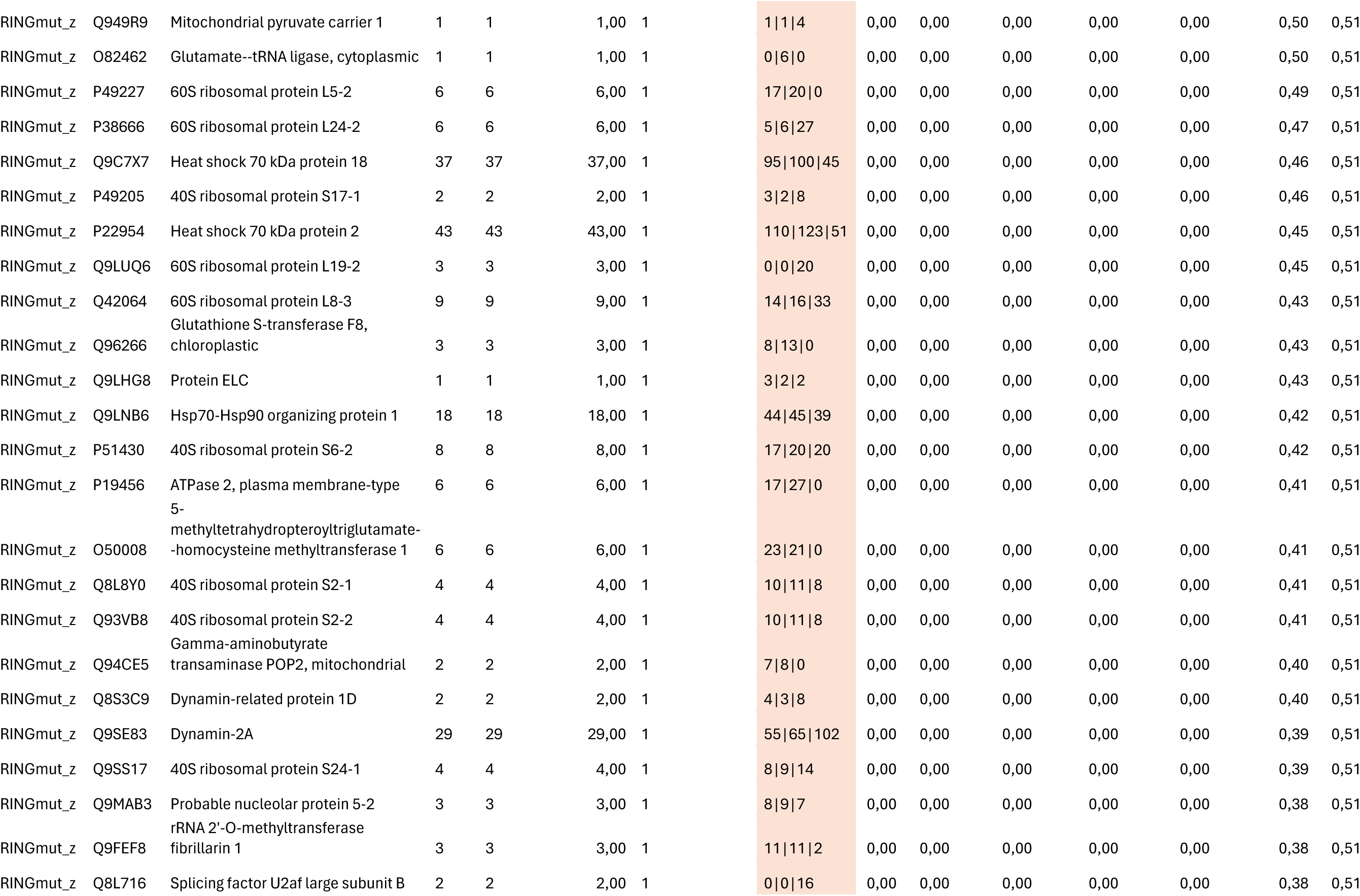
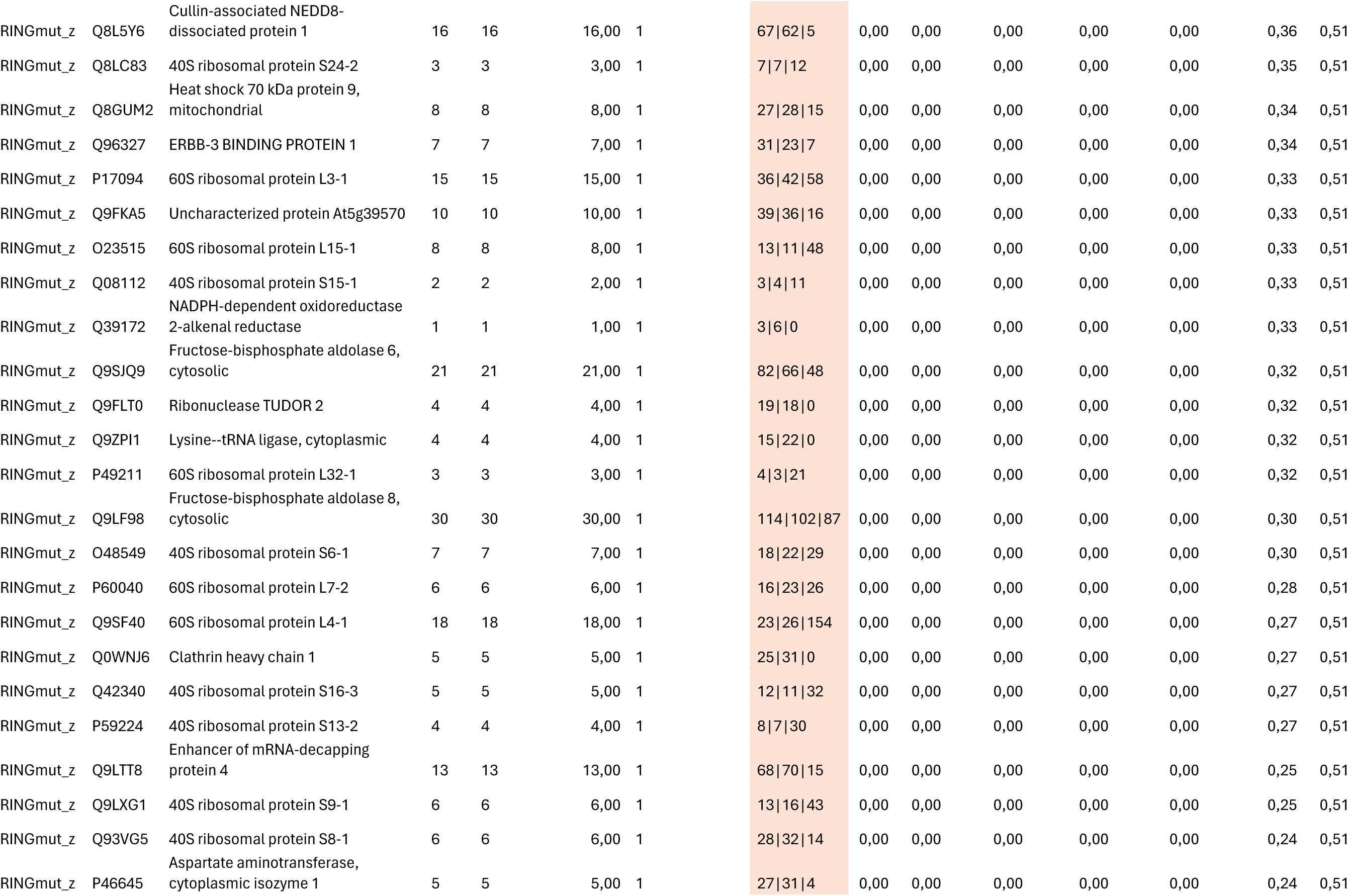
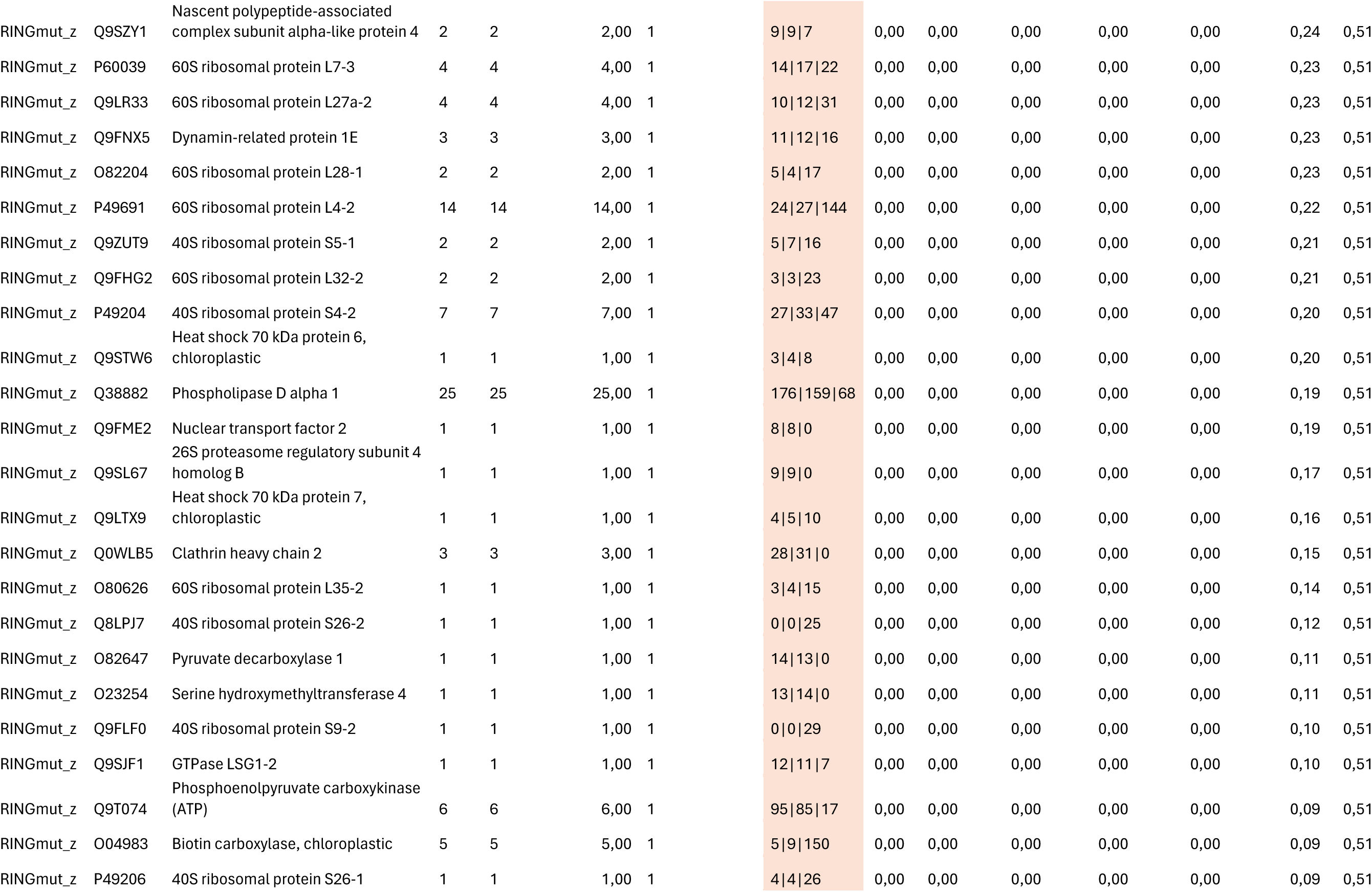
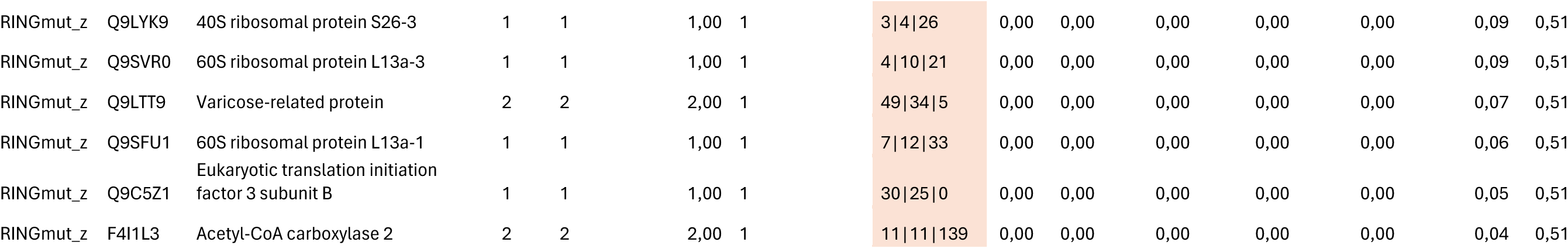

## REFERENCES

Adachi S, Minamisawa K, Okushima Y, Inagaki S, Yoshiyama K, Kondou Y, Kaminuma E, Kawashima M, Toyoda T, Matsui M, et al. 2011. Programmed induction of endoreduplication by DNA double-strand breaks in Arabidopsis. Proc Natl Acad Sci U S A 108(24): 10004–10009.

Bardoczy V, Geczi V, Sawasaki T, Endo Y, Meszaros T. 2008. A set of ligation-independent in vitro translation vectors for eukaryotic protein production. BMC Biotechnol 8: 32.

Beasley SA, Hristova VA, Shaw GS. 2007. Structure of the Parkin in-between-ring domain provides insights for E3-ligase dysfunction in autosomal recessive Parkinson’s disease. Proc Natl Acad Sci U S A 104(9): 3095–3100.

Berckmans B, Vassileva V, Schmid SP, Maes S, Parizot B, Naramoto S, Magyar Z, Alvim Kamei CL, Koncz C, Bogre L, et al. 2011. Auxin-dependent cell cycle reactivation through transcriptional regulation of Arabidopsis E2Fa by lateral organ boundary proteins. Plant Cell 23(10): 3671–3683.

Biedermann S, Harashima H, Chen P, Heese M, Bouyer D, Sofroni K, Schnittger A. 2017. The retinoblastoma homolog RBR1 mediates localization of the repair protein RAD51 to DNA lesions in Arabidopsis. EMBO J 36(9): 1279–1297.

Blilou I, Xu J, Wildwater M, Willemsen V, Paponov I, Friml J, Heidstra R, Aida M, Palme K, Scheres B. 2005. The PIN auxin efflux facilitator network controls growth and patterning in Arabidopsis roots. Nature 433(7021): 39–44.

Bourbousse C, Vegesna N, Law JA. 2018. SOG1 activator and MYB3R repressors regulate a complex DNA damage network in Arabidopsis. Proc Natl Acad Sci U S A 115(52): E12453–E12462.

Branon TC, Bosch JA, Sanchez AD, Udeshi ND, Svinkina T, Carr SA, Feldman JL, Perrimon N, Ting AY. 2018. Efficient proximity labeling in living cells and organisms with TurboID. Nat Biotechnol 36(9): 880–887.

Bueso E, Ibanez C, Sayas E, Munoz-Bertomeu J, Gonzalez-Guzman M, Rodriguez PL, Serrano R. 2014a. A forward genetic approach in Arabidopsis thaliana identifies a RING-type ubiquitin ligase as a novel determinant of seed longevity. Plant Sci 215-216: 110–116.

Bueso E, Rodriguez L, Lorenzo-Orts L, Gonzalez-Guzman M, Sayas E, Munoz-Bertomeu J, Ibanez C, Serrano R, Rodriguez PL. 2014b. The single-subunit RING-type E3 ubiquitin ligase RSL1 targets PYL4 and PYR1 ABA receptors in plasma membrane to modulate abscisic acid signaling. Plant J 80(6): 1057–1071.

Chen P, De Winne N, De Jaeger G, Ito M, Heese M, Schnittger A. 2023. KNO1-mediated autophagic degradation of the Bloom syndrome complex component RMI1 promotes homologous recombination. EMBO J 42(10): e111980.

Chen P, Sjogren CA, Larsen PB, Schnittger A. 2019. A multi-level response to DNA damage induced by aluminium. Plant J 98(3): 479–491.

Clay DE, Fox DT. 2021. DNA Damage Responses during the Cell Cycle: Insights from Model Organisms and Beyond. Genes (Basel) 12(12).

Cools T, Iantcheva A, Maes S, Van den Daele H, De Veylder L. 2010. A replication stress-induced synchronization method for Arabidopsis thaliana root meristems. Plant J 64(4): 705–714.

Cruz-Ramirez A, Diaz-Trivino S, Blilou I, Grieneisen VA, Sozzani R, Zamioudis C, Miskolczi P, Nieuwland J, Benjamins R, Dhonukshe P, et al. 2012. A bistable circuit involving SCARECROW-RETINOBLASTOMA integrates cues to inform asymmetric stem cell division. Cell 150(5): 1002–1015.

Culligan KM, Robertson CE, Foreman J, Doerner P, Britt AB. 2006. ATR and ATM play both distinct and additive roles in response to ionizing radiation. Plant J 48(6): 947–961.

Damodaran S, Strader LC. 2019. Indole 3-Butyric Acid Metabolism and Transport in Arabidopsis thaliana. Front Plant Sci 10: 851.

Damodaran S, Strader LC. 2024. Factors governing cellular reprogramming competence in Arabidopsis adventitious root formation. Dev Cell.

Davis OM, Ogita N, Inagaki S, Takahashi N, Umeda M. 2016. DNA damage inhibits lateral root formation by up-regulating cytokinin biosynthesis genes in Arabidopsis thaliana. Genes Cells 21(11): 1195–1208.

Dunn SD. 1986. Effects of the modification of transfer buffer composition and the renaturation of proteins in gels on the recognition of proteins on Western blots by monoclonal antibodies. Anal Biochem 157(1): 144–153.

Fernandez MA, Belda-Palazon B, Julian J, Coego A, Lozano-Juste J, Inigo S, Rodriguez L, Bueso E, Goossens A, Rodriguez PL. 2020. RBR-Type E3 Ligases and the Ubiquitin-Conjugating Enzyme UBC26 Regulate Abscisic Acid Receptor Levels and Signaling. Plant Physiol 182(4): 1723–1742.

Furukawa T, Curtis MJ, Tominey CM, Duong YH, Wilcox BW, Aggoune D, Hays JB, Britt AB. 2010. A shared DNA-damage-response pathway for induction of stem-cell death by UVB and by gamma irradiation. DNA Repair (Amst) 9(9): 940–948.

Gombos M, Raynaud C, Nomoto Y, Molnar E, Brik-Chaouche R, Takatsuka H, Zaki A, Bernula D, Latrasse D, Mineta K, et al. 2023. The canonical E2Fs together with RETINOBLASTOMA-RELATED are required to establish quiescence during plant development. Commun Biol 6(1): 903.

Harashima H, Sugimoto K. 2016. Integration of developmental and environmental signals into cell proliferation and differentiation through RETINOBLASTOMA-RELATED 1. Curr Opin Plant Biol 29: 95–103.

Hekkelman ML, de Vries I, Joosten RP, Perrakis A. 2023. AlphaFill: enriching AlphaFold models with ligands and cofactors. Nat Methods 20(2): 205–213.

Herbst J, Li QQ, De Veylder L. 2024. Mechanistic insights into DNA damage recognition and checkpoint control in plants. Nat Plants 10(4): 539–550.

Horvath BM, Kourova H, Nagy S, Nemeth E, Magyar Z, Papdi C, Ahmad Z, Sanchez-Perez GF, Perilli S, Blilou I, et al. 2017. Arabidopsis RETINOBLASTOMA RELATED directly regulates DNA damage responses through functions beyond cell cycle control. EMBO J 36(9): 1261–1278.

Jumper J, Evans R, Pritzel A, Green T, Figurnov M, Ronneberger O, Tunyasuvunakool K, Bates R, Zidek A, Potapenko A, et al. 2021. Highly accurate protein structure prediction with AlphaFold. Nature 596(7873): 583–589.

Kosarev P, Mayer KF, Hardtke CS. 2002. Evaluation and classification of RING-finger domains encoded by the Arabidopsis genome. Genome Biol 3(4): RESEARCH0016.

Laible M, Boonrod K. 2009. Homemade site directed mutagenesis of whole plasmids. J Vis Exp(27).

Lang L, Pettko-Szandtner A, Tuncay Elbasi H, Takatsuka H, Nomoto Y, Zaki A, Dorokhov S, De Jaeger G, Eeckhout D, Ito M, et al. 2021. The DREAM complex represses growth in response to DNA damage in Arabidopsis. Life Sci Alliance 4(12).

Lang L, Schnittger A. 2020. Endoreplication - a means to an end in cell growth and stress response. Curr Opin Plant Biol 54: 85–92.

LaRocque JR, McVey M. 2023. DNA Damage Response Mechanisms in Model Systems. Genes (Basel) 14(7).

Lee DJ, Park JY, Ku SJ, Ha YM, Kim S, Kim MD, Oh MH, Kim J. 2007. Genome-wide expression profiling of ARABIDOPSIS RESPONSE REGULATOR 7(ARR7) overexpression in cytokinin response. Mol Genet Genomics 277(2): 115–137.

Marin I. 2010. Diversification and Specialization of Plant RBR Ubiquitin Ligases. PLoS One 5(7): e11579.

Mathur J, Koncz C. 1998. Establishment and maintenance of cell suspension cultures. Methods Mol Biol 82: 27–30.

Mladek C, Guger K, Hauser MT. 2003. Identification and characterization of the ARIADNE gene family in Arabidopsis. A group of putative E3 ligases. Plant Physiol 131(1): 27–40.

Nagy SK, Kallai BM, Andras J, Meszaros T. 2020. A novel family of expression vectors with multiple affinity tags for wheat germ cell-free protein expression. BMC Biotechnol 20(1): 17.

Nemeth K, Salchert K, Putnoky P, Bhalerao R, Koncz-Kalman Z, Stankovic-Stangeland B, Bako L, Mathur J, Okresz L, Stabel S, et al. 1998. Pleiotropic control of glucose and hormone responses by PRL1, a nuclear WD protein, in Arabidopsis. Genes Dev 12(19): 3059–3073.

Nisa MU, Huang Y, Benhamed M, Raynaud C. 2019. The Plant DNA Damage Response: Signaling Pathways Leading to Growth Inhibition and Putative Role in Response to Stress Conditions. Front Plant Sci 10: 653.

Ogita N, Okushima Y, Tokizawa M, Yamamoto YY, Tanaka M, Seki M, Makita Y, Matsui M, Okamoto-Yoshiyama K, Sakamoto T, et al. 2018. Identifying the target genes of SUPPRESSOR OF GAMMA RESPONSE 1, a master transcription factor controlling DNA damage response in Arabidopsis. Plant J 94(3): 439–453.

Orosa-Puente B, Spoel SH. 2022. Harnessing the ubiquitin code to respond to environmental cues. Essays Biochem 66(2): 111–121.

Pedroza-Garcia JA, Xiang Y, De Veylder L. 2022. Cell cycle checkpoint control in response to DNA damage by environmental stresses. Plant J 109(3): 490–507.

Raina A, Sahu PK, Laskar RA, Rajora N, Sao R, Khan S, Ganai RA. 2021. Mechanisms of Genome Maintenance in Plants: Playing It Safe With Breaks and Bumps. Front Genet 12: 675686.

Reidt W, Wurz R, Wanieck K, Chu HH, Puchta H. 2006. A homologue of the breast cancer-associated gene BARD1 is involved in DNA repair in plants. EMBO J 25(18): 4326–4337.

Romero-Barrios N, Vert G. 2018. Proteasome-independent functions of lysine-63 polyubiquitination in plants. New Phytol 217(3): 995–1011.

Salvi E, Rutten JP, Di Mambro R, Polverari L, Licursi V, Negri R, Dello Ioio R, Sabatini S, Ten Tusscher K. 2020. A Self-Organized PLT/Auxin/ARR-B Network Controls the Dynamics of Root Zonation Development in Arabidopsis thaliana. Dev Cell 53(4): 431–443 e423.

Schonrock N, Exner V, Probst A, Gruissem W, Hennig L. 2006. Functional genomic analysis of CAF-1 mutants in Arabidopsis thaliana. J Biol Chem 281(14): 9560–9568.

Sjogren CA, Bolaris SC, Larsen PB. 2015. Aluminum-Dependent Terminal Differentiation of the Arabidopsis Root Tip Is Mediated through an ATR-, ALT2-, and SOG1-Regulated Transcriptional Response. Plant Cell 27(9): 2501–2515.

Spratt DE, Walden H, Shaw GS. 2014. RBR E3 ubiquitin ligases: new structures, new insights, new questions. Biochem J 458(3): 421–437.

Strader LC, Culler AH, Cohen JD, Bartel B. 2010. Conversion of endogenous indole-3-butyric acid to indole-3-acetic acid drives cell expansion in Arabidopsis seedlings. Plant Physiol 153(4): 1577–1586.

Strader LC, Wheeler DL, Christensen SE, Berens JC, Cohen JD, Rampey RA, Bartel B. 2011. Multiple facets of Arabidopsis seedling development require indole-3-butyric acid-derived auxin. Plant Cell 23(3): 984–999.

Su S, Zhang Y, Liu P. 2020. Roles of ubiquitination and SUMOylation in DNA damage response. Current issues in molecular biology 35(1): 59–84.

Szurman-Zubrzycka M, Jedrzejek P, Szarejko I. 2023. How Do Plants Cope with DNA Damage? A Concise Review on the DDR Pathway in Plants. Int J Mol Sci 24(3).

Takahashi H, Nozawa A, Seki M, Shinozaki K, Endo Y, Sawasaki T. 2009. A simple and high-sensitivity method for analysis of ubiquitination and polyubiquitination based on wheat cell-free protein synthesis. BMC Plant Biol 9: 39.

Takahashi N, Inagaki S, Nishimura K, Sakakibara H, Antoniadi I, Karady M, Ljung K, Umeda M. 2021. Alterations in hormonal signals spatially coordinate distinct responses to DNA double-strand breaks in Arabidopsis roots. Sci Adv 7(25).

Takahashi N, Ogita N, Koike T, Nishimura K, Inagaki S, Umeda M. 2022. Local induction of IAA5 and IAA29 promotes DNA damage-triggered stem cell death in Arabidopsis roots. bioRxiv: 2022.2009. 2002.506394.

Teo G, Liu G, Zhang J, Nesvizhskii AI, Gingras AC, Choi H. 2014. SAINTexpress: improvements and additional features in Significance Analysis of INTeractome software. J Proteomics 100: 37–43.

Varadi M, Anyango S, Deshpande M, Nair S, Natassia C, Yordanova G, Yuan D, Stroe O, Wood G, Laydon A, et al. 2022. AlphaFold Protein Structure Database: massively expanding the structural coverage of protein-sequence space with high-accuracy models. Nucleic Acids Res 50(D1): D439–D444.

M, Li X, Luo S, Fan B, Zhu C, Chen Z. 2020. Coordination and crosstalk between autophagosome and multivesicular body pathways in plant stress responses. Cells 9(1): 119.

Xie L, Lang-Mladek C, Richter J, Nigam N, Hauser MT. 2015. UV-B induction of the E3 ligase ARIADNE12 depends on CONSTITUTIVELY PHOTOMORPHOGENIC 1. Plant Physiol Biochem 93: 18–28.

Xuan W, Audenaert D, Parizot B, Moller BK, Njo MF, De Rybel B, De Rop G, Van Isterdael G, Mahonen AP, Vanneste S, Beeckman T. 2015. Root Cap-Derived Auxin Pre-patterns the Longitudinal Axis of the Arabidopsis Root. Curr Biol 25(10): 1381–1388.

Yi D, Alvim Kamei CL, Cools T, Vanderauwera S, Takahashi N, Okushima Y, Eekhout T, Yoshiyama KO, Larkin J, Van den Daele H, et al. 2014. The Arabidopsis SIAMESE-RELATED cyclin-dependent kinase inhibitors SMR5 and SMR7 regulate the DNA damage checkpoint in response to reactive oxygen species. Plant Cell 26(1): 296–309.

Yoo SD, Cho YH, Sheen J. 2007. Arabidopsis mesophyll protoplasts: a versatile cell system for transient gene expression analysis. Nat Protoc 2(7): 1565–1572.

Yoshiyama K, Conklin PA, Huefner ND, Britt AB. 2009. Suppressor of gamma response 1 (SOG1) encodes a putative transcription factor governing multiple responses to DNA damage. Proc Natl Acad Sci U S A 106(31): 12843–12848.

Yoshiyama KO, Kobayashi J, Ogita N, Ueda M, Kimura S, Maki H, Umeda M. 2013a. ATM-mediated phosphorylation of SOG1 is essential for the DNA damage response in Arabidopsis. EMBO Rep 14(9): 817–822.

Yoshiyama KO, Sakaguchi K, Kimura S. 2013b. DNA damage response in plants: conserved and variable response compared to animals. Biology (Basel) 2(4): 1338–1356.

Yu C, Hou L, Huang Y, Cui X, Xu S, Wang L, Yan S. 2023. The multi-BRCT domain protein DDRM2 promotes the recruitment of RAD51 to DNA damage sites to facilitate homologous recombination. New Phytol 238(3): 1073–1084.

## Supporting Information References

Bardóczy, V., Géczi, V., Sawasaki, T., Endo, Y., & Mészáros, T. (2008). A set of ligation-independent in vitro translation vectors for eukaryotic protein production. BMC biotechnology, 8(1), 1–7.

Branon, T. C., Bosch, J. A., Sanchez, A. D., Udeshi, N. D., Svinkina, T., Carr, S. A., … & Ting, A. Y. (2018). Efficient proximity labeling in living cells and organisms with TurboID. Nature biotechnology, 36(9), 880–887.

Fischer, R., & Kessler, B. M. (2015). Gel-aided sample preparation (GASP)—A simplified method for gel-assisted proteomic sample generation from protein extracts and intact cells. Proteomics, 15(7), 1224–1229.

Haider, S. R., Reid, H. J., & Sharp, B. L. (2012). Tricine-sds-page. Protein electrophoresis: methods and protocols, 81–91.

Hubner, N. C., Bird, A. W., Cox, J., Splettstoesser, B., Bandilla, P., Poser, I., … & Mann, M. (2010). Quantitative proteomics combined with BAC TransgeneOmics reveals in vivo protein interactions. Journal of Cell Biology, 189(4), 739–754.

Jankovics, F., Bence, M., Sinka, R., Faragó, A., Bodai, L., Pettkó-Szandtner, A., … & Erdélyi, M. (2018). Drosophila small ovary gene is required for transposon silencing and heterochromatin organization, and ensures germline stem cell maintenance and differentiation. Development, 145(23), dev170639.

Kobayashi, K., Suzuki, T., Iwata, E., Nakamichi, N., Suzuki, T., Chen, P., … & Ito, M. (2015). Transcriptional repression by MYB 3R proteins regulates plant organ growth. The EMBO journal, 34(15), 1992–2007.

Laible, M., & Boonrod, K. (2009). Homemade site directed mutagenesis of whole plasmids. JoVE (Journal of Visualized Experiments), (27), e1135.

Mathur, J., & Koncz, C. (1998). Establishment and maintenance of cell suspension cultures. Arabidopsis Protocols, 27–30.

Nagy, S. K., Darula, Z., Kállai, B. M., Bogre, L., Bánhegyi, G., Medzihradszky, F., … & Mészáros, T. (2015). Activation of AtMPK9 through autophosphorylation that makes it independent of the canonical MAPK cascades.

Nagy, S. K., Kállai, B. M., András, J., & Mészáros, T. (2020). A novel family of expression vectors with multiple affinity tags for wheat germ cell-free protein expression. BMC biotechnology, 20, 1–9.

Németh, K., Salchert, K., Putnoky, P., Bhalerao, R., Koncz-Kálmán, Z., Stankovic-Stangeland, B., … & Koncz, C. (1998). Pleiotropic control of glucose and hormone responses by PRL1, a nuclear WD protein, in Arabidopsis. Genes & development, 12(19), 3059–3073.

Studier, F. W. (2005). Protein production by auto-induction in high-density shaking cultures. Protein expression and purification, 41(1), 207–234.

Yoo, Sang-Dong, Young-Hee Cho, and Jen Sheen. Arabidopsis mesophyll protoplasts: a versatile cell system for transient gene expression analysis. Nature protocols 2.7 (2007): 1565–1572.

